# V- and V_L_-Scores Uncover Viral Signatures and Origins of Protein Families

**DOI:** 10.1101/2024.10.24.619987

**Authors:** Kun Zhou, James C. Kosmopoulos, Etan Dieppa Colón, Peter John Badciong, Karthik Anantharaman

## Abstract

Viruses are key drivers of microbial diversity, nutrient cycling, and co-evolution in ecosystems, yet their study is hindered due to challenges in culturing. Traditional gene-centric methods, which focus on a few hallmark genes like for capsids, miss much of the viral genome, leaving key viral proteins and functions undiscovered. Here, we introduce two powerful annotation-free metrics, V-score and V_L_-score, designed to quantify the “virus-likeness” of protein families and genomes and create an open-access searchable database, ‘V-Score-Search’. By applying V- and V_L_-scores to public databases (KEGG, Pfam, and eggNOG), we link 38−77% of protein families with viruses, a 9−16x increase over current estimates. These metrics outperform existing approaches, enabling precise detection of viral genomes, prophages, and host-derived auxiliary viral genes (AVGs) from fragmented sequences, and significantly improving genome binning. Remarkably, we identify up to 17x more AVGs, dominated by non-metabolic proteins of unknown function. This innovation unlocks new insights into virus signatures and host interactions, with wide-ranging implications from genomics to biotechnology.

## MAIN

Viruses are indispensable components of the biosphere. By their sheer abundance in microbiomes and ecosystems and their high genetic diversity^1^, viruses have the ability to regulate populations^2^, facilitate nutrient cycling^3^, promote genetic diversity^4^, and drive co-evolutionary dynamics^5^. In spite of their importance, viruses are difficult to culture in the laboratory necessitating advances in computational approaches to study uncultured viruses. Understanding viral genomes and proteins is crucial for grasping their diversity and understanding their roles in ecosystems. This knowledge helps unravel the complexity of life and advances biotechnological applications like vaccines and phage therapy.

Traditionally, virus-specific genes, including hallmark genes such as for capsid proteins, have been considered the definitive signatures of viral genomes and used for identifying and characterizing viral genomes^6-8^. However, hallmark genes account for a small portion of viral genomes^9^. Genome or metagenome fragments often do not contain hallmark genes, making it difficult to identify and classify viruses using traditional gene-centric approaches. As a result, many viral genomes remain unidentified, leading to a significant loss of information and a growing recognition of the need to overcome these limitations in viral discovery and protein annotation.

Annotating viral genes and predicting their functions provide clues about the nature of viral sequences and protein families. We reasoned that analyzing entire viral genomes, even when fragmented, with functional annotations could break convention and yield innovative viral signatures. Here we introduce the concepts of V-scores and V_L_-scores that are quantitative metrics to serve as a virus-like signature for differentiating between viral and non-viral protein families and genomes. We demonstrate specific use cases of V-scores and V_L_-scores in virus identification, prophage discovery, annotation of host-derived and metabolic proteins on viral genomes, and virus genome binning. Finally, to facilitate adoption of our approach, we created a publicly available database of V-scores and V_L_-scores associated with every protein cluster or family in five widely used public databases (https://anantharamanlab.github.io/V-Score-Search/) including Prokaryotic Virus Remote Homologous Groups (PHROG), Virus Orthologous Groups (VOG), Kyoto Encyclopedia of Genes and Genomes (KEGG), Protein Families Database (Pfam), and evolutionary genealogy of genes: Non-supervised Orthologous Groups (eggNOG). We propose that V-scores and V_L_-scores will serve as a metric to define the likelihood of protein families being detected in viruses and enable diverse applications associated with viral genomics, ecology, and evolution.

## RESULTS

### Assessment of protein families for virus-like proteins

We used 18,435,589 viral proteins sourced from diverse viruses to construct associations between viruses and protein families (**Fig. 1a**). Each protein family (i.e., clusters of similar proteins represented under a single annotation in databases, which includes proteins of unknown function) was assigned a V-score and a V_L_-score, representing metrics of virus association when the protein family had significant hits to viral proteins (see details in Methods and in Supplementary Tables S1-5). We identified cutoffs of V-score = 0.01 and V_L_-score = 0 to define viral proteins with high certainty. High V-scores and V_L_-scores indicated a strong association with viral proteins, whereas low scores suggested a weaker association. Protein families associated with viral proteins constituted approximately 76.9%, 52.1%, and 38.7% of the total protein families in KEGG (20,005), Pfam (10,835), and eggNOG (135,509), respectively (**Fig. 1b**). In contrast, current estimates of viral protein entries in KEGG, Pfam, and eggNOG are limited, representing a very small fraction (<10%) (**Supplementary Fig. S1**). Our analysis substantially increases the number of protein families in public databases associated with viruses and significantly improves the overall representation of viral proteins in these databases. This increase in viral representation will facilitate better understanding of viral roles in ecosystems, their interactions with hosts, and their evolutionary dynamics.

**Fig. 1.**
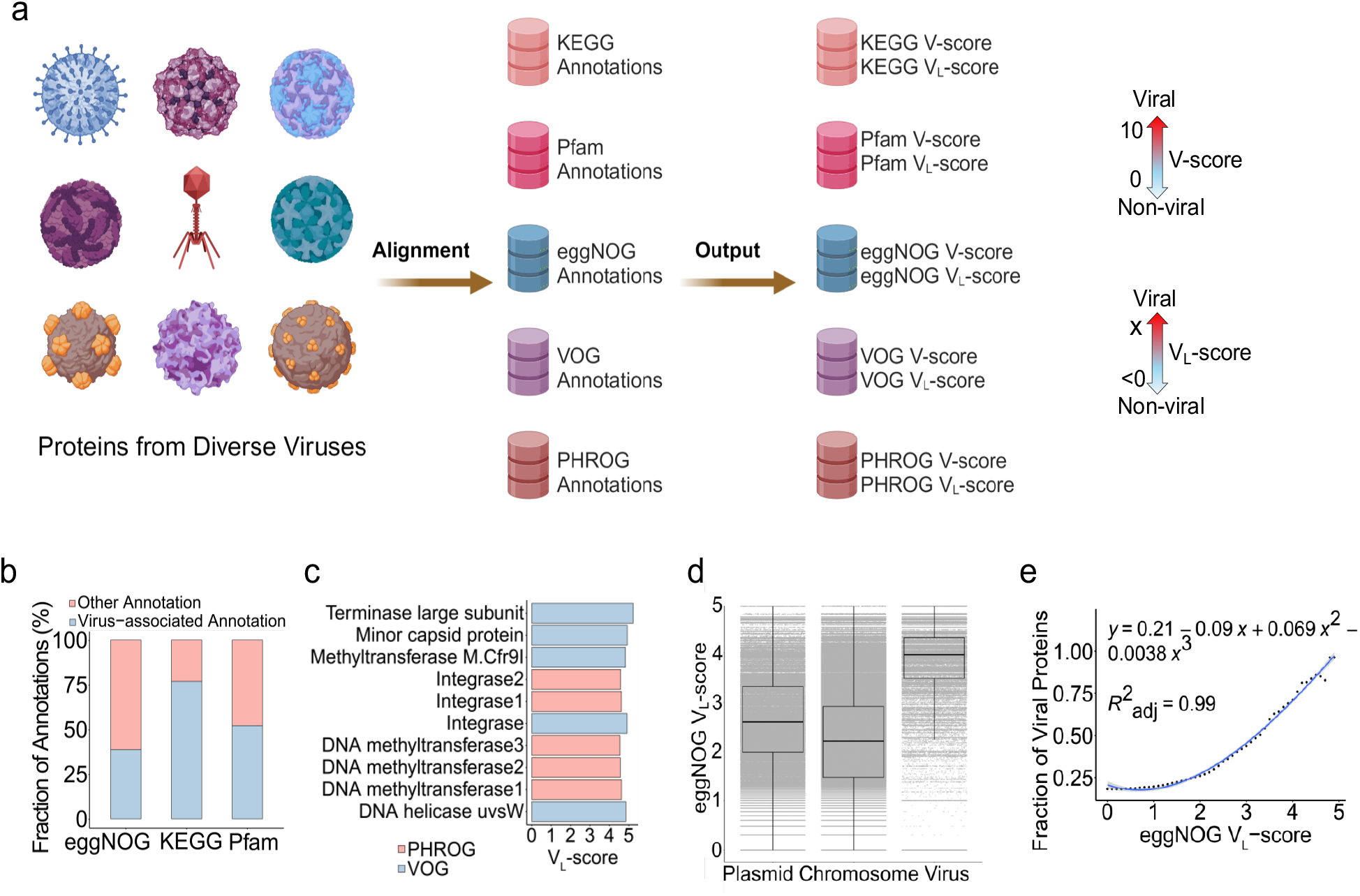
Concepts of V-score and V_L_-score. **a**, Workflow of V-score and V_L_-score generation. Nine representatives of viral taxa are shown here for the diverse viruses used in the study. A scale for V_L_-scores and V_L_-scores is displayed by two-sided arrows going from 0 to 10 and <0 to X, respectively, suggesting low scores indicate non-viral and high-scores indicate viral. **b**, Frequency of virus-associated annotations with V-score ≥ 0.01 and/or V_L_-score ≥ 0. **c**, Top five annotations associated with viruses based on V_L_-scores. **d**, Distribution of eggNOG V_L_-score across proteins from prokaryotic chromosomes (*n* = 7,561,596), plasmids (*n* = 437,241) and prokaryotic viruses (*n* = 83,664). The horizontal line that splits the box represents the median, upper and lower sides of the box represent upper and lower quartiles, whiskers are 1.5 times the interquartile ranges and data points beyond whiskers are considered potential outliers. **e**, Relationship between the fraction of viral proteins used in (d) and eggNOG V_L_-score. The generation of the fraction of viral proteins from the comparison between plasmids, chromosomes, and viruses is illustrated in Supplementary **Fig. S10**.

Next, we hypothesized that the associative nature of V-scores and V_L_-scores could also reflect gene frequencies in viral communities. Towards this, we used the PHROG and VOG protein families that provide valuable resources for characterizing viral proteins. We determined that the range of V-scores and V_L_-scores were associated with patterns of gene frequencies with high scores indicating frequent distributions and low scores indicating infrequent distributions. For example, according to PHROG and VOG V_L_-scores, methyltransferase-coding genes were frequently distributed in viral communities (Fig. 1c), which was also evidenced by the high V_L_-scores for these protein families in KEGG, Pfam, and eggNOG (e.g., KEGG V_L_-score = 4.8 and Pfam V_L_-score = 4.7). This approach will allow for the identification of new viral hallmark proteins and other proteins commonly encountered on viruses but whose function is currently not known. In contrast, protein families with very low V-scores and V_L_-scores, e.g., host-derived proteins, metabolic proteins, and hypothetical proteins with V-scores of 0.01, indicated the presence of viral proteins that are rare in communities and may confer specialized functions more likely to be involved in niche-specific interactions^10^.

Interestingly, V_L_-scores of eggNOG protein families revealed the likelihood of viral origin of different protein families. V_L_-scores revealed a significant difference between viral and non-viral proteins when comparing viral proteins to those found in plasmids and prokaryotic chromosomes (**Fig. 1d**). The proportion of viral proteins in a protein family increased with higher eggNOG V_L_-scores, demonstrating a clear relationship between scores and the probability of viral origin (**Fig. 1e)**. High V_L_-scores (>4) indicated that the protein families are likely virus-specific, while low V_L_-scores (<2.2) suggest non-viral origin (**Fig. 1e**). This finding offers a promising approach for the differentiation between viral and non-viral proteins, extending beyond simple gene presence or absence and incorporating quantitative assessment. Such metrics could be particularly useful in cases where traditional methods struggle, such as in distinguishing viral genes embedded within plasmids^11^ or identifying viral elements within bacterial genomes^12, 13^. Additionally, these quantitative metrics for protein families can also be applied for the differentiation of viral and non-viral genome sequences using combined V_L_-scores or V-scores across different proteins.

### Generation of AV-scores and AV_L_-scores for viral differentiation and prediction

To build upon our understanding of V-scores and V_L_-scores from the protein to genome-scales, we posited that the association and frequency of V-scores and V_L_-scores may confer features on viral genomes that distinguish them from other organisms. To test on this, we investigated a whole-genome catalog of 5,800 viral, 50,523 plasmid, and 4,813 prokaryotic genomes and developed the concepts of average V-score (AV-score) and average V_L_-score (AV_L_-score) (See methods for details) (**Fig. 2a**). We proposed that AV- and AV_L_-scores represented the average scores of protein families across an entire genome and would thus be representative of the overall virus-like character of a given genome. We determined that prokaryotic viruses had significantly higher medians of AV-scores (3.602−9.515) and AV_L_-scores (1.802−3.830) compared to plasmids and prokaryotic chromosomes regardless of annotation databases (*p-value* < 10^−5^). Interestingly, viral genome fragments (1−15kb) extracted from whole genomes also displayed significantly higher medians (see examples of KEGG and Pfam AV-scores and AV_L_-scores in **Supplementary Fig. S2** and **S3**, respectively). The higher median scores for viral genomes suggest that this metric could capture features unique to viruses, making it highly effective for identifying viral genomes in mixed communities such as metagenomes of viruses, plasmids, and chromosomes. To validate this, we conducted polynomial regression analyses on the fraction of viral genomes within mixed metagenomes containing viruses, plasmids, and chromosomes at various cutoffs of AV-scores and AV_L_ -scores for both whole genomes and genome fragments (**Supplementary Tables S6-9**). At the whole-genome level, the fraction of viral genomes increased with higher AV-score and AV_L_ - scores (for VOG) (**Fig. 2b**). Similarly, at the fragment level, the fraction of viral genomes increased with higher AV-score cutoffs for KEGG and Pfam (**Supplementary Fig. S4** and **S5**). From regression analyses (**Fig. 2b**), whole genomes with AV-scores/ AV_L_ -scores exceeding the corresponding cutoffs (e.g., a VOG AV-score of 2, which surpasses the VOG AV-score cutoff of 1.93) were predicted to be viral with a 70% probability (likely viral) or a 90% probability (most likely viral) (see detailed cutoffs in **Supplementary Table S10**). For genome fragments, only the AV-scores of VOG, PHROG, KEGG, and Pfam were able to generate cutoffs predictive of viral genomes with a 70% or 90% probability (**Supplementary Fig. S4−7**). Given that cutoffs may vary with fragment size, different cutoffs were established for corresponding sizes (**Supplementary Table S10**). Overall, the concepts of AV-scores and AV_L_-scores offer novel insights into genome signatures, traditionally defined by k-mer frequency^14^ or single-copy signature genes^15^. The cutoffs for AV-scores and AV_L_-scores, used to differentiate between viral and non-viral genomes, may prove valuable for viral identification in metagenomic studies. Overall, these metrics address limitations of conventional gene-centric and alignment-dependent methods^8, 16-18^.

**Fig. 2.**
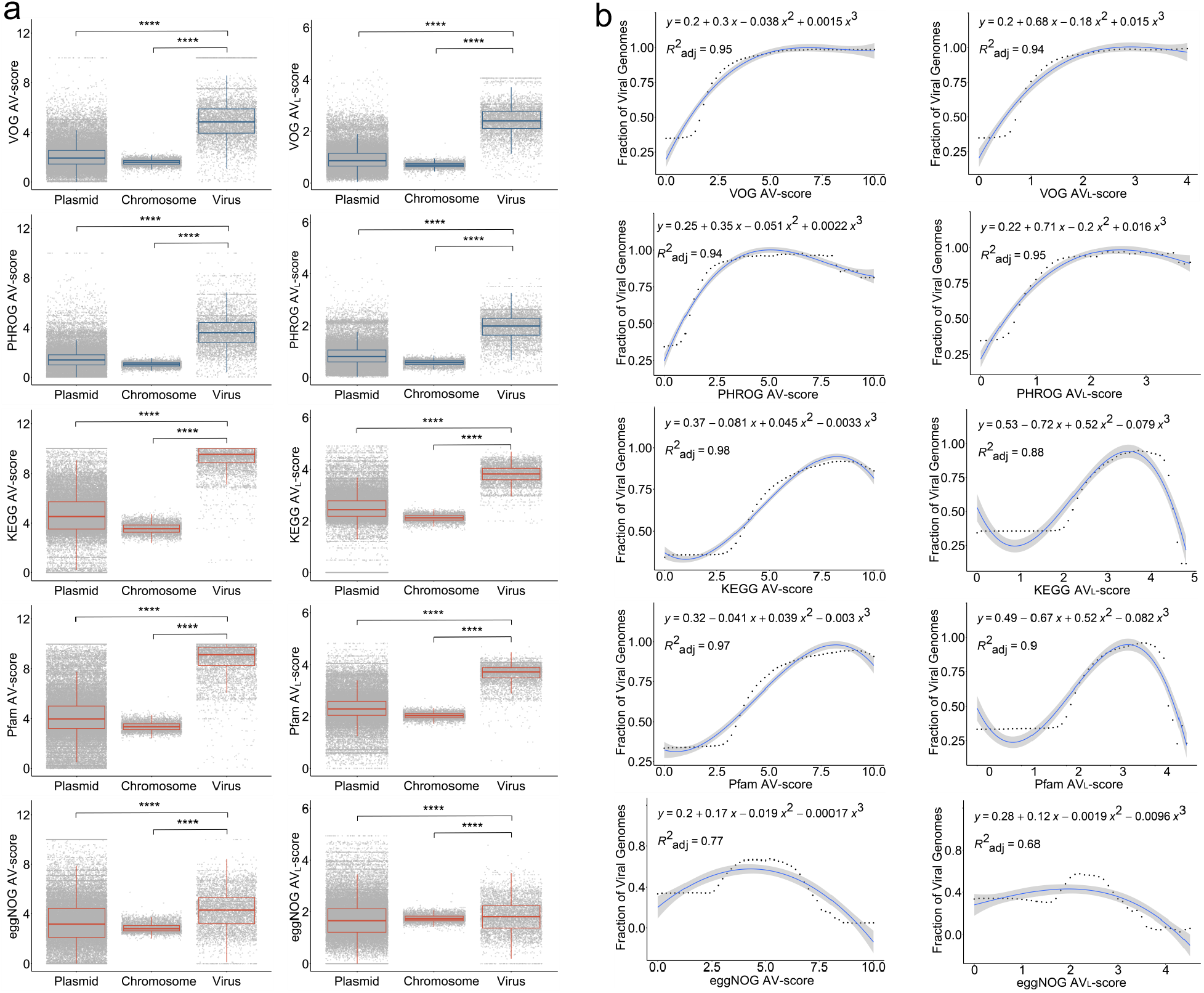
Concepts of Average V-score (AV-score), Average V_L_-score (AV_L_-score), and cutoffs of AV-score and AV_L_-score. **a**, Distribution of AV-score and AV_L_-score of prokaryotic chromosomes (*n* = 4,813) and the genomes of plasmids (*n* = 50,523) and prokaryotic viruses (*n* = 5,800). The blue boxes denote the AV-scores and AV_L_-scores of VOG and PHROG. The red boxes denote the AV-scores and AV_L_-scores of KEGG, Pfam, and eggNOG. The horizontal line that splits the box is the median, the upper and lower sides of the box are upper and lower quartiles, whiskers are 1.5 times the interquartile ranges and data points beyond whiskers are considered potential outliers. An ANOVA test was used to show differences between three means are significant (*p* < 2.2 × 10^−16^). **** denotes *p* < 10^−4^. **b**, Relationship between the fraction of viral genomes used in (a) and the AV-scores and AV_L_-scores. In this study, we define the fraction of viral genomes as the probability that a given genome sequence is viral. The dots on the dotted line represent the actual values of the fraction of viral genome sequences, while the blue lines indicate the predicted values. The process for generating the fraction of viral genome sequences is identical to the method used for generating the fraction of viral proteins, as illustrated in Supplementary **Fig. S10**.

### Maximizing identification of viral genomes

To evaluate the potential of AV-scores and AV_L_-scores for applications in metagenomics, we analyzed a dataset of 39 host-associated metagenomes. By applying AV-score cutoffs (with a 70% probability of being viral) for genome fragments of varying sizes, derived from KEGG, Pfam, VOG, or PHROG, we identified 13,167 viral sequences of low, medium, and high quality (**Fig. 3a**). Of these, 2,064 sequences overlapped with those identified using geNomad which is a virus identifier dependent on virus-specific markers^8^ (**Supplementary Fig. S8a**). Notably, for medium- and high-quality sequences, the AV-score-based approach outperformed geNomad, identifying more than 1,000 high-quality viral sequences—approximately seven times more than geNomad identified (**Supplementary Fig. S8b**).

**Fig. 3.**
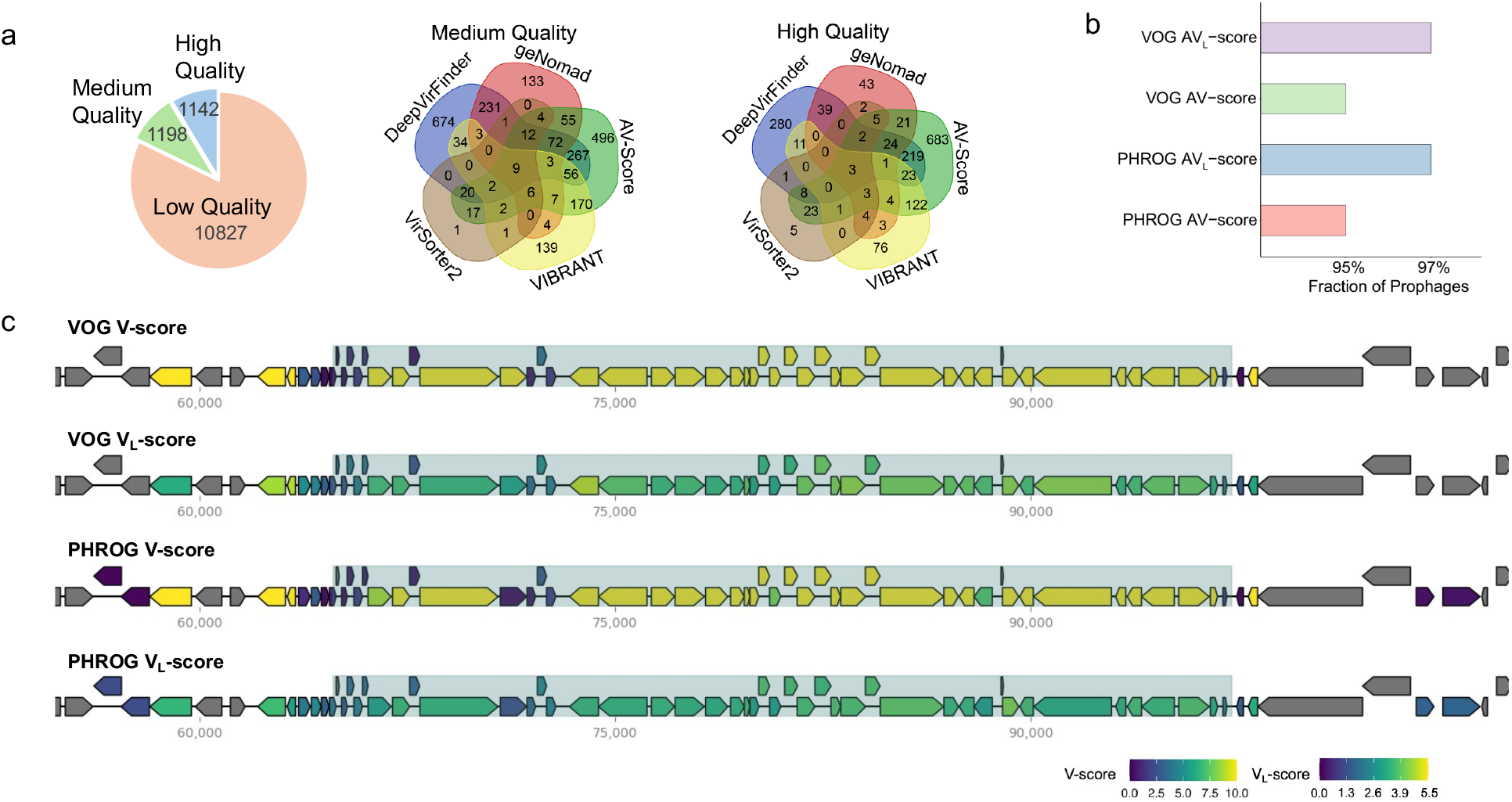
Application of V-scores, V_L_-scores, AV-scores, and AV_L_-scores for viral identification in genomes and metagenomes. **a**, Number of sequences identified with AV-scores and AV_L_-scores and four commonly used software. For medium- and high-quality sequences, as assessed by CheckV, the overlap between the five approaches was illustrated using Venn diagrams, showing the number of shared and specific sequences identified by different methods. **b**, Fraction of prophages in a database that have AV-scores and AV_L_-scores above corresponding cutoffs for viral-like determination. **c**, Distribution of V-scores and V_L_-scores for genes within a verified *Escherichia coli* prophage and its adjacent host sequences. Prophage regions are shaded for emphasis.

Additionally, the AV-score-based method surpassed other conventional tools, including machine learning-dependent DeepVirFinder^19^, VIBRANT^17^, a hybrid approach incorporating machine learning and protein similarity, and VirSorter2^18^, in identifying high-quality sequences (Fig. 3a). Moreover, compared to previous studies on sponge-associated microbiomes^20, 21^, we identified 129 viral sequences of medium or higher quality—more than 15 times the number of viral genomes (7 sequences) previously predicted using VirSorter2^7^. Most of the high-quality viral genomes identified by the AV-score approach are specific to AV-score, indicating that this method can uncover viral genomes that other tools may not recognize (**Fig. 3a**). These findings suggest that the usage of AV-scores and AV_L_-scores can detect many viral sequences that traditional, viral-specific gene-dependent methods may overlook. Overall, the application of AV-scores and AV_L_-scores as metrics for genome differentiation offers a novel and powerful tool for identifying viral genomes in metagenomic studies.

We further tested the potential of this approach for prophage identification and assessment. The results showed that over 95% of sequences in a prophage database used by a popular prophage identification tool, PHASTER^22^ (65,668 prophages), had AV-scores and AV_L_-scores above our suggested cutoffs for whole genomes (70% probability, based on VOG and PHROG scores) (**Fig. 3b**). Additionally, clear boundaries between a verified Escherichia coli prophage and its adjacent host sequences were delineated by relatively low V-scores and V_L_-scores using VOG and PHROG (**Fig. 3c**). Furthermore, the higher AV-scores observed for VOG, PHROG, Pfam, KEGG, and eggNOG families in prophages (see **Supplementary Fig. S9**) strongly support the idea that AV-scores and/or AV_L_-scores are useful in identifying prophage boundaries when combined with sliding window approaches (e.g., a 10 kb sliding window^23^). In addition to AV-scores and AV_L_-scores, V_L_-scores may also be valuable for determining boundaries, as a gene with an eggNOG V_L_-score greater than 4 has over a 70% probability of being viral (Fig. 1e). Accurately predicting prophage boundaries has long been a challenge^24, 25^, possibly due to the presence of auxiliary metabolic genes (AMGs) in phages^26, 27^ or the ability of phages to be transposable and encode serine-integrases rather than tyrosine integrases^24^. Given their ability to distinguish viral from non-viral genes and sequences, AV-scores, AV_L_-scores, and V_L_-scores may offer highly precise methods for boundary recognition.

### Advancing the identification of auxiliary genes in viral genomes

Despite recent efforts, the vast majority of viral proteins (>80%) have no known function which has hindered our understanding of the roles of viruses in ecosystems and microbiomes. V-scores and V_L_-scores as quantitative metrics display a property of measuring the frequency of individual protein families among viral genomes in public databases. Leveraging this property through the development of hidden Markov models for protein families, we assessed their effectiveness in identifying AVGs, including AMGs on viral genomes. AVGs are virus-encoded genes of prokaryotic origin that are not essential for viral propagation processes such as genome replication, lysis, or capsid assembly, while AMGs are auxiliary genes that are associated with metabolic roles^28^. Such genes likely provide a fitness benefit to the virus encoding them^28-30^. Identifying AVGs is a particularly difficult problem compounded by host-associated contamination and the host-derived nature of these genes. Given their importance due to the increasing recognition of auxiliary genes involved in human and environmental microbiomes^30-34^, we investigated whether V-scores and V_L_-scores could effectively identify auxiliary genes.

To test this hypothesis, we evaluated the ability of V-scores, V_L_-scores, AV-scores, and AV_L_-scores to identify 17 experimentally verified AMGs. We first distinguished AMGs from host-encoded metabolic genes and non-auxiliary genes by using V-scores and VL-scores (**Fig. 4a and 4b**). We then averaged the V_L_-scores of all KEGG or Pfam protein families across entire scaffolds, establishing a scaffold Pfam/KEGG AV_L_-score of 3 as optimal for differentiating viral from host scaffolds (**Fig. 4c**). Our workflow effectively detected AMGs (**Fig. 4d**). We achieved a sensitivity of 97.71% and a false positivity rate of 2.29% using a database of biochemically characterized AMGs (experimentally verified) for benchmarking (see details in **Supplementary Table S11**). Community standards for analyzing AMGs recommend verifying that a virally encoded AMG is flanked both upstream and downstream by hallmark genes^35, 36^. This check ensures that metabolic genes identified from proviral sequences are not in regions of host contamination, however, this standard hinders AMG recall for non-proviruses. The requirement for verification significantly reduced sensitivity to 66% (when verified with genes having V-scores of 10) and to 2.67% (when verified with hallmark genes), while also increasing the false discovery rate to 30% when using hallmark gene verification (**Fig. 4d, Supplementary Tables S11, S12**). The ability of V-scores and V_L_-scores to confidently identify viral proteins circumvents the need to identify hallmark proteins. Therefore V-scores offer a novel methodology for verifying that AMGs encoded by proviruses are not the result of host contamination.

**Fig. 4.**
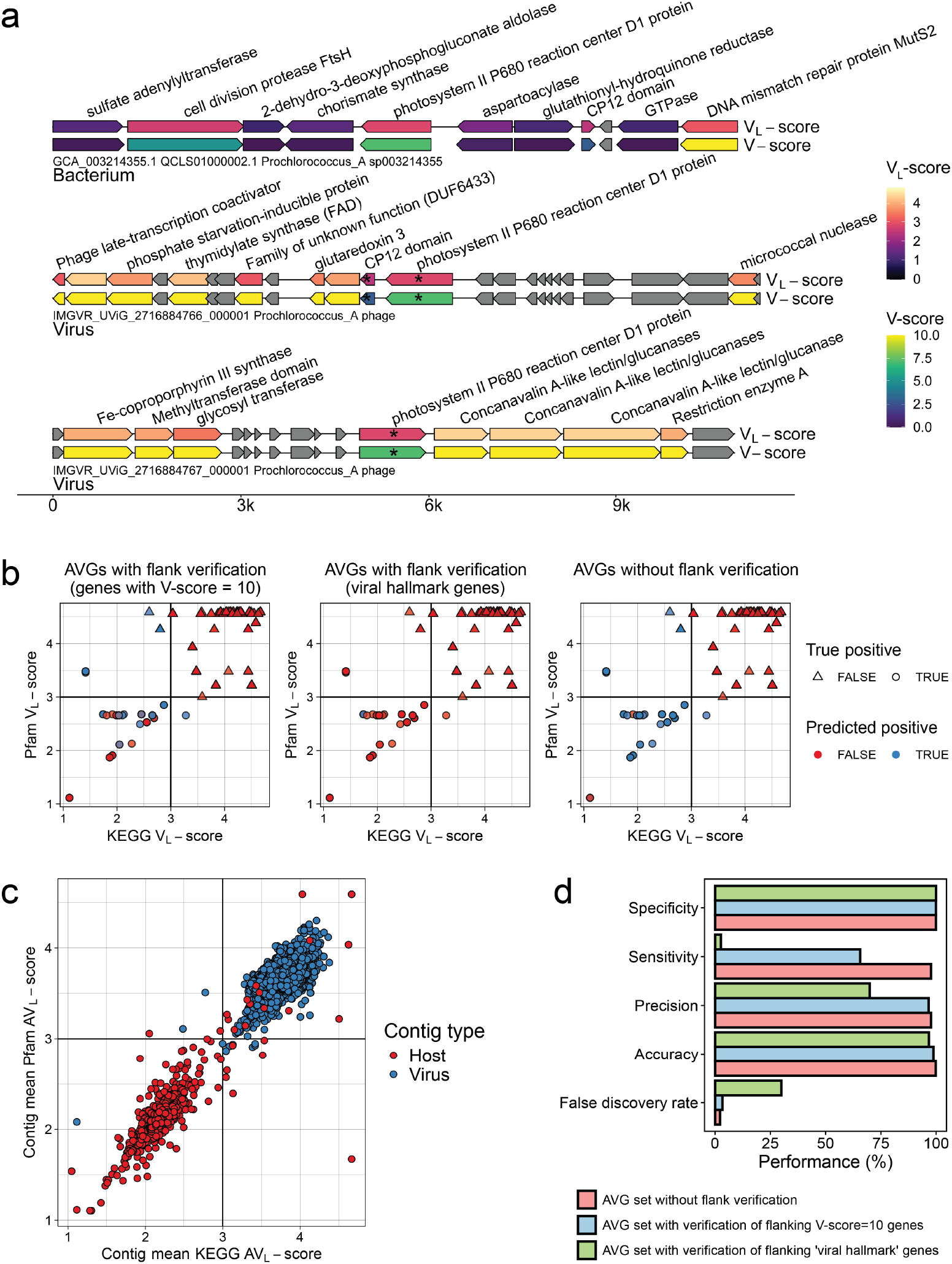
Application of V-scores, V_L_-scores, and AV_L_-scores to auxiliary gene (AVG) detection. **a**, V-scores and V_L_-scores reveal AMGs in viral genomes and distinguish AMGs from host-encoded metabolic genes. Genes with an asterisk (*) were predicted as AMGs using the described workflow (see Methods). **b**, Establishing optimal Pfam V_L_-score / KEGG V_L_-score combinations to distinguish viral auxiliary vs. non-auxiliary genes. Points represent individual genes in our database of viral and host genomes that had both Pfam^5^ and KEGG^6^ annotations matching to either the database of the 17 AMGs or 10 non-AMG protein families. Genes marked as potentially auxiliary have a maximum KEGG and Pfam V_L_-scores of 3, as indicated by the vertical and horizontal lines. **c**, Establishing the optimal Pfam/KEGG AV_L_-score of query scaffolds to distinguish viral vs. host genomes. Points represent individual genes, plotted by the AV_L_-score of all Pfam or KEGG annotations encoded by the gene’s origin scaffold. Vertical and horizontal lines represent the chosen scaffold AV_L_-score used to distinguish viral from host scaffolds (> 3: virus, < 3: host). Points are colored by the actual scaffold type of the gene’s origin (host or virus). **d**, Performance of the proposed AMG identification workflow. Performance was evaluated based on the confusion matrices in Supplementary Table S12.

Leveraging this advantage, we were able to predict a significantly larger number of auxiliary genes from 5,116 high-quality viral genomes, providing deeper insights into viral functions. Our workflow (with verified flanking genes with V-score=10) identified a total of 27,442 viral genes likely to be auxiliary and the workflow without verification predicted 34,015 auxiliary genes (4.85% of all viral genes in our test dataset and 16.50% of all annotated viral genes) (**Supplementary Table S13**). Notably, non-metabolic AVGs comprise a substantial majority, accounting for 89%, while auxiliary metabolic genes represent a small subset, making up only 11% (**Fig. 5a**). The identified AVGs included genes encoding various metabolic enzymes, antibiotic resistance proteins, transporters, DNA/RNA replication proteins, transposases/recombinases, nucleases/endonucleases, and uncharacterized/hypothetical proteins. These AVGs serve diverse functions including metabolism, genetic information processing, environmental information processing, and cellular processes (**Fig. 5b**; **Supplementary Table S13**). Some of the genes have been considered auxiliary, for example, the genes encoding D-3-phosphoglycerate dehydrogenase for carbon metabolism^26^, S-adenosylmethionine decarboxylase for amino acid metabolism^37^, and alpha-L-fucosidase for glycan degradation^38^. Notably, our study predicted numerous auxiliary genes that were typically overlooked in previous studies of auxiliary genes. For instance, over 700 viral auxiliary genes related to toxin-antitoxin systems were identified. These systems, which are typically used by hosts as a defense mechanism against viral infections^39, 40^, may be employed by viruses to enhance their ability to infect host organisms^39, 41, 42^, contributing to viral evolution in the ongoing virus-host arms race. Additionally, the presence of many genes with unknown functions suggests that there are still numerous unexplored roles for viruses, likely with important ecosystem or microbiome contexts.

**Fig. 5.**
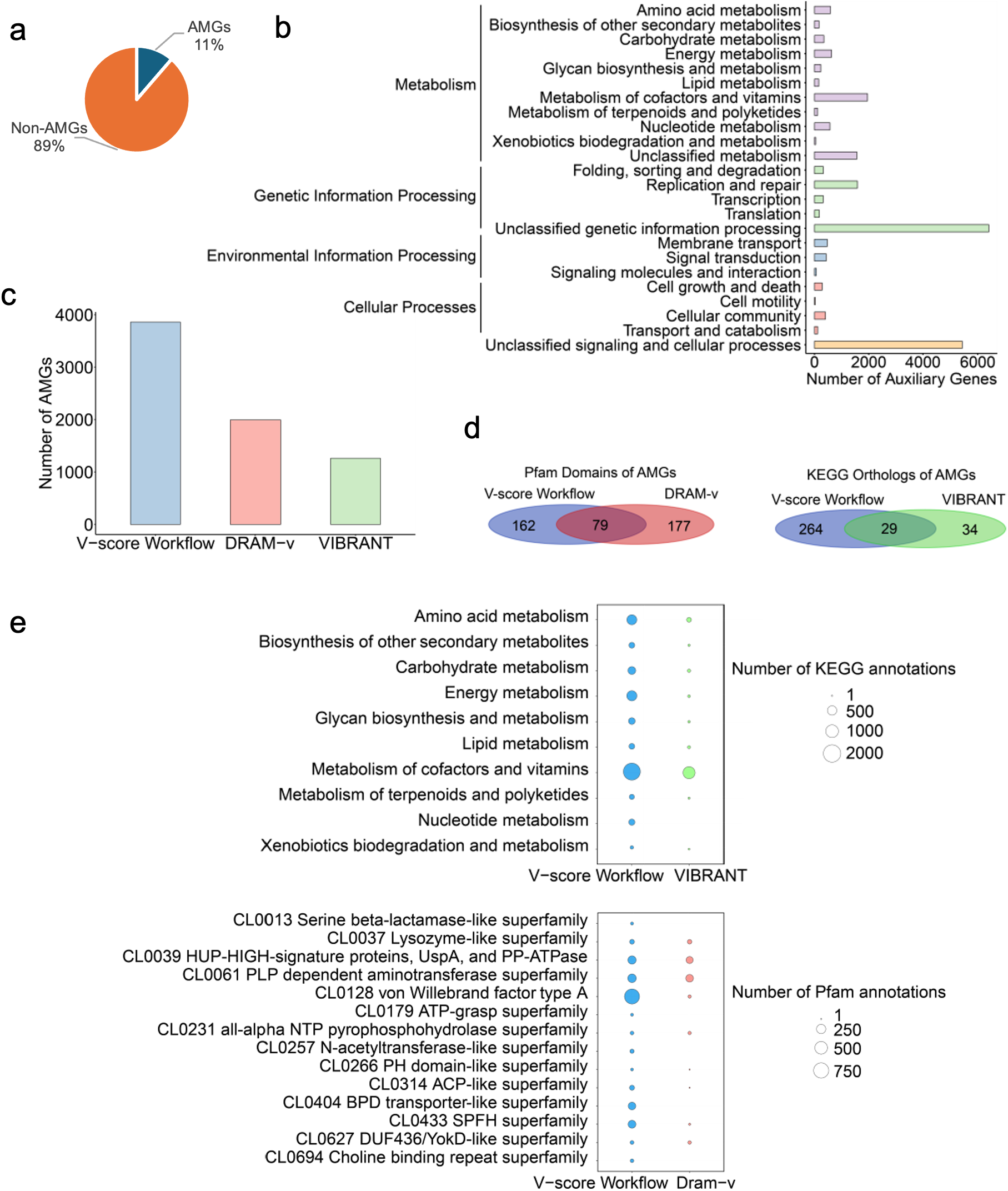
Auxiliary genes identified in the study and comparison with existing methods. **a**, auxiliary gene composition. AMG: auxiliary metabolic genes. **b**, Potential functions of auxiliary genes with annotations detected using the V-score workflow. Purple bars represent categories within Metabolism, green bars denote Genetic Information Processing, blue bars indicate Environmental Information Processing, pink bars correspond to Cellular Processes, and orange bars represent unclassified signaling and cellular processes. **c**, Number of AMGs identified by the V-score workflow compared to other existing methods, including DRAM-v and VIBRANT. **d**, Overlap and unique Pfam domains or KEGG orthologs of AMGs identified by the V-score workflow, DRAM-v, and VIBRANT. **e**, Comparison of the number of KEGG or Pfam annotations of AMGs identified using the V-score workflow, DRAM-v, and VIBRANT. Please note that VIBRANT exclusively outputs results that contain KEGG annotations, while DRAM-v mainly generated Pfam annotations for AMGs identified in the study.

In comparison to other existing approaches, our workflow significantly outperformed widely used approaches including VIBRANT^17^ and DRAM-v^35^, as demonstrated by the identification of AMGs. When applied to the same set of viral genomes, our V-score workflow identified 3,859 AMGs (**Fig. 5c**; **Supplementary Table S13**), while VIBRANT and DRAM-v identified only 1,261 and 1,993 AMGs, respectively (**Fig. 5c**; **Supplementary Tables S14** and **S15**). Notably, only a small fraction of Pfam domains or KEGG orthologs of AMGs were commonly identified by three approaches (**Fig. 5d**), with most AMGs being unique to each method. This suggests that our V-score workflow reveals novel functions that are often overlooked by existing AMG detection tools. Some unique metabolic enzymes uncovered by our method include the serine beta-lactamase-like superfamily (Pfam clan accession: CL0013), ATP-grasp superfamily, N-acetyltransferase-like superfamily, and Choline binding repeat superfamily (**Fig. 5e**). Furthermore, our workflow outperformed VIBRANT, as shown by the higher number of AMGs identified across all KEGG categories (Fig. 5d). Collectively, these findings demonstrate that the V-score-based approach can detect a greater number of potential AVGs with high precision.

### Signatures of population differentiation and enhancing genome binning strategies

Characterizing new viral species in complex systems is crucial for understanding how microbial interactions impact the spread of diseases and their development and impact on health^43^. AV-scores and AV_L_-scores capture the association and frequency of viral genomes, as well as their differentiation from other genomes. Leveraging these signatures, we assessed whether AV-score and AV_L_-score analyses could effectively recover viral metagenome-assembled genomes (vMAGs) from a mixed metagenome. Prior to this assessment, we evaluated the ability of AV-scores and AV_L_-scores to cluster population genomes, to verify their relevance and effectiveness in the context of genome binning. We analyzed a dataset of 11 viral species that were available in the NCBI RefSeq database. We found that the similar viral species had very similar AV-scores or AV_L_-scores, while different species exhibited distinct scores (**Fig. 6a**). This highlights the reliability and accuracy of these metrics for viral genome classification and identification of novel species. For instance, changes in the gut phage population have been repeatedly linked to various gastrointestinal diseases^44-46^. The application of AV-scores or AV_L_-scores into gut phage population studies would provide opportunity to differentiate viral populations in complex host-associated systems and contribute to uncover certain disease-related viral species.

**Fig. 6.**
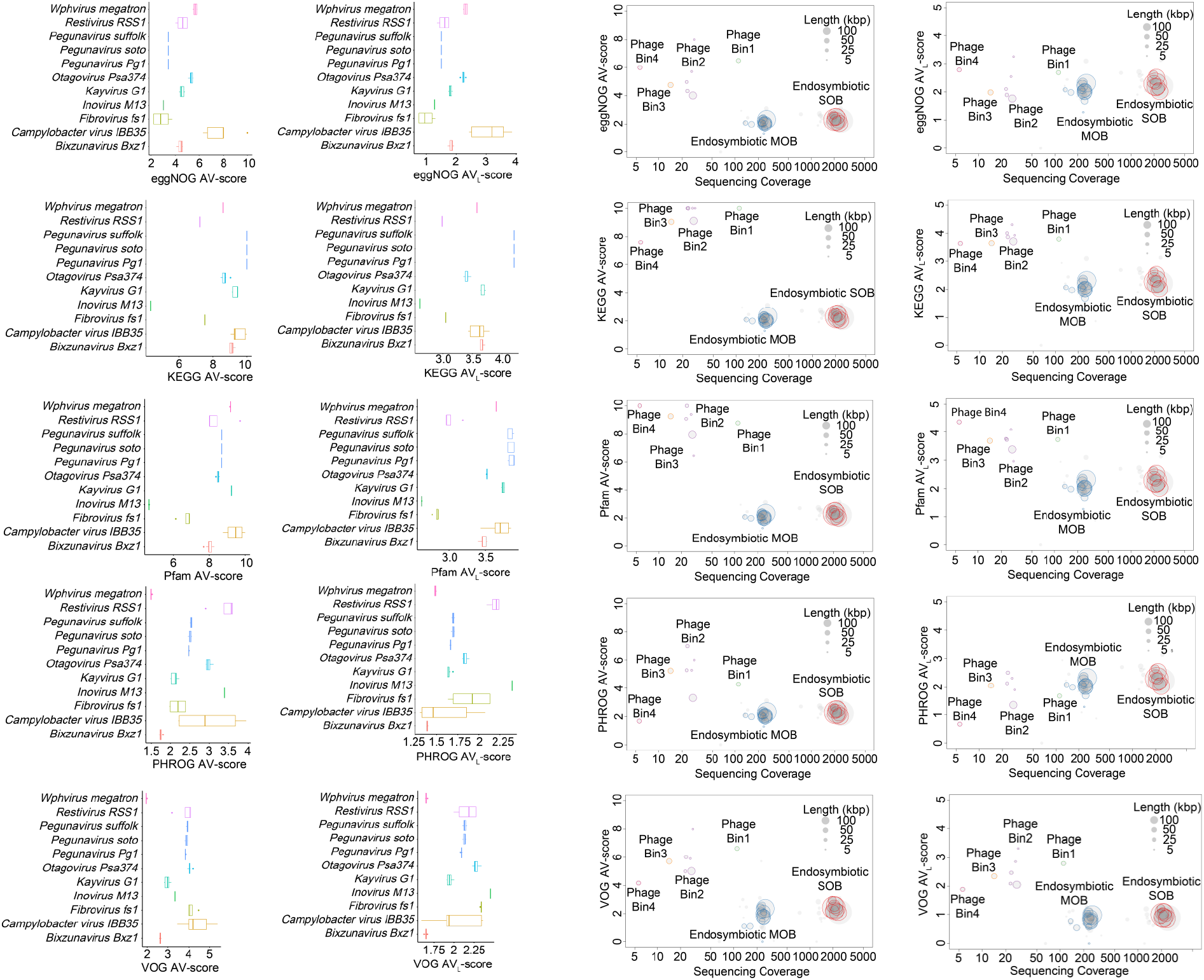
Application of AV-scores, and AV_L_-scores to metagenomic binning. **a**, Viral population differentiation with AV-scores and AV_L_-scores. Viral species include *Bixzunavirus Bxz1* (n = 13), *Campylobacter* virus IBB35 (n = 5), *Fibrovirus fs1* (n = 4), *Inovirus M13* (n = 8), *Kayvirus G1* (n = 15), *Otagovirus Psa374* (n = 7), *Pegunavirus Pg1* (n = 6), *Pegunavirus soto* (n = 5), *Pegunavirus Suffolk* (n = 6), *Restivirus RSS1* (n = 4), and *Wphvirus megatron* (n = 4). The horizontal line that splits the box is the median, the upper and lower sides of the box are upper and lower quartiles, whiskers are 1.5 times the interquartile ranges and data points beyond whiskers are considered potential outliers. **b**, Genome binning with AV-scores, AV_L_-scores, and sequencing coverage for a snail-associated metagenome. SOB: sulfur-oxidizing bacteria; MOB: methane-oxidizing bacteria.

AV-scores and AV_L_-scores facilitate species clustering and even strain-level differentiation, as demonstrated by the distinct separation of viral populations based on AV-scores and AV_L_-scores of VOG and PHROG (**Fig. 6a**). AV-scores and AV_L_-scores can therefore be effective metrics for differentiating microbial and viral species or strains and facilitating genome binning in metagenomic studies. We next tested a host-associated metagenome. The analysis of a deep-sea snail microbiome using AV-scores, AV_L_-scores, and sequencing coverage demonstrated the effectiveness of these metrics in genome binning of microbes and viruses (**Fig. 6b**). We observed clear clustering of four phage genome bins and two bacterial chromosome bins, which was consistent with a prior study^47^, thereby highlighting the capability of these metrics to differentiate between viral and bacterial genomes accurately. This approach could complement current tools, such as vRhyme^48^, and enhance the construction of vMAGs that more accurately represent the true composition of viruses within a sample. Significantly, this approach would reduce the overestimation of viral diversity that can result from the assumption that a single genome fragment represents an uncultivated viral genome (UViG) or a viral population^49, 50^.

## DISCUSSION

In conclusion, V-scores, V_L_-scores, AV-scores, and AV_L_-scores represent powerful quantitative metrics that describe the virus-like nature and origin of protein families and genomes. These metrics can serve as the foundation of new tools to advance viral genomics, ecology, and evolutionary analyses. By enabling open and public distribution of these scores ((https://anantharamanlab.github.io/V-Score-Search/), we propose that they will propagate broadly in microbiology. Our approach allows for citation of these scores using databases identifiers like for KEGG, Pfam etc or using protein annotations. For example, a picornavirus capsid protein (PF00073) has a V-score of 10 implying a strong virus association while a Hepatitis C virus capsid protein (PF01543) has a V-score of 1 implying a weaker virus association, presumably because its proteins domains are not specific to capsids.

The versatility of these scores allows for their incorporation into diverse genomics tools such as for genome binning, genome completion, virus identification in complex datasets, and identification of AMGs. These scores can enhance genome binning strategies by providing an additional layer of resolution in separating viral from non-viral sequences. This capability is especially valuable in metagenomic studies, where the accurate classification of sequences is critical for understanding the composition and dynamics of microbial communities. By integrating metrics like AV-scores and AV_L_-scores, researchers could develop more refined tools for viral identification, potentially leading to the discovery of novel viral genomes and a deeper understanding of virus-host interactions. The broader implication of this approach is that it allows for more nuanced and data-driven differentiation between viral and non-viral entities at both the gene and genome levels. This could revolutionize how we identify and characterize viruses in complex biological systems, offering new insights into viral evolution, diversity, and function. The quantitative nature of the metrics also opens up possibilities for automating and scaling viral genome study across large datasets, for example the completeness assessment of linear viral genome in cases where identifiable terminal repeats are absent^6^, making it an invaluable resource in the field of viral (meta)genomics.

## METHODS AND MATERIALS

### Viral protein database construction

Viral protein sequences were downloaded from public databases (accessed January 2024), including the National Center for Biotechnology Information (NCBI) RefSeq database, the Virus Orthologous Groups (VOG) database (version 221, https://fileshare.csb.univie.ac.at/vog/), the Prokaryotic Virus Remote Homologous Groups (PHROG) database^51^, and the IMG/VR Viral Resources v4.1^52^. Protein sequences from IMG/VR Viral Resources were filtered and we only retained high-quality and medium-quality viral sequences that were assessed by CheckV v1.0.1^53^. To dereplicate proteins, MMseqs2 linclust version 13.45111^54^ was used with an identity cutoff of 95% (custom parameters: --min-seq-id 0.95 --cluster-mode 2 --cov-mode 1 -c 1.0), and generated non-redundant 18,435,589 protein sequences.

### Annotation profile database selection

To construct a wide range of associations between annotation profiles and viral proteins, a diverse collection of profile databases was selected. The profile databases included Kyoto Encyclopedia of Genes and Genomes (KEGG) KOfam (version 2024-01-01)^55^ that is a customized Hidden Markov Models (HMMs) profile collection of KEGG Orthologs, Pfam-A (release 36.0)^56^ database of a large collection of diverse protein families, and eggNOG (version 5.0)^57^ that is a database of non-supervised orthologs created from a large number of various organisms. Two additional curated viral ortholog collections are the VOG (release 221, vogdb.org) and PHROG both of which were constructed based on remote homology.

### V–score and V_L_-score generation

The V-score and V_L_-score for each annotation profile in the KEGG, Pfam, eggNOG, PHROG, and VOG databases was determined based on the number of significant hits (E-value ≤ 10^−5^) identified by hmmsearch (HMMER 3.4)^58^ and MMseqs2. For V-score, the resulting number was divided by 100, with a maximum limit set at 10 after division. For V_L_-score, the resulting number was scaled down using the common logarithm (base 10) without a maximum limit. In the case of annotations containing viral keywords including “virus”, “viral”, “phage”, “portal”, “terminase”, “spike”, “capsid”, “sheath”, “tail”, “coat”, “virion”, “lysin”, “holin”, “base plate”, “lysozyme”, “head”, “structural”, or “Viral protein families”, protein families/annotations were assigned adjusted V-score of 1 and V_L_-score of 2 if the original V-score was less than 1 and V_L_-score less than 2. Each annotation profile is given a V-score and a V_L_-score, serving as metrics for virus association. It is important to note that the V-scores do not consider virus specificity or association with non-viruses and have been manually adjusted to prioritize viral hallmark genes.

### Databases of chromosomes, plasmids, and viral genomes for AV-score and AV_L_-score generation

Databases of prokaryotic chromosomes, plasmid sequences, and prokaryotic viral genomes were constructed for the generation of AV-score and AV_L_-score. Prokaryotic genomes (release 214) were downloaded from the Genome Taxonomy Database (GTDB; gtdb.ecogenomic.org)^59, 60^. We assessed the quality of each genome with a quality score (score = completeness − 5 × contamination − 0.05 × no. scaffolds)^8^, genomes of each GTDB family with the highest quality score were selected as family representatives to reduce computational load and taxonomic bias. As a result, 4,304 bacterial and 509 archaeal genomes were selected to be used in the following analyses. Then, provirus and provirus-like sequence regions were identified with VirSorter2 version 2.2.4 and VIBRANT version 1.2.1 and removed from the selected prokaryotic genomes. Additionally, plasmid sequences (sequence headers containing the word “plasmid”) were removed from the selected prokaryotic genomes. For plasmids and prokaryotic viruses, 50,523 plasmid sequences were downloaded from the PLSDB database version 2023_11_23^61^ and viral genomes were downloaded from the NCBI RefSeq database^62^ (retrieved in January 2024). To retrieve prokaryotic viral genomes, the GenBank dabase division PHG was used to filter bacterial and archaeal viruses in the RefSeq database. Finally, 5,800 genomes of prokaryotic viruses were retained.

### Generation of AV-score and AV_L_-score

Databases of prokaryotic chromosomes, plasmids, and prokaryotic viruses constructed above were used to calculate the AV-score and AV_L_-score for each genome. Each whole genome of prokaryotic viruses, plasmids, and chromosomes were randomly split into non-overlapping, non-redundant genome fragments at length from 1 to 15 kb to simulate metagenome-assembled sequences. Proteins of each whole genome and split genome fragment were predicted using Prodigal V2.6.3 (parameters: -m -p meta)^63^. Hmmsearch^58^ (HMMER 3.4, parameter: -E 10^−5^) was used to match the proteins of prokaryotic viruses, plasmids, prokaryotes to the HMM profiles of KEGG, VOG, and Pfam. EggNOG-mapper version 2.1.12 (parameters: -m mmseqs --evalue 10^−5^)^64^ was used to annotate the proteins with the eggNOG database. MMseqs2 (parameter: E-value ≤ 10^−5^) was employed to search the predicted proteins against the PHROG database. Only the hit with the highest score was kept. Post this, V-score and V_L_-score of KEGG, VOG, eggNOG, Pfam, and PHROG were assigned to each protein. For comparison between viruses, plasmids, and chromosomes, AV-score and AV_L_-score were calculated for each whole genome and genome fragment. The AV-score and AV_L_-score of KEGG, Pfam, and eggNOG were expressed as:

AV-score = (Sum of V-score of Proteins with Significant Hits) / (Number of Proteins with Significant Hits);

AV_L_-score = (Sum of V_L_-score of Proteins with Significant Hits) / (Number of Proteins with Significant Hits).

The AV-score and AV_L_-score of PHROG and VOG were calculated as:

AV-score= (Sum of V-score of Proteins with Significant Hits) / (Total Number of Proteins Encoded in a Genome);

AV_L_-score = (Sum of V_L_-score of Proteins with Significant Hits) / (Total Number of Proteins Encoded in a Genome).

### Generation of cutoffs of V_L_-score, AV-score, and AV_L_-score for viral-like protein/genome determination

To predict the probability of a protein or a genome sequence being viral, the cutoff (see the definition of cutoff in Supplementary Fig. S10) of the V_L_-score, AV-score, and AV_L_-score generated above was examined to determine the probability. The cutoff of the AV-score was set from 0 to 10 with steps of 0.2. The cutoff of the V_L_-score/AV_L_-score was set from 0 to 5 with step 0.1. The probability of a protein/genome being viral was represented by the fraction of normalized viral proteins/genomes (Nv) compared with normalized plasmids (Np) and chromosomes (Nc) at each cutoff. The fraction at each cutoff was expressed as:

For proteins:

Fraction = Nv / (Nv+Np+Nc)

Nv = (Number of viral proteins with scores above cutoff) / (Total number of viral proteins)

Np = (Number of plasmid proteins with scores above cutoff) / (Total number of plasmid proteins)

Nc = (Number of chromosome proteins with scores above cutoff) / (Total number of chromosome proteins)

For genome sequences:

Fraction = Nv/(Nv+Np+Nc)

Nv = (Number of viral sequences with scores above cutoff) / (Total number of viral sequences)

Np = (Number of plasmid sequences with scores above cutoff) / (Total number of plasmid sequences)

Nc = (Number of chromosome sequences with scores above cutoff) / (Total number of chromosome sequences)

Polynomial regression with the smoothing method “lm” was used to predict the best-fit curve that matches the pattern of the cutoff and probability. The cutoffs for the probability of 70% and 90% were predicted according to estimated polynomial regression equations. If a protein or genome sequence has a score above the cutoff for the probability of 70%, this protein or sequence was determined as a “likely” viral-like protein or sequence. If a protein or genome sequence has an AV-score above the cutoff for the probability of 90%, this protein or sequence was determined as a “most likely” viral-like sequence.

### Applying cutoffs to the identification of viral sequences

Metagenomes from host-associated microbiomes were analyzed as a use case to demonstrate the application of viral genome identification. Raw Illumina reads of one snail-associated metagenome^47^, three sponge-associated metagenomes^20, 21^, three human-associated metagenomes^65^, and 32 coral-associated metagenomes^66^ were retrieved from NCBI (BioProject accessions: PRJNA612619 for snail, PRJNA552185 for sponge, PRJNA763232 for human, PRJNA574146 for coral). The downloaded reads were then trimmed using Trimmomatic^67^ (version 0.36) with custom settings (ILLUMINACLIP: TruSeq3-PE.fa:2:30:10 LEADING:3 TRAILING:3 SLIDINGWINDOW:4:15 MINLEN:40). Trimmed reads from the sponge-, human-, and snail-associated microbiomes were assembled with MEGAHIT^68^ version 1.2.9 using default parameters, while reads from coral-associated microbiomes were assembled using SPAdes^69^ version 3.11.1 with custom settings (--meta, k-mer sizes varied from 51 to 91, with a 10-mer step size). The assembled metagenomes were then functionally annotated using VOG, PHROG, KEGG, and Pfam via Hmmsearch (HMMER 3.4, parameter: -E 10-5) and MMseqs2 (E-value ≤ 10-5). AV-scores for VOG, PHROG, KEGG, and Pfam were subsequently calculated for each sequence. Predicted viral genomes were identified based on the following criteria: (1) sequences with at least one AV-score (from VOG, PHROG, KEGG, or Pfam) exceeding the corresponding cutoffs for each fragment size (e.g., a PHROG AV-score > 4.24 or a VOG AV-score > 4.91 for a 2.5 kb scaffold; detailed cutoffs by fragment size are provided in Supplementary Table S10). For sequences larger than 15 kb, cutoffs for 14−15 kb fragments were used. (2) Sequences meeting criterion (1) were further filtered for completeness >0%, as assessed by CheckV^53^ v1.0.13. In parallel, geNomad^8^ v1.7.411, VirSorter2^18^ v2.2.3, VIBRANT^17^ v1.2.0, and DeepVirFinder^19^ v1.0, (score ≥ 0.75, *p* < 0.05), were used to identify viral sequences from the host-associated metagenomes, allowing for a comparison between the V-score-based and specific gene- or hallmark- or machine learning-based viral identification methods. For consistency, viral sequences identified by geNomad, VirSorter2, VIBRANT, and DeepVirFinder were also required to have completeness >0%, as assessed by CheckV v1.0.13.

### Applying cutoffs to the assessment of proviral sequences

Cutoffs of AV-scores and AV_L_-scores of whole genomes in Supplementary Table S10 were used for the assessment on proviral sequences by estimating the consistency of our method with a custom prophage database. The custom prophage database developed by Arndt *et al*.^22^ were downloaded from PHASTER (https://phaster.ca/databases). Then prophage sequences in the database were functionally annotated with VOG and PHROG using Hmmsearch (HMMER 3.4, parameter: -E 10^−5^) and MMseqs2 (E-value ≤10^−5^), followed by the calculation of the AV-scores and the AV_L_-scores of VOG and PHROG for each prophage. Any prophage sequences with an AV-score or AV_L_-score above their corresponding cutoff were considered consistent with the prophage database.

To show a potential application in prophage boundary identification, one experimentally verified provirus, *Enterobacteria* phage P88^70^, and its host were selected and downloaded from NCBI (*Escherichia coli* GenBank: GCA_001005685.1). Proteins of prophage and host genomes were predicted using Prodigal V2.6.3 (parameters: -m -p meta)^63^. Hmmsearch^58^ (HMMER 3.4, parameter: -E 10^−5^) was used to match the proteins of prophages and hosts to the HMM profiles of VOG. MMseqs2 with a custom parameter (E-value ≤ 10^−5^) was used to search prophage and host proteins against the PHROG database. Only the best hit to each protein was retained. Then V-score and V_L_-score of VOG and PHROG were assigned to each protein, followed by calculating AV-score and AV_L_-score for each prophage and adjacent host sequence. The gene feature plots of prophages were generated and visualized with DNA Features Viewer^71^.

### Database construction for benchmarking on AMGs identification

We assembled a database of 17 KEGG and Pfam HMM profiles (V_L_-scores < 3 for KEGG annotations or V_L_-scores < 3 for Pfam annotations) representing AMGs experimentally demonstrated to affect host metabolism^72-76^ (Supplementary Table S16) and a database of 10 selected HMMs that represent non-AMGs (Supplementary Table S17). From IMG/VR v4^52^, we compiled a database of 5,116 high-quality^50^ viral genomes (Supplementary Table S18) containing the 17 experimentally verified AMGs, the 10 non-AMGs, and genomes with neither to obtain a representative sample. We ensured each viral genome had a known host genus, and compiled a database of 180 host genomes (containing homologs of the 17 experimentally verified AMGs) representing the known host genera (Table S13). We used GeNomad^8^ v1.7.4 to predict viral scaffolds in the 180 host genomes and removed viral scaffolds binned in host genome assemblies (Supplementary Table S19).

Open reading frames in all virus and host genomes were identified and translated using pyrodigal-gv^8, 63^ v0.3.1 (github.com/althonos/pyrodigal-gv). Translated proteins were aligned to Pfam-A^56^ v36.0 HMMs and KEGG^77^ KO HMMs using pyhmmer^58, 78^ v0.10.10 hmmsearch^58^ with a maximum e-value of 1e-05. For proteins aligning to multiple HMM profiles within the same database, the highest scoring alignment was reported. Each protein with a Pfam or KEGG functional annotation was assigned its corresponding Pfam or KEGG V_L_-score and V-score.

### Workflow for AMGs identification

Using the database of 17 KEGG and Pfam HMM profiles, we identified potential AMGs by searching for each protein with Pfam V_L_-score < 3 or KEGG V_L_-score < 3 and with Pfam and KEGG V-scores < 10. We distinguished AMGs from host-encoded metabolic genes by averaging the V_L_-scores of all KEGG or Pfam annotations in entire scaffolds, establishing a minimum scaffold Pfam/KEGG AV_L_-score of 3 as optimal for differentiating viral from host scaffolds. Thus, for a gene flagged as a potential AMG using our predefined V_L_-score and V-score cutoffs, we also required that the scaffold encoding the gene have an AV_L_-score > 3 for Pfam/KEGG annotations and AV-score > 4.81 for KEGG annotations or AV-score > 4.39 for Pfam annotations.

It is recommended by community standards for AMG analysis that a potential AMG should be validated by ensuring it is flanked on both the upstream and downstream sides by hallmark genes^35, 36^. However, given the poor annotation rate of virus proteins, this also impacts the identification of AMGs. Here, we conducted our flanking verification approach by running our AMG identification workflow using viral hallmark genes to verify flanking regions of potential AMGs. We defined viral hallmark genes in our KEGG and Pfam HMM databases as previously described^79^; any HMM profile with an annotation/description containing any of the following keywords: virion structure (truncated from *structure* to account for matches to the terms “structure” or “structural”), capsid, portal, tail, and terminase. A list of KEGG and Pfam HMMs defined as viral hallmark genes this way are provided in Supplementary Table S20. In parallel, we verified that AMGs identified with our workflow were flanked on both sides by at least one gene with a V-score of 10 within 10 kb of the AMG, recognizing that viral genes with unknown functions may still be characteristically viral. The verification approach may not be necessary when analyzing complete or cultured viral genomes, so we report results with and without flank verification.

### Assessment on performance of the workflow for AMGs identification

To assess the performance of our workflow, we established true positives and negatives for AMGs in our test genome dataset. A gene encoded by a viral scaffold with an annotation in the experimentally verified AMG database was considered a true positive, while any host-encoded gene in the experimentally verified AMG database was considered a true negative. Genes encoded on viral scaffolds with annotations matching any of 10 selected HMMs that represent non-AMGs were also considered true negatives. Any other gene, encoded on a known host or viral genome, that was not annotated with the experimentally verified AMG database or non-AMG database was not considered a true positive or negative.

In addition to the true positives and negatives, we predicted positives and negatives. To ensure that we did not analyze viral genes in host genomes, all genes encoded on host scaffolds predicted as viral were removed before we predicted the positives and negatives of our AMG identification workflow. Predicted positives were any gene, encoded on a known host or viral scaffold, that met the following criteria: (1) the gene has a Pfam V_L_-score < 3 or a KEGG V_L_-score < 3, (2) the gene has a Pfam V-score < 10 or a KEGG V-score < 10, (3) the gene is encoded on a scaffold with a Pfam AV_L_-score > 3 or a KEGG AV_L_-score > 3, (4) the gene is encoded on a scaffold with a Pfam AV-score > 4.39 or a KEGG AV-score > 4.81, (5) the gene is flanked to the left and right by at least one other gene with a V-score of 10 within a 10 kb distance (only applies to results reporting prediction “with flank verification”). Any gene with an annotation belonging to the AMG database or the non-AMG database that did not meet these criteria was considered a predicted negative. Genes without annotations to the non-AMG or the AMG database were not predicted as positives or negatives. The counts of true positives, true negatives, predicted positives, and predicted negatives were used to construct the confusion matrices in Supplementary Table S12.

### Identification of auxiliary genes using our workflow and other existing approaches

We assembled a dataset of 5,116 high-quality viral genomes from IMG/VR v4^52^ (Supplementary Table S18). All viral genes were evaluated for potential auxiliary functions using the AMG identification workflow, both with and without flank verification. Genes annotated under KEGG’s “sulfur relay system” or “metabolic pathways” category, excluding those related to nucleotide metabolism or sulfonate transport system substrate-binding proteins, were considered potential AMGs. Additionally, auxiliary genes with KEGG and PFAM annotations were cross-referenced against a viral AMG database^35^, which includes experimentally verified AMGs from previous studies^26, 37, 72-76, 80, 81^. PFAM and KEGG accessions associated with AMGs were retrieved, and ORFs containing these accessions were retained and integrated into the AMG dataset. To compare our approach with other existing tools to identify AMGs, we ran VIBRANT^17^ with the “annoVIBRANT” implementation (github.com/AnantharamanLab/annoVIBRANT) and DRAM-v^35^ on the same set of high-quality viral genomes. For DRAM-v only the AMGs with a score of 1 were retained, which indicates the presence of at least one hallmark gene on both sides, suggesting the gene is likely viral.

### Visualization of V_L_-scores, and V-scores of phage and host genomes containing *psbA*

We visualized the genomic context of one predicted AMG, the photosystem II P680 reaction center D1 protein (*psbA* KO K02703), in viral and host genomes. We identified one *Prochlorococcus* host genome (GenBank GCA_003214355.1) and two viral genomes (IMGVR_UViG_2716884766_000001 and IMGVR_UViG_2716884767_000001) encoding *psbA* (Supplementary Table S18) predicted by IMG/VR to be *Prochlorococcus* phages. We plotted genes within localized regions of these genomes using the R package gggenomes^82^ v1.0.0 using annotations, V_L_-scores, and V-scores obtained as described above.

### Viral species differentiation based on AV-score and AV_L_-score

Reference prokaryotic viruses were used for assessment on viral population differentiation based on AV-score and AV_L_-score. Lineage of the reference viruses was downloaded from virushostdb (https://www.genome.jp/virushostdb). According to the lineage information of each viral RefSeq genome, 11 species of reference prokaryotic viruses were selected (each species with ≥ 4 genomes). Viral species include *Bixzunavirus Bxz1, Campylobacter* virus IBB35, *Fibrovirus fs1, Inovirus M13, Kayvirus G1, Otagovirus Psa374, Pegunavirus Pg1, Pegunavirus soto, Pegunavirus Suffolk, Restivirus RSS1*, and *Wphvirus megatron*. Viral genomes were annotated with databases of VOG, PHROG, KEGG, Pfam, and eggNOG using Hmmsearch (HMMER 3.4, parameter: -E 10^−5^), MMseqs2 (parameter: E-value ≤ 10^−5^), or EggNOG-mapper version 2.1.12 (parameters: -m mmseqs --evalue 10^−5^). In the following, the AV-score and AV_L_-score of each genome were calculated. Detailed information of NCBI RefSeq accessions and AV-score and AV_L_-score of viral genomes was provided in Supplementary Table S21.

### Metagenome binning with AV-score and AV_L_-score

The metagenome of deep-sea snail (*Gigantopelta aegis*) microbiome^47^ was analyzed as a use case to show an application in genome binning. Raw Illumina reads of the snail *G. aegis* metagenome were retrieved from NCBI (BioProject accession: PRJNA612619). Then the downloaded reads were trimmed by Trimmomatic (version 0.36)^67^ with custom setting (ILLUMINACLIP: TruSeq3-PE.fa:2:30:10 LEADING:3 TRAILING:3 SLIDINGWINDOW:4:15 MINLEN:40). Scaffolds of the genomes of two bacterial endosymbionts and four phages infecting the endosymbionts were mapped to the trimmed reads with Bowtie2 version 2.3.4^83^ and SAMtools version 1.6^84^ to calculate sequencing coverage. Additionally, the microbial genomes were functionally annotated with VOG, PHROG, KEGG, Pfam, and eggNOG with Hmmsearch (HMMER 3.4, parameter: -E 10^−5^), MMseqs2 (E-value ≤10^−5^), or EggNOG-mapper version 2.1.12 (parameters: -m mmseqs --evalue 10^−5^), followed by the calculation of AV-score and AV_L_-score for each scaffold in a genome. Finally, we manually binned bacterial and phage scaffolds (length ≥5 kb) following a previously described approach^85^ on the basis of AV-score and AV_L_-score, sequencing depth, phage hallmark genes, and bacterial conserved single-copy genes.

## Conflict of interest

The authors declare no competing interests.

## Correspondence

Correspondence and requests for materials should be addressed to Karthik Anantharaman.

## Funding

This research was supported by the National Science Foundation under grant number DBI2047598 and National Institute of General Medical Sciences of the National Institutes of Health under award number R35GM143024 to KA. JCK was supported by an NSF Graduate Research Fellowship.

## Acknowledgments

We thank members of the Anantharaman Laboratory for discussions and feedback on this manuscript.

## Author contributions

Conceptualization: KZ and KA. Methodology: KZ and KA. Open access software: KZ and PJB. Validation: KZ, JCK, EDC. Formal analysis: KZ, JCK. Investigation: KZ, JCK, EDC and KA. Resources: KA. Data curation: KZ and JCK. Original draft: KZ and KA. Writing-review and editing: KZ, JCK, EDC, PJB and KA. Visualization: KZ and JCK. Supervision: KA. Project administration: KA. Funding acquisition: KA.

## Notes

### Competing Interest Statement

The authors have declared no competing interest.

