## Supplementary Figures for "V- and V_L_-Scores Uncover Viral Signatures and Origins of Protein Families"

for

Percentage of Annotations

|  | Non-viral | Viral |
| --- | --- | --- |
| KEGG | 24117 | 1900 |
| Pfam | 19598 | 1197 |
| eggNOG | 341431 | 8318 |

**Fig. S1.** Proportion of viral annotation entries in the current KEGG, Pfam, and eggNOG databases. Viral entries in each database represent less than 10% of the total.

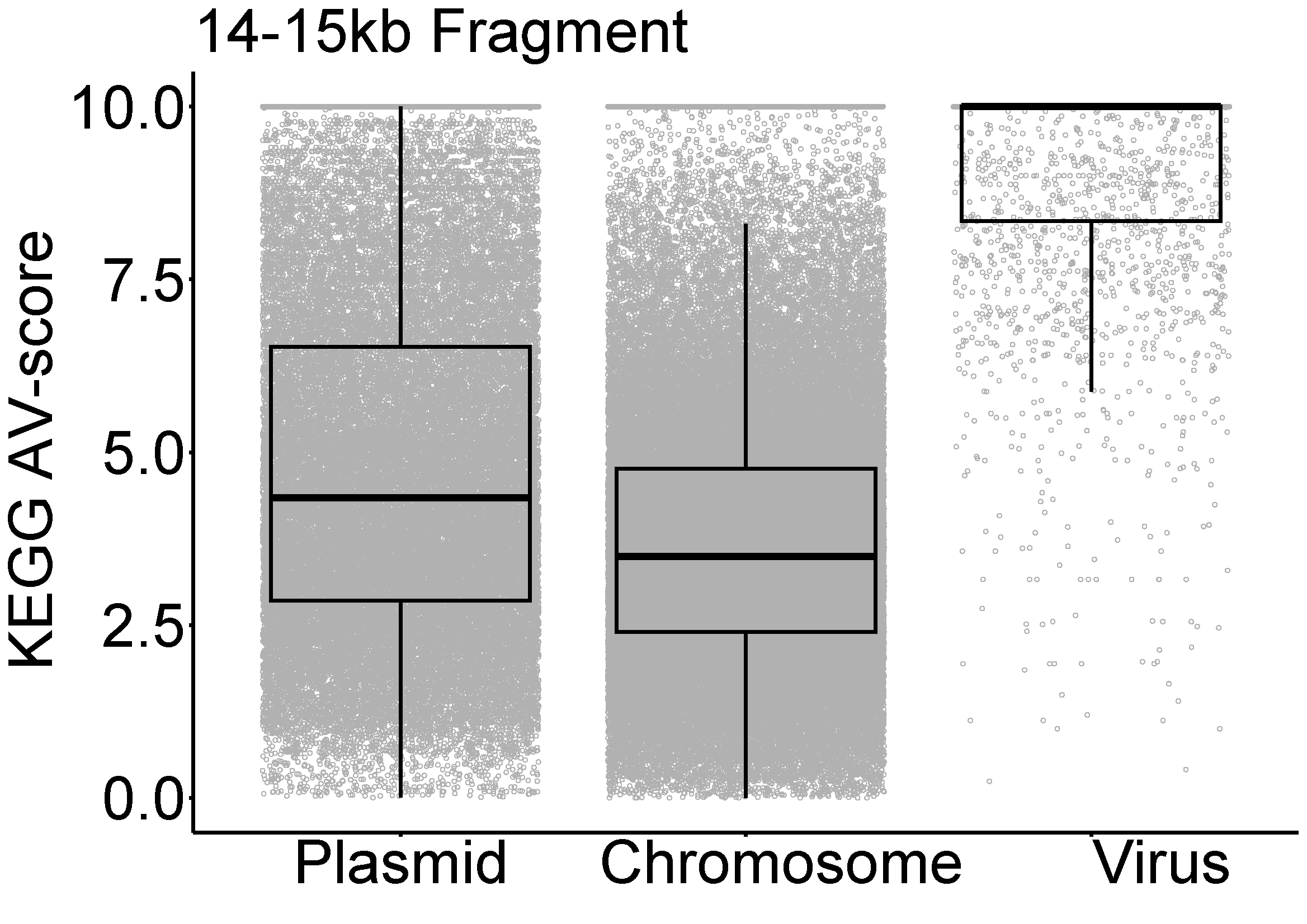

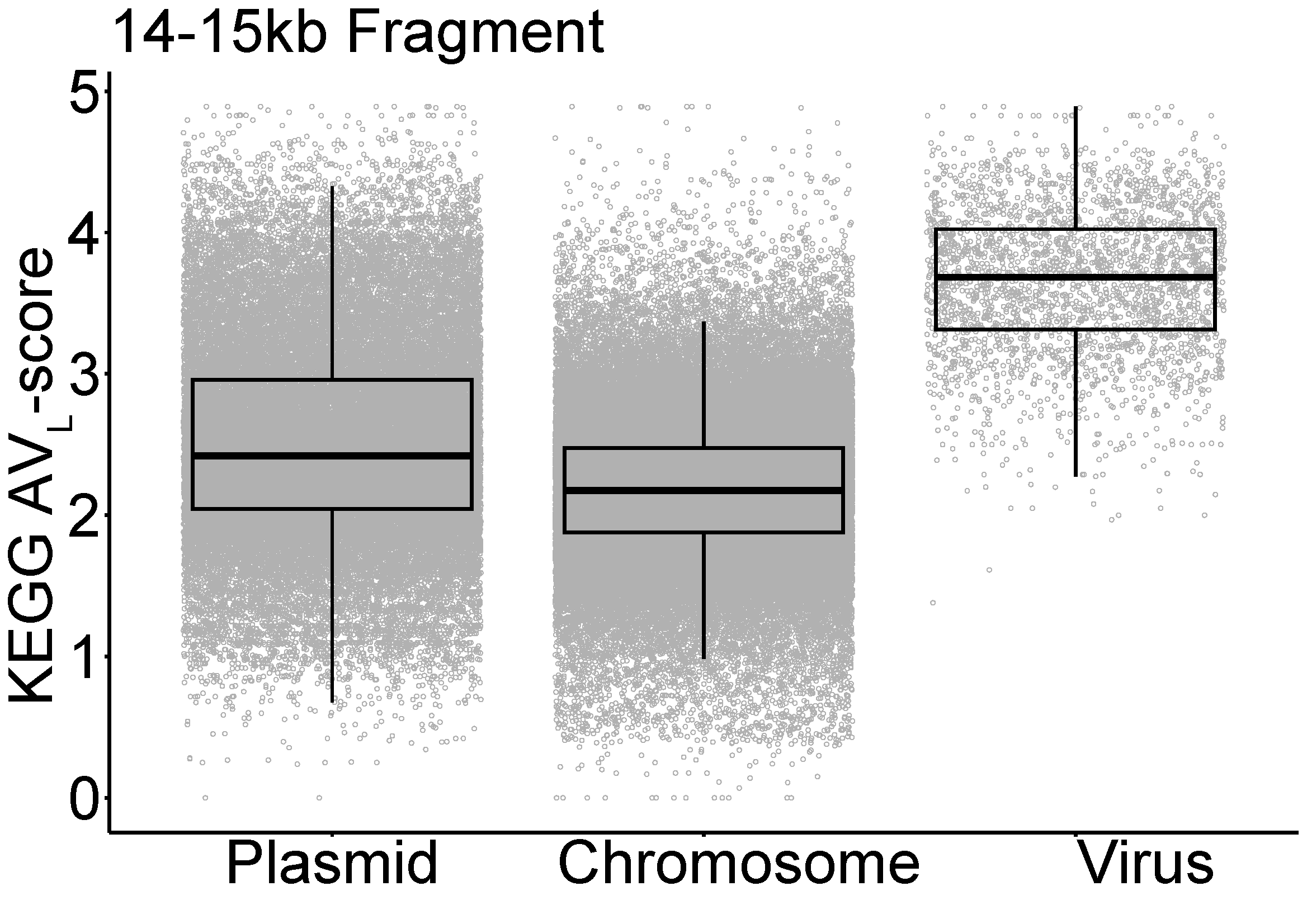

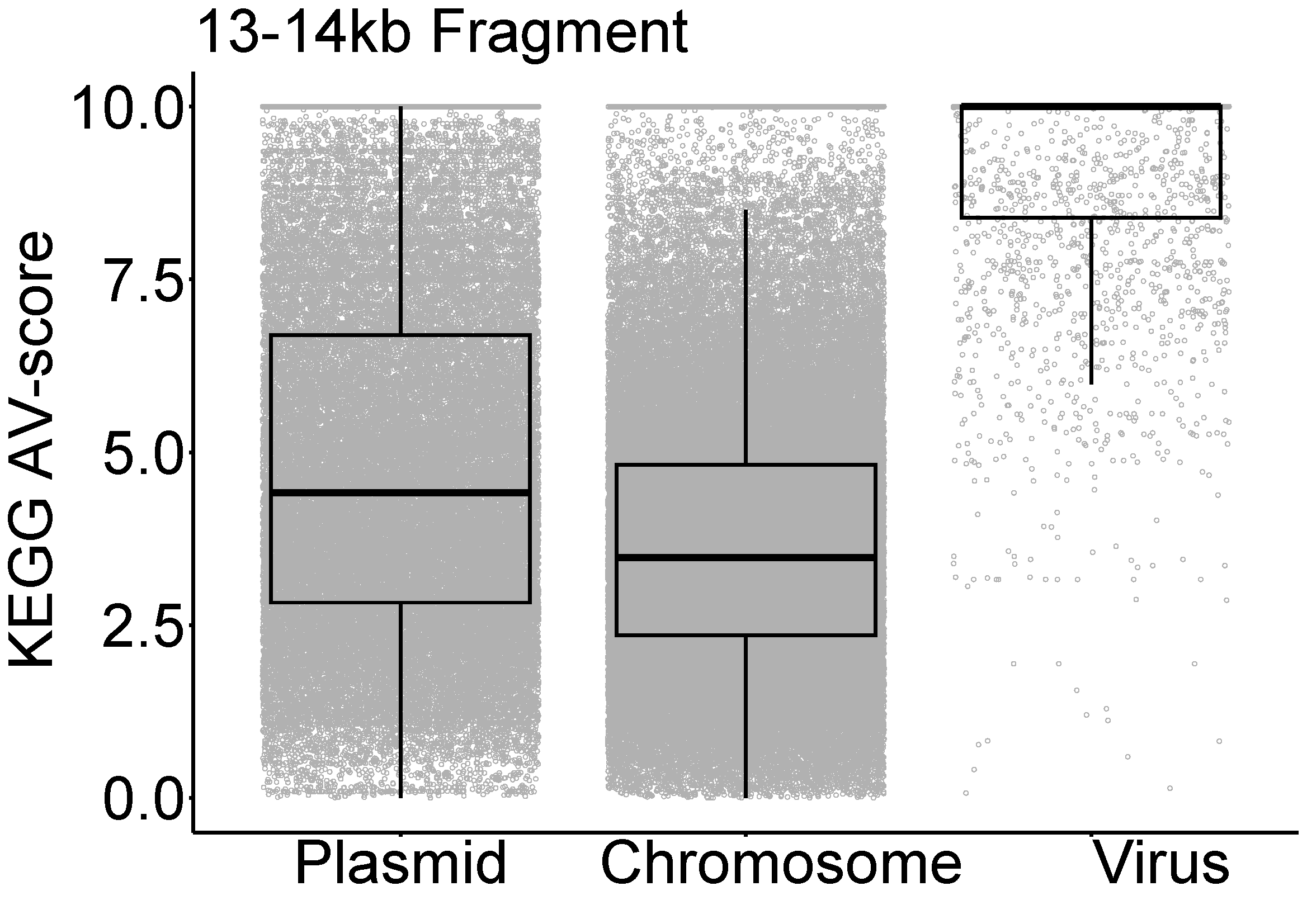

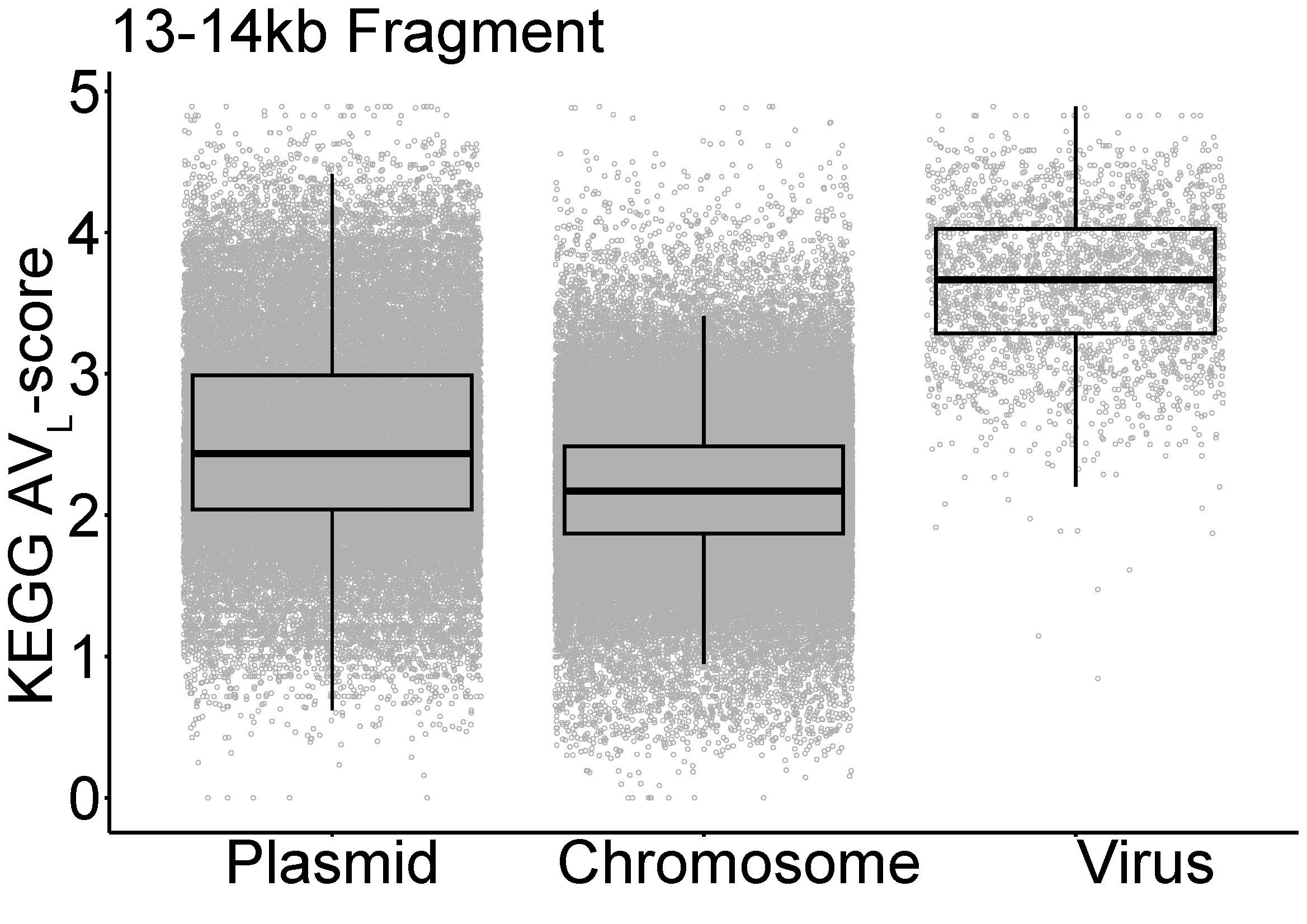

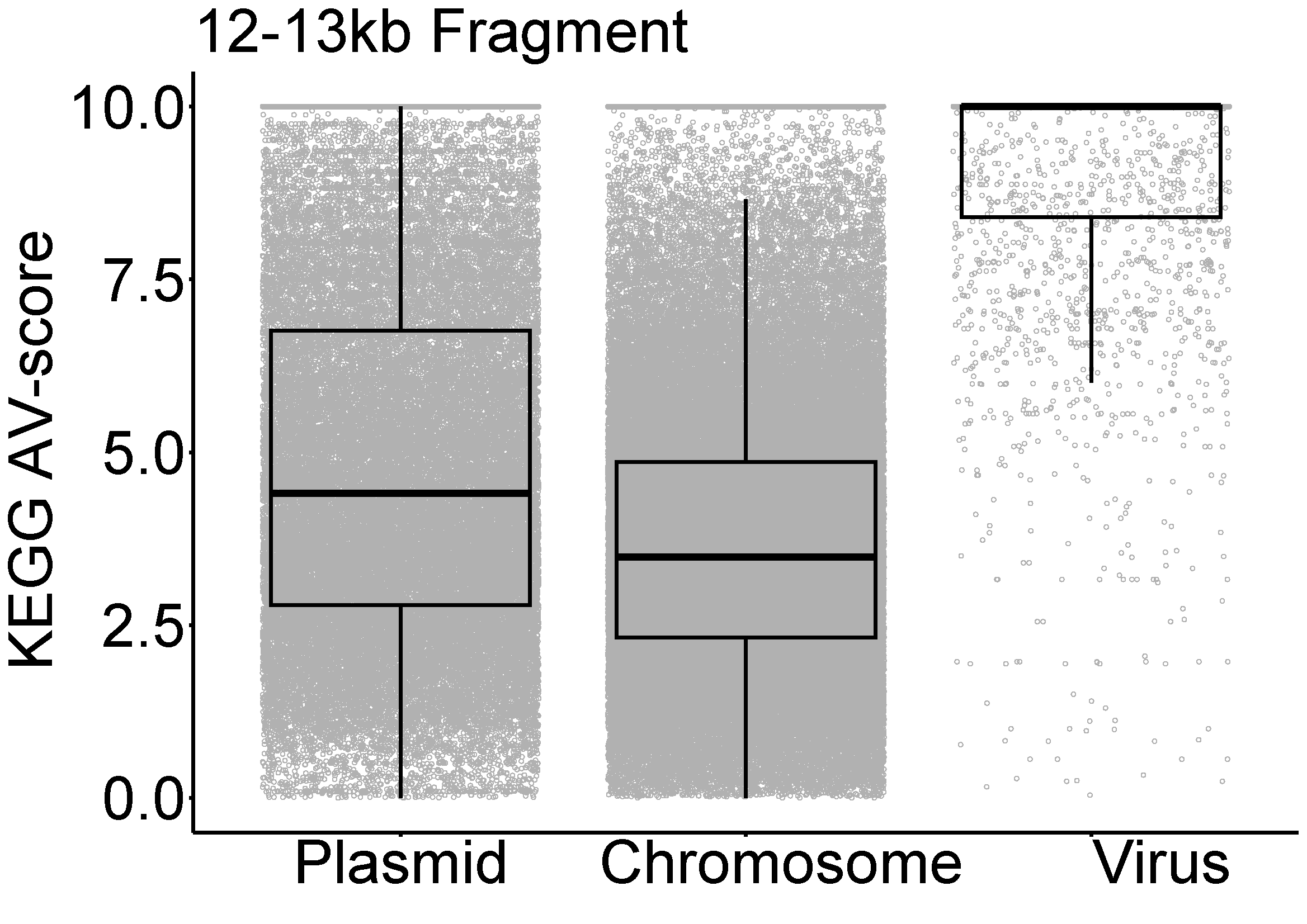

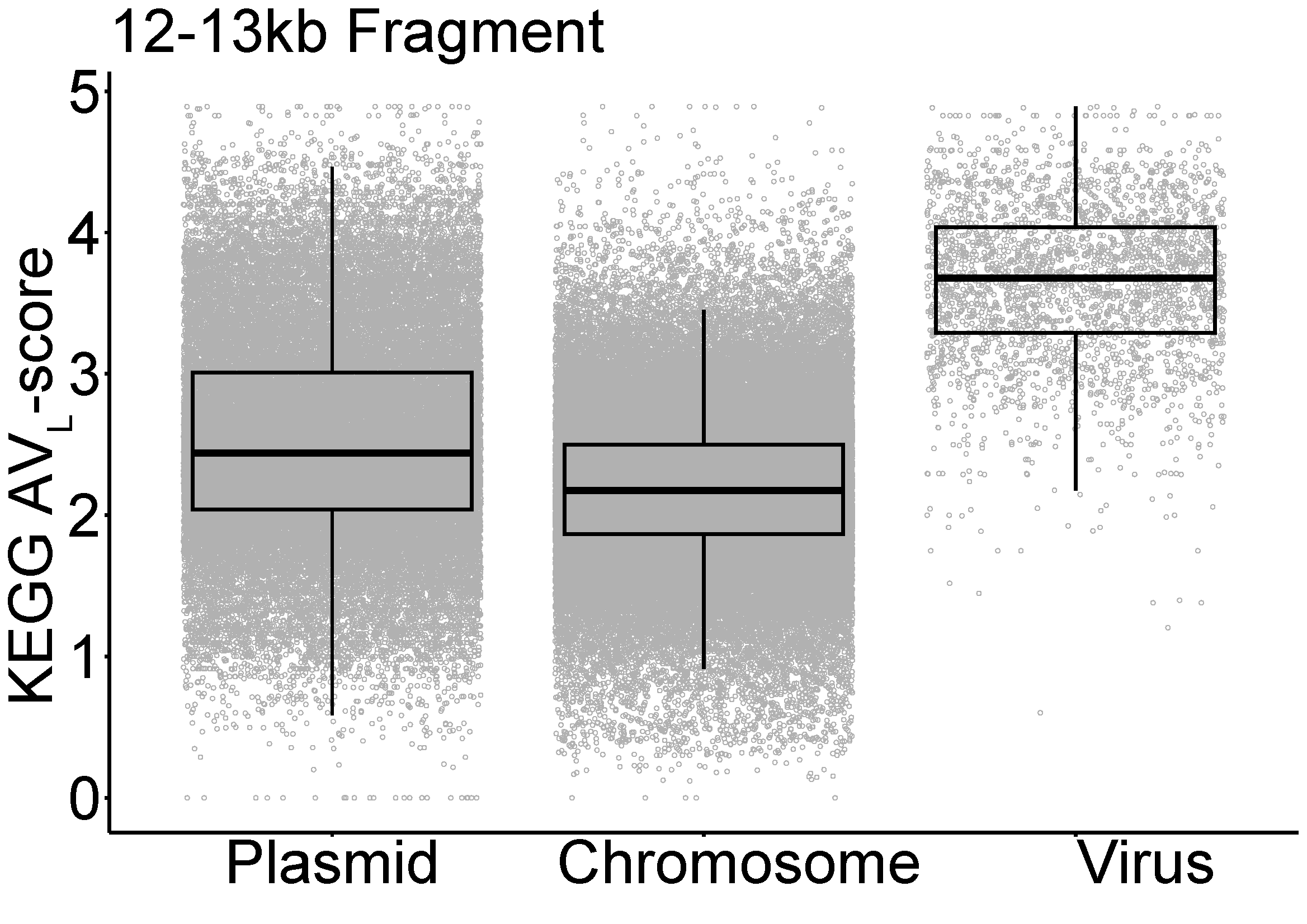

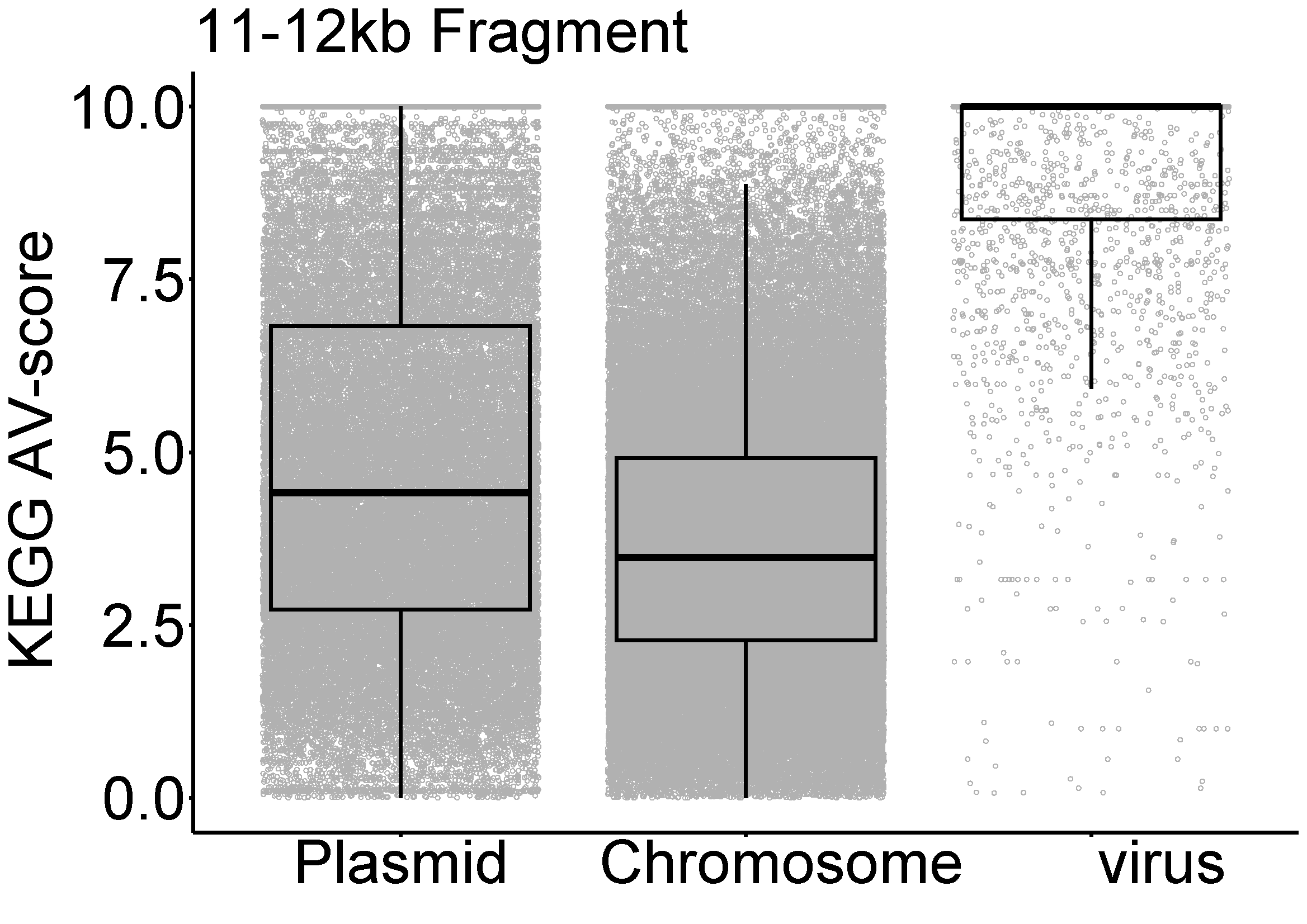

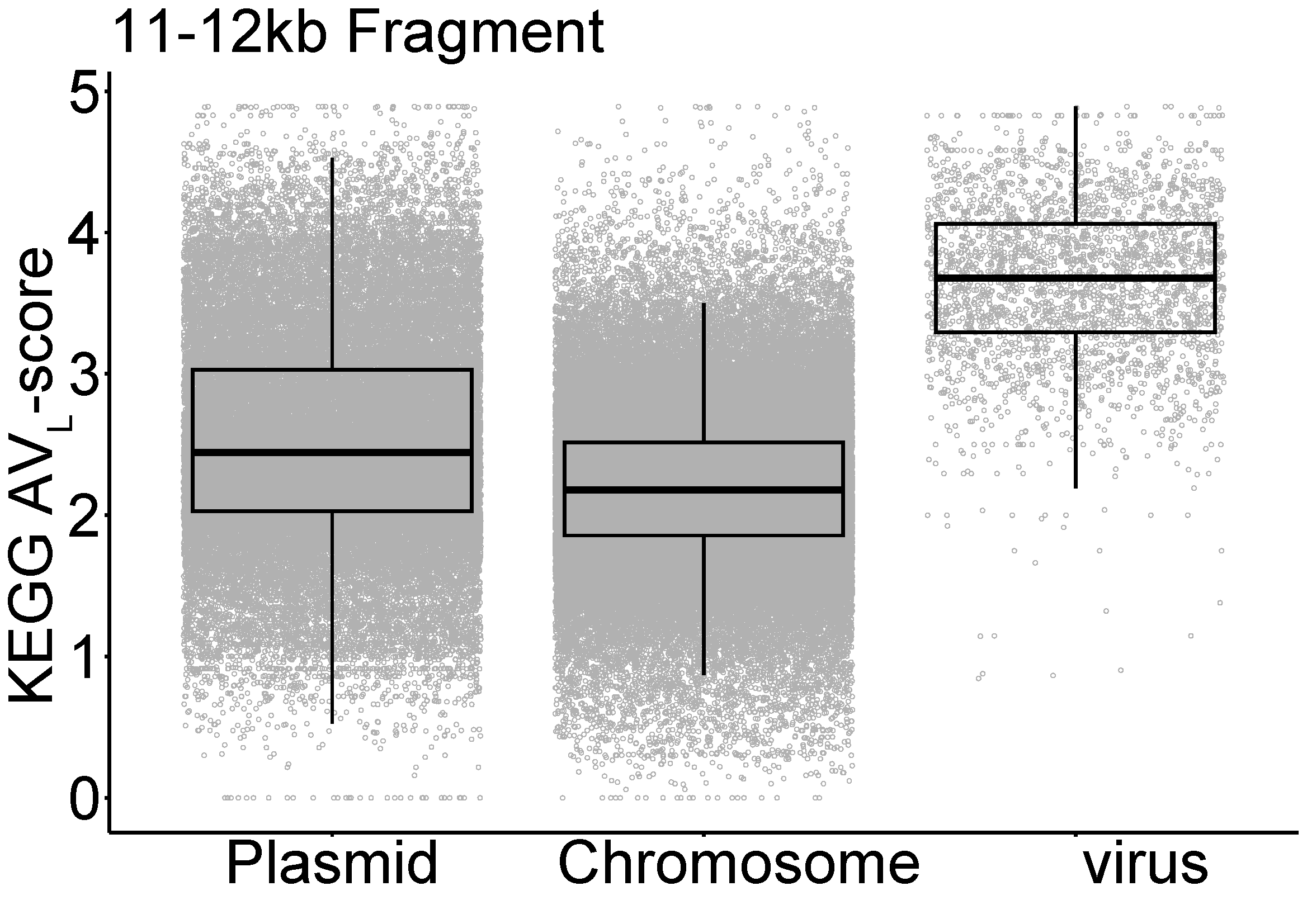

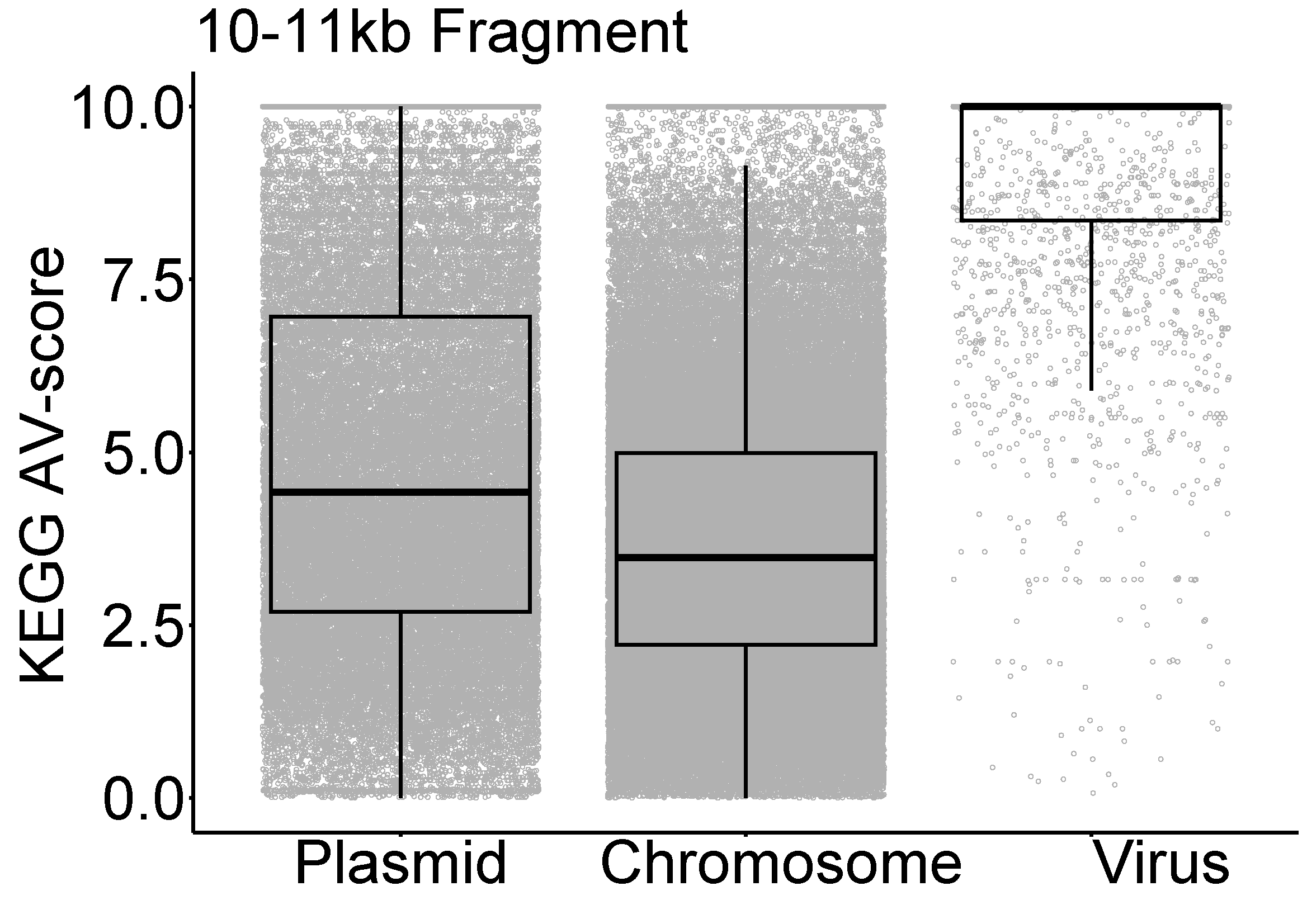

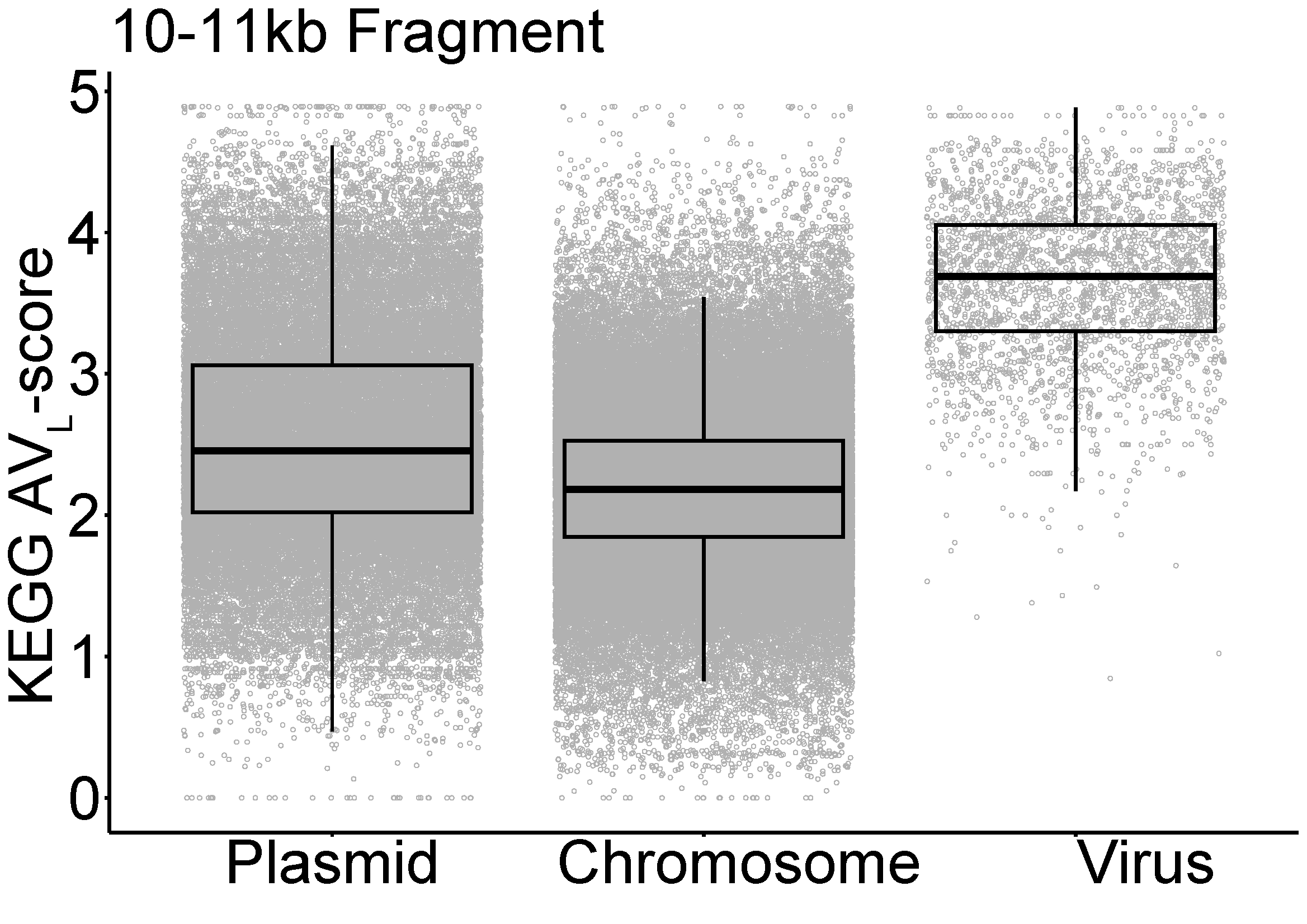

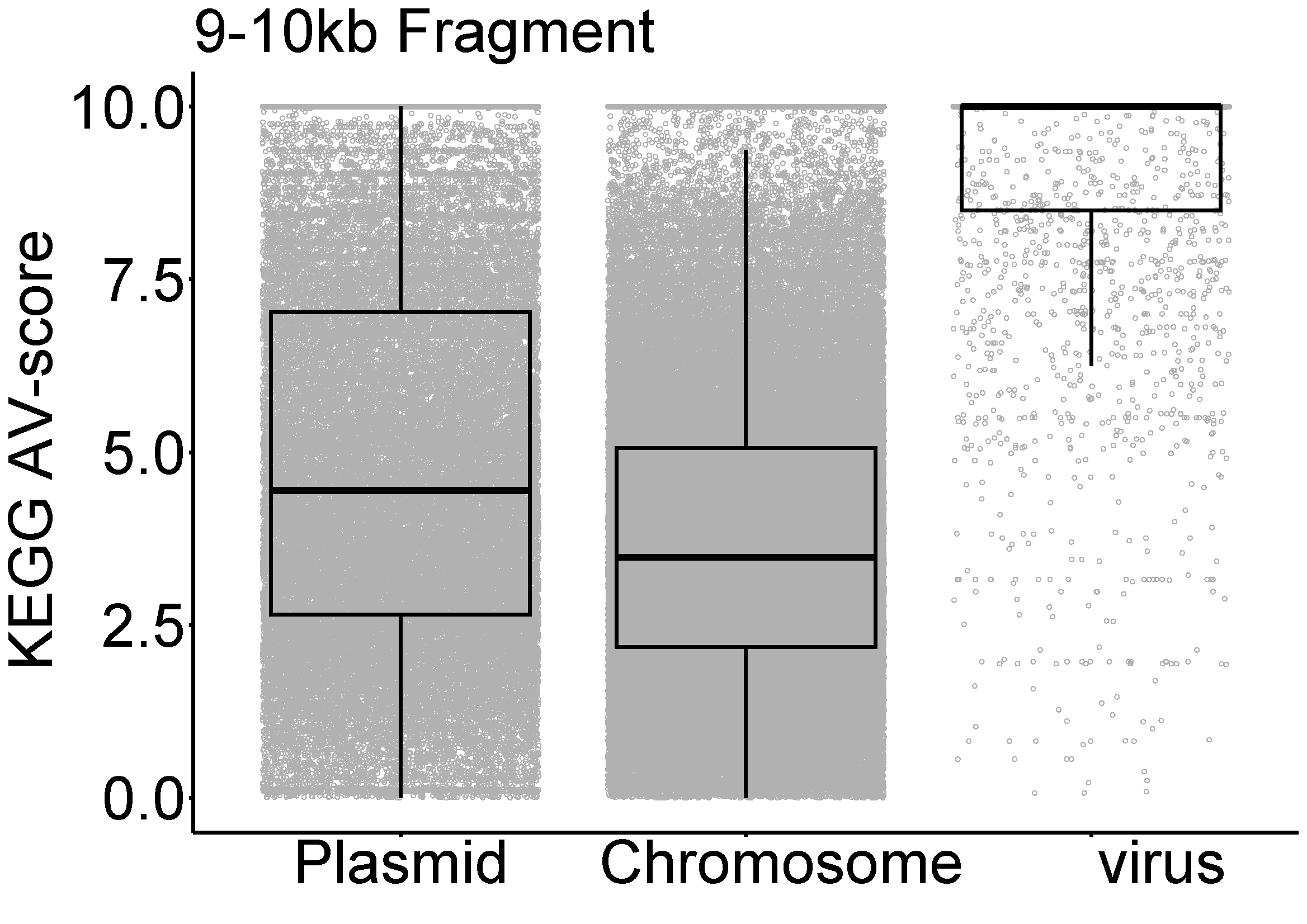

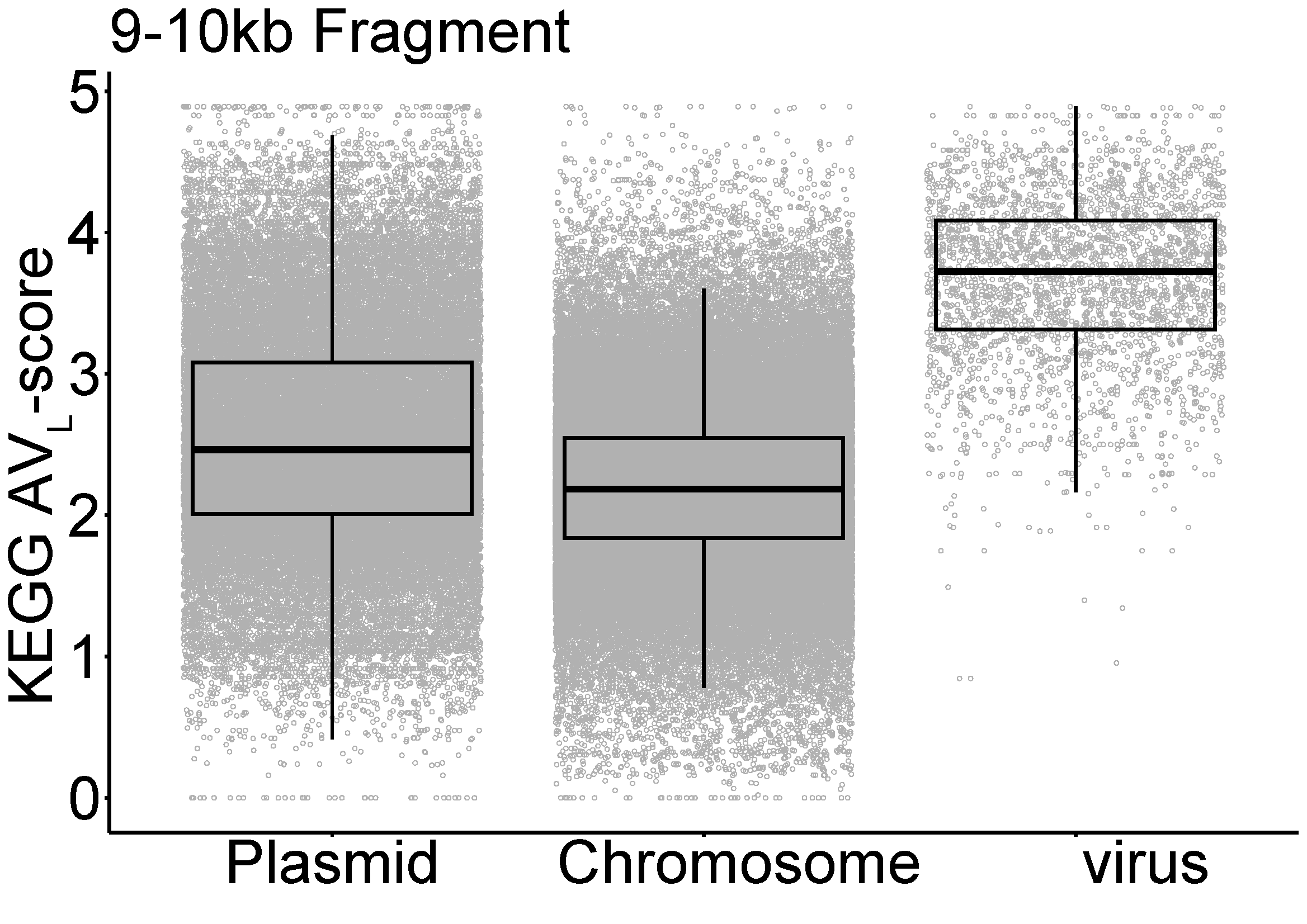

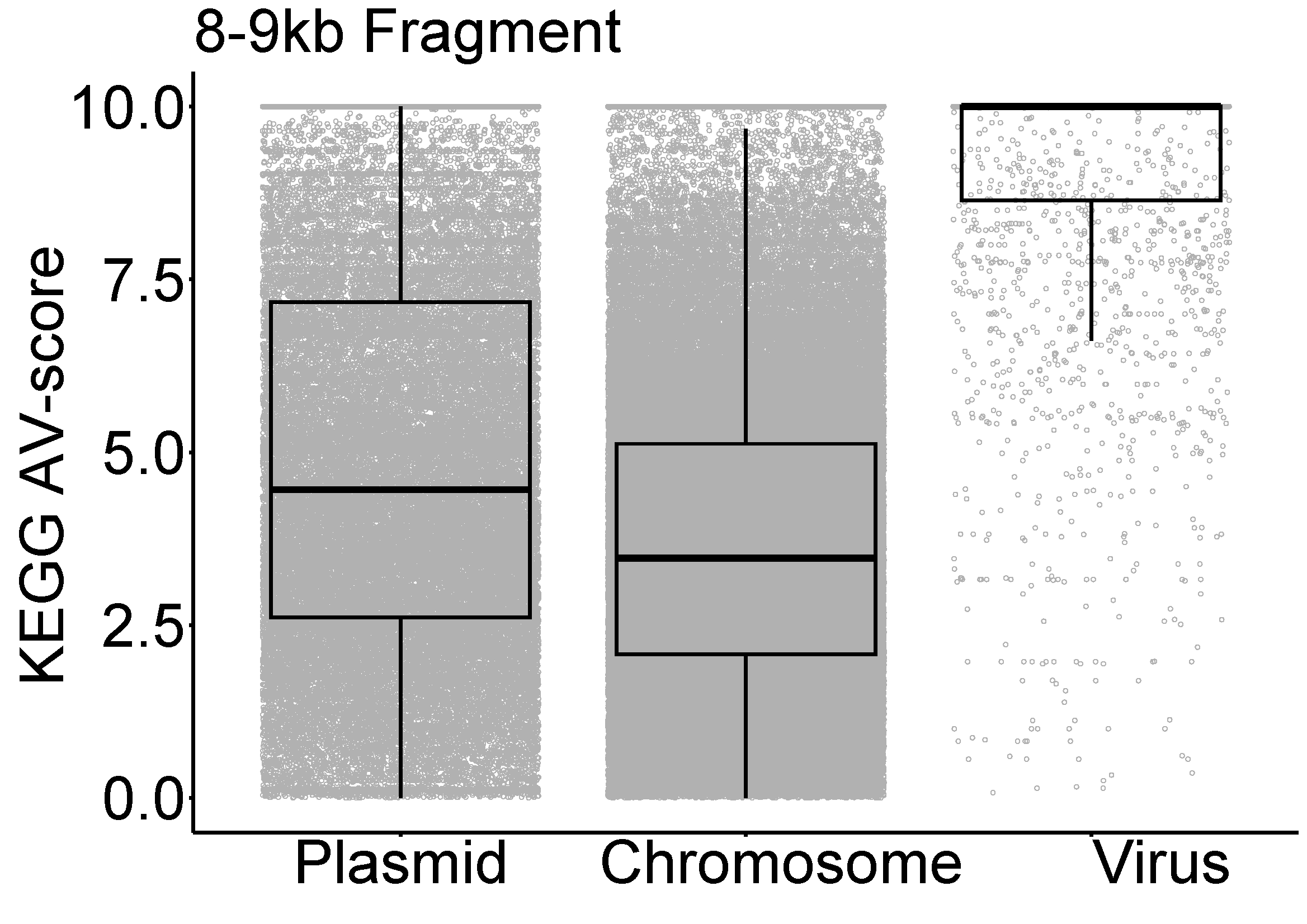

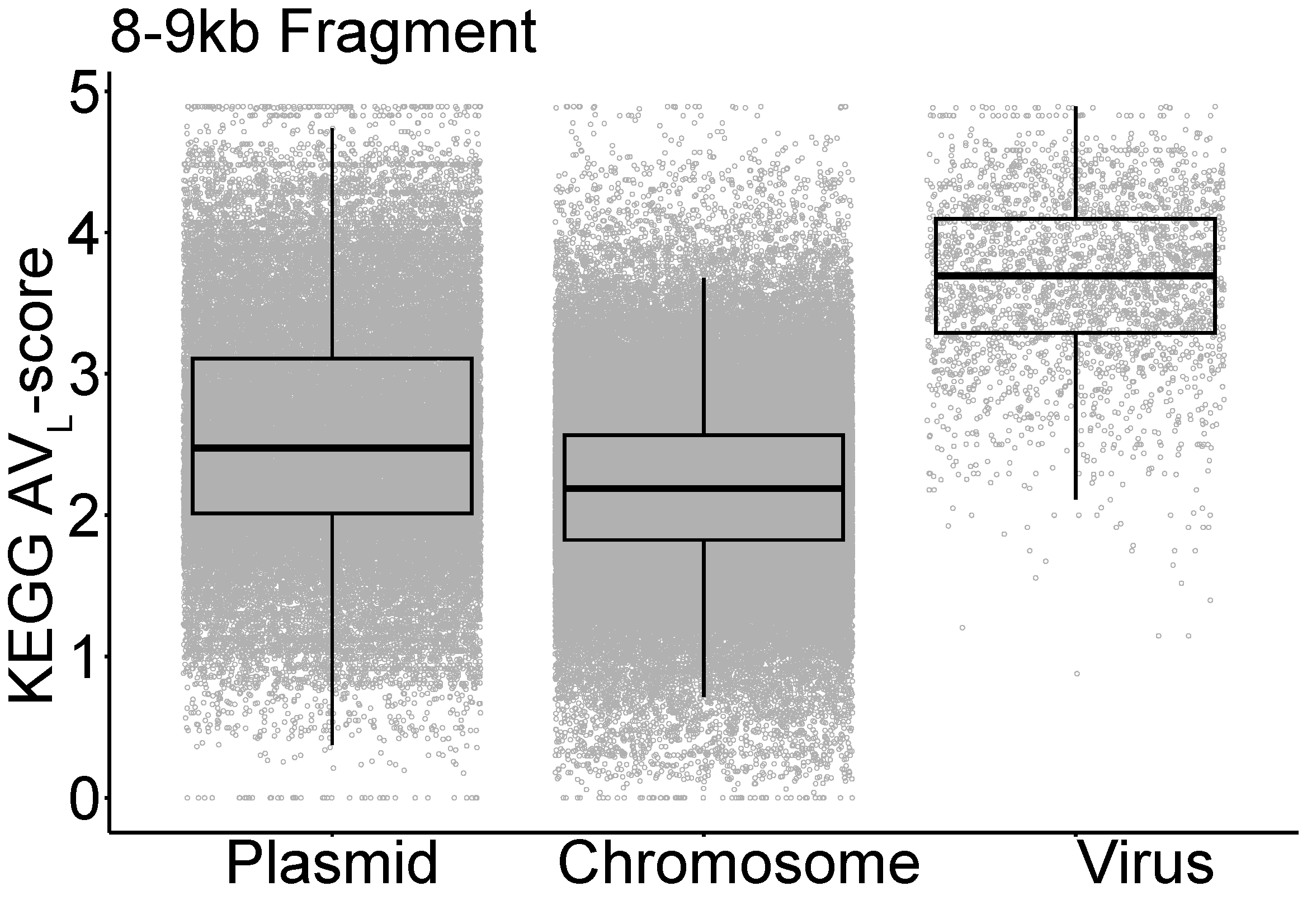

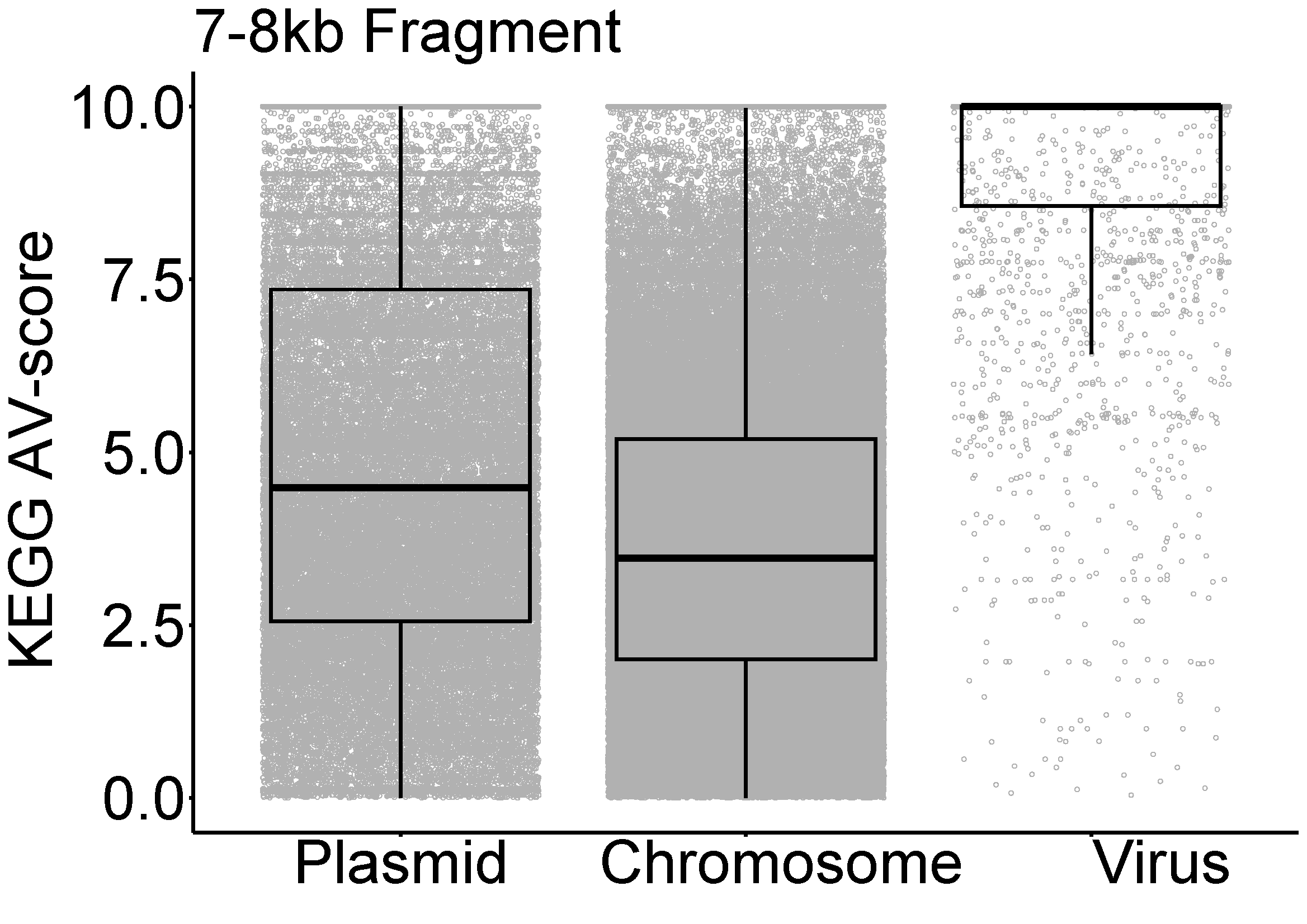

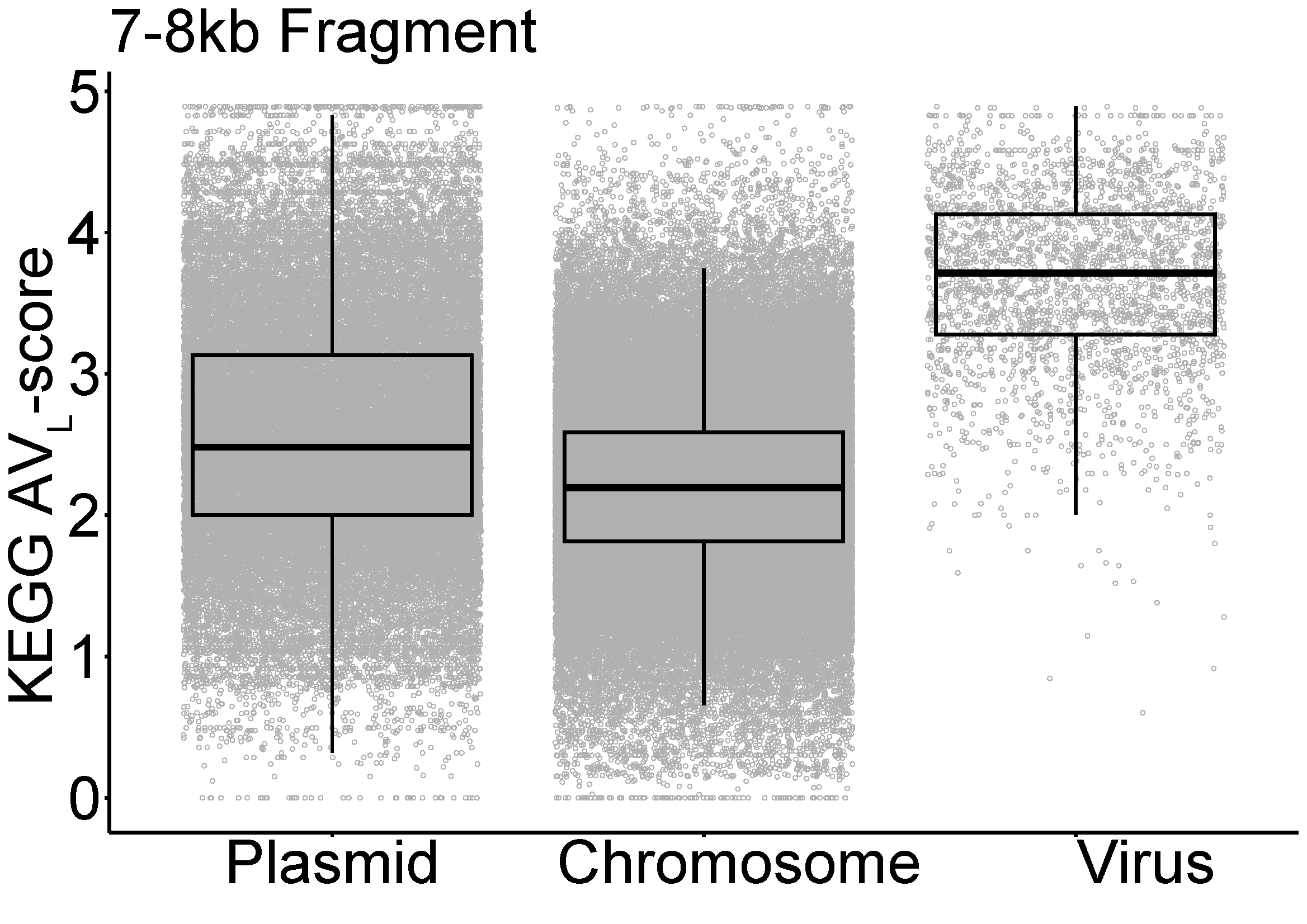

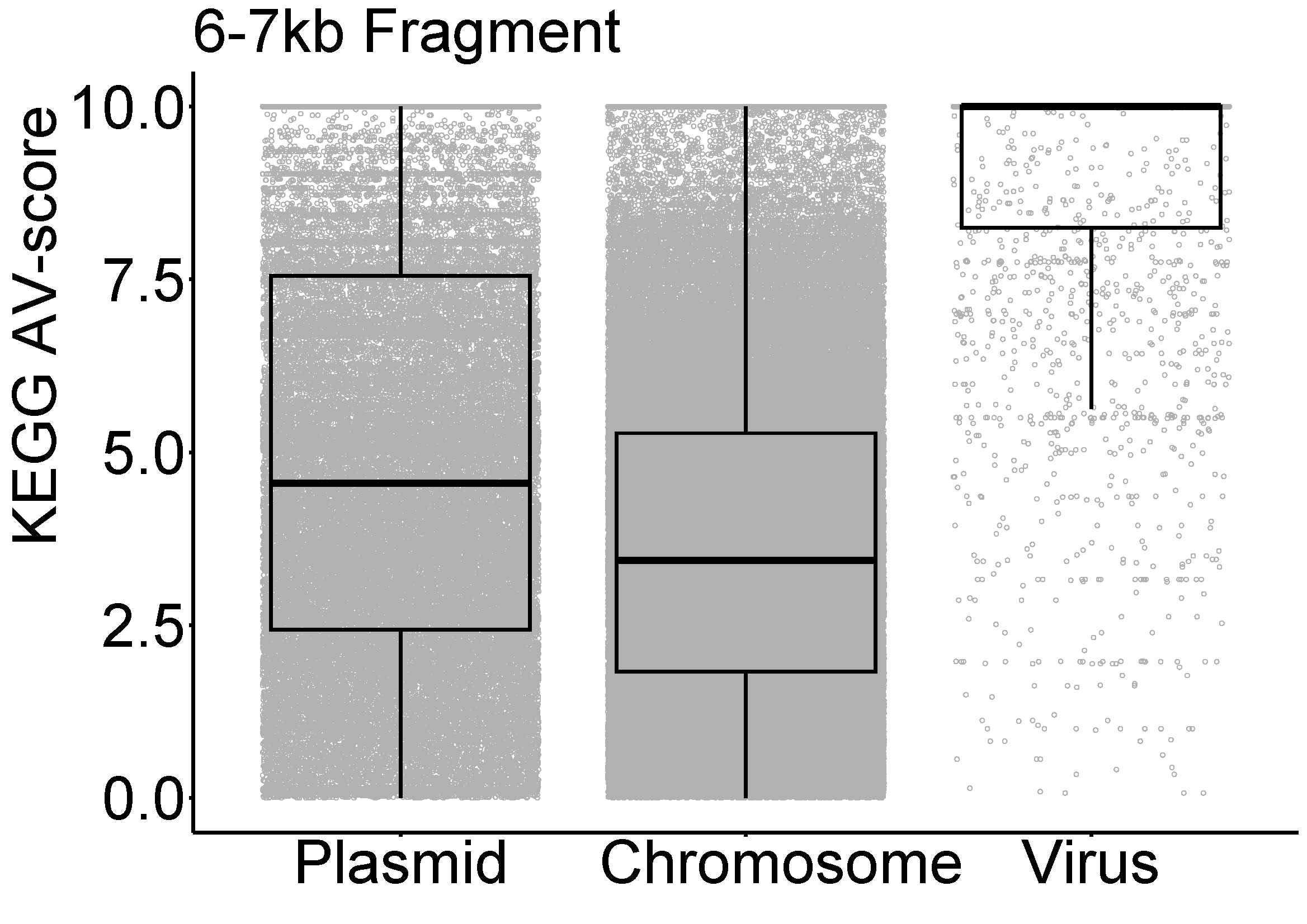

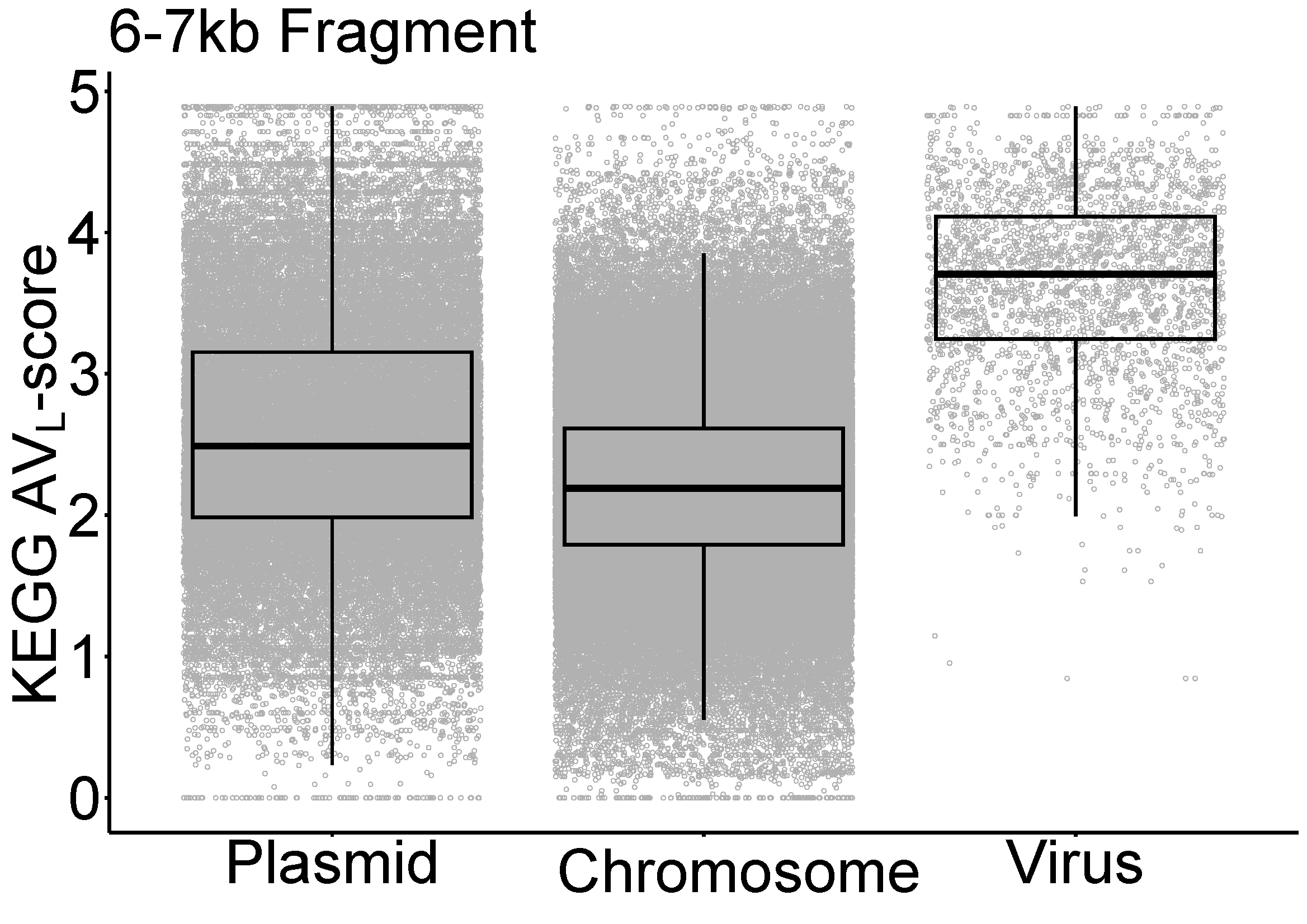

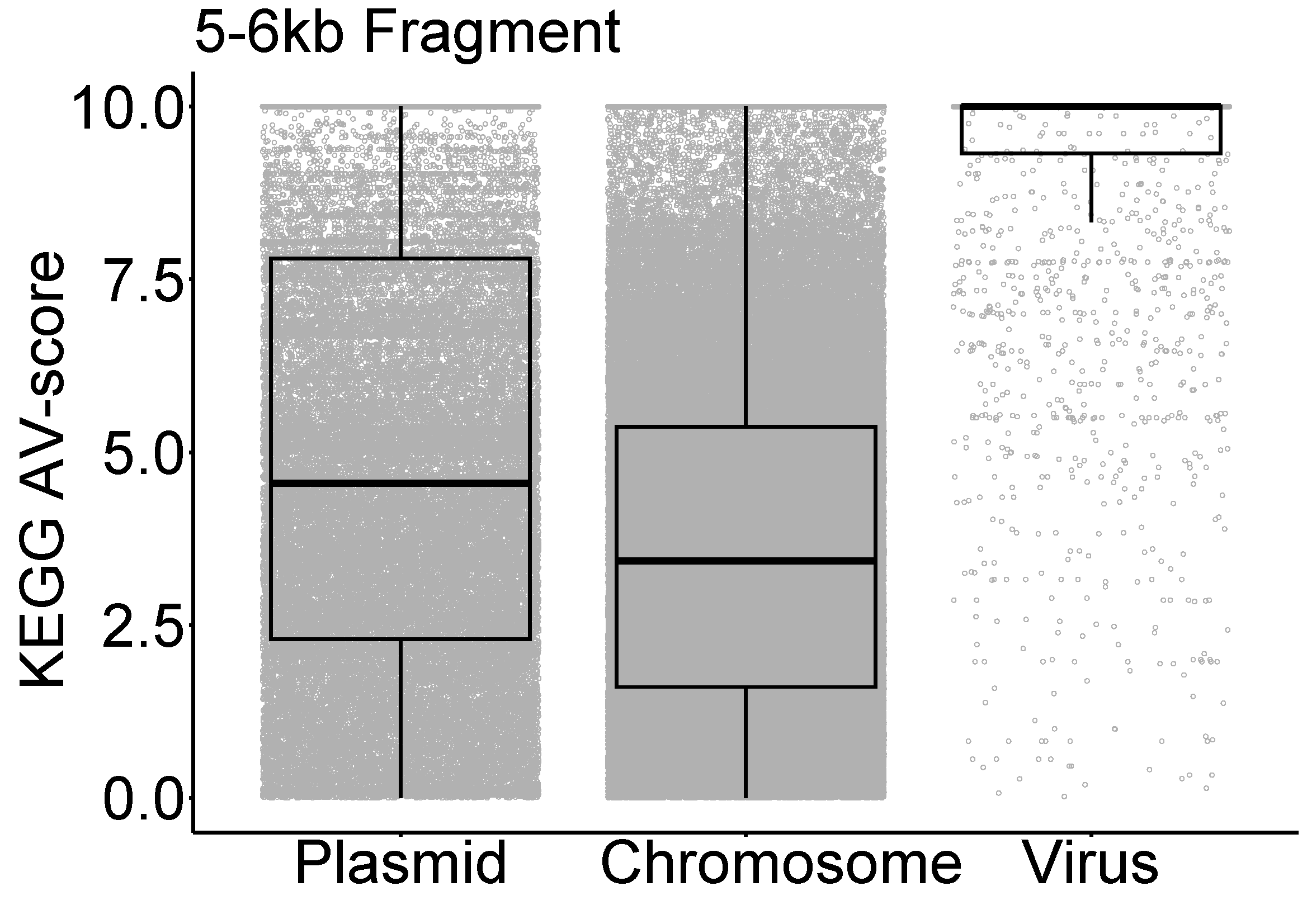

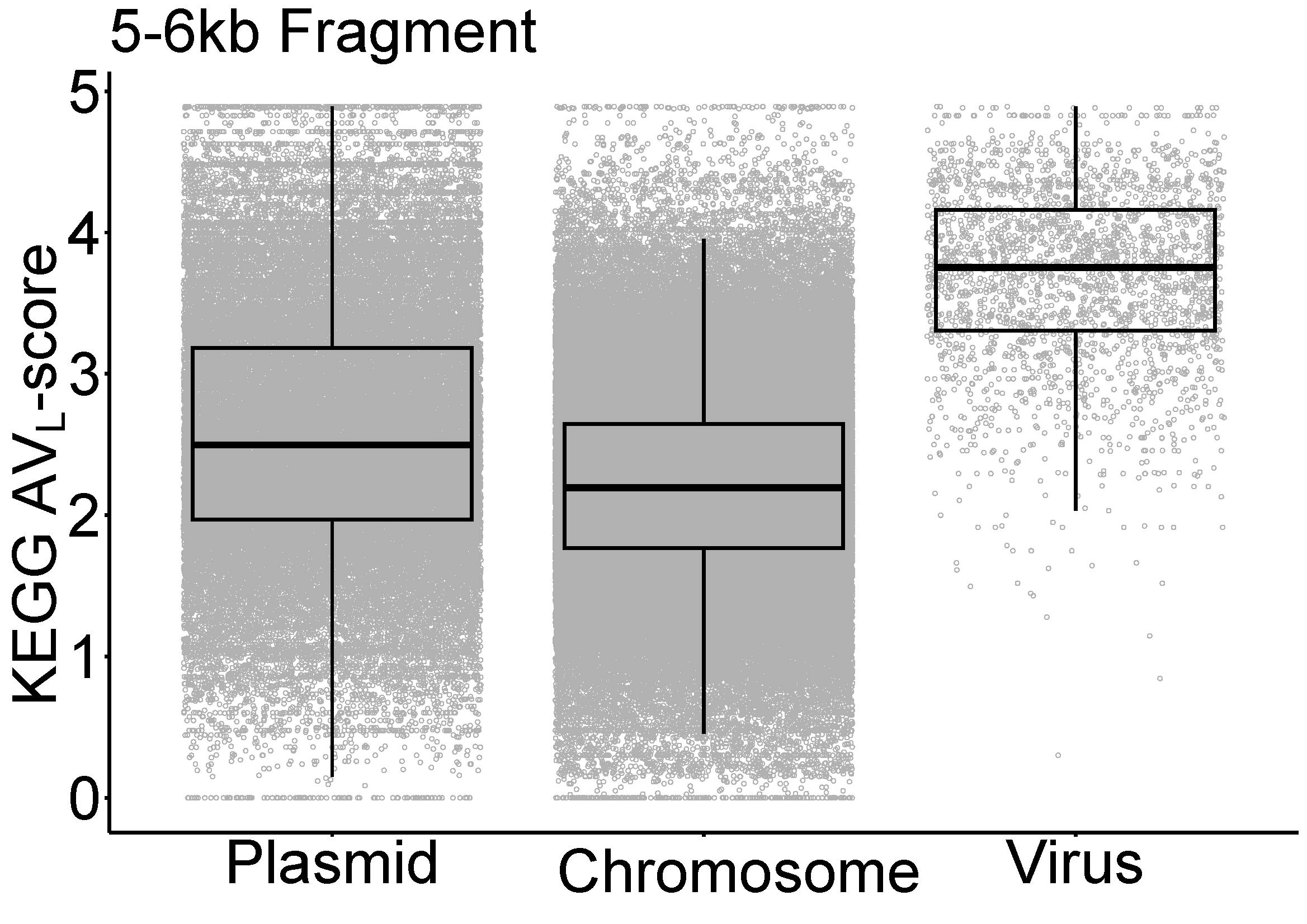

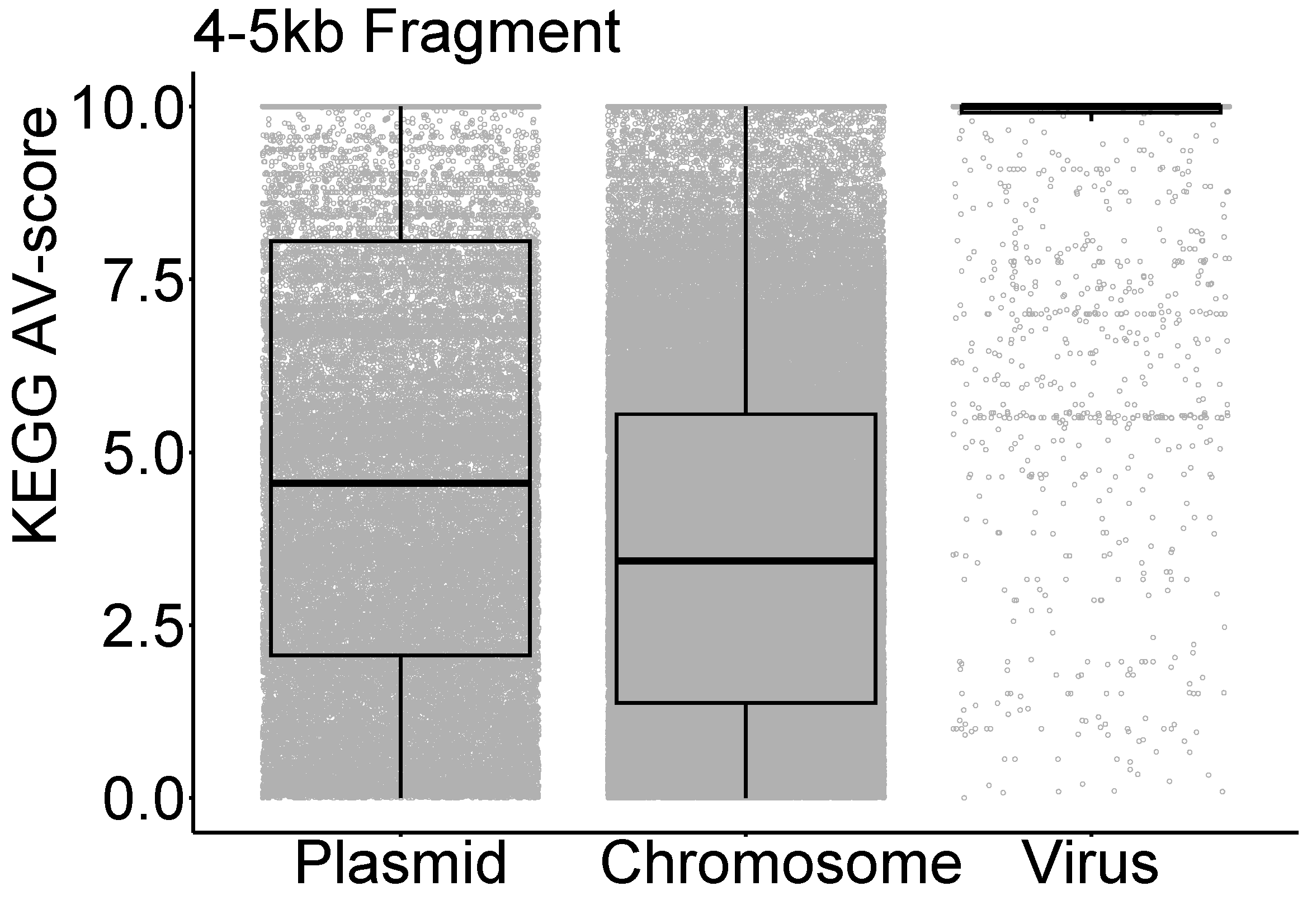

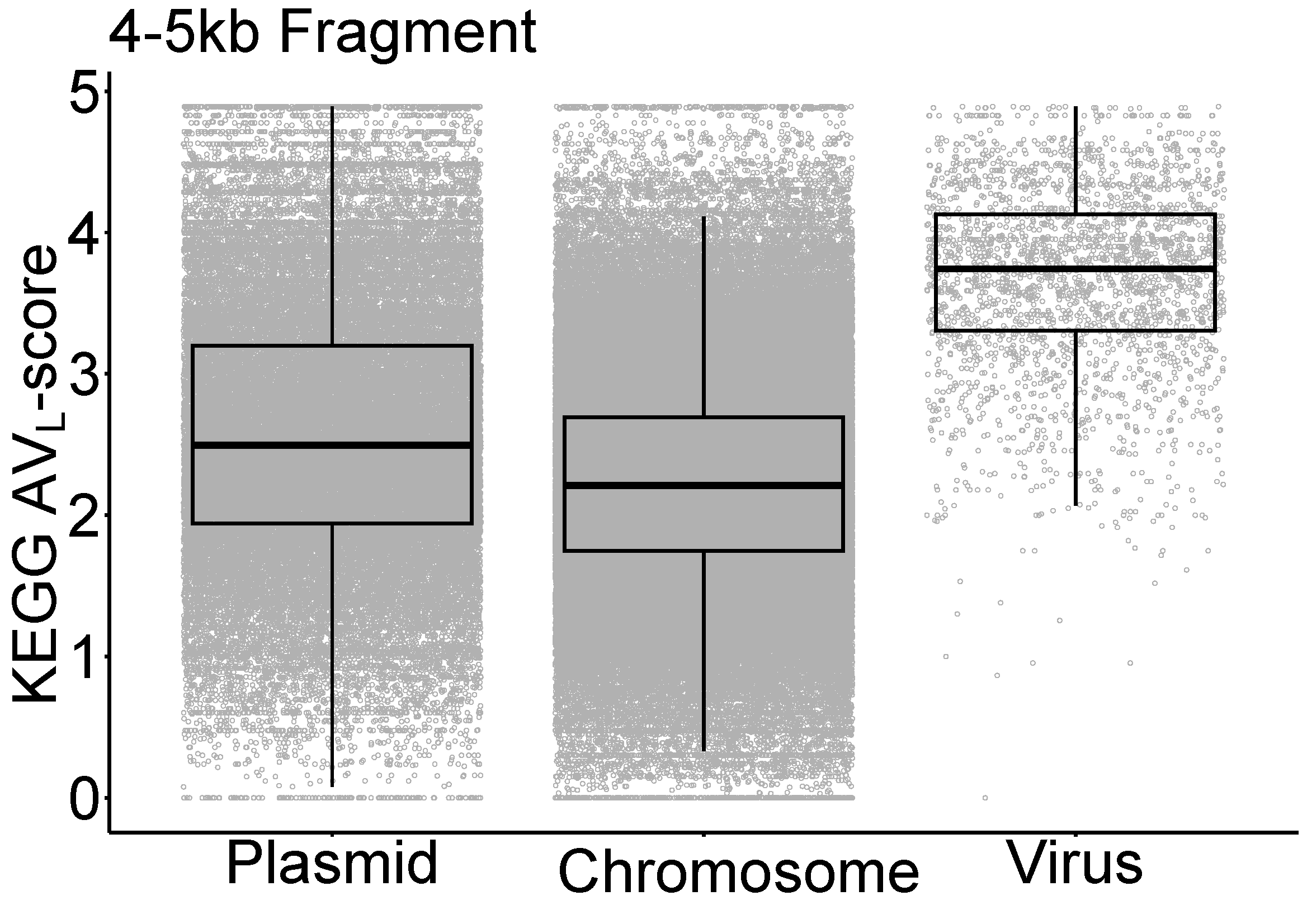

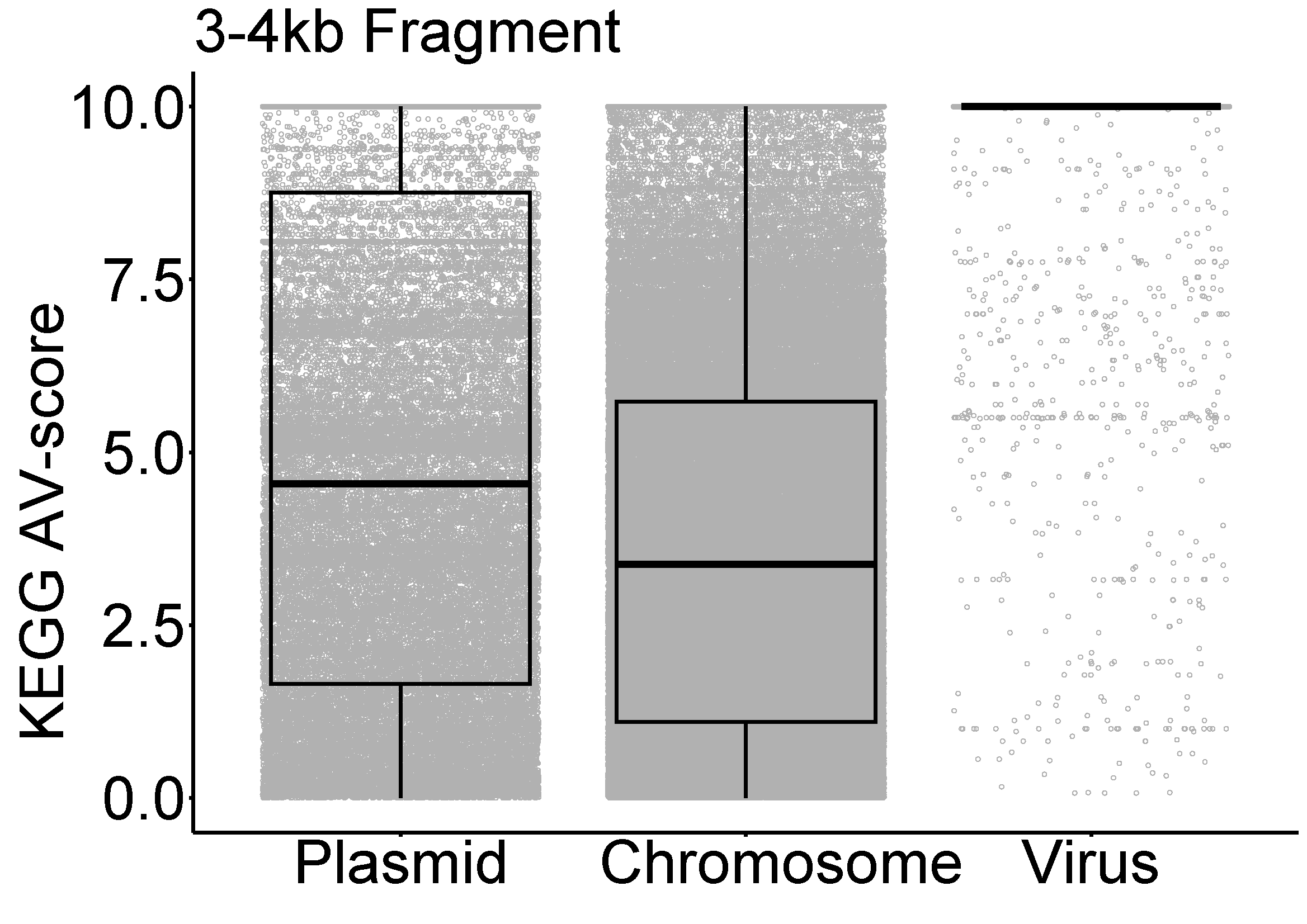

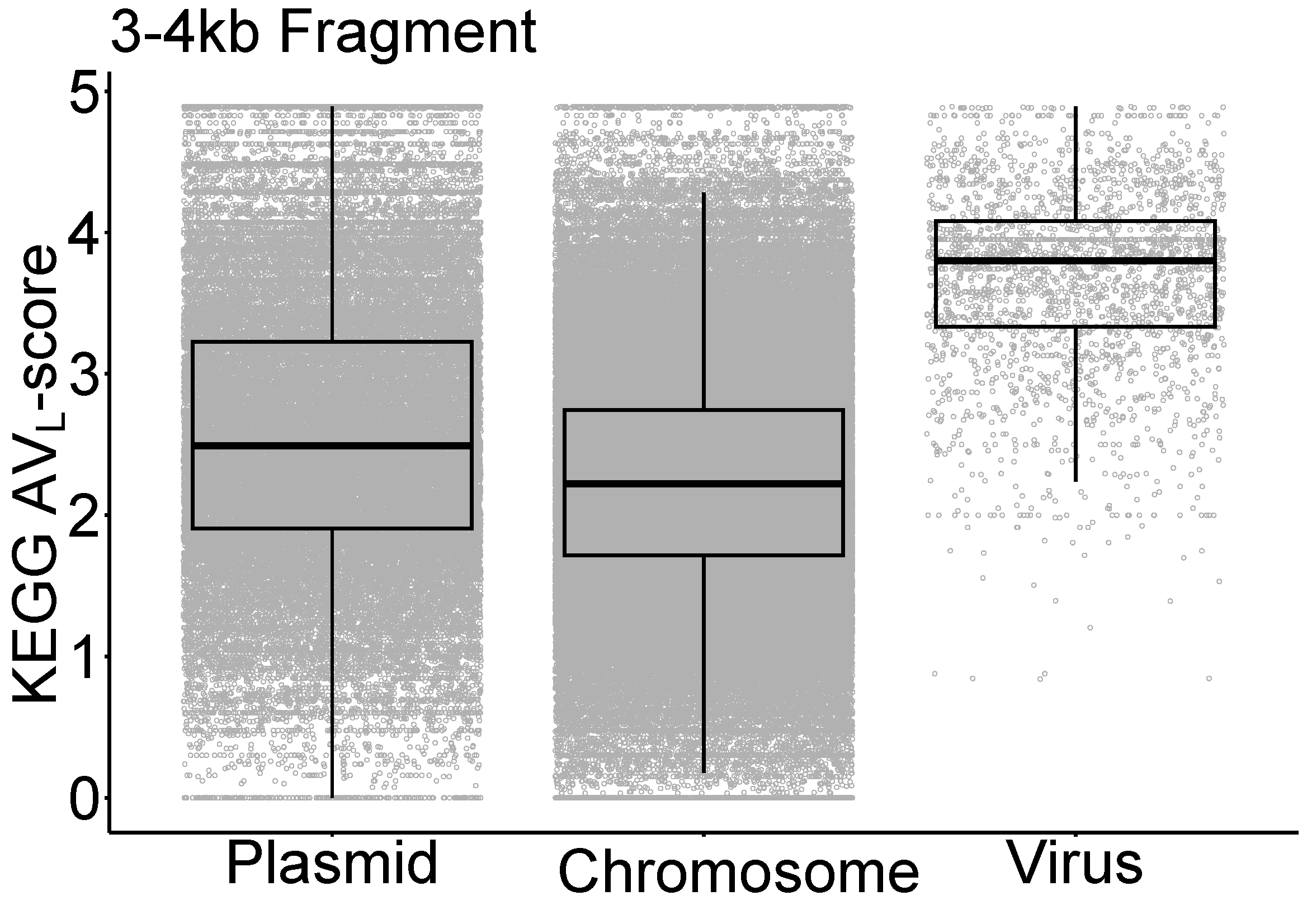

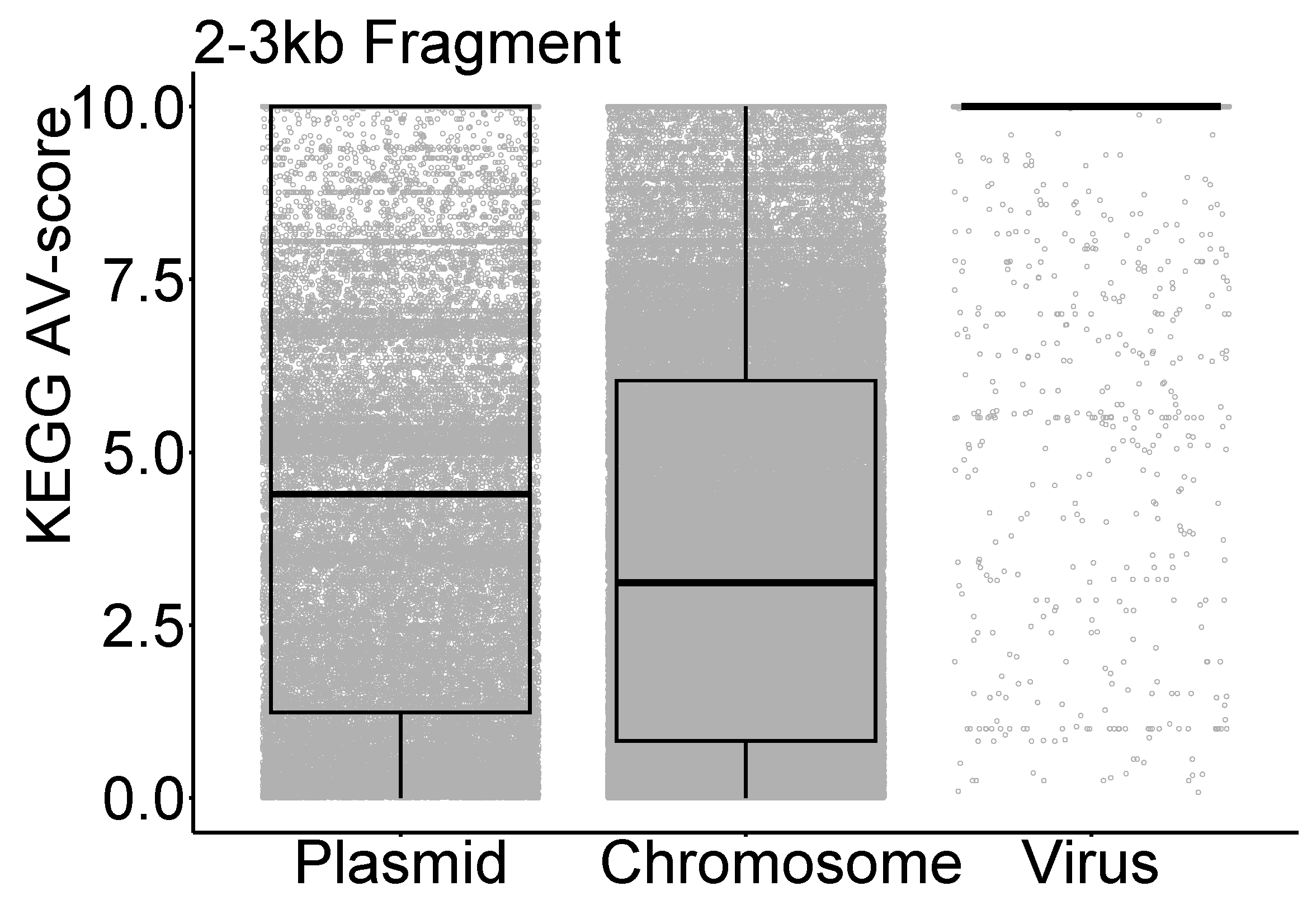

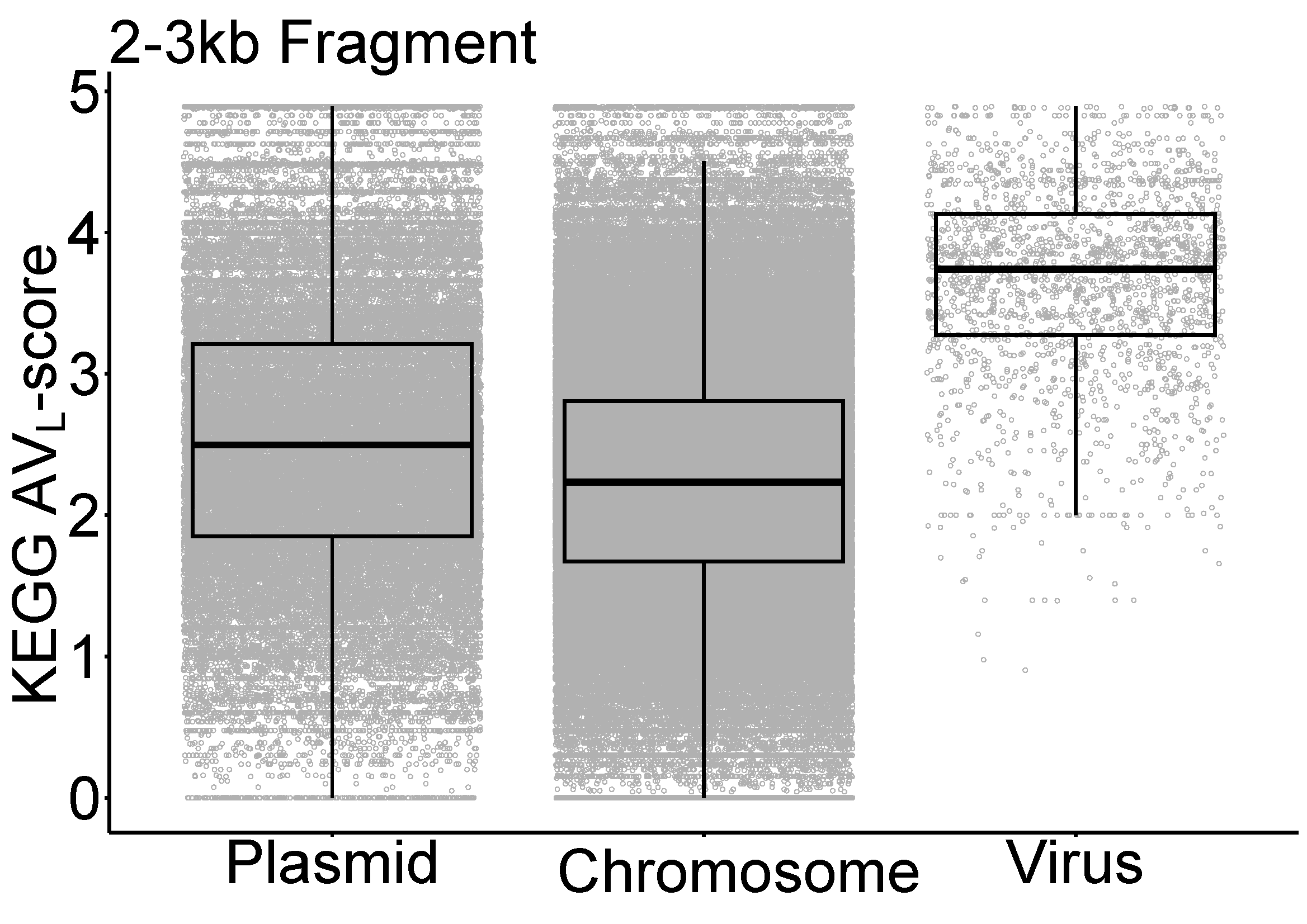

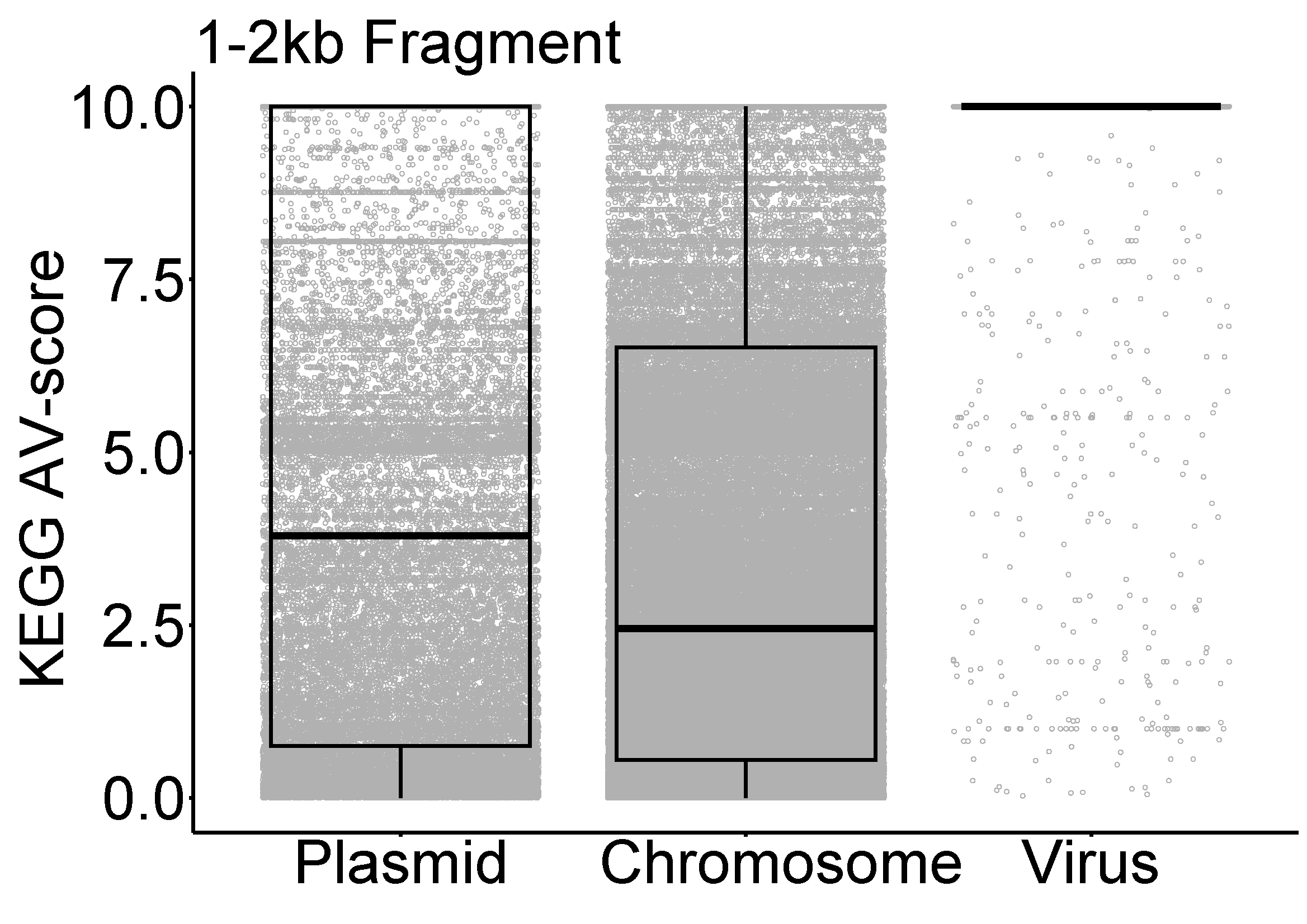

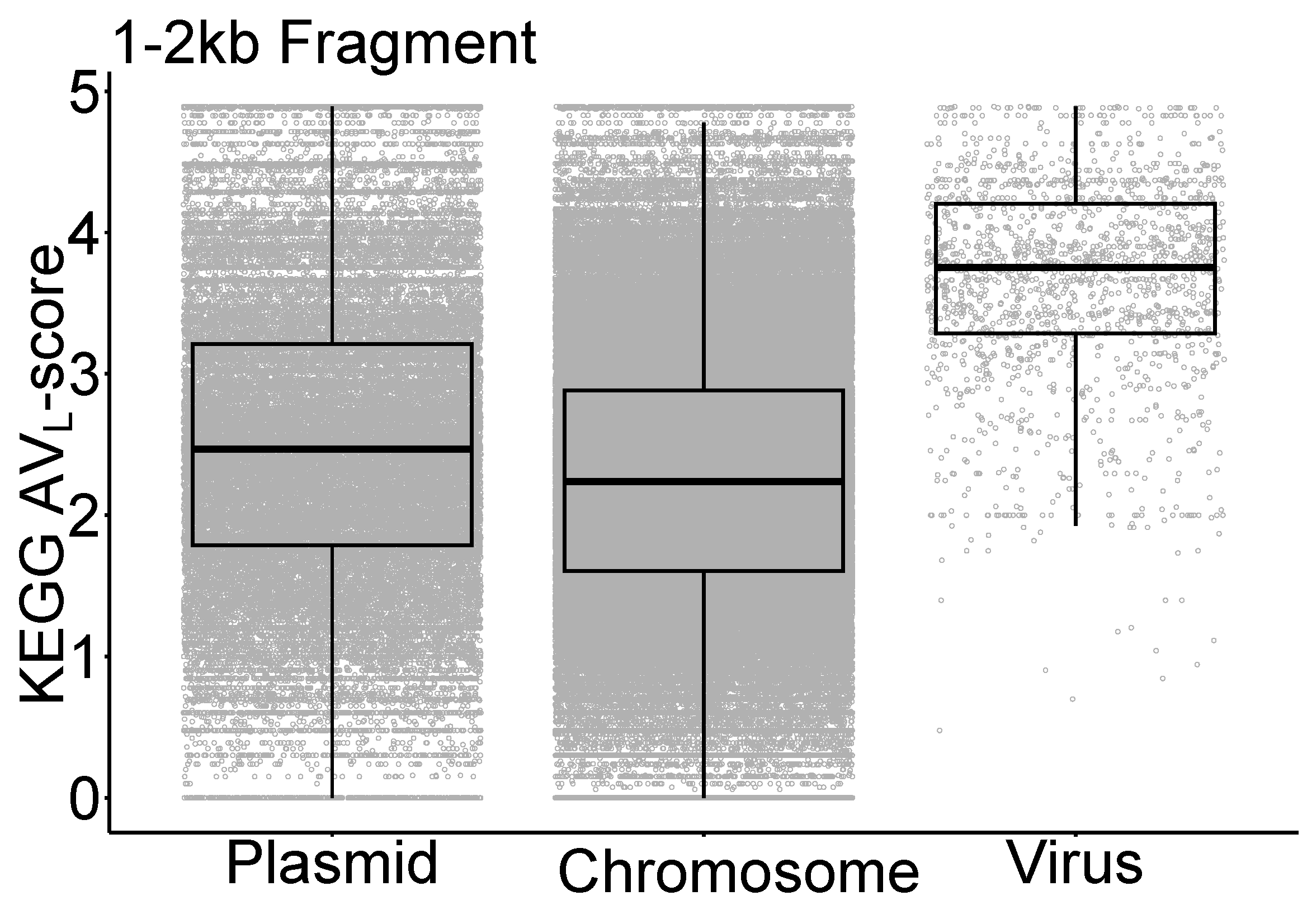

**Fig. S2.** Distribution of KEGG AV-score and AV_L_-score of split prokaryotic chromosome fragments (*n* = 1975,048), genome fragments of plasmids (*n* = 681,614) and prokaryotic viruses (*n* = 48,880). The horizontal line that splits the box is the median, the upper and lower sides of the box are upper and lower quartiles, whiskers are 1.5 times the interquartile ranges and data points beyond whiskers are considered potential outliers.

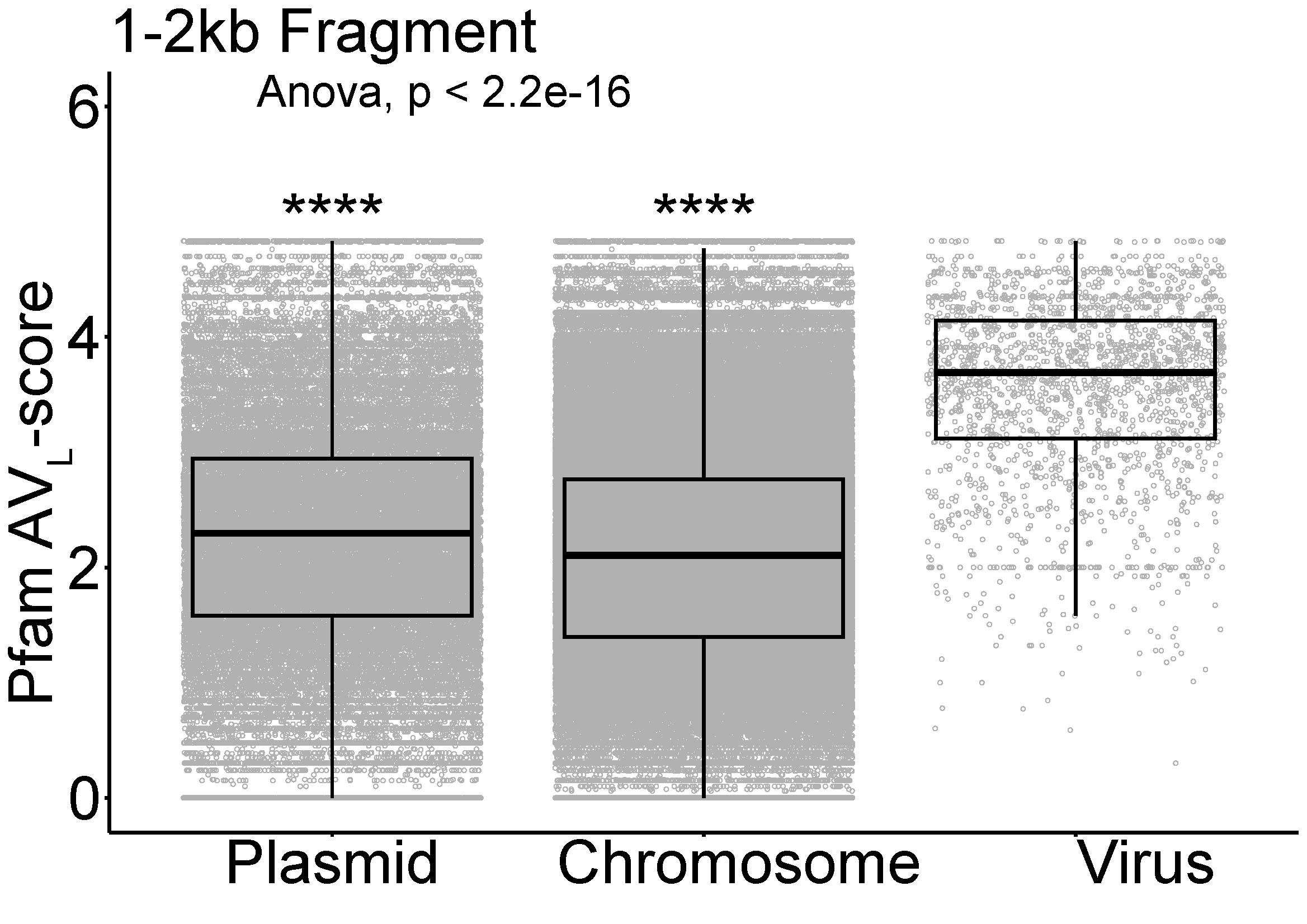

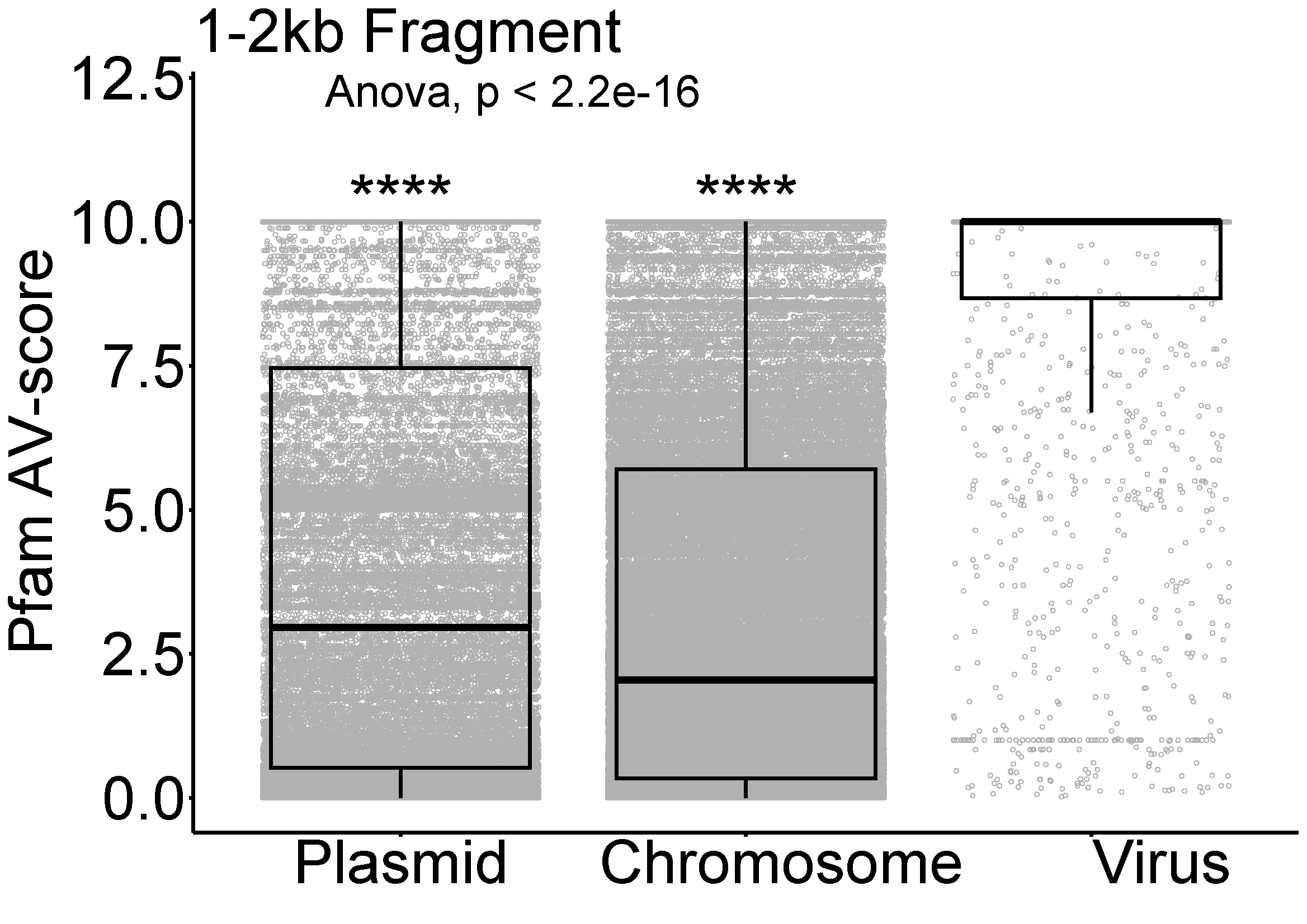

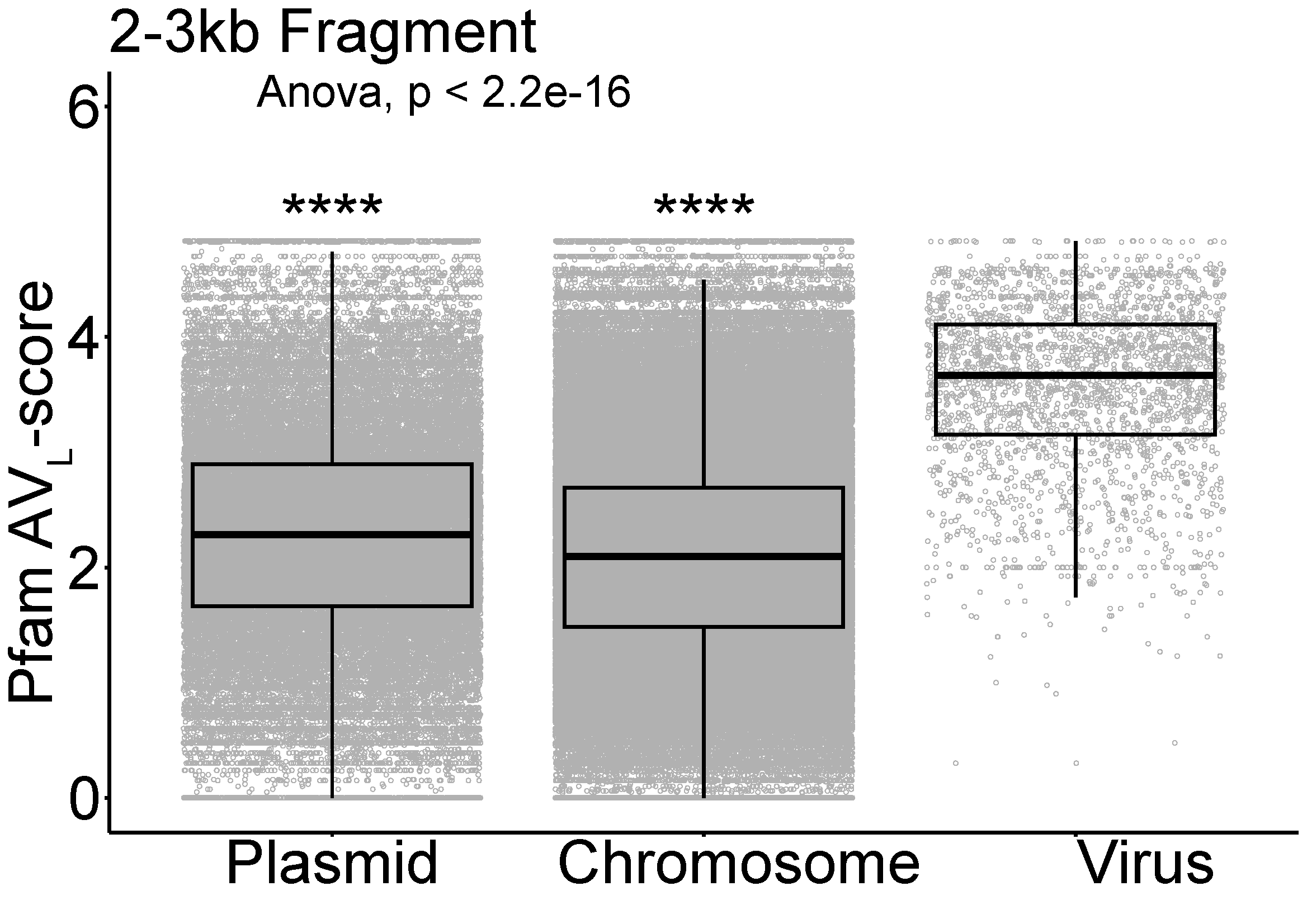

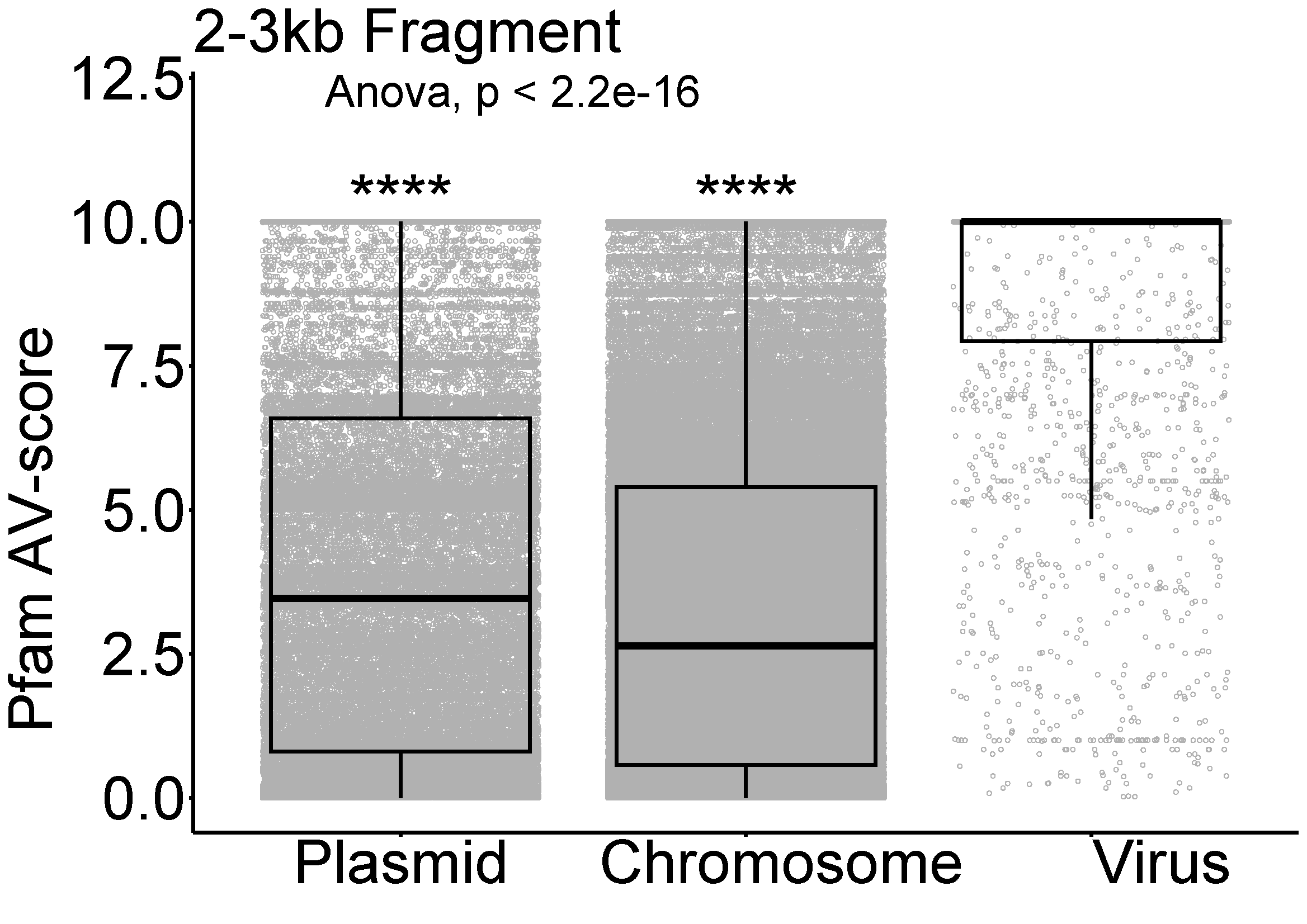

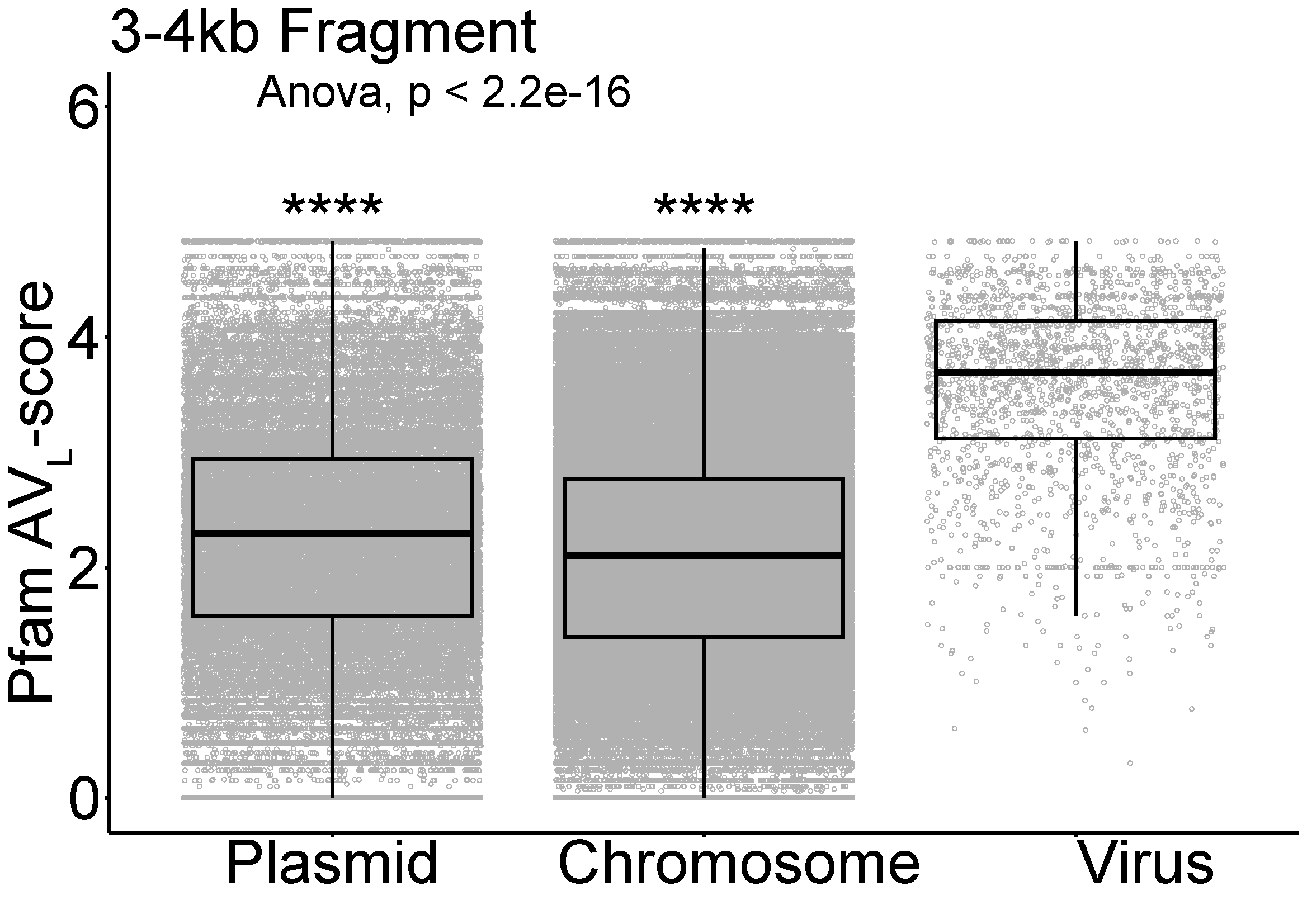

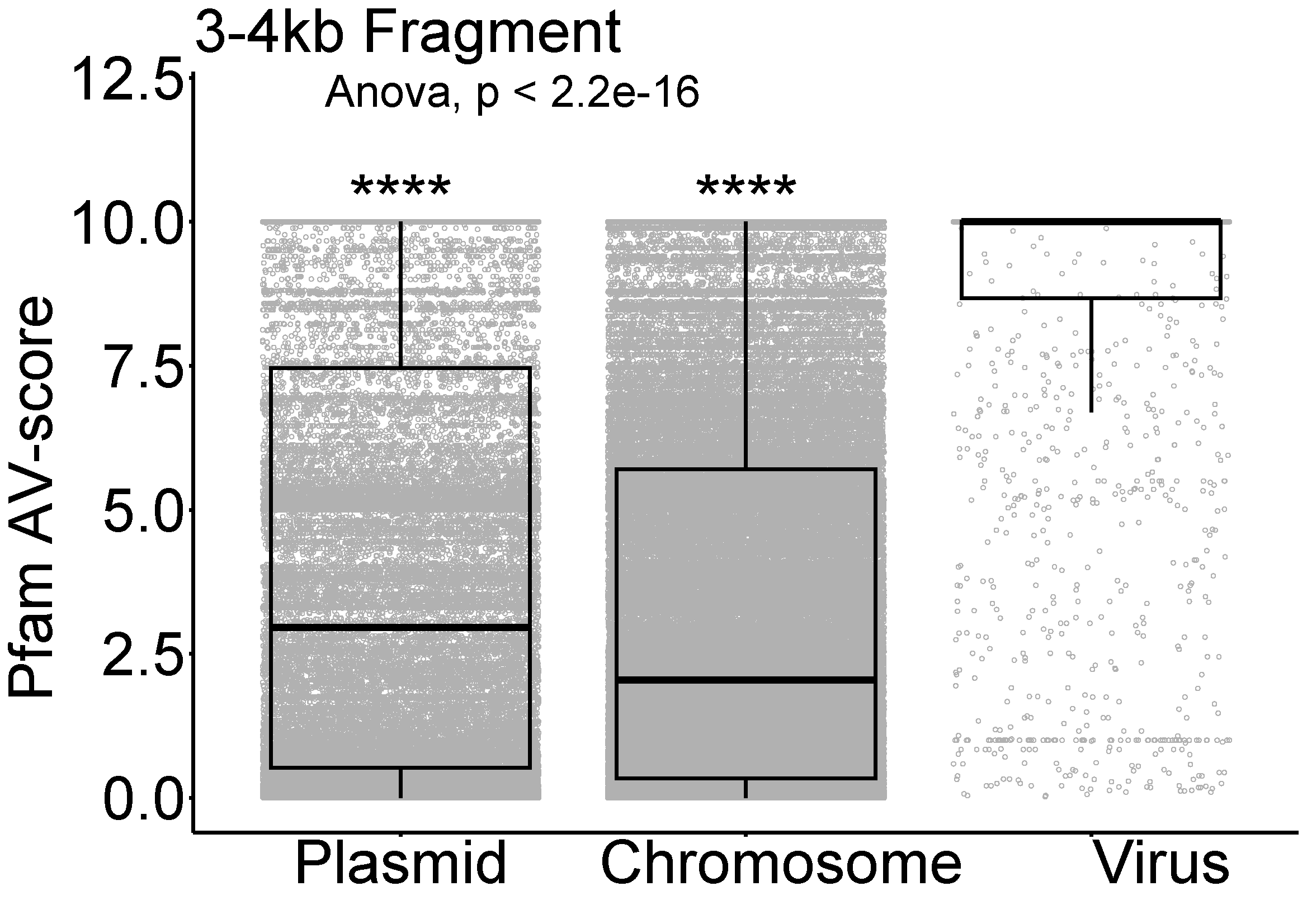

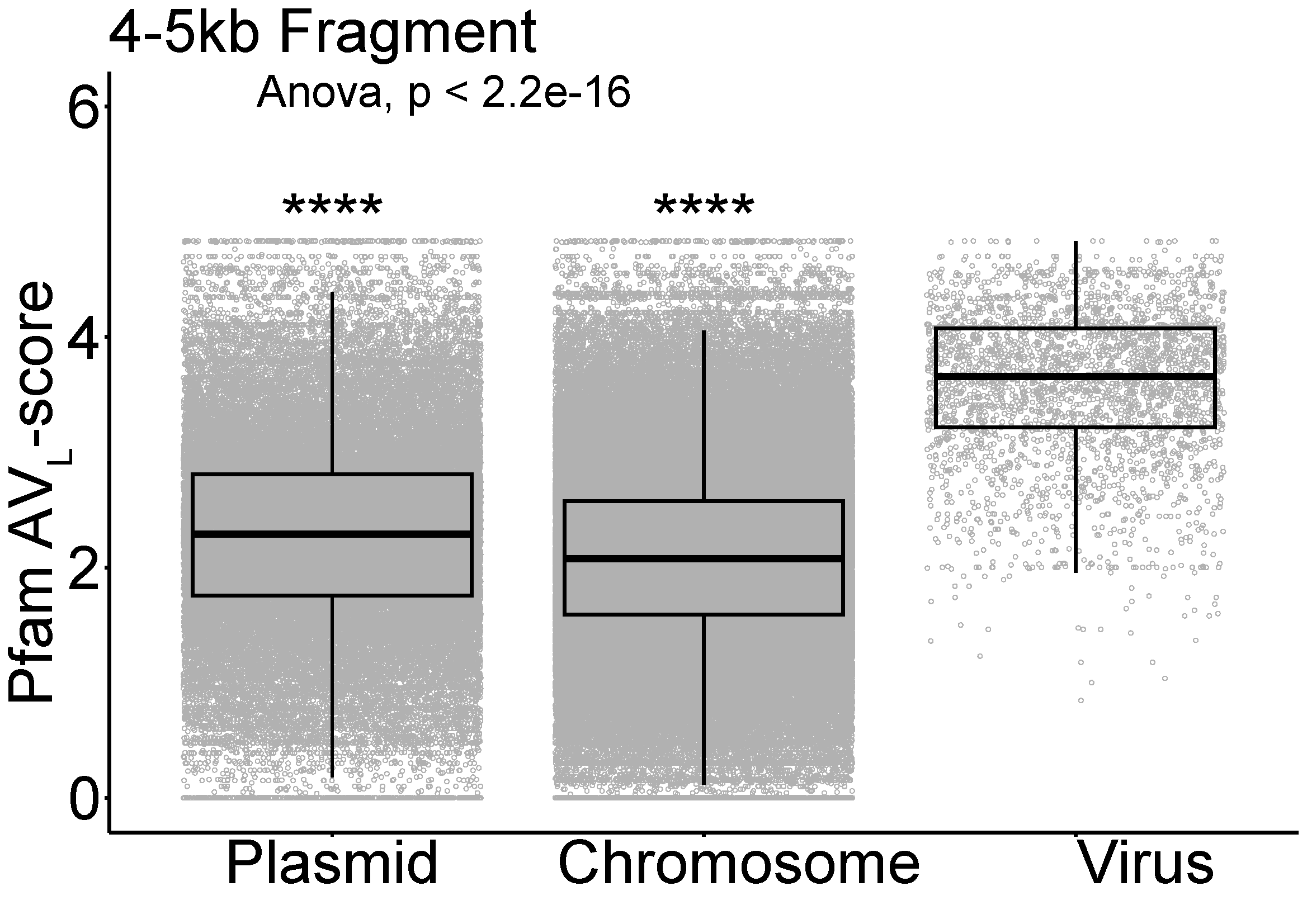

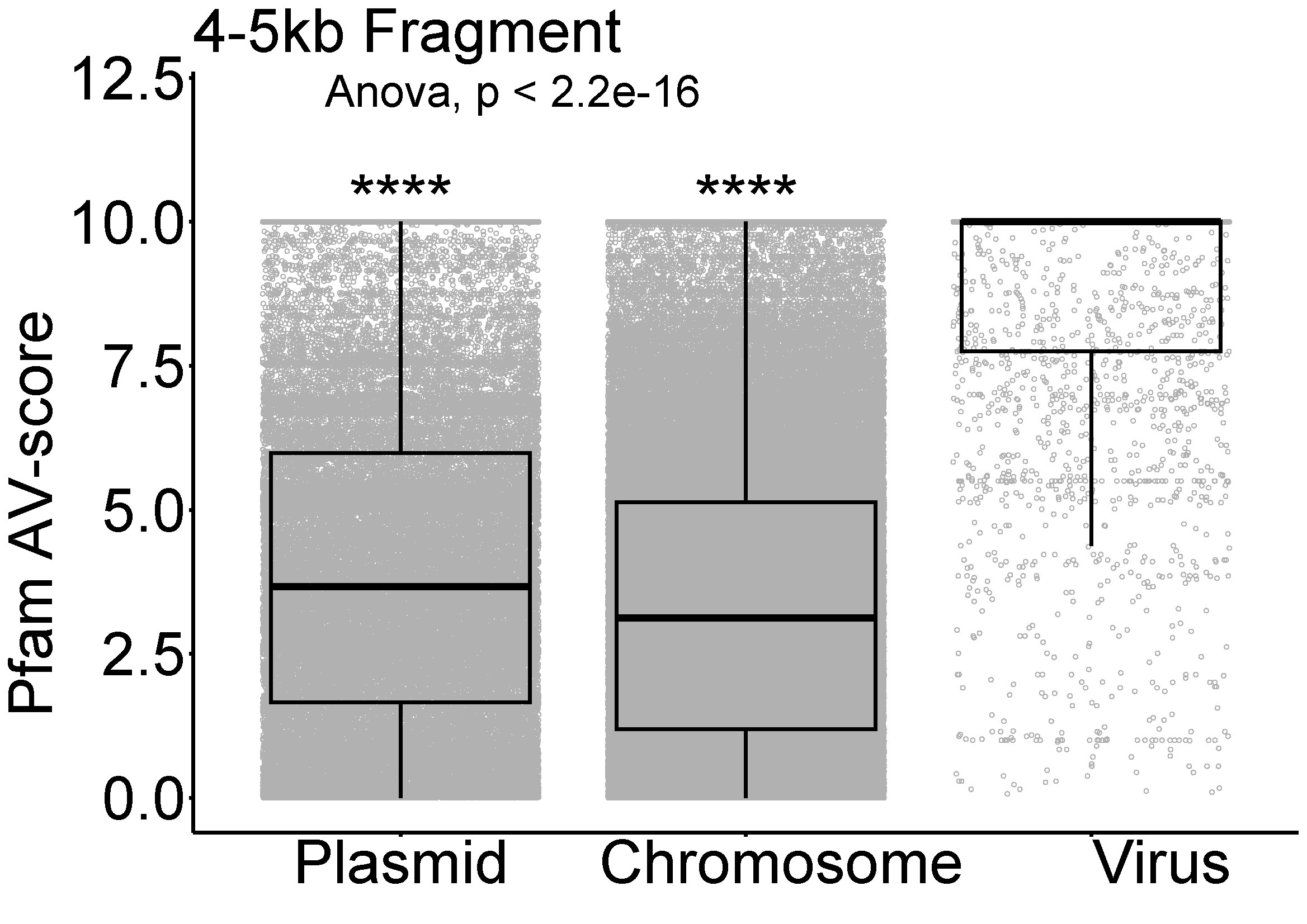

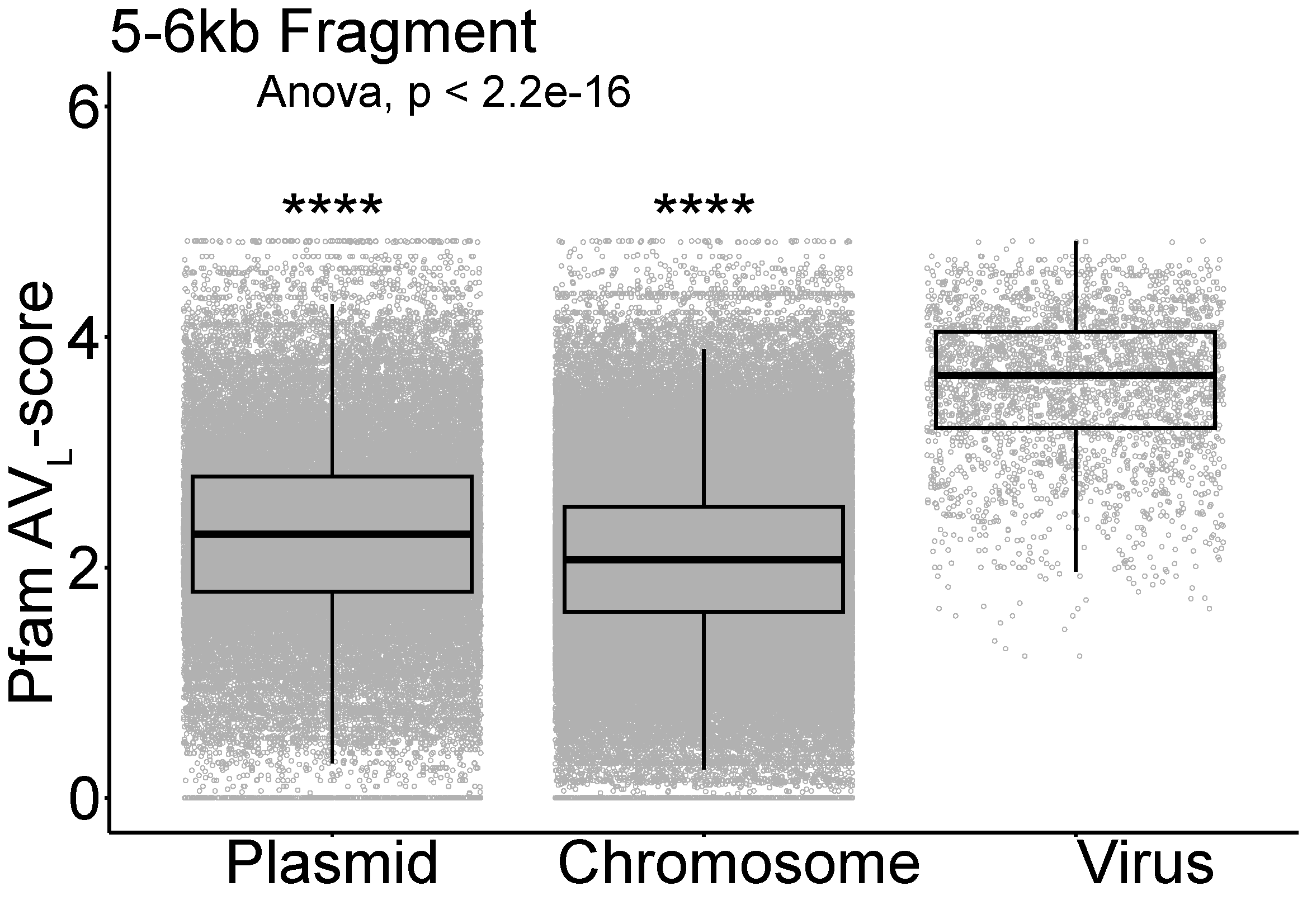

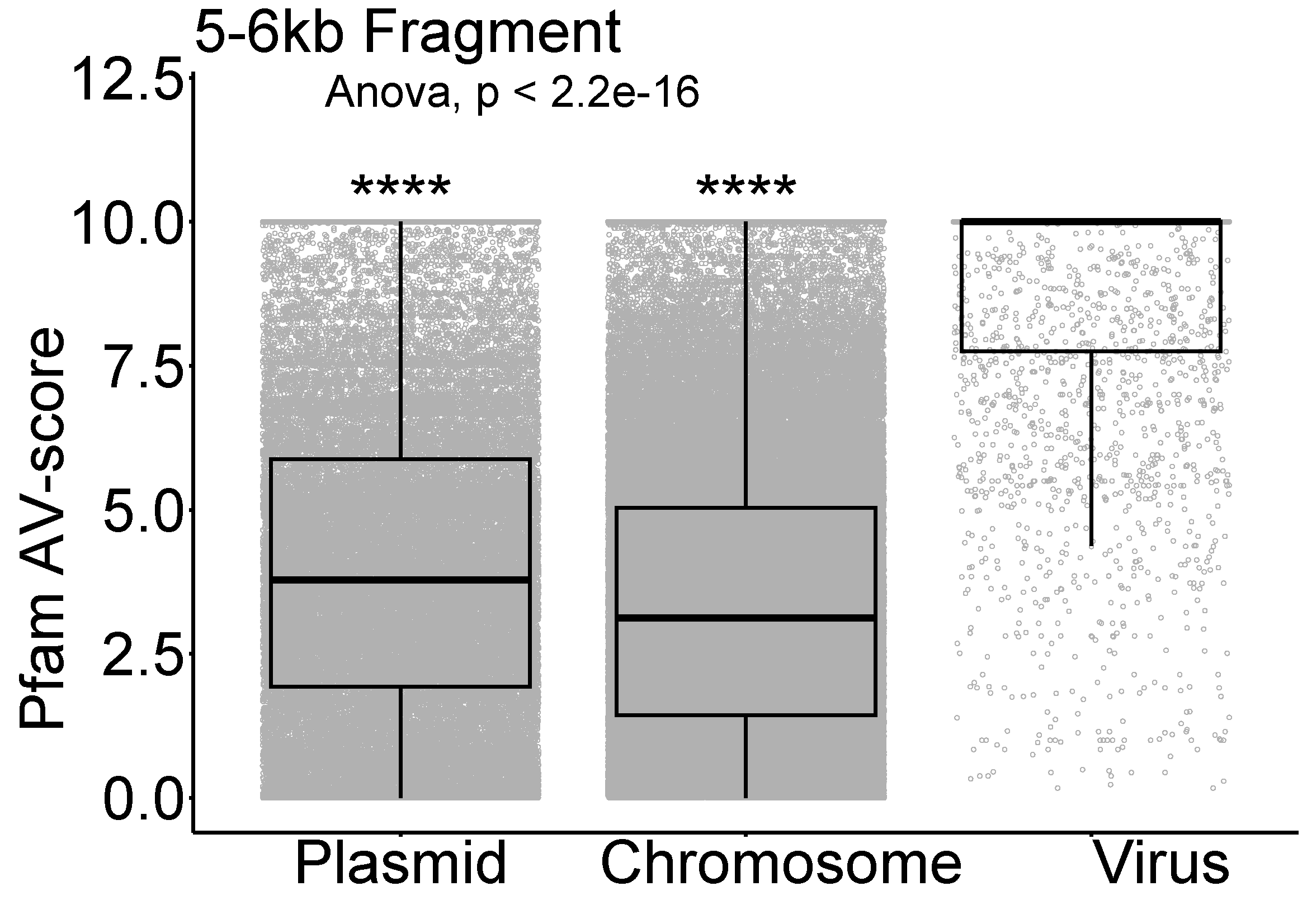

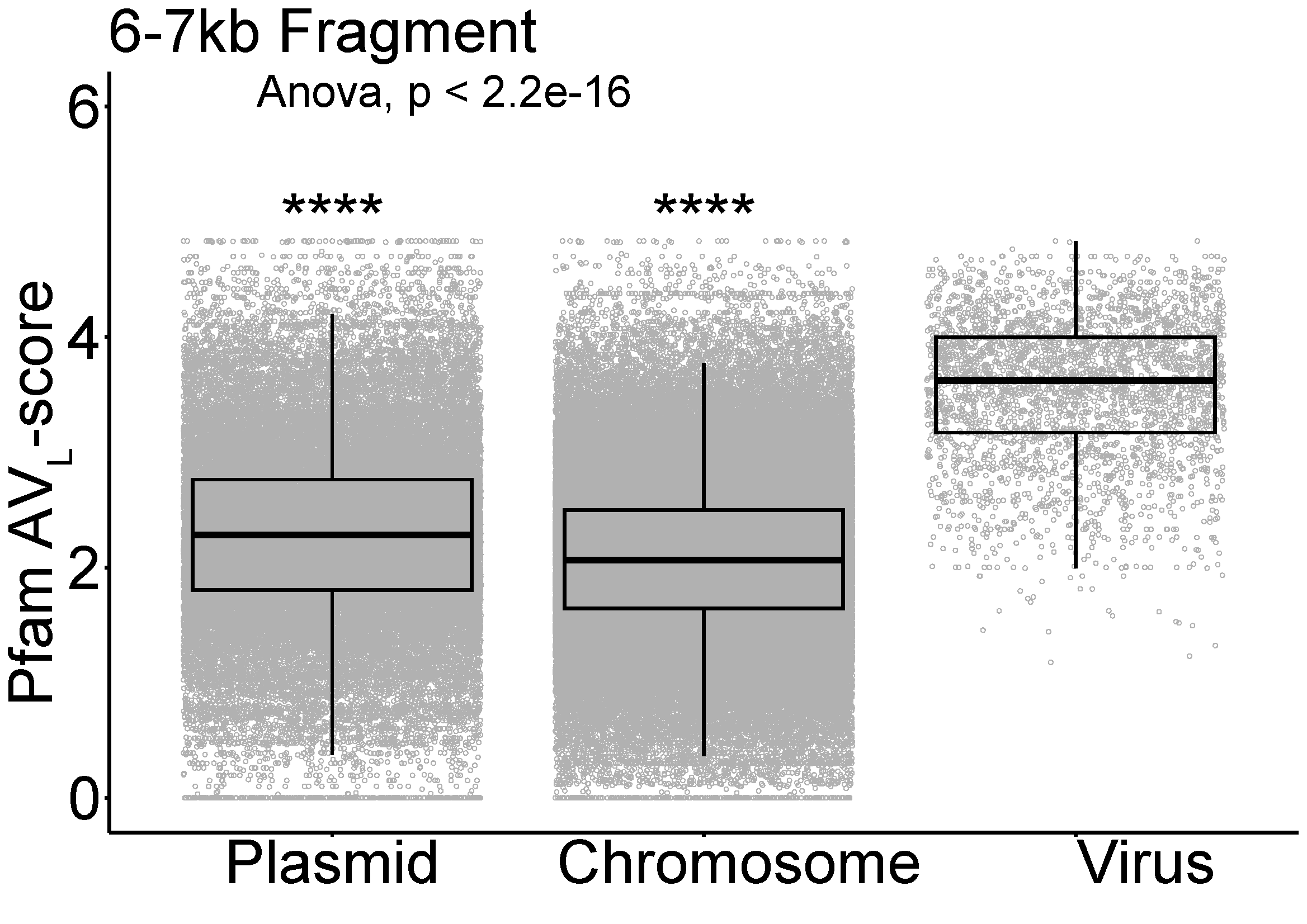

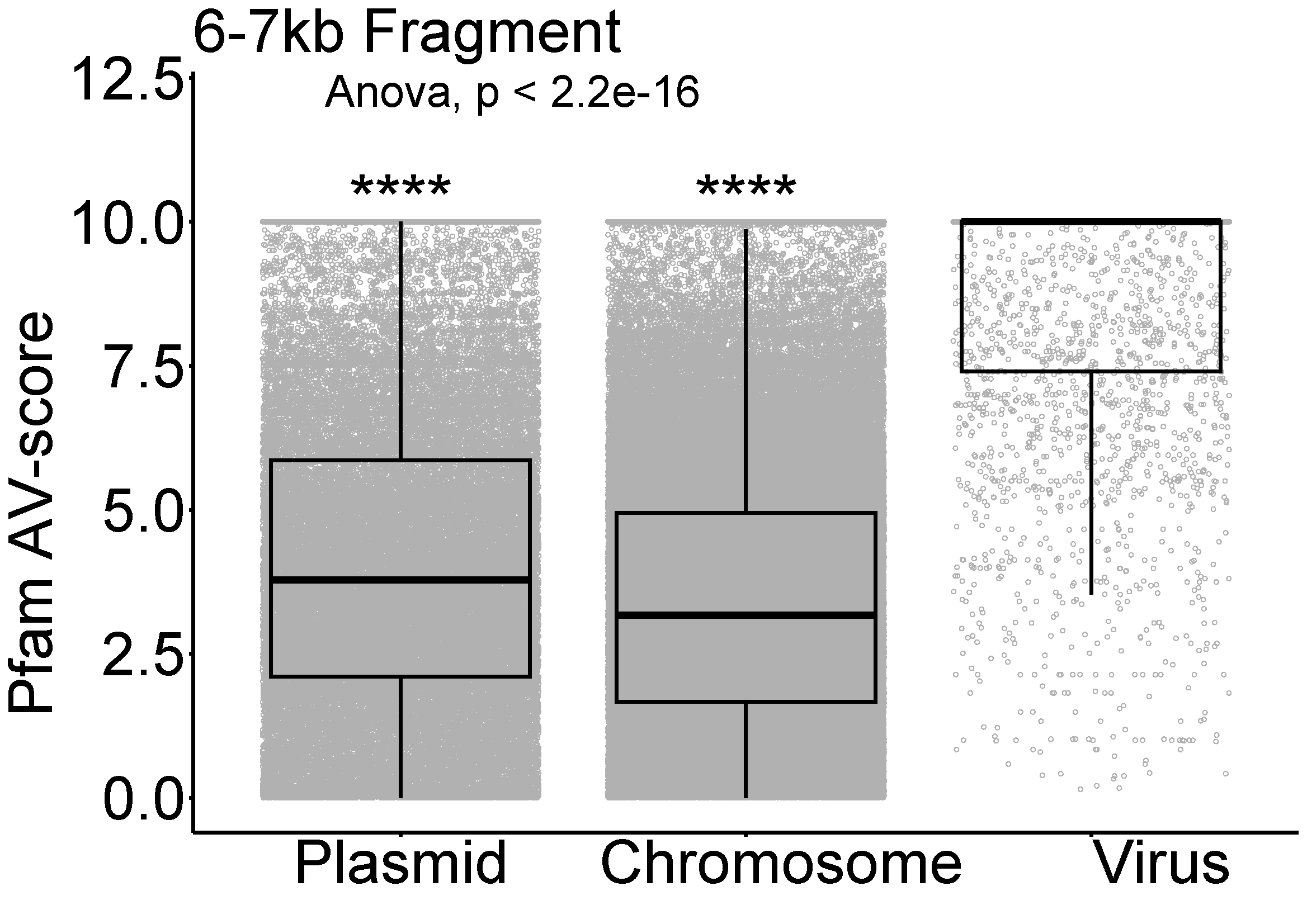

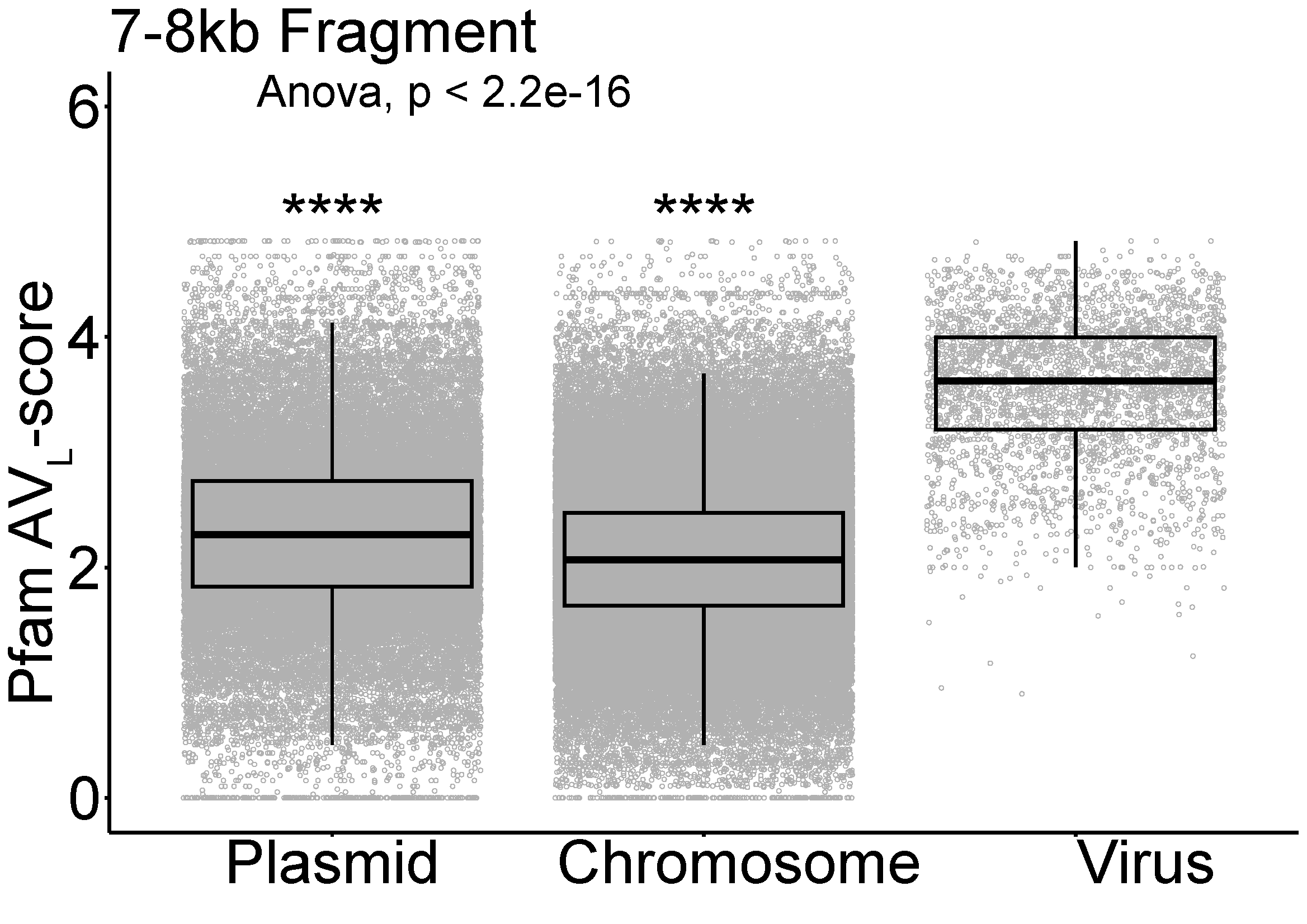

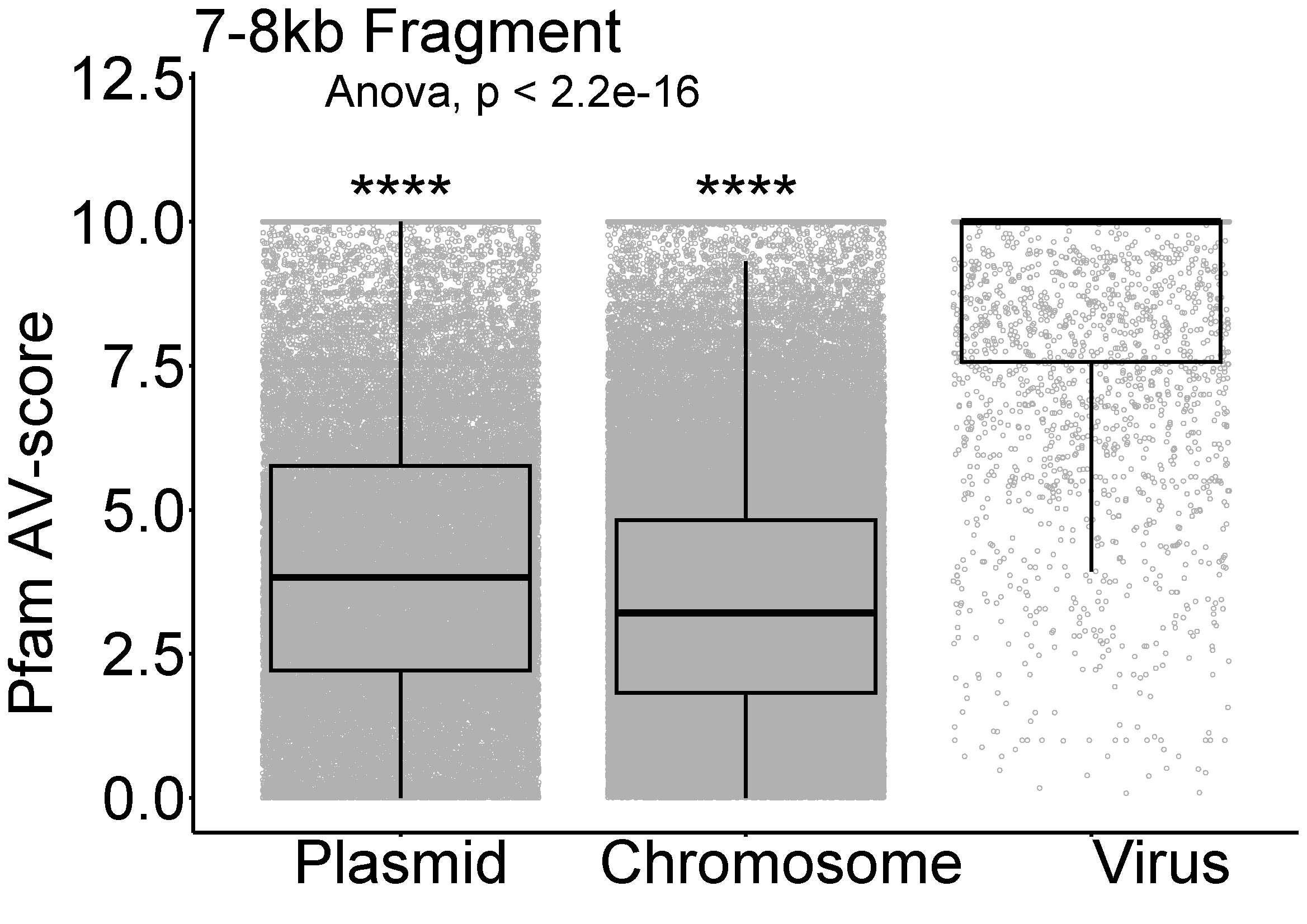

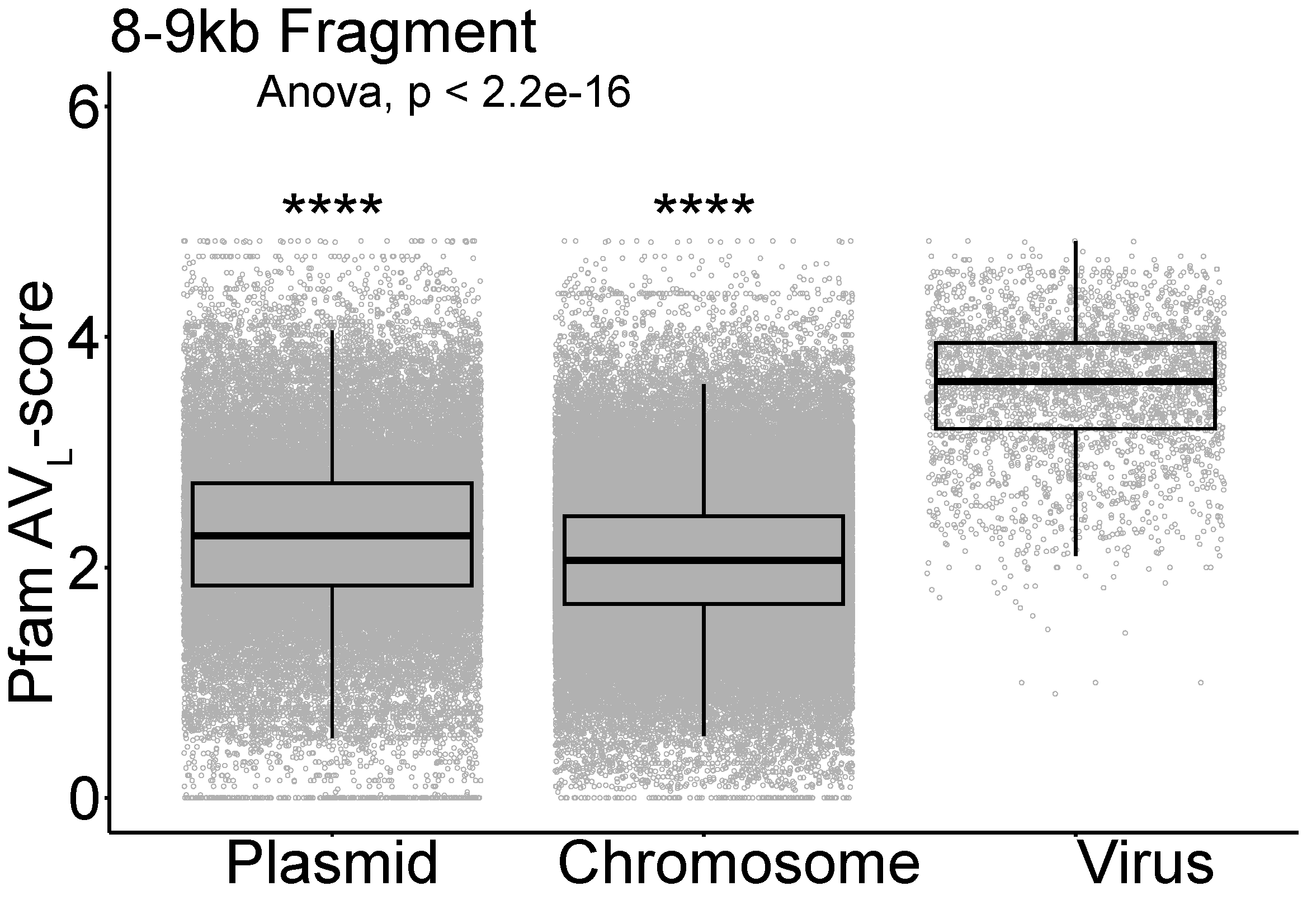

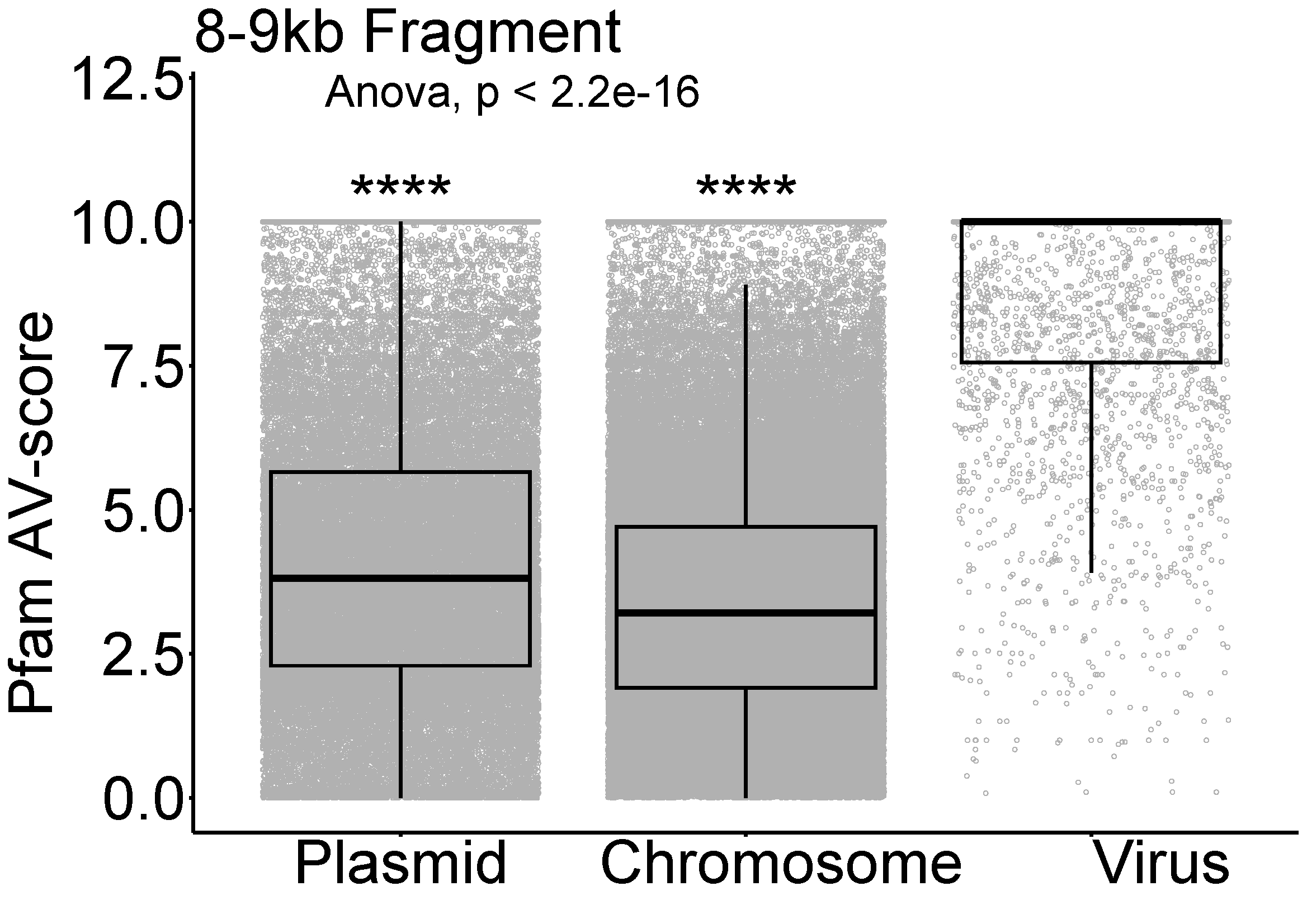

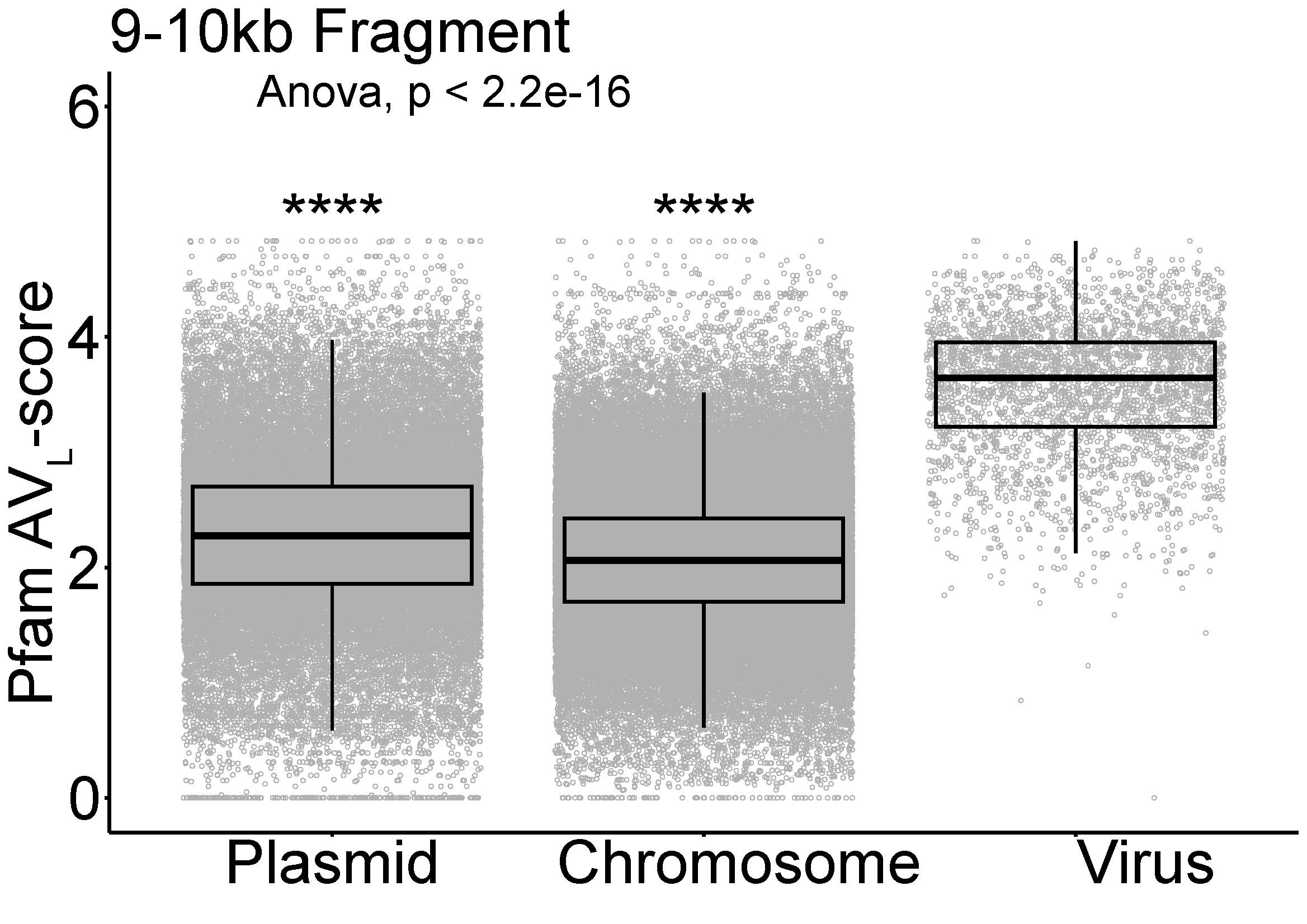

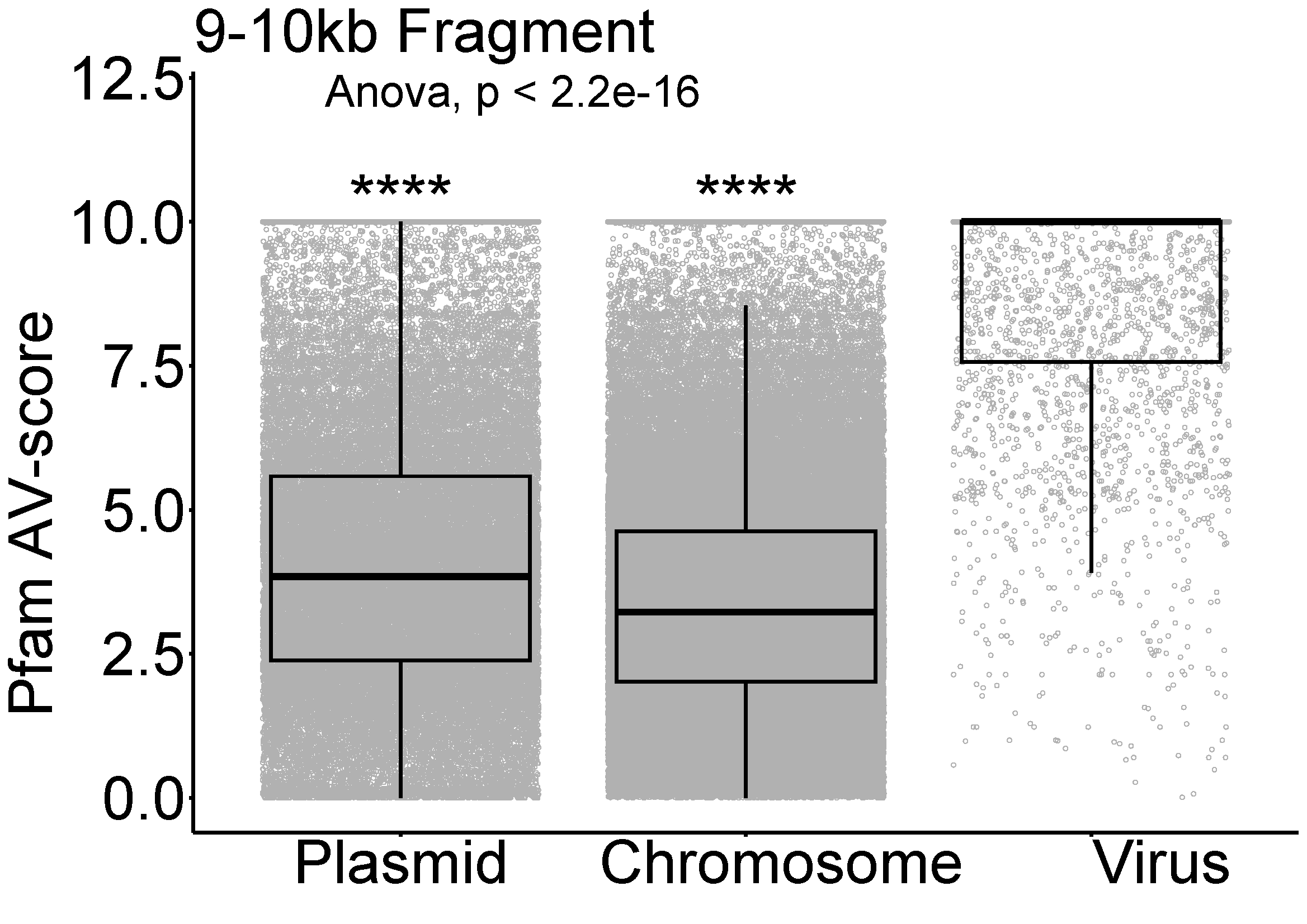

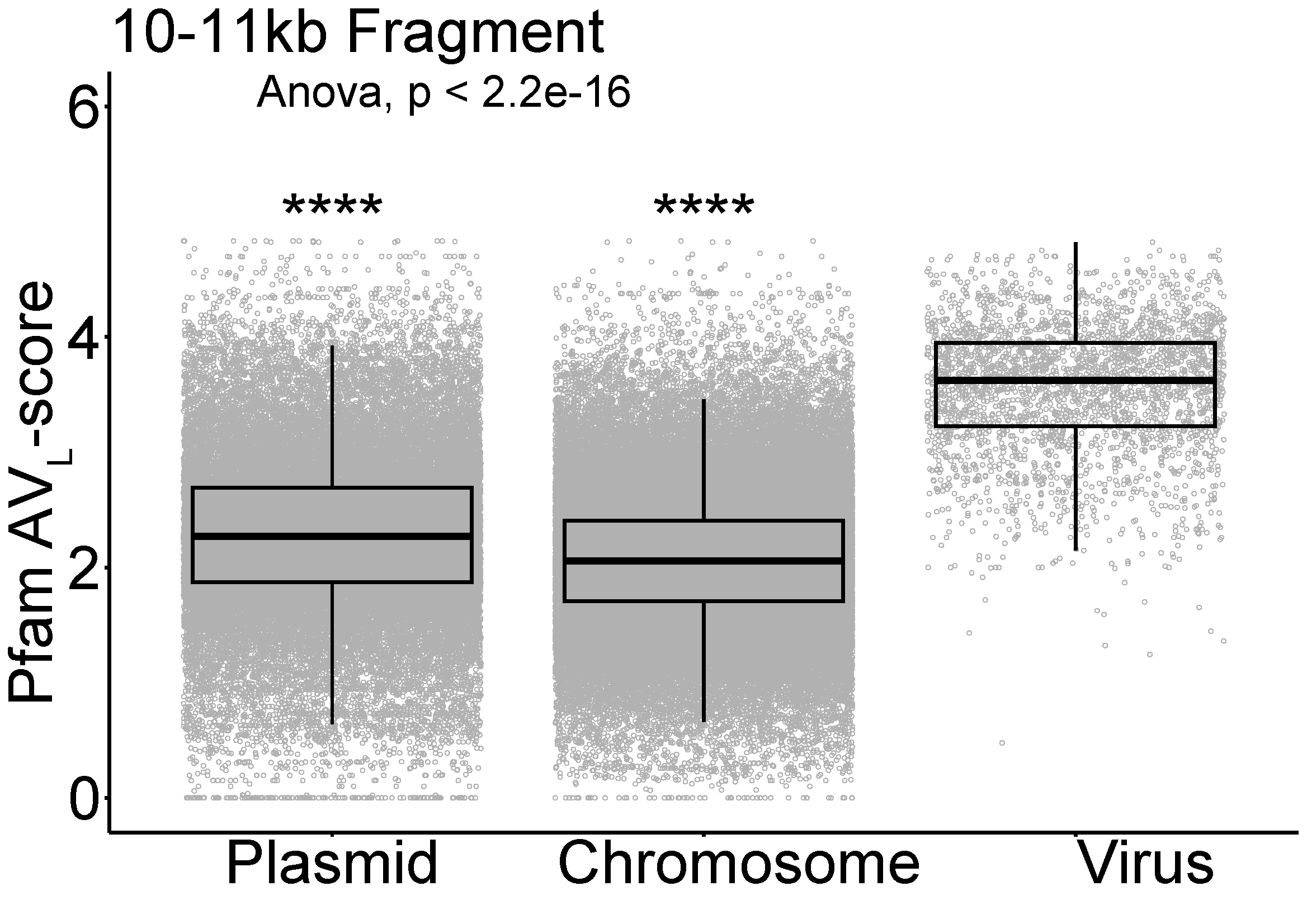

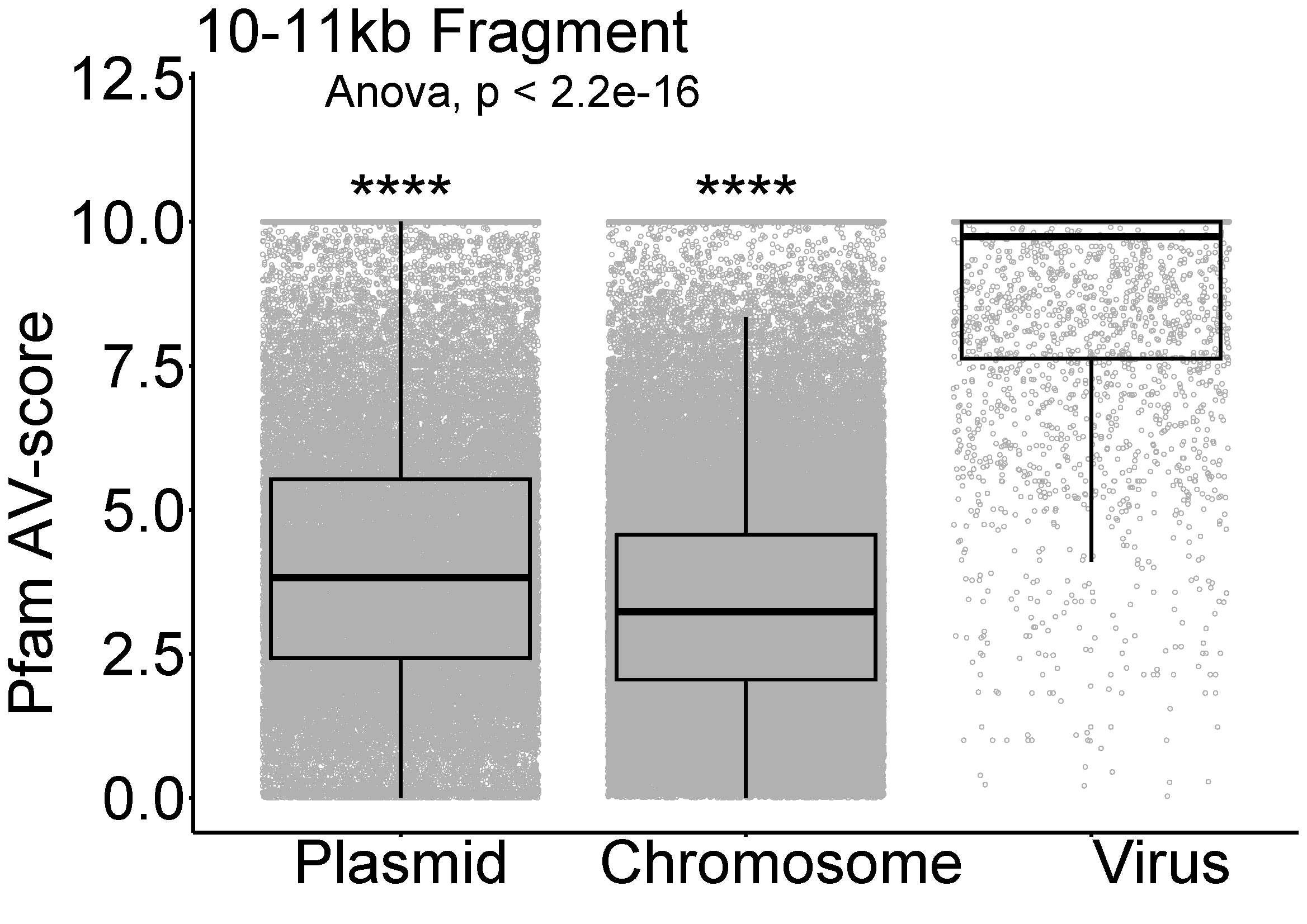

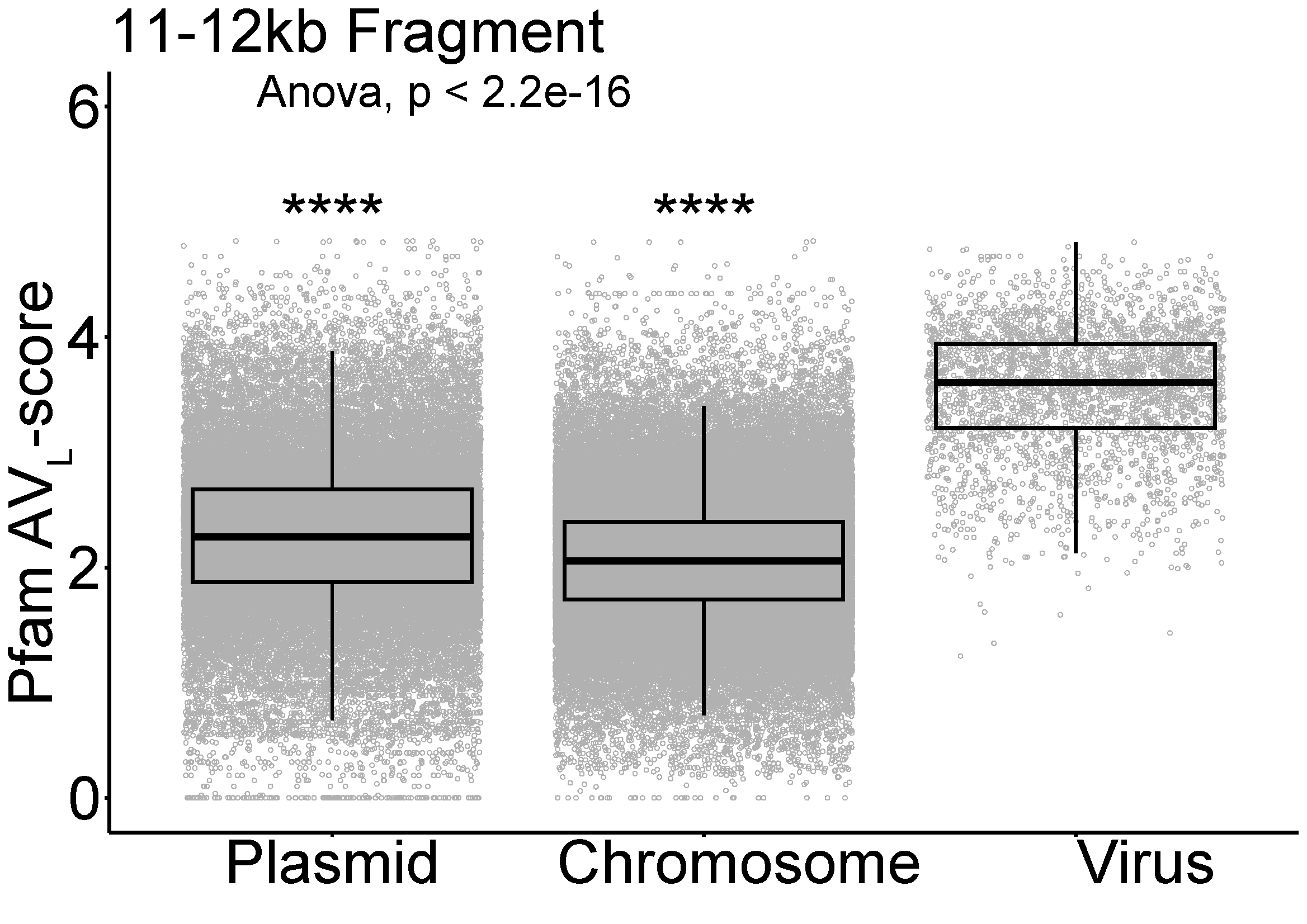

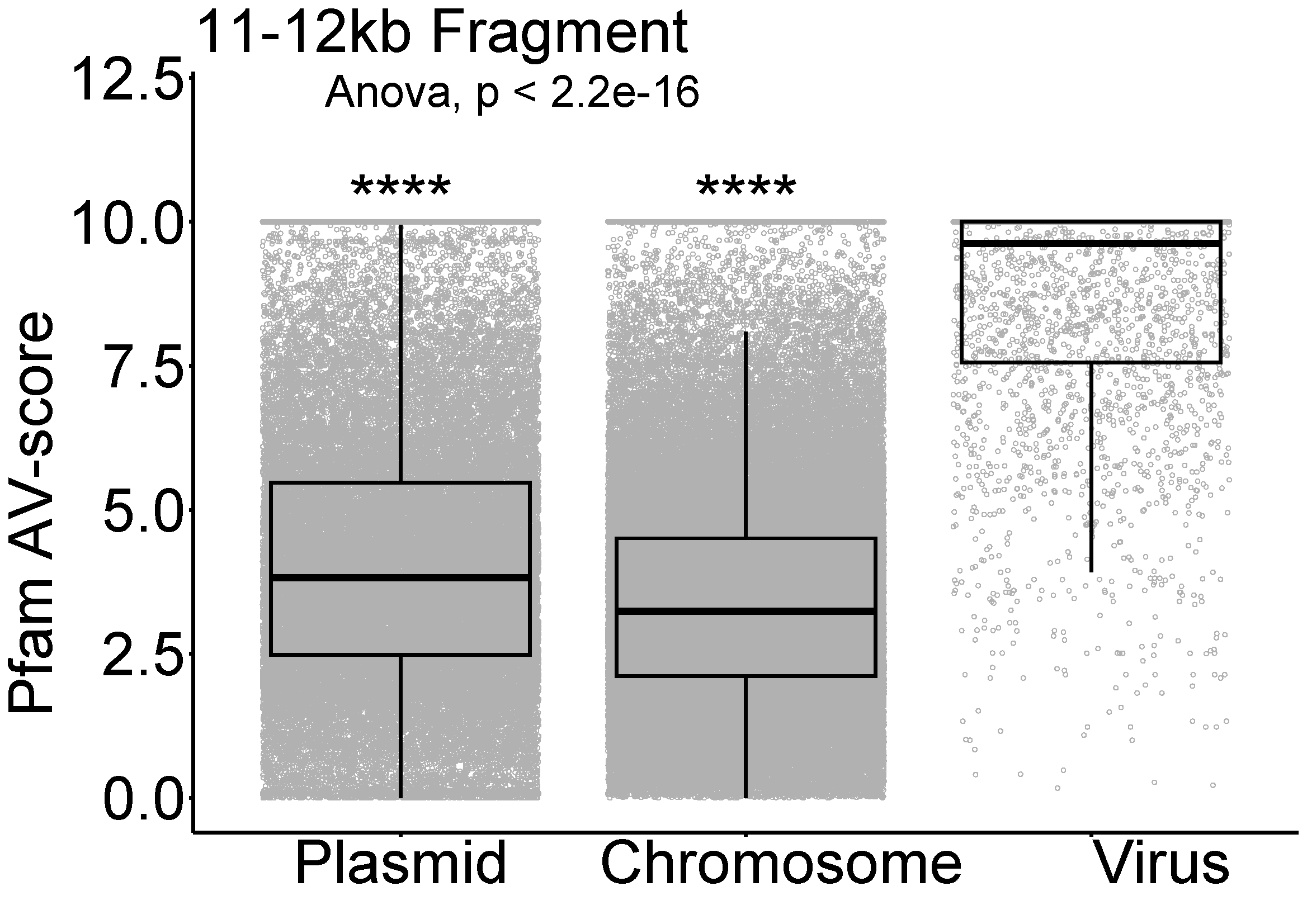

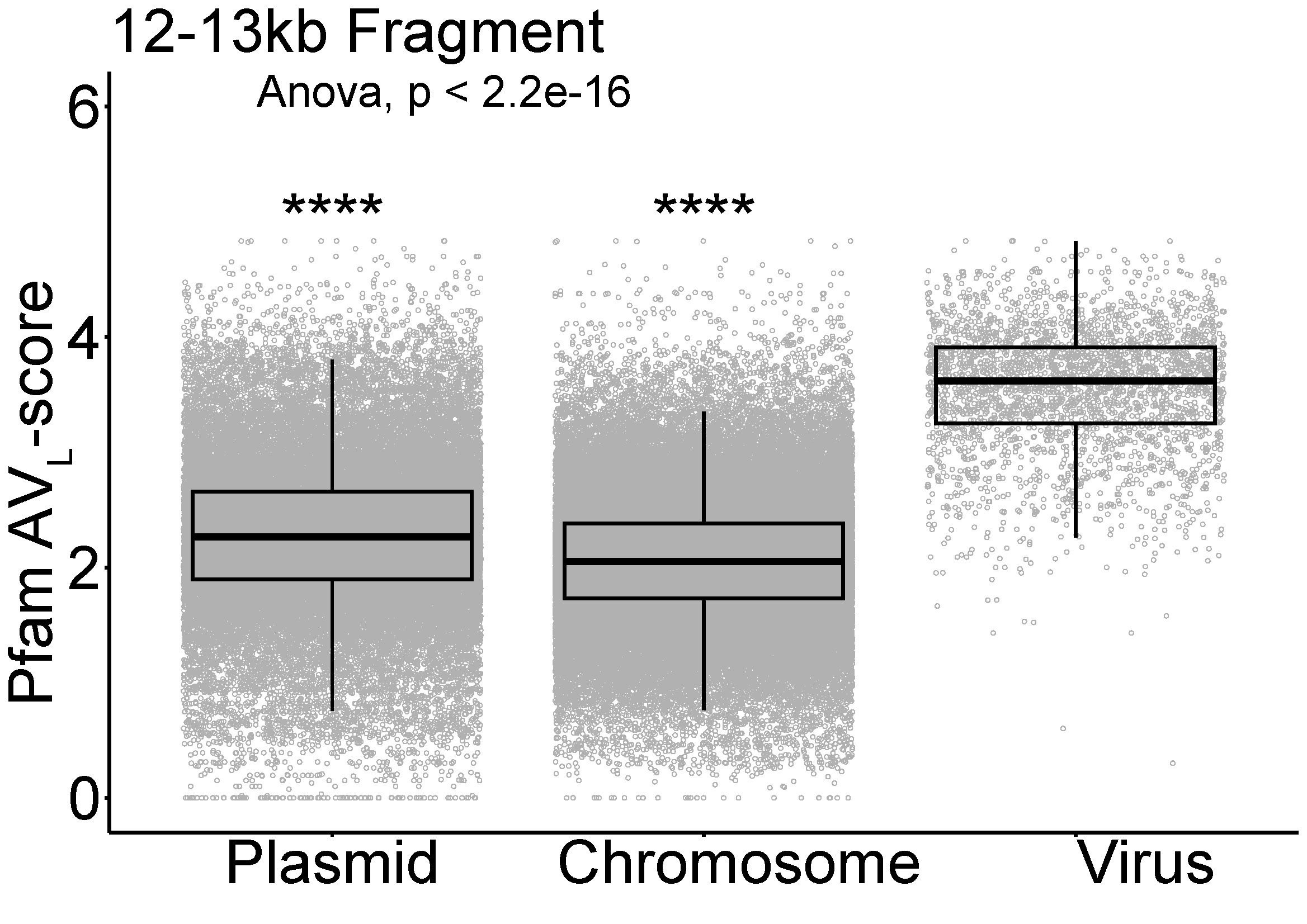

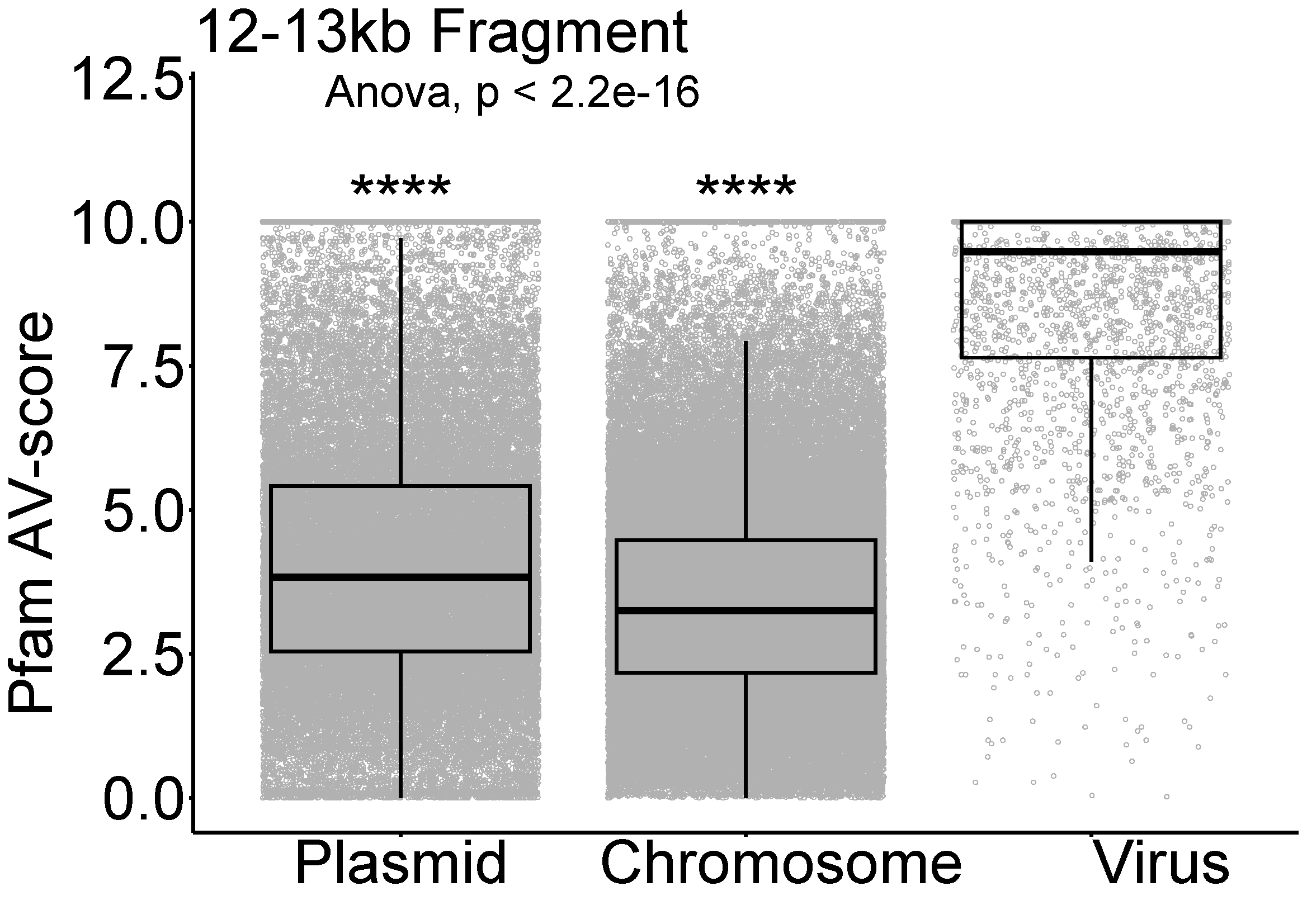

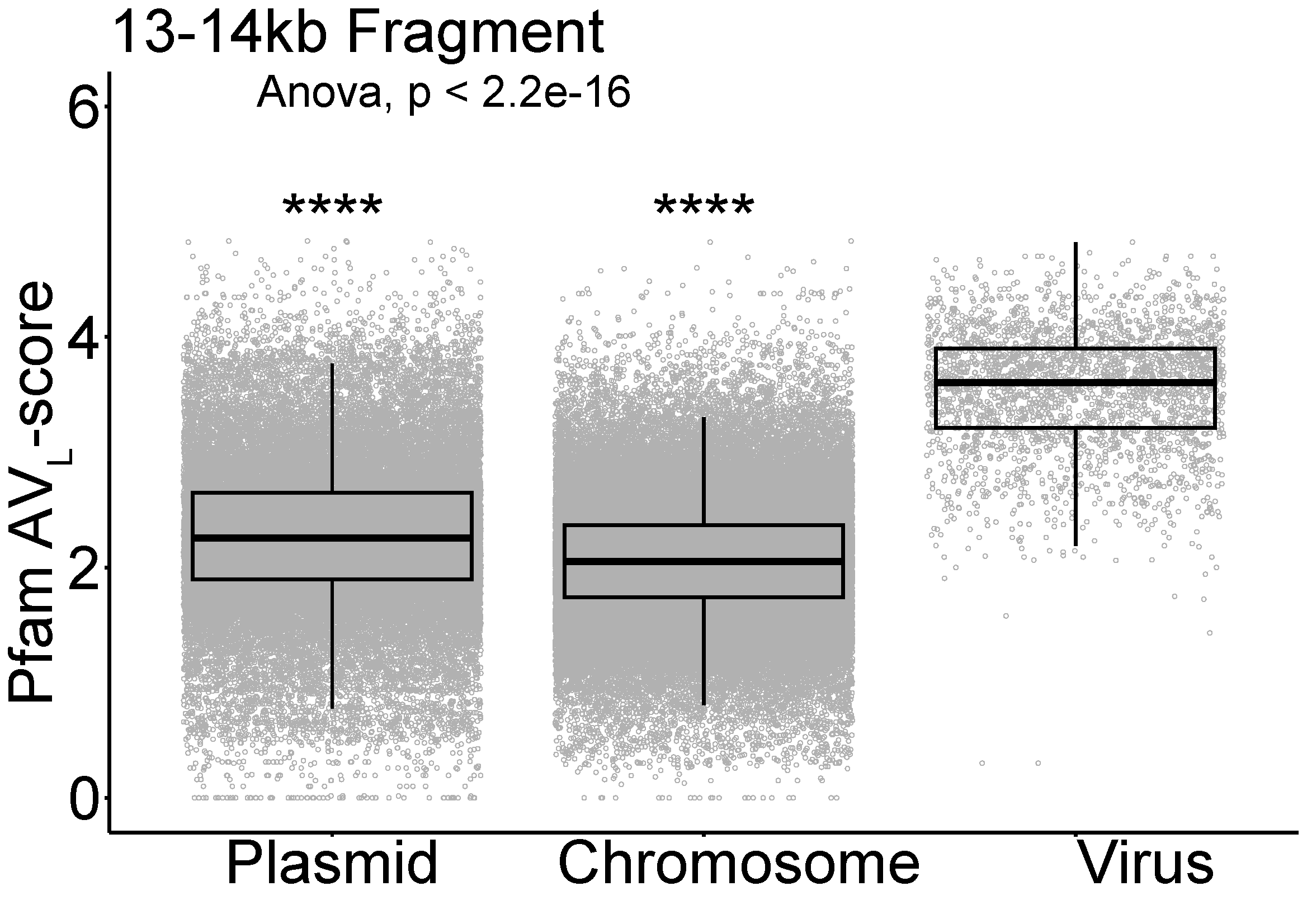

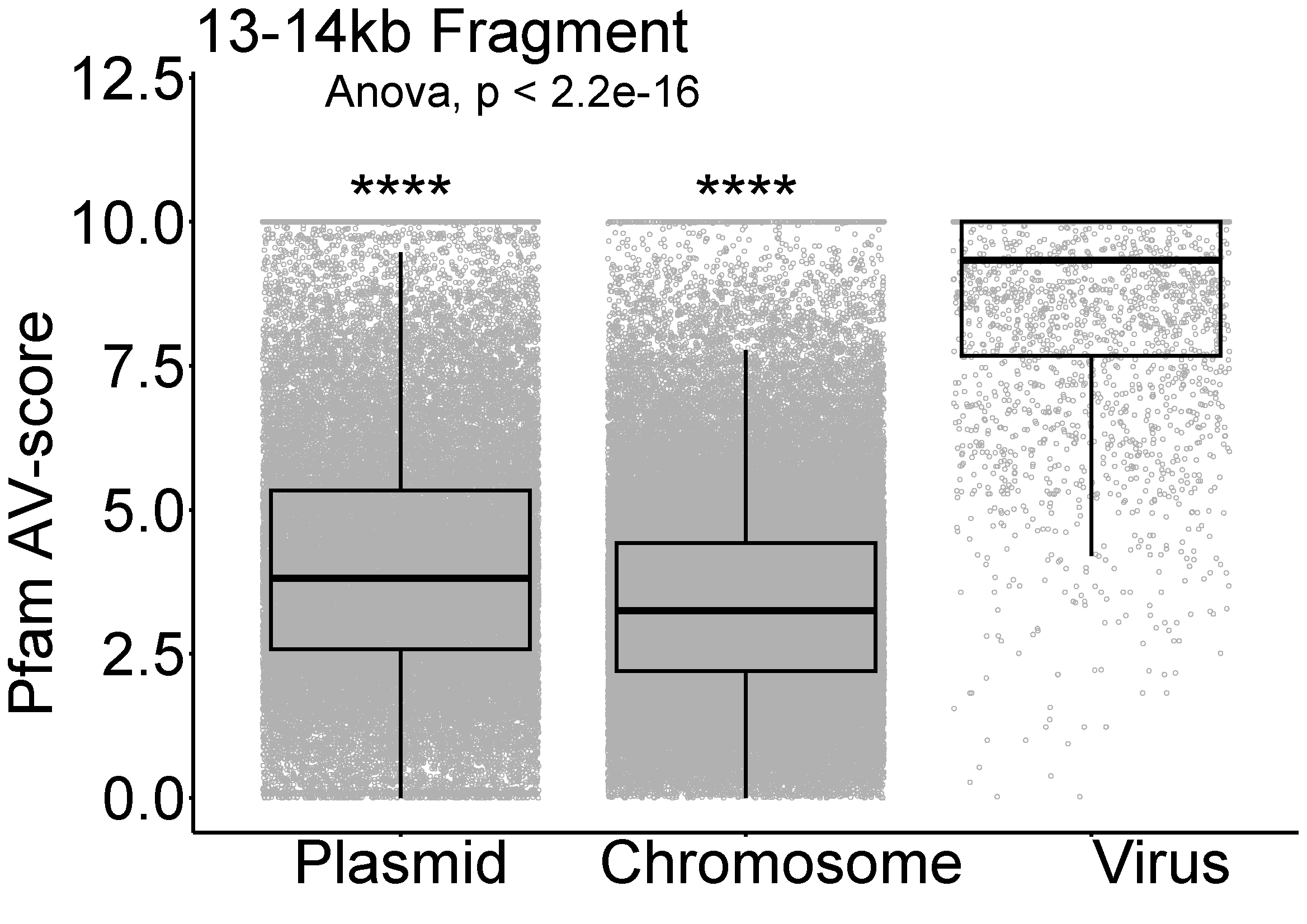

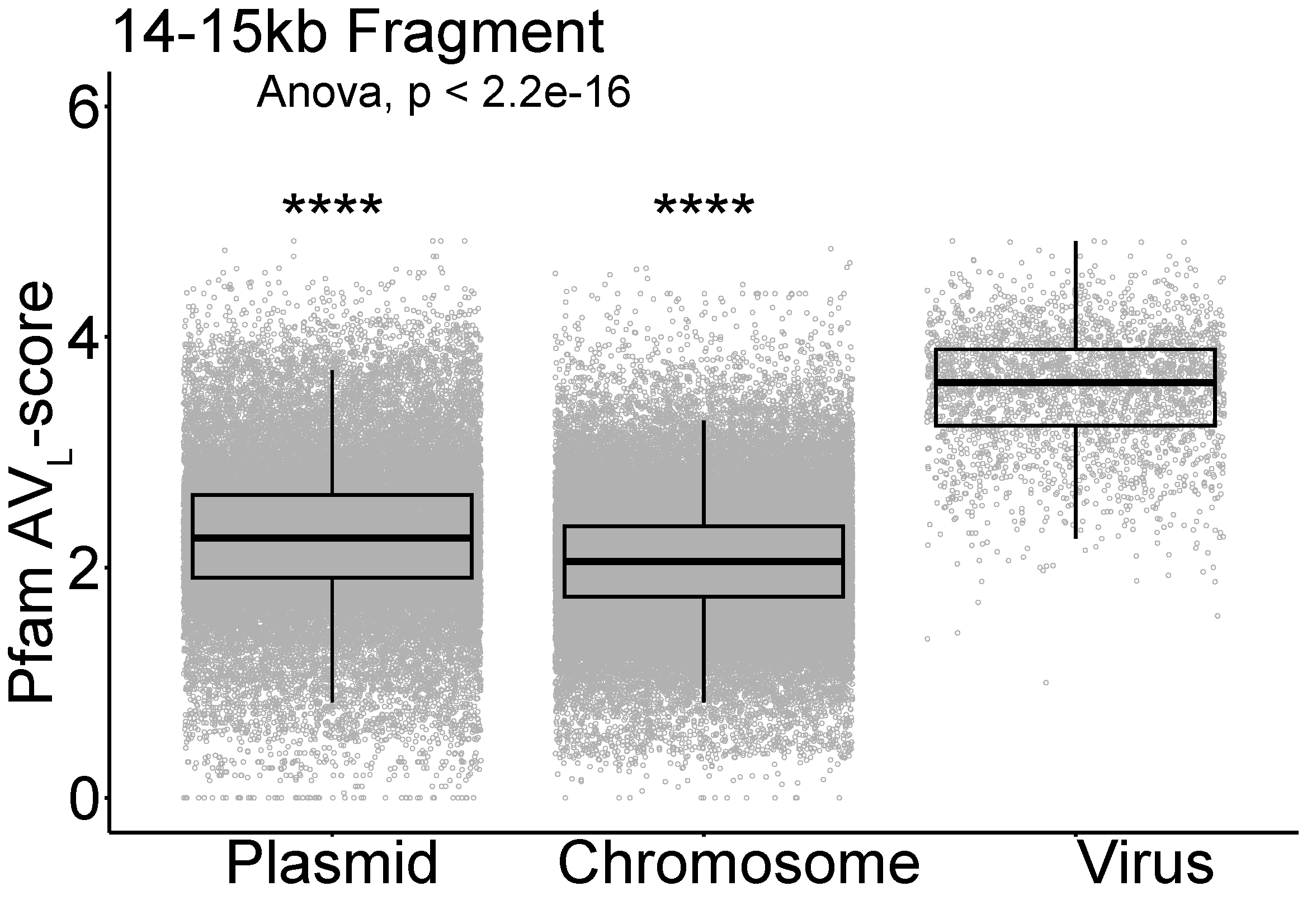

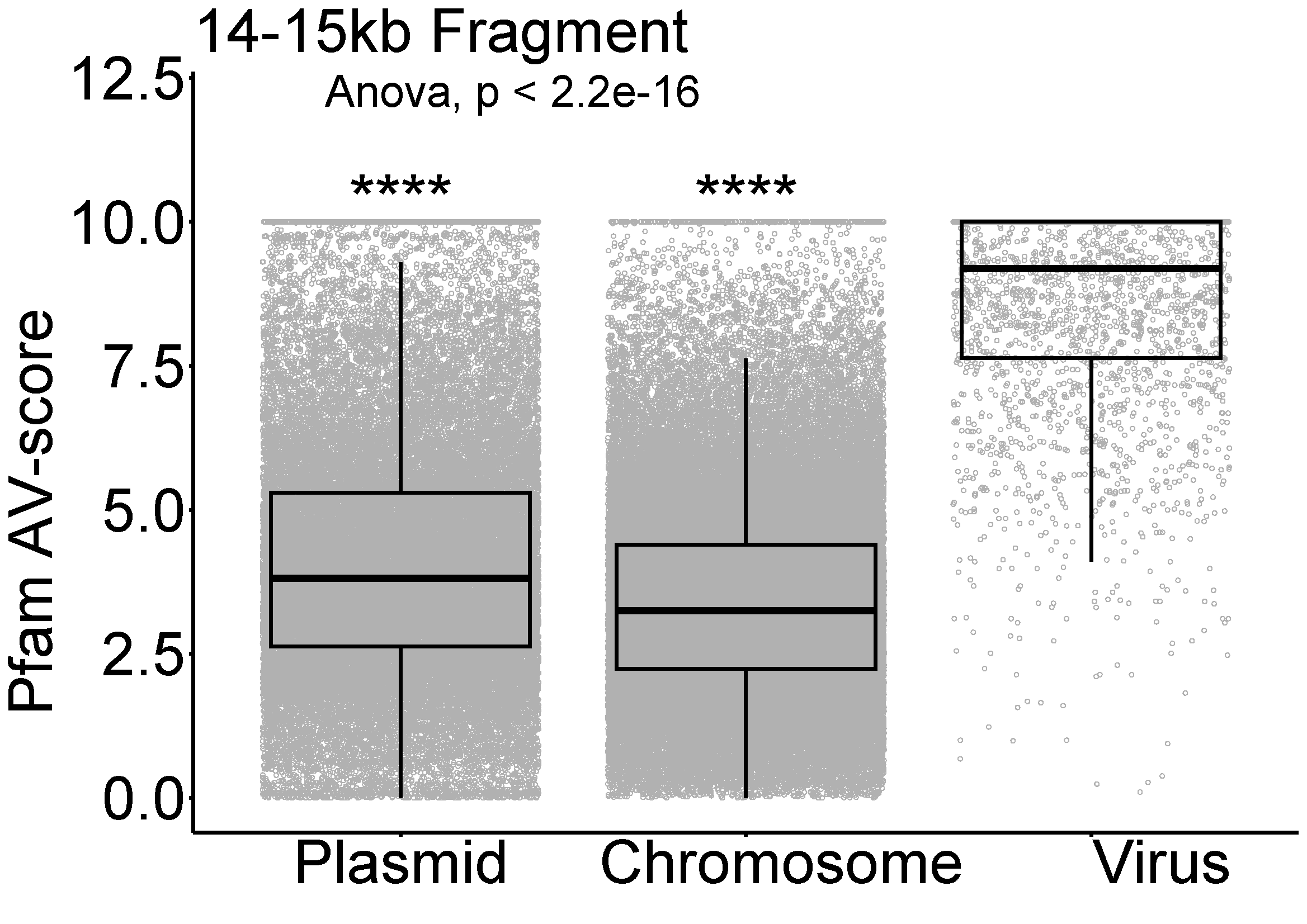

**Fig. S3.** Distribution of Pfam AV-score and AV_L_-score of split prokaryotic chromosome fragments (*n* = 1975,048), genome fragments of plasmids (*n* = 681,614) and prokaryotic viruses (*n* = 48,880). The horizontal line that splits the box is the median, the upper and lower sides of the box are upper and lower quartiles, whiskers are 1.5 times the interquartile ranges and data points beyond whiskers are considered potential outliers. An ANOVA test was used to show differences between three means are significant (*p* < 2.2 × 10^-16^). The asterisks (****) indicate a comparison with viruses, with a significance level of *p* < 10^-4^.

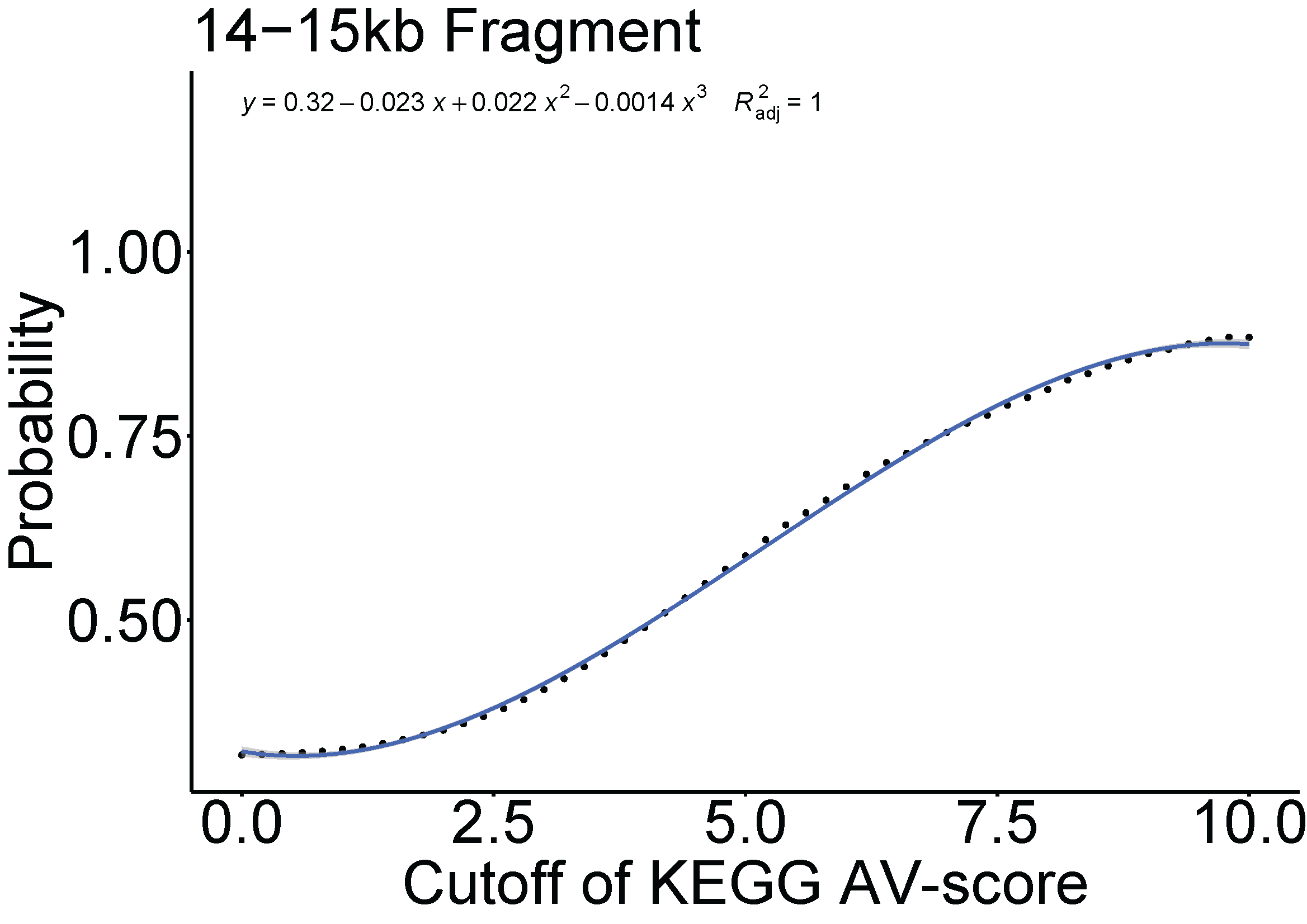

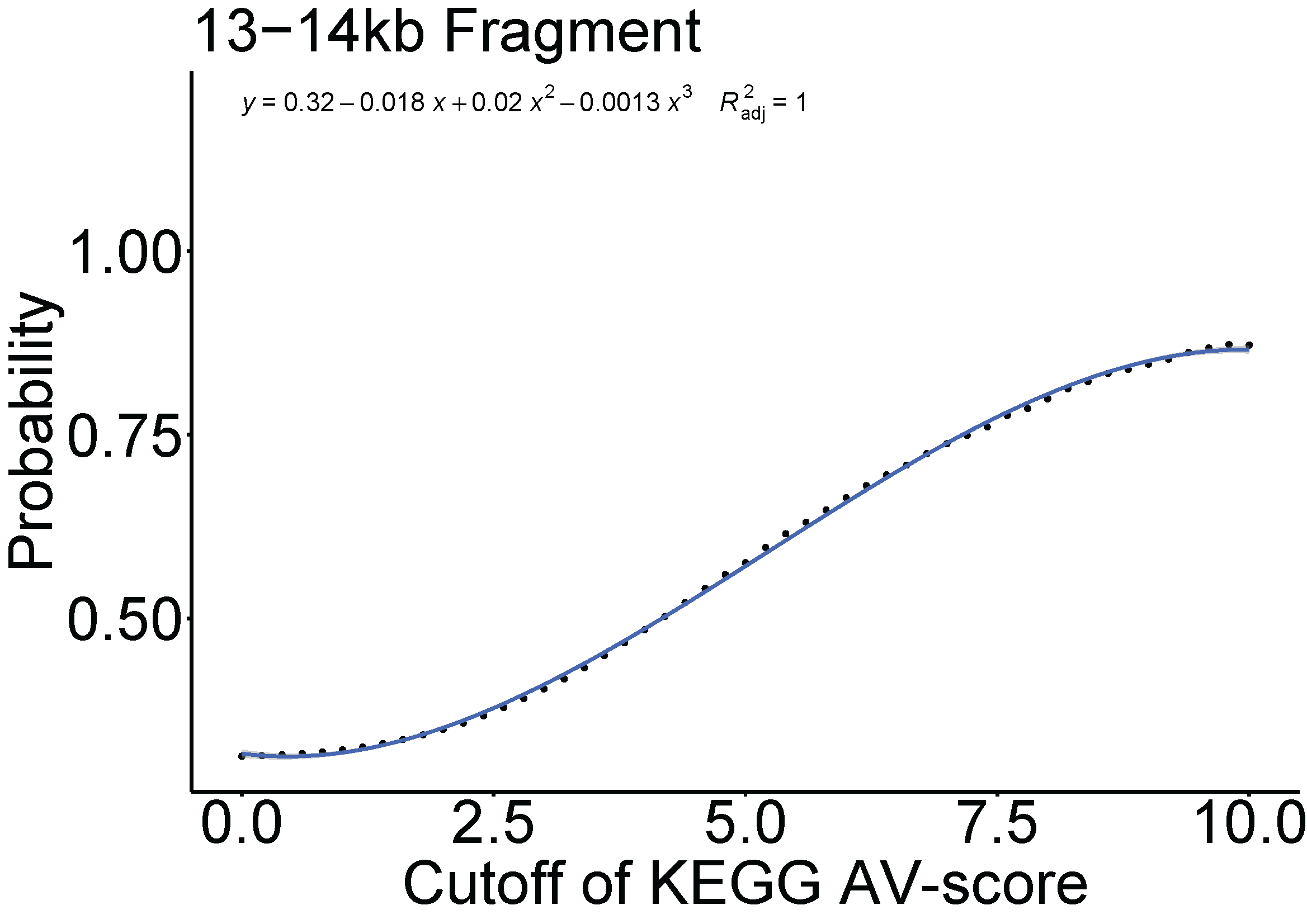

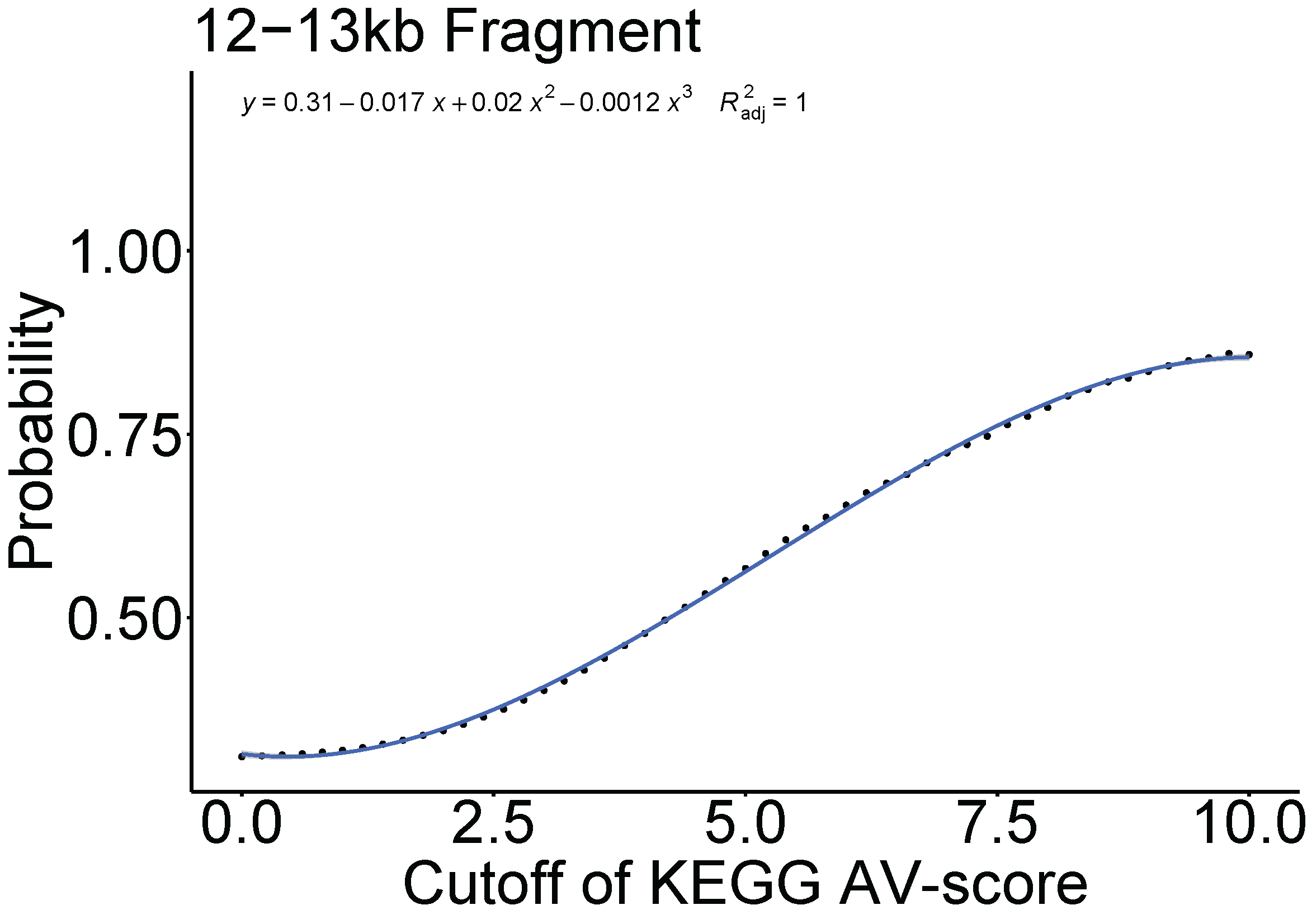

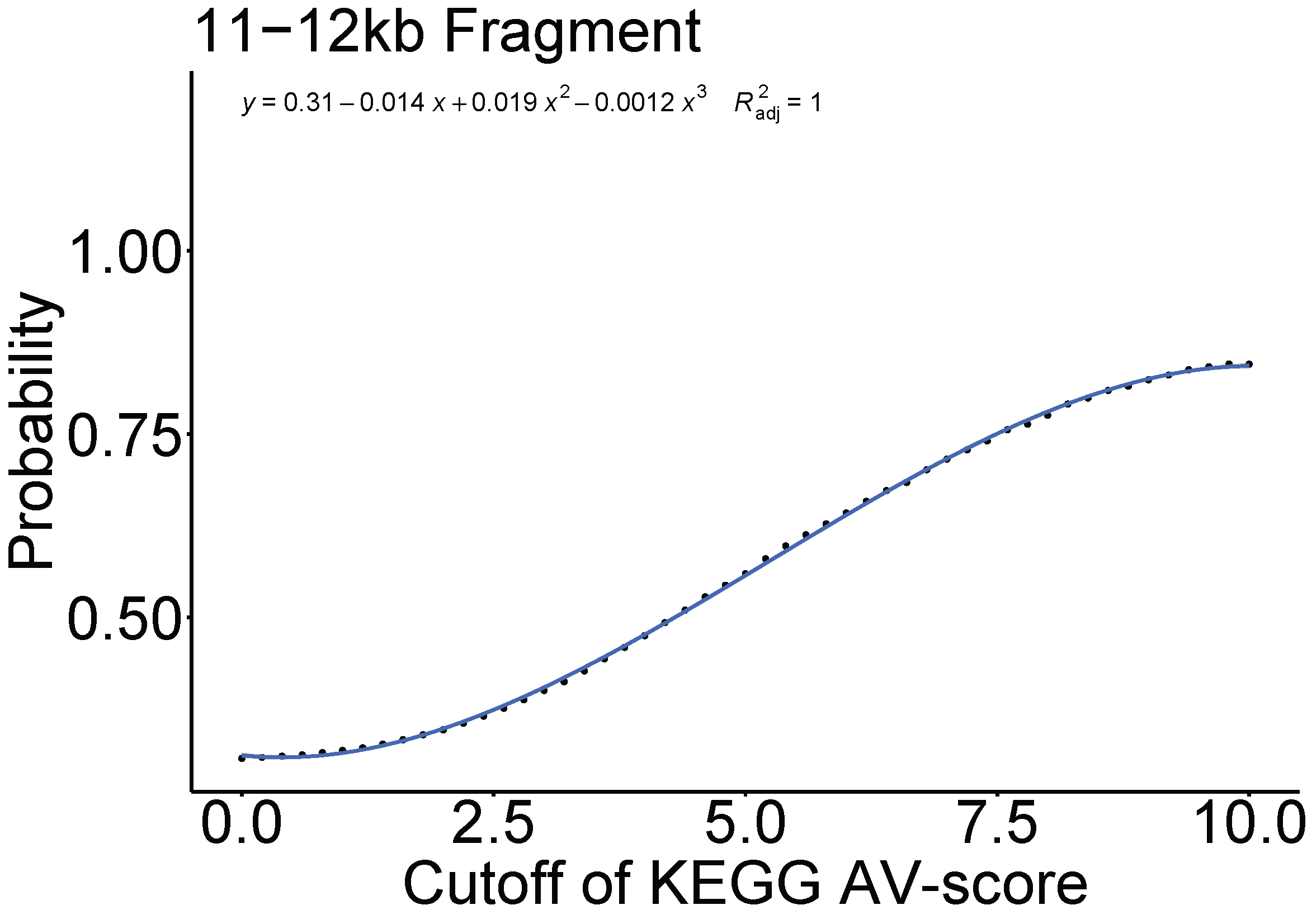

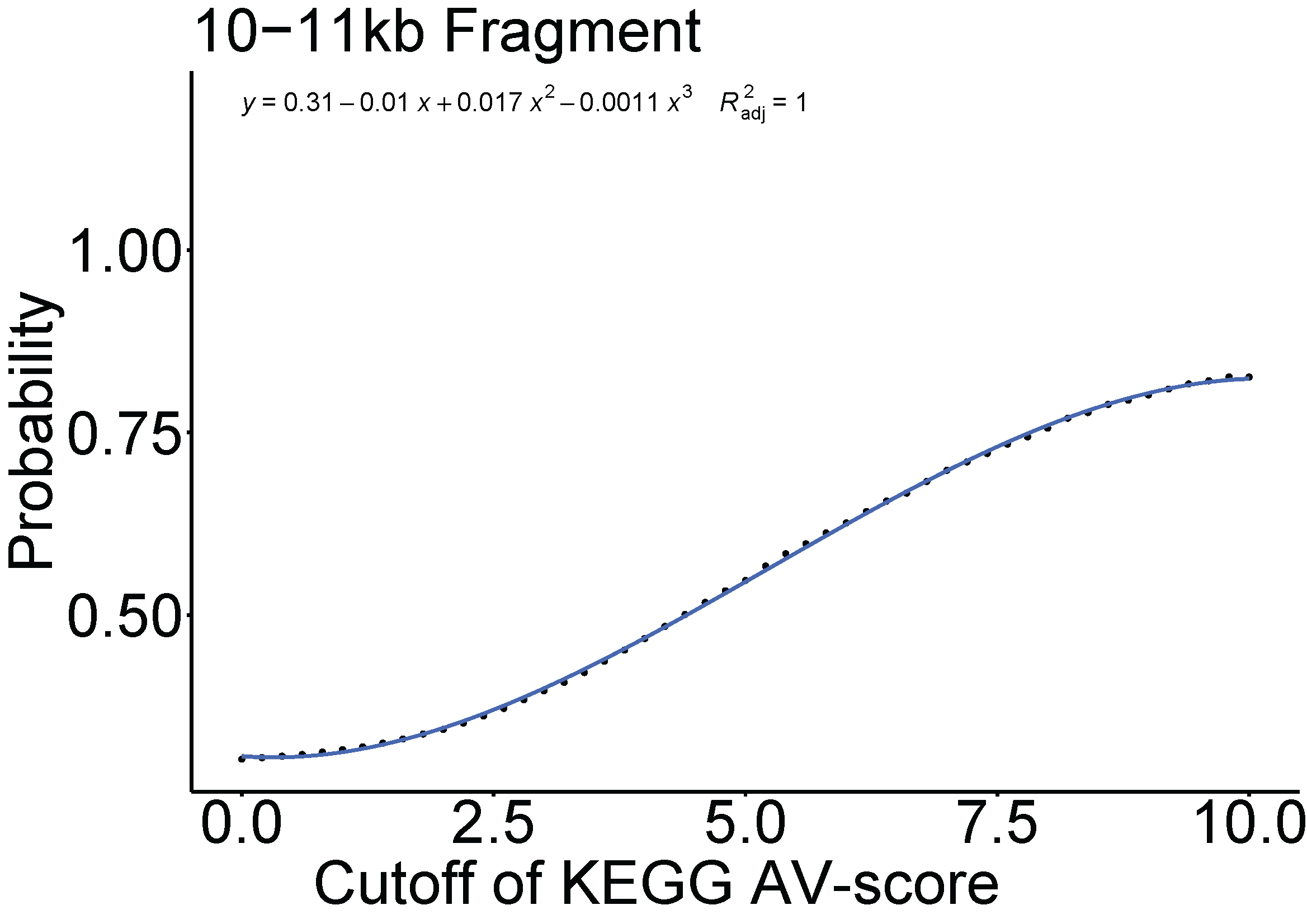

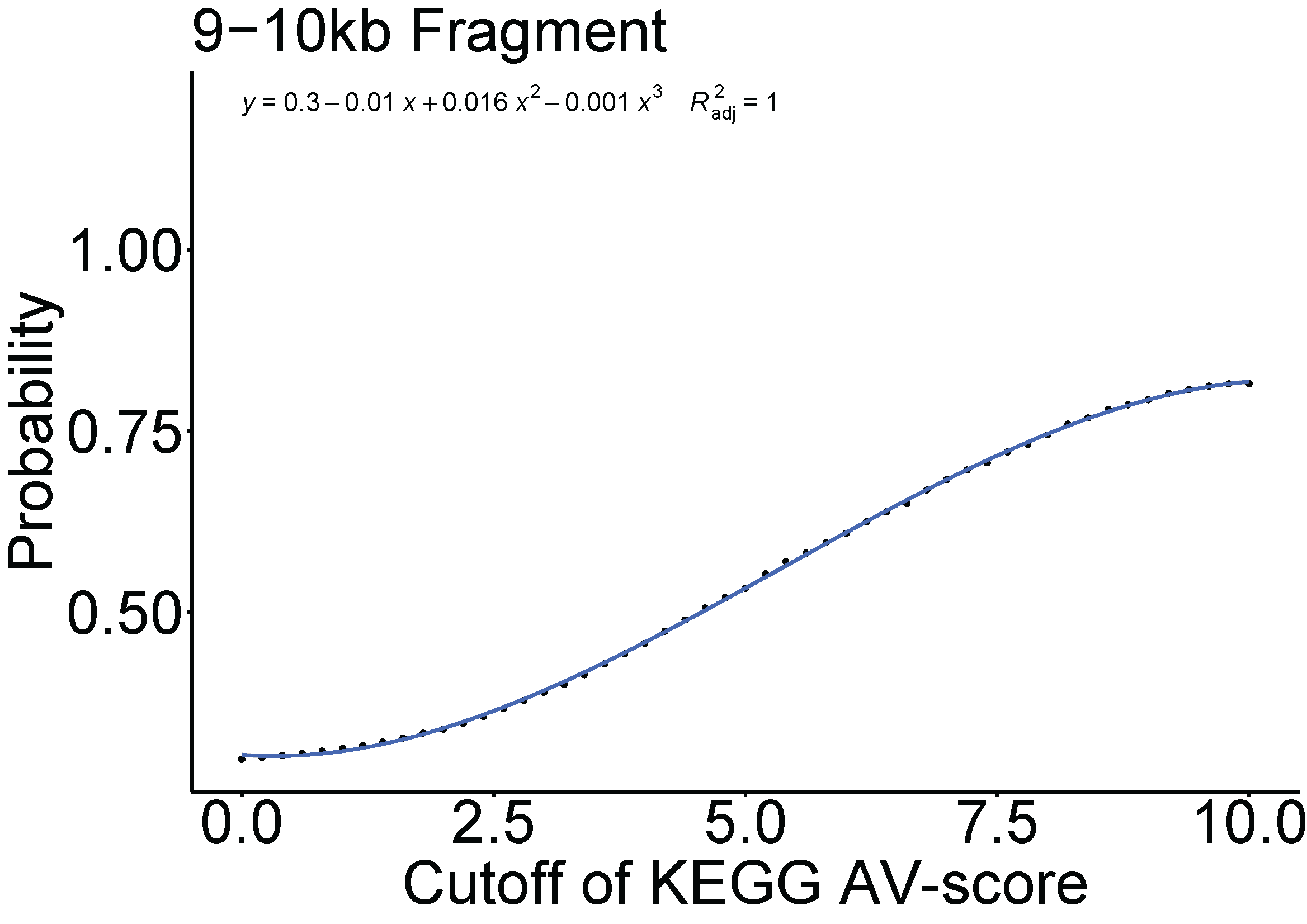

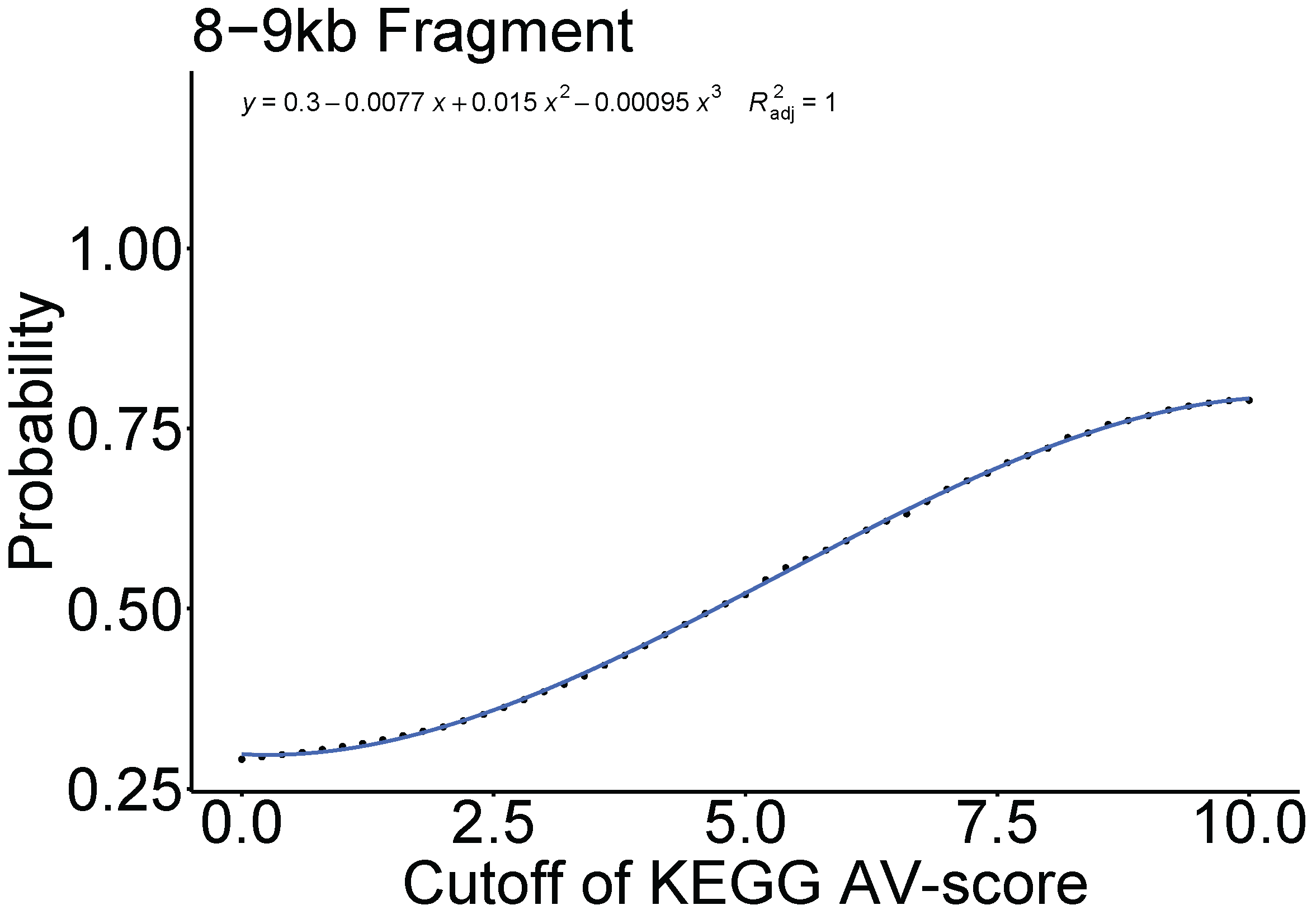

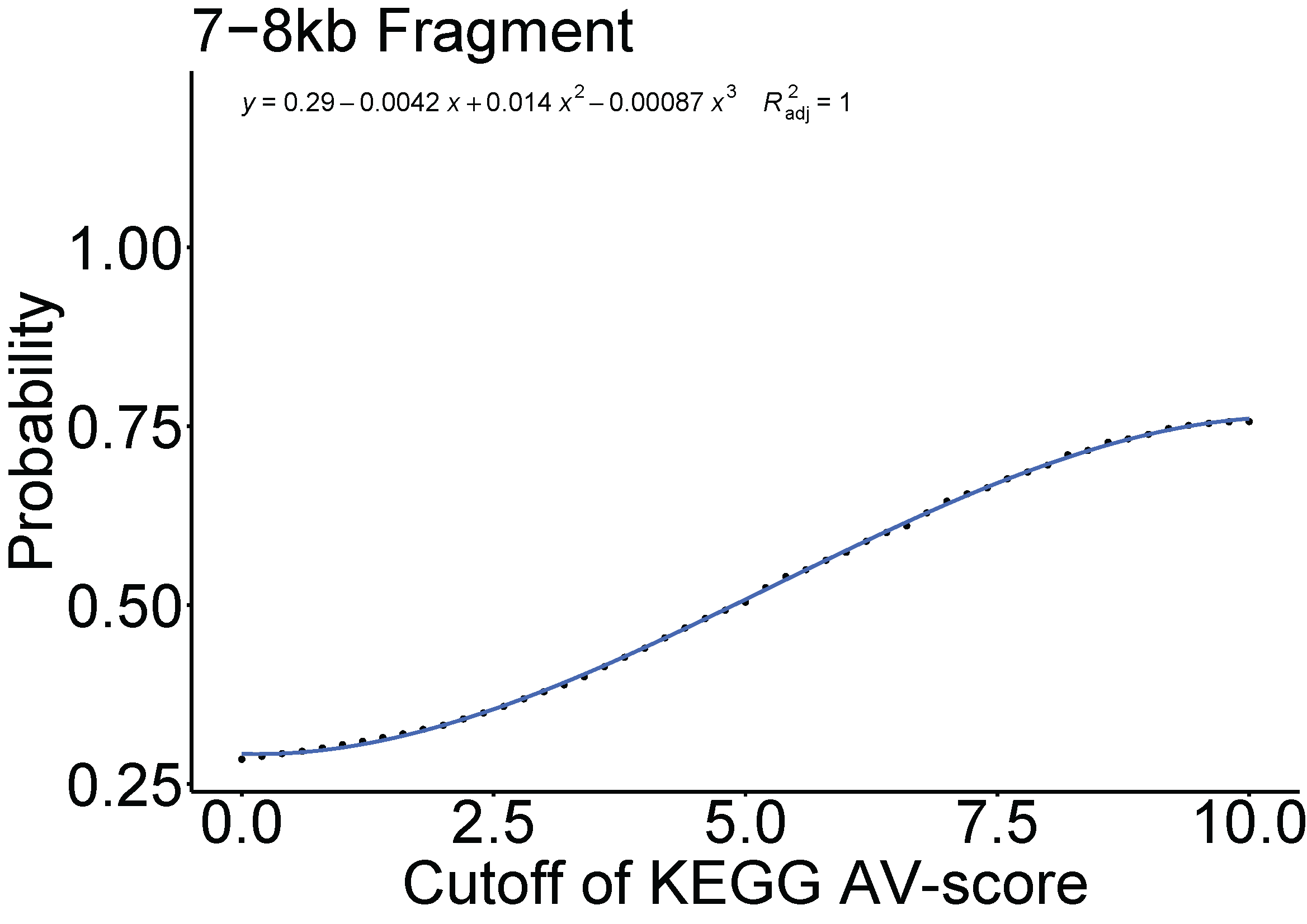

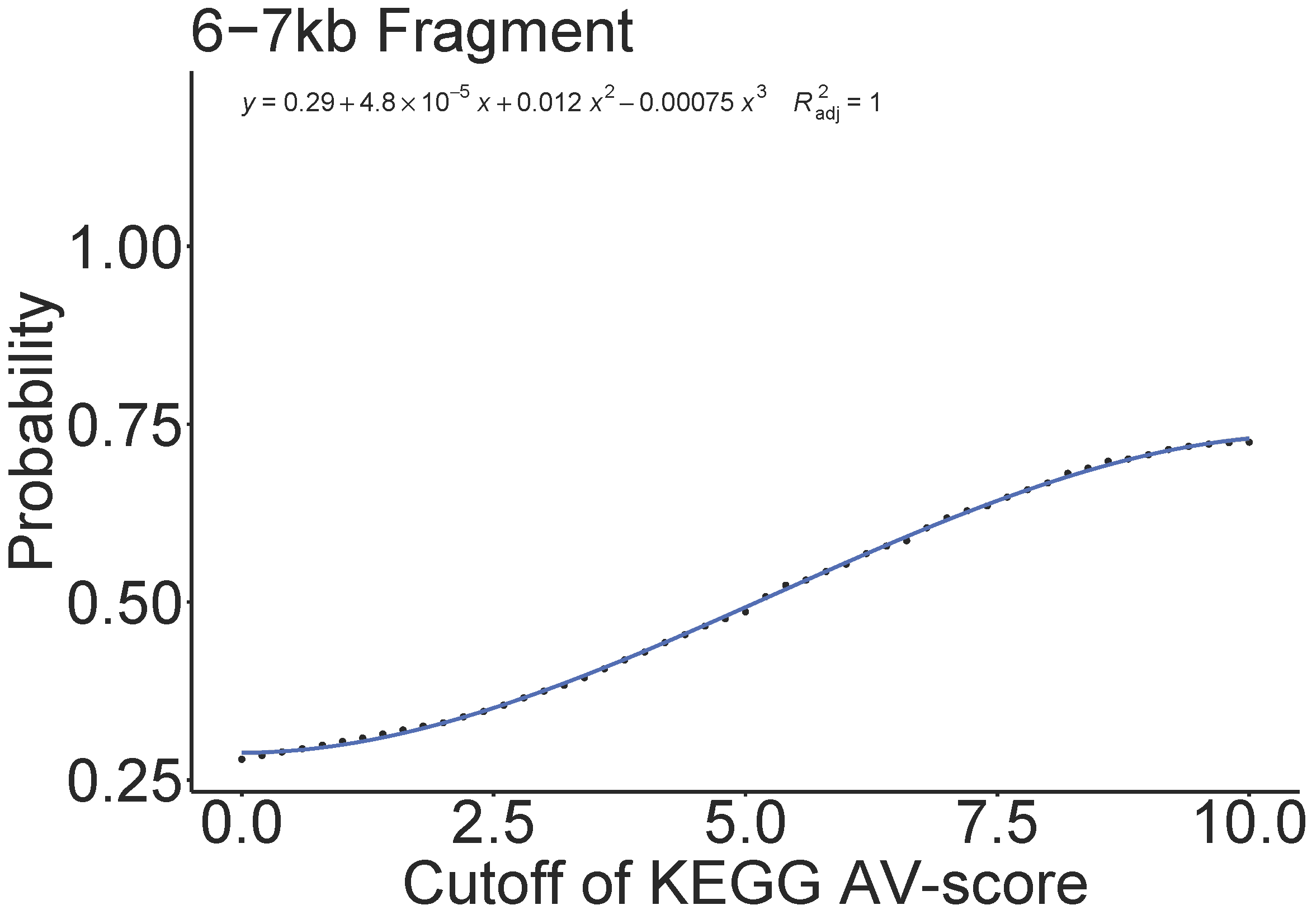

**Fig. S4.** Relationship between the cutoffs (see the definition of cutoff in Fig. S10) of the KEGG AV-score and AV_L_-score of whole genomes used in Fig. S2 and S3 and the fraction of viral genomes (here was defined as probability) above certain cutoffs.

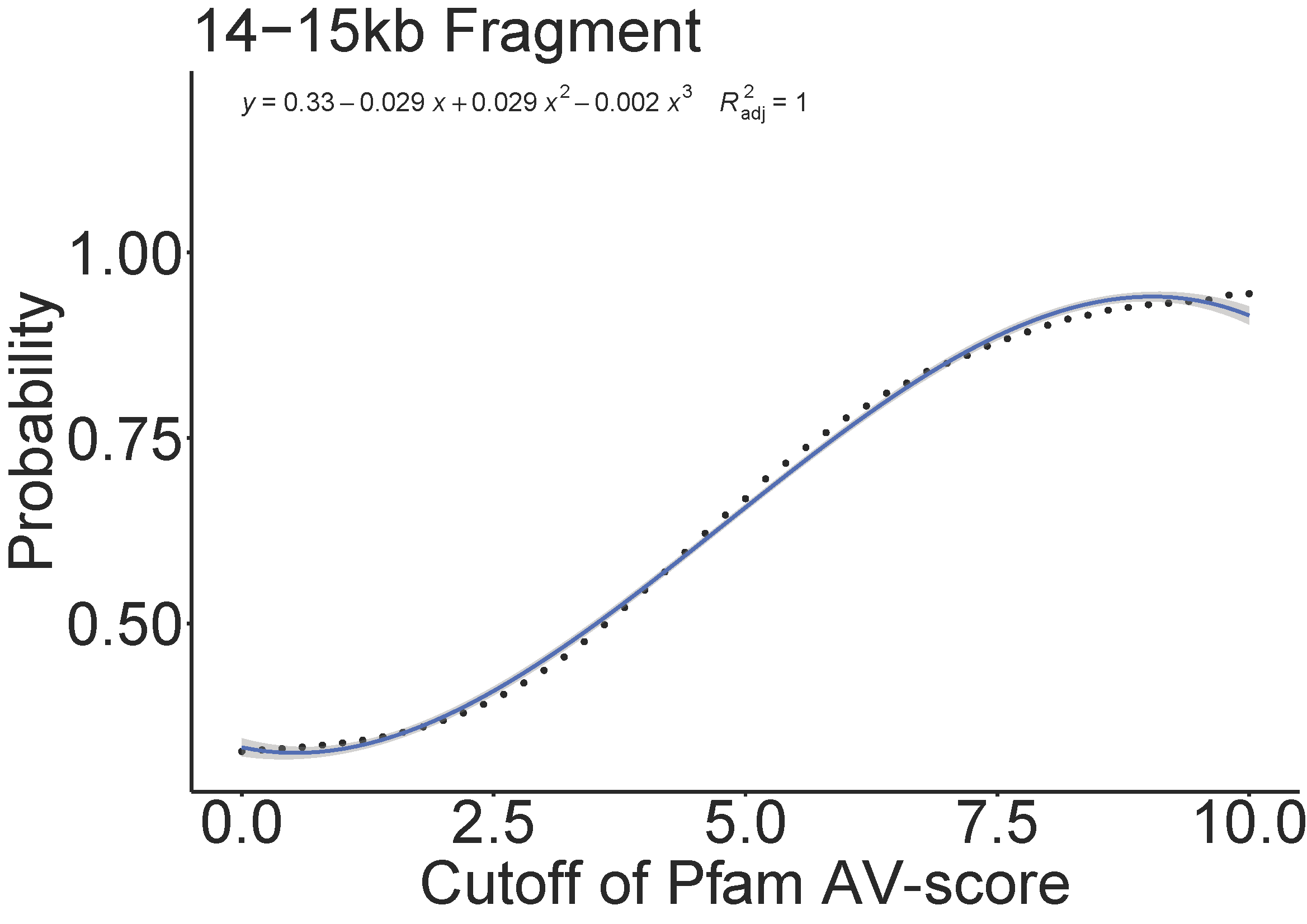

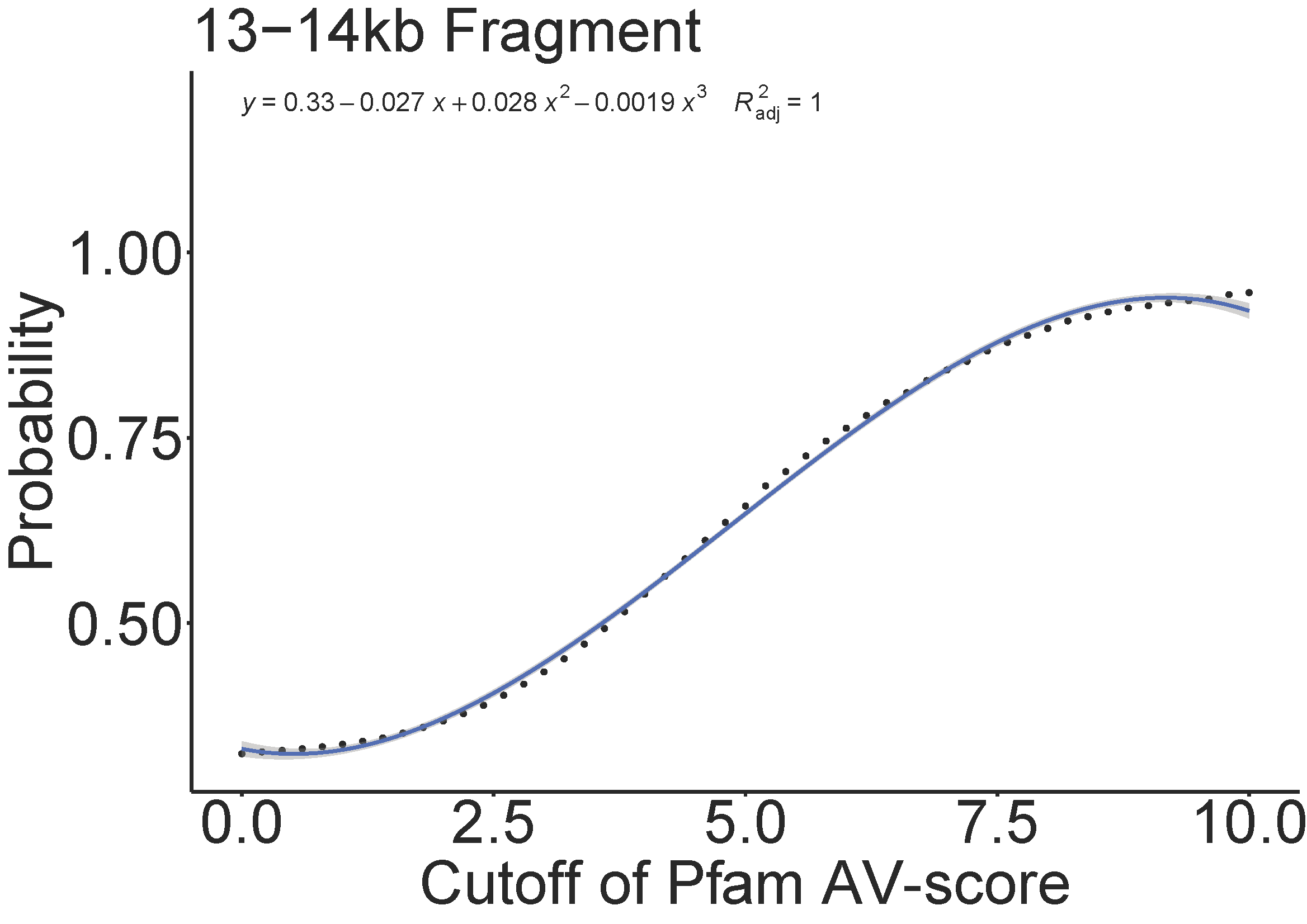

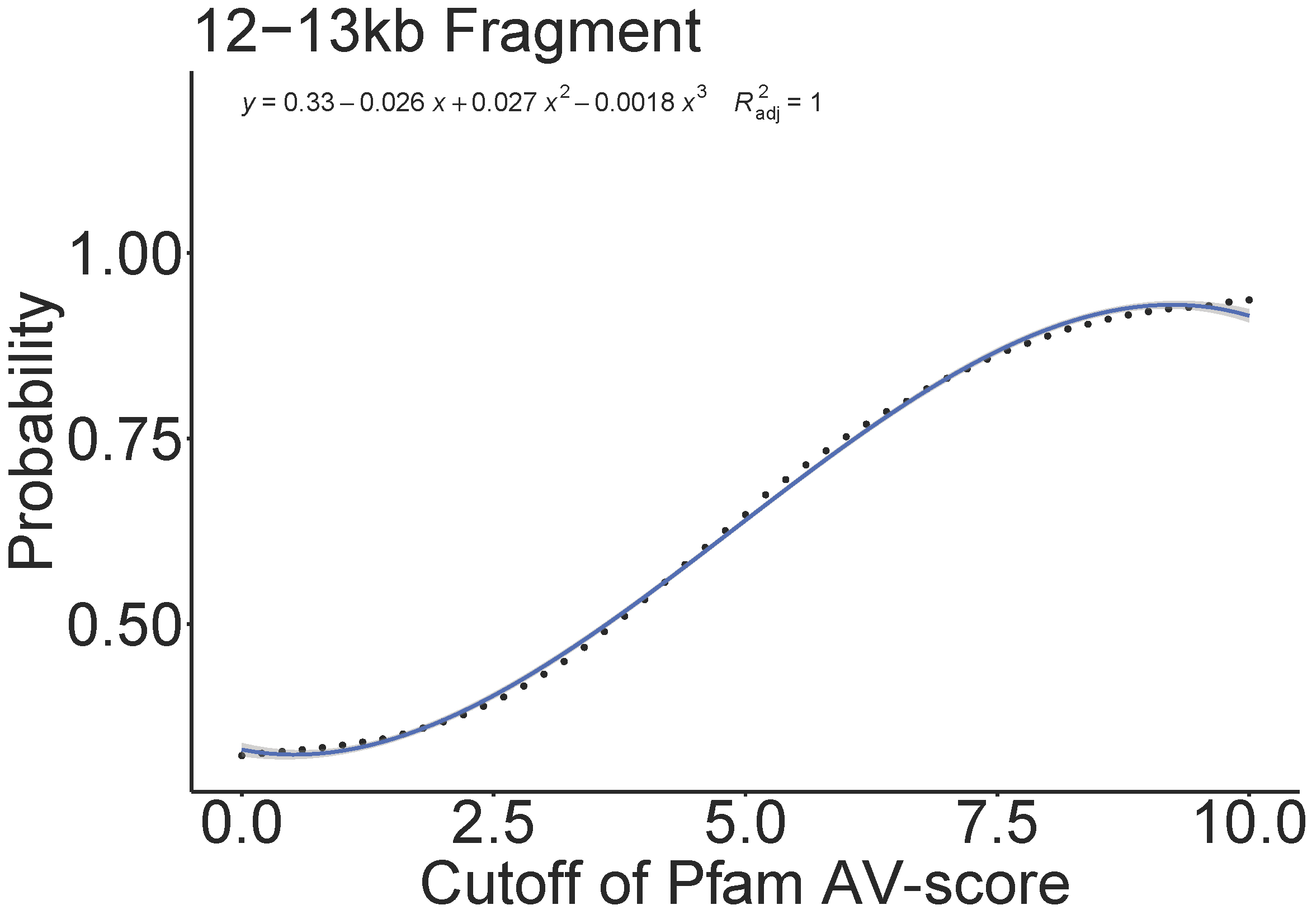

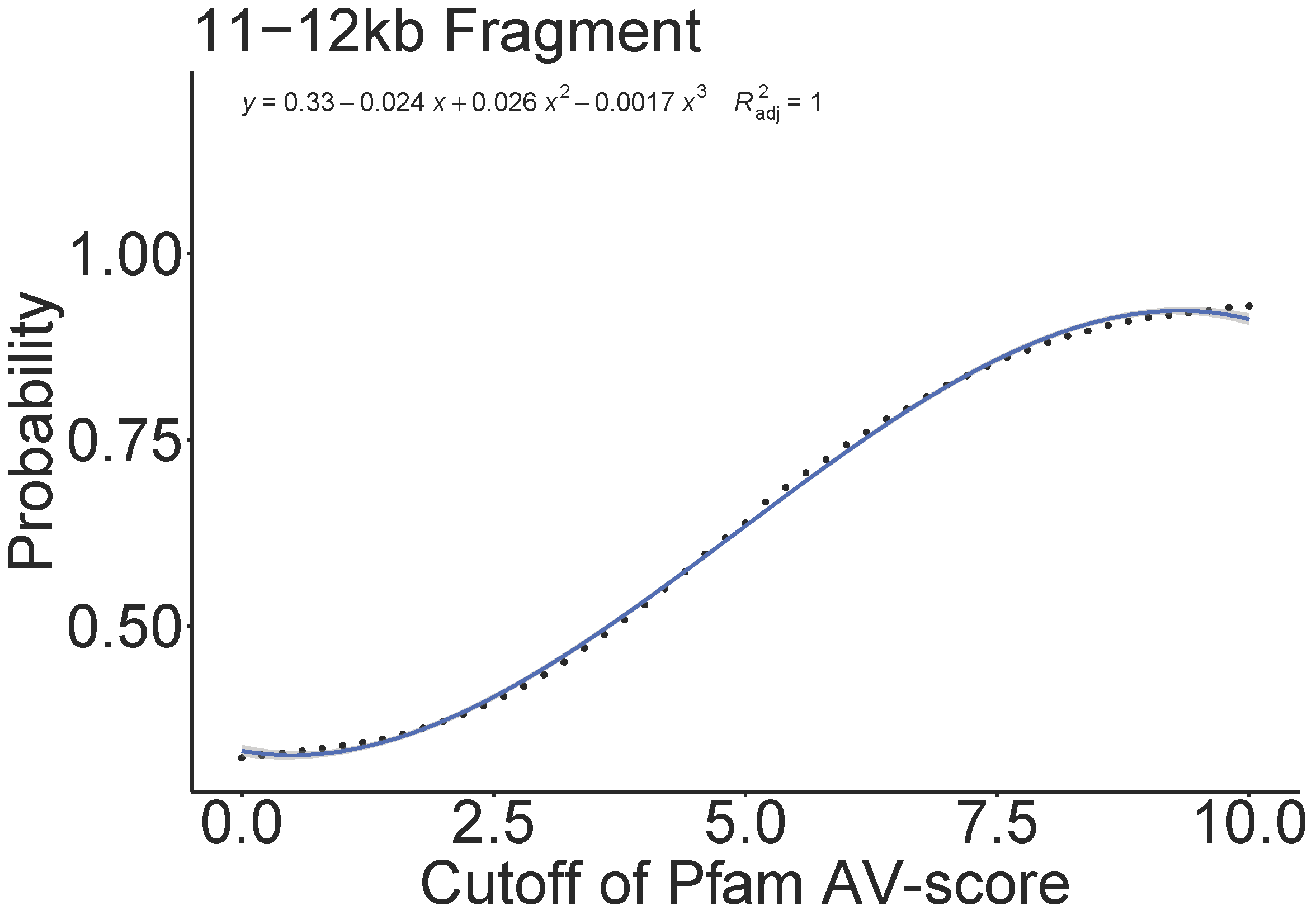

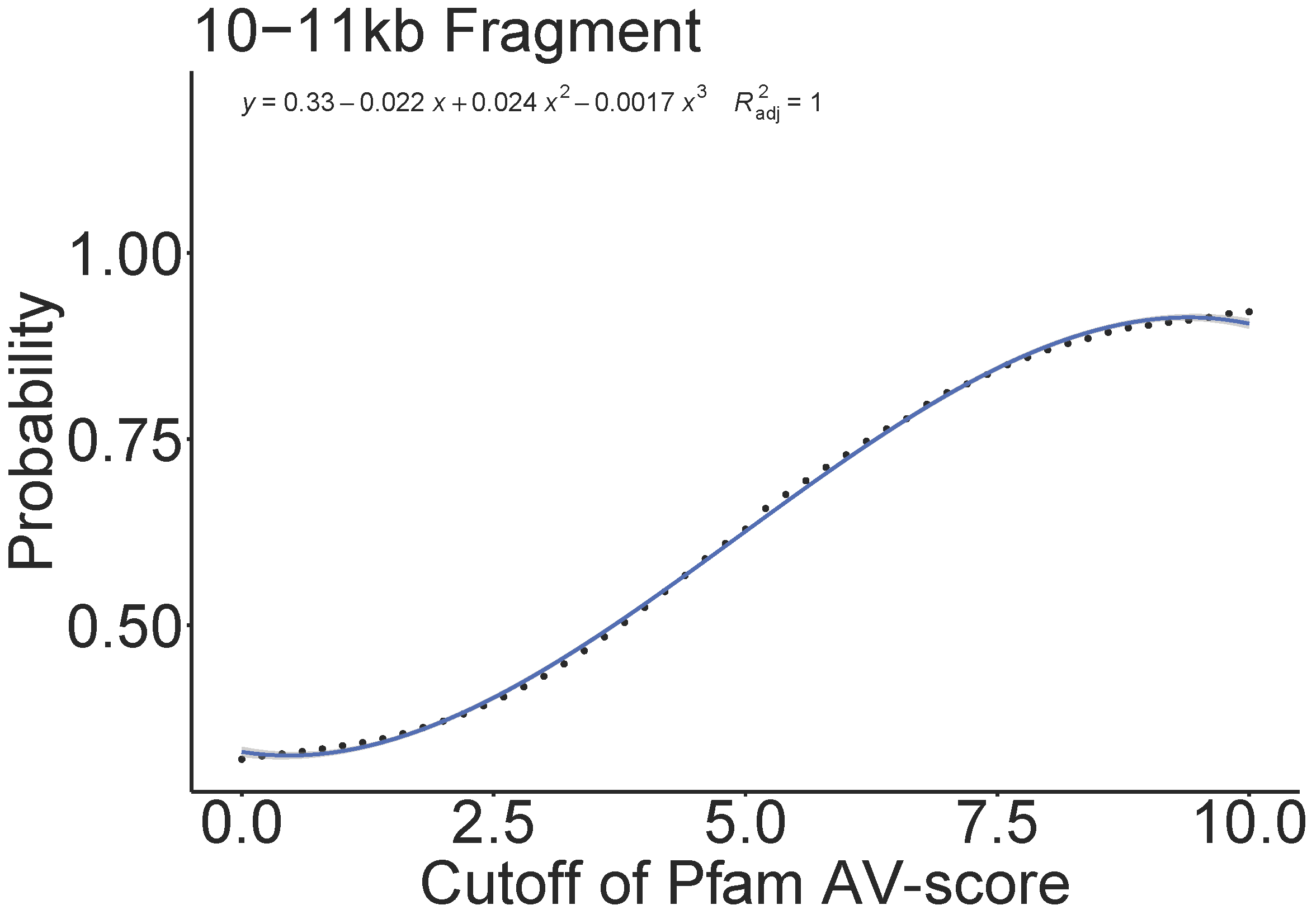

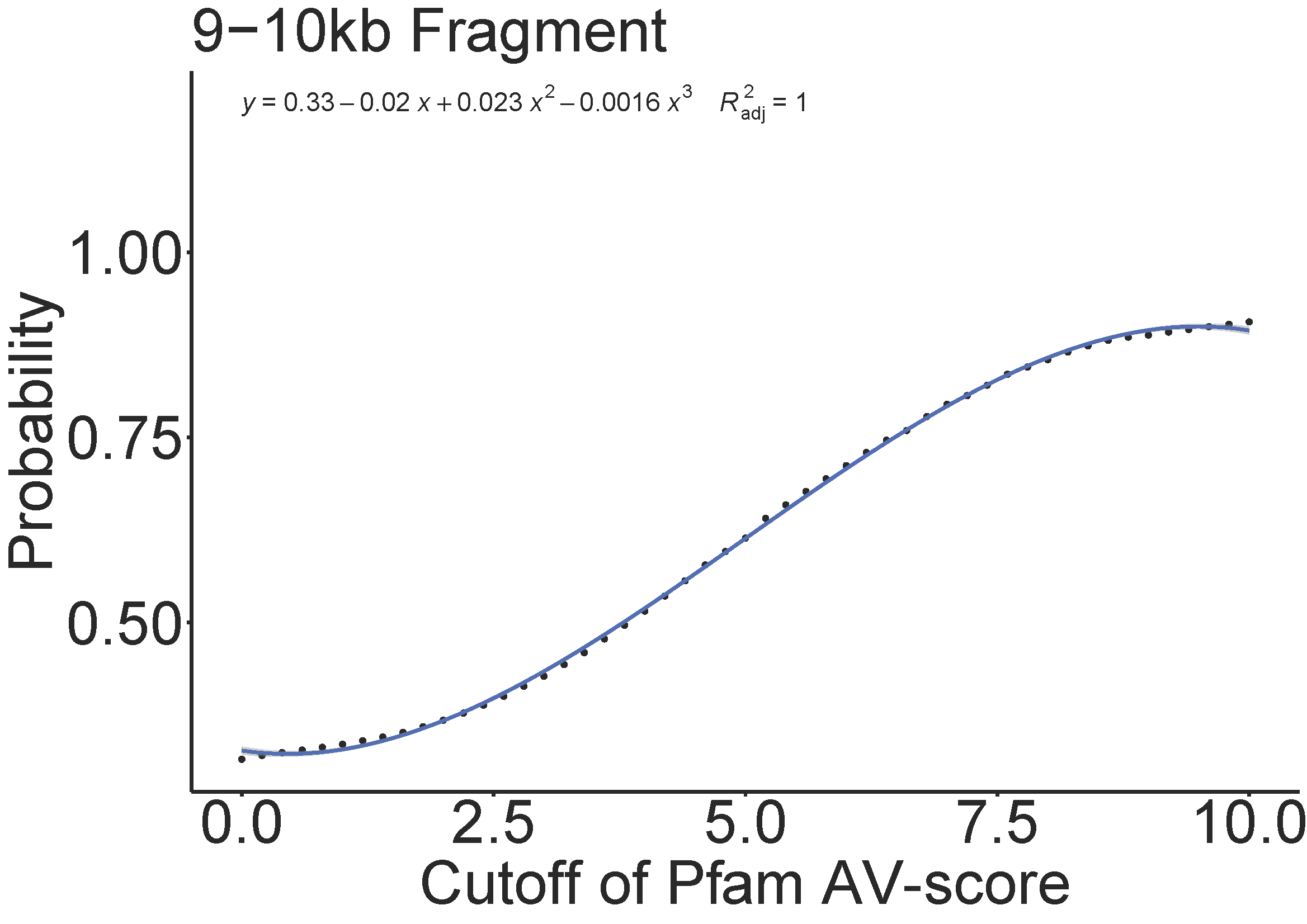

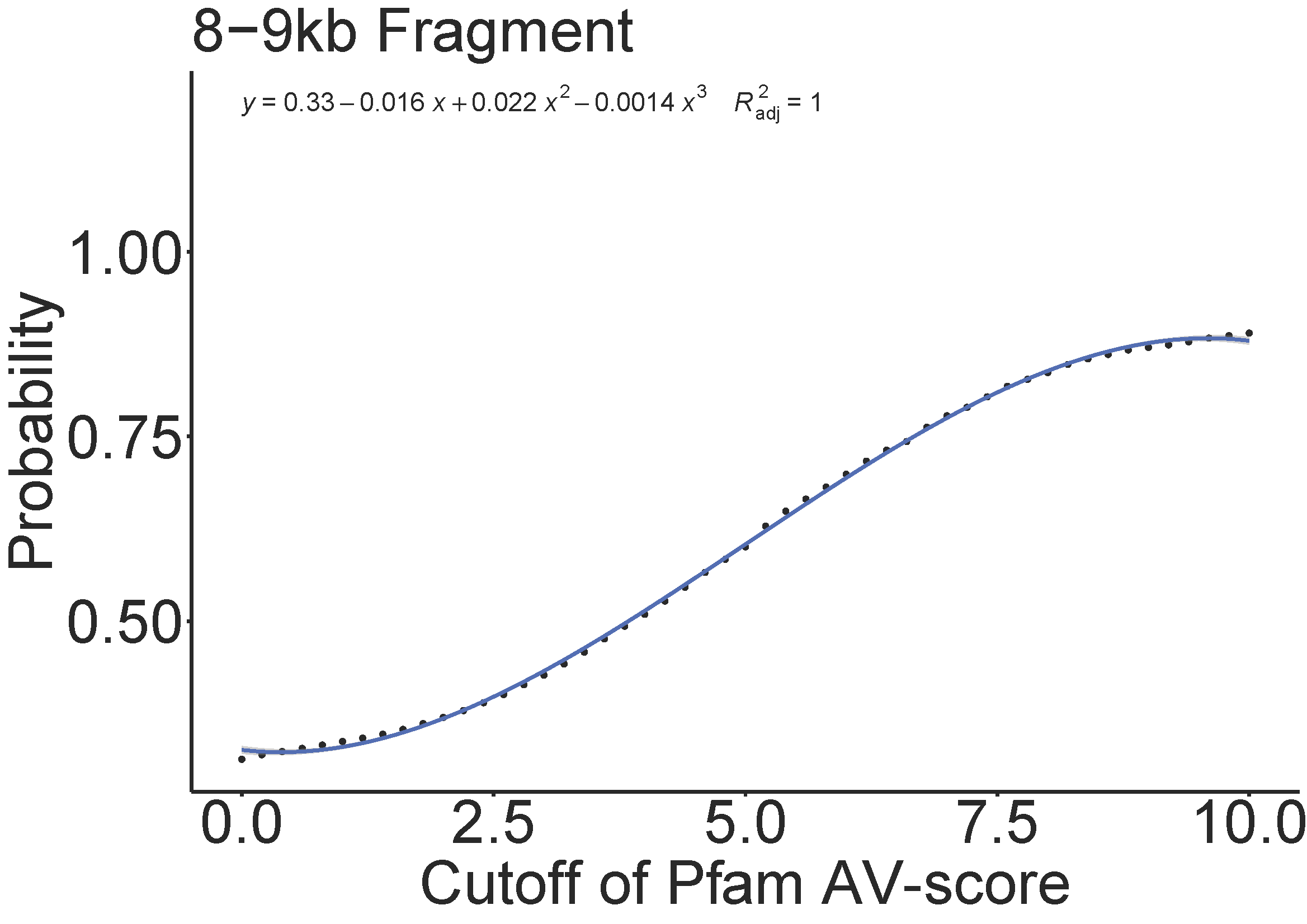

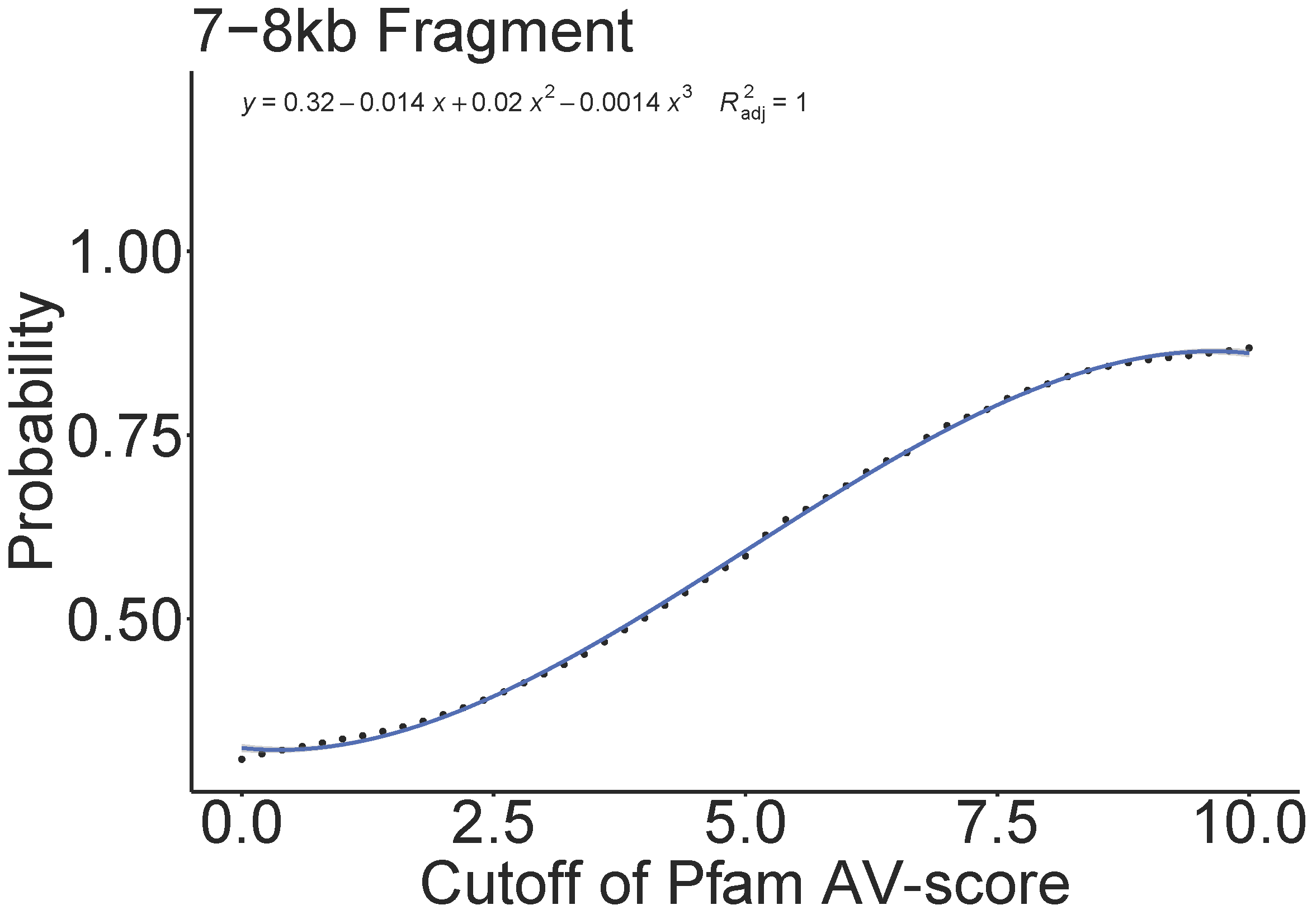

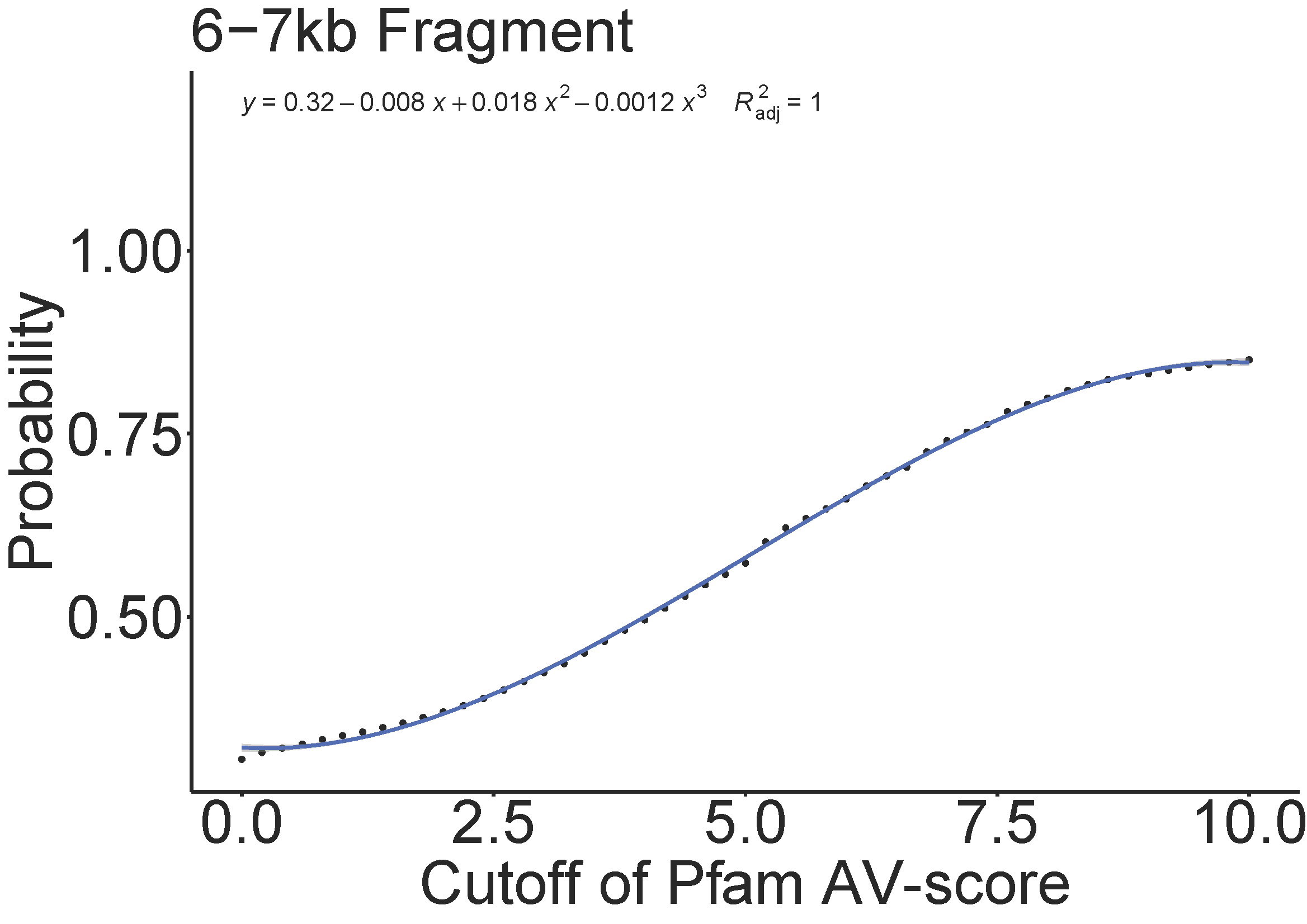

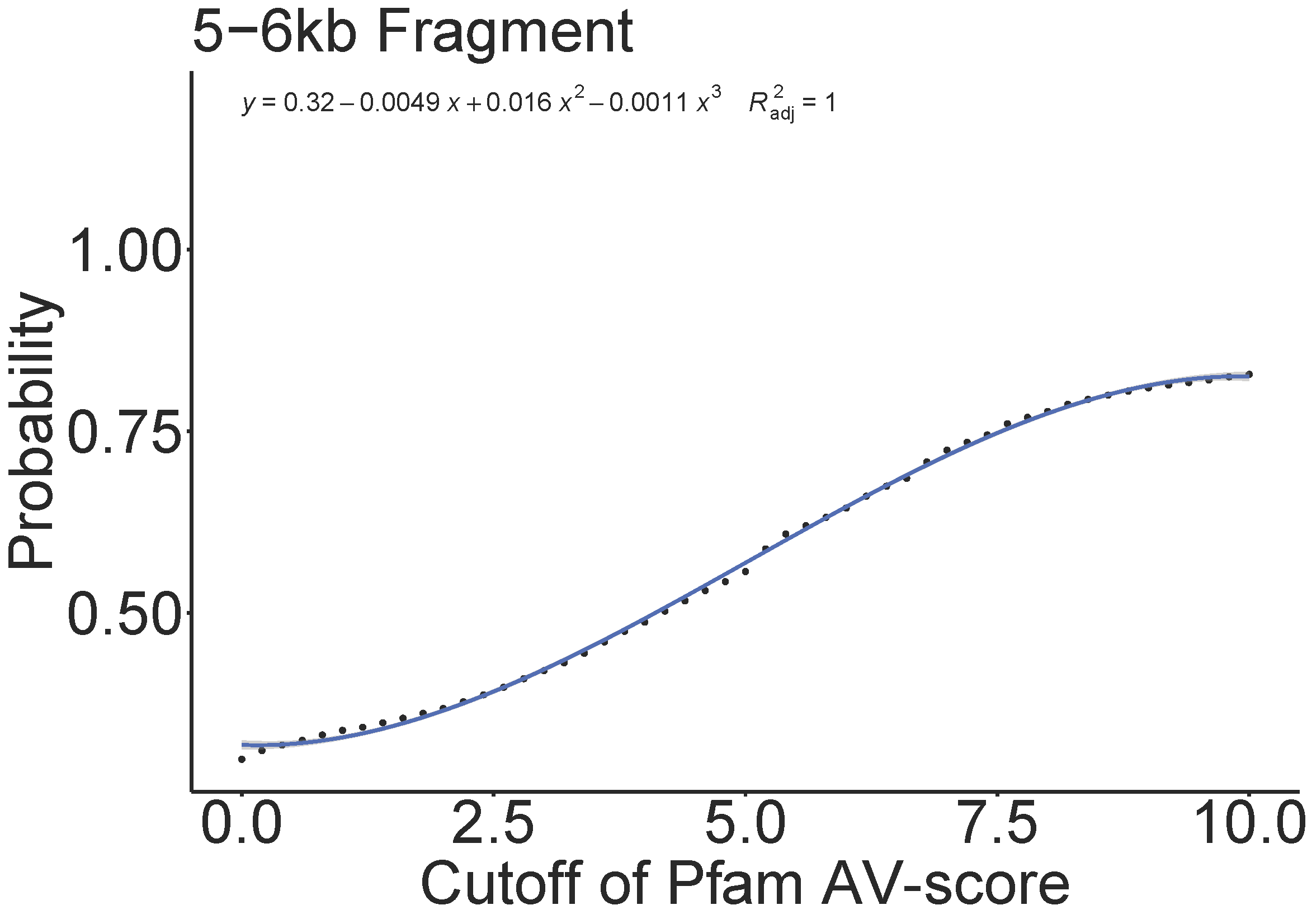

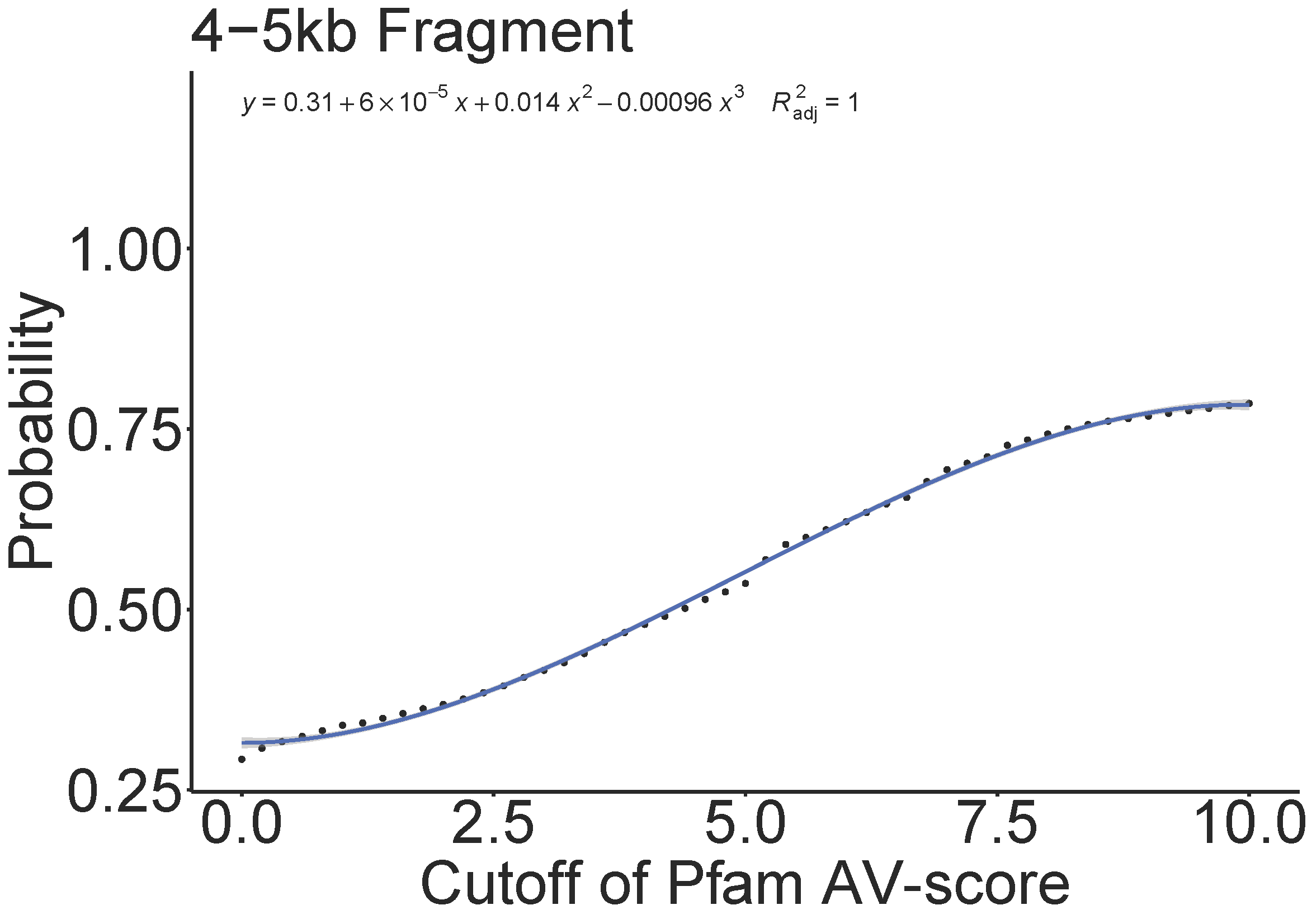

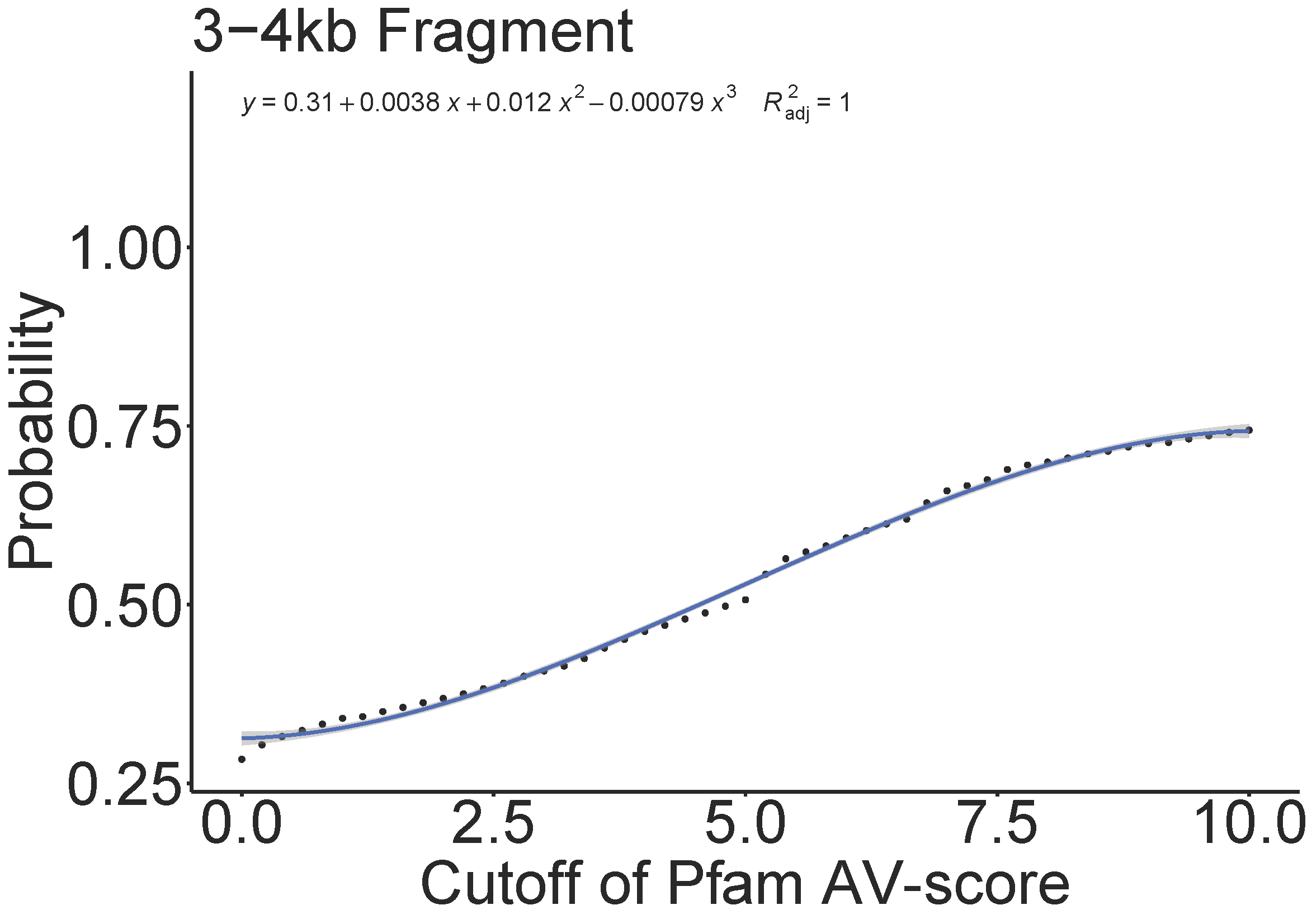

**Fig. S5.** Relationship between the cutoffs (see definition of cutoff in Fig. S10) of the Pfam AV-score and AV_L_-score of whole genomes used in Fig. S2 and S3 and the fraction of viral genomes (here was defined as probability) above certain cutoffs.

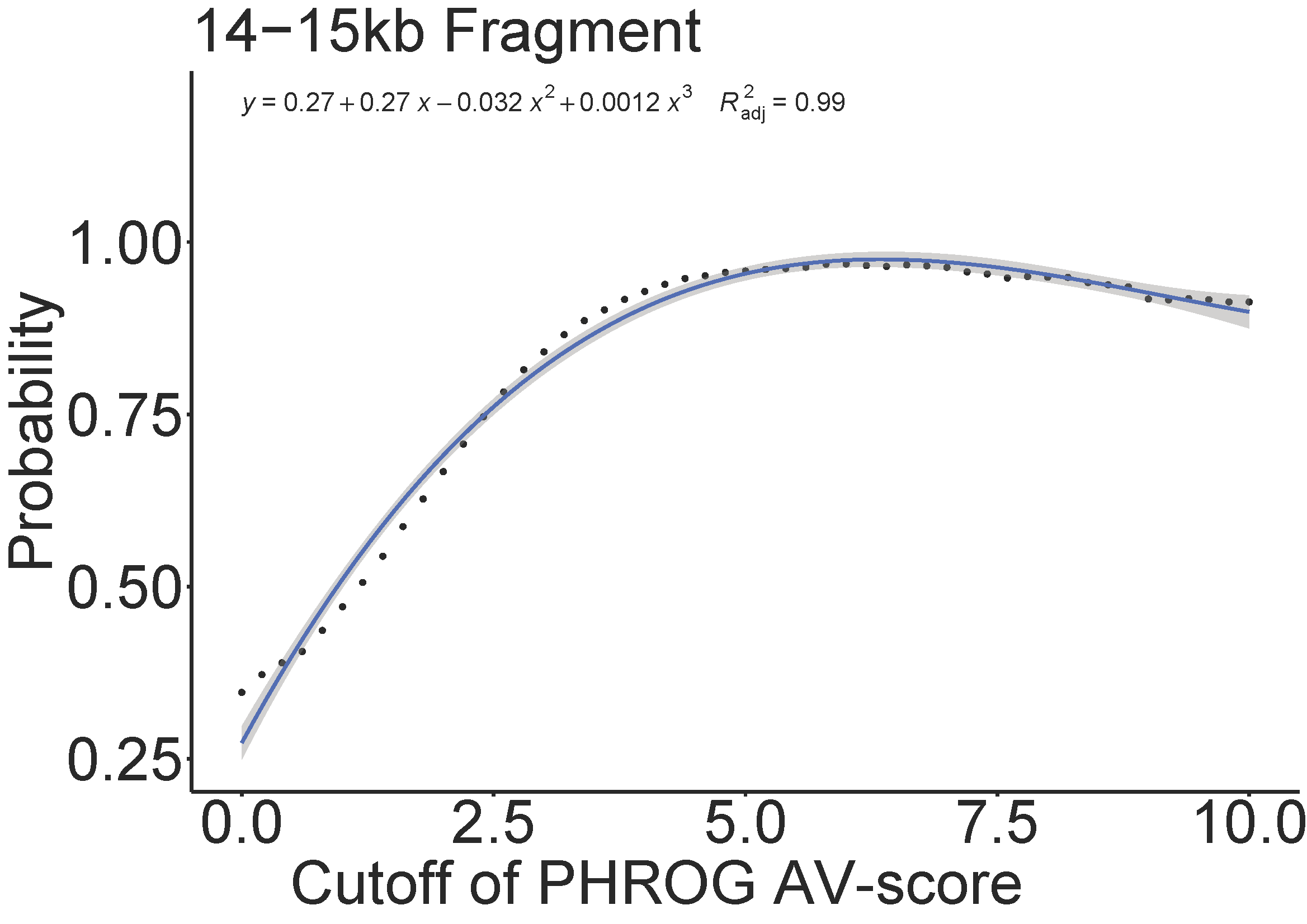

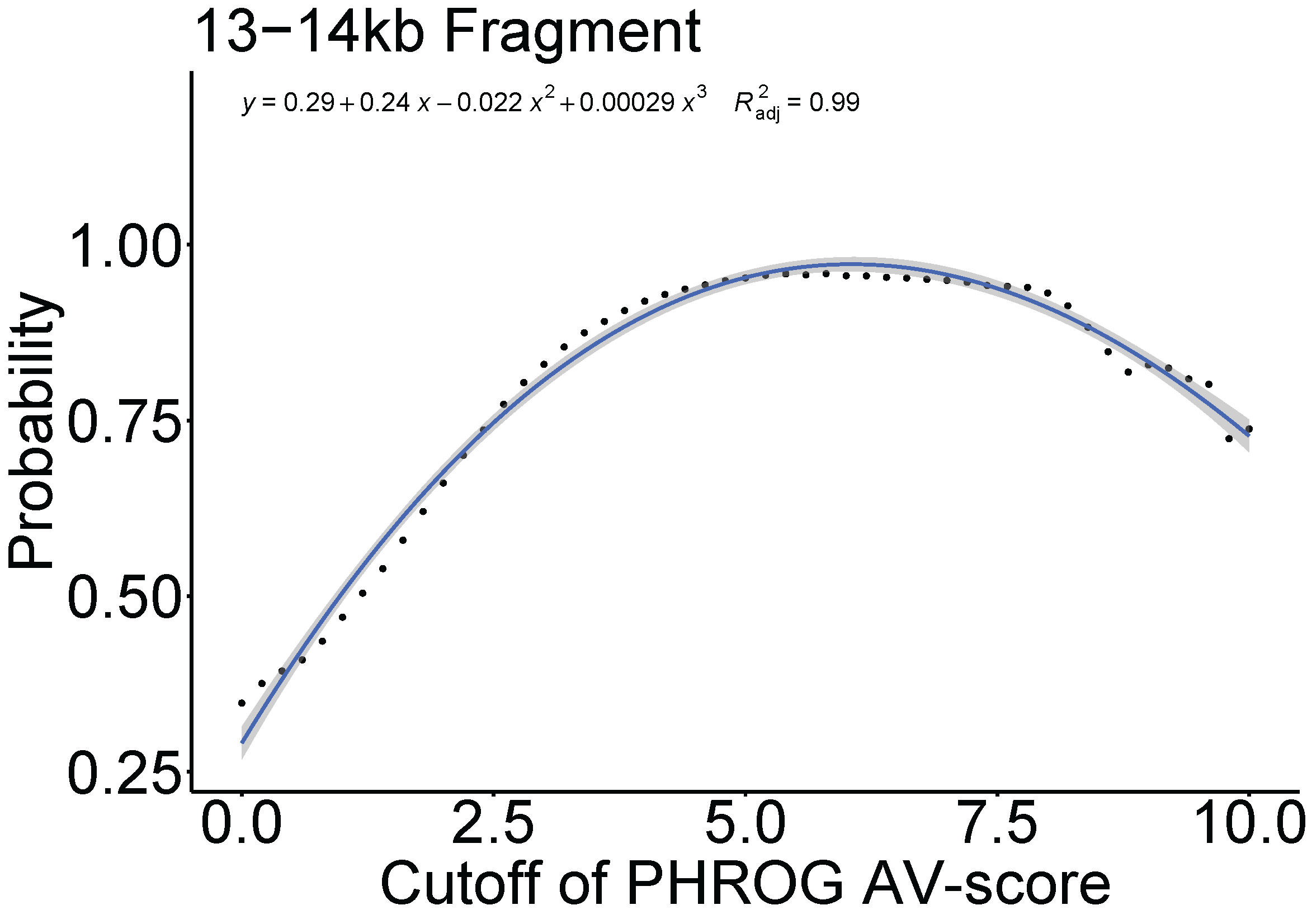

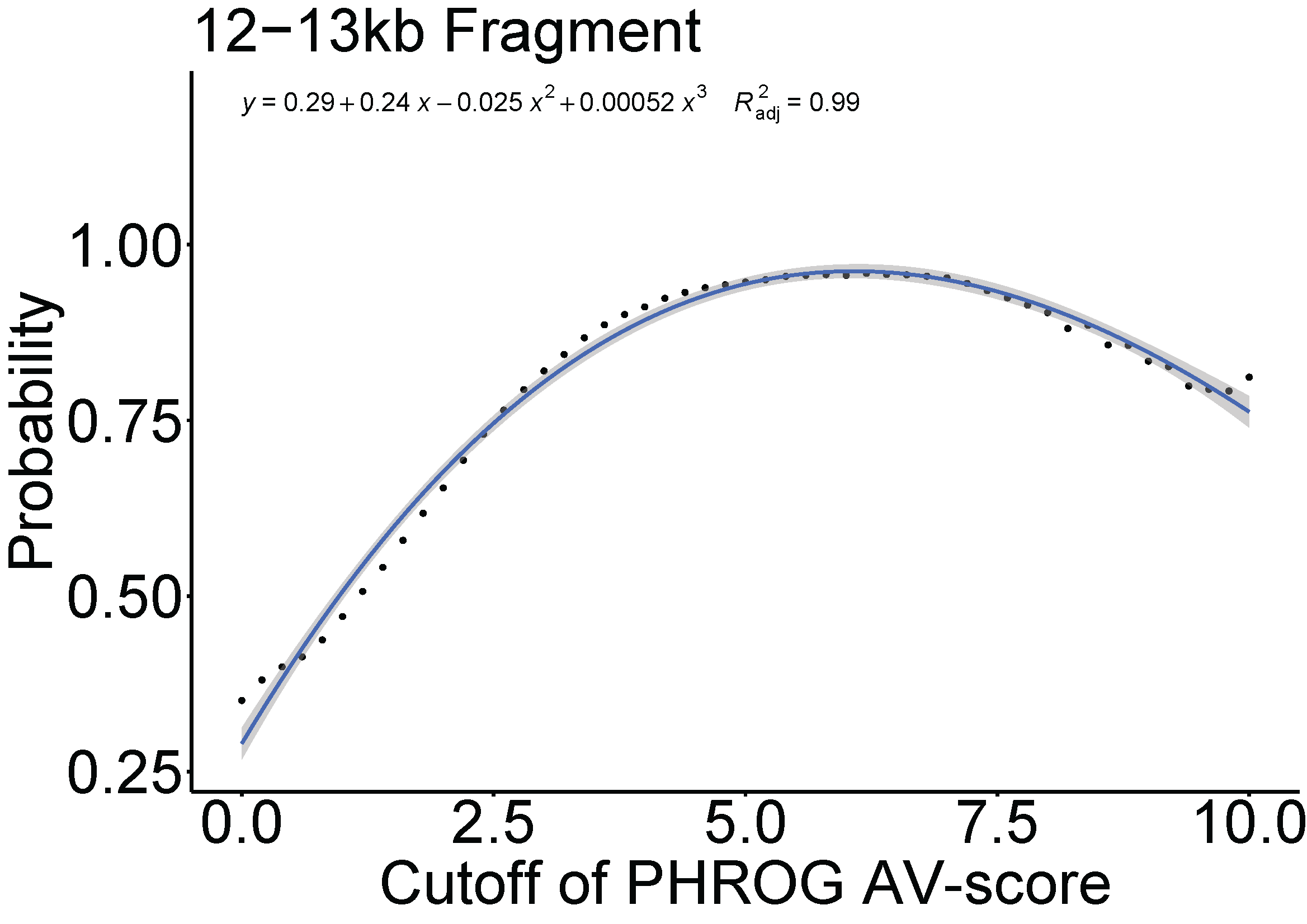

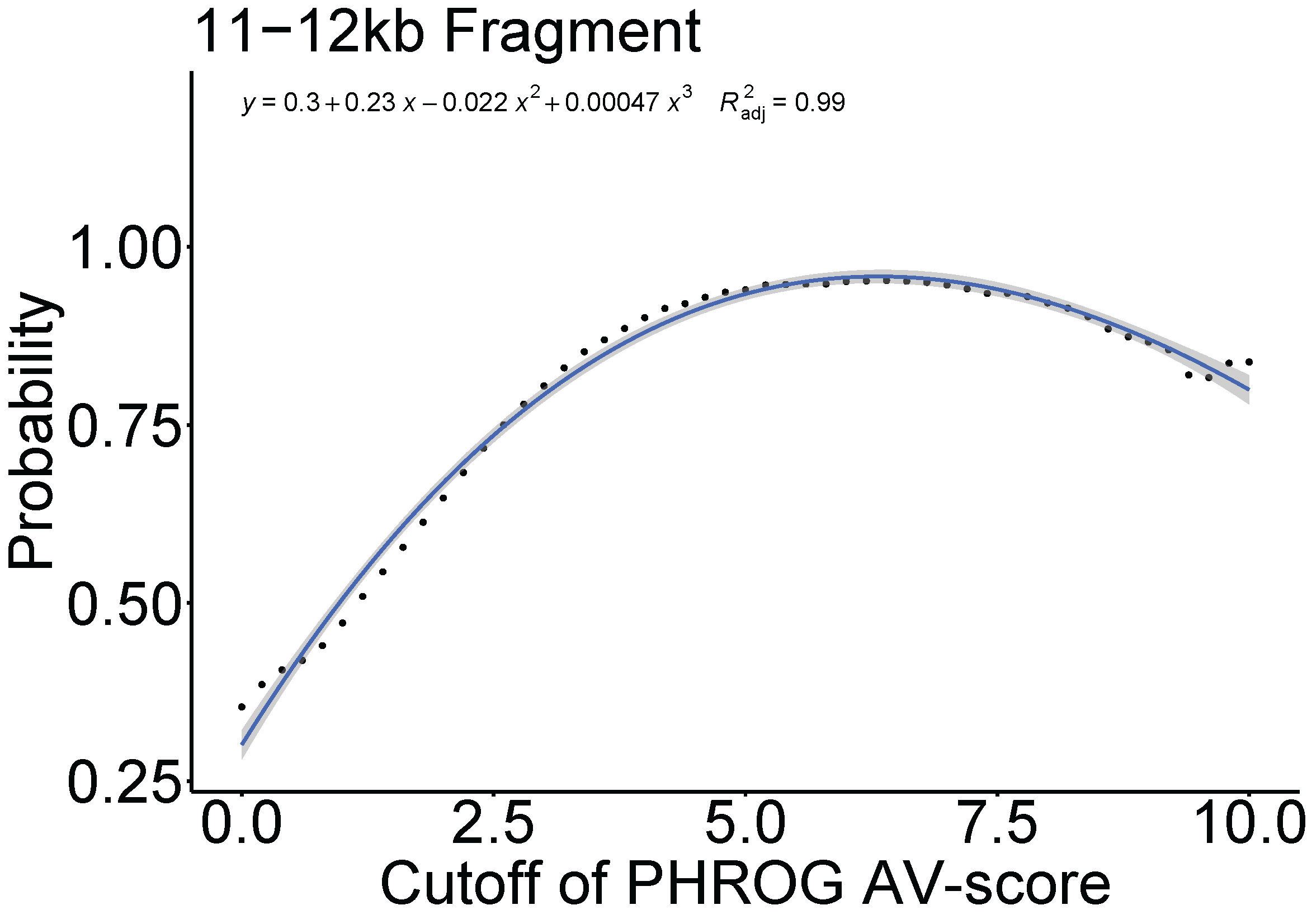

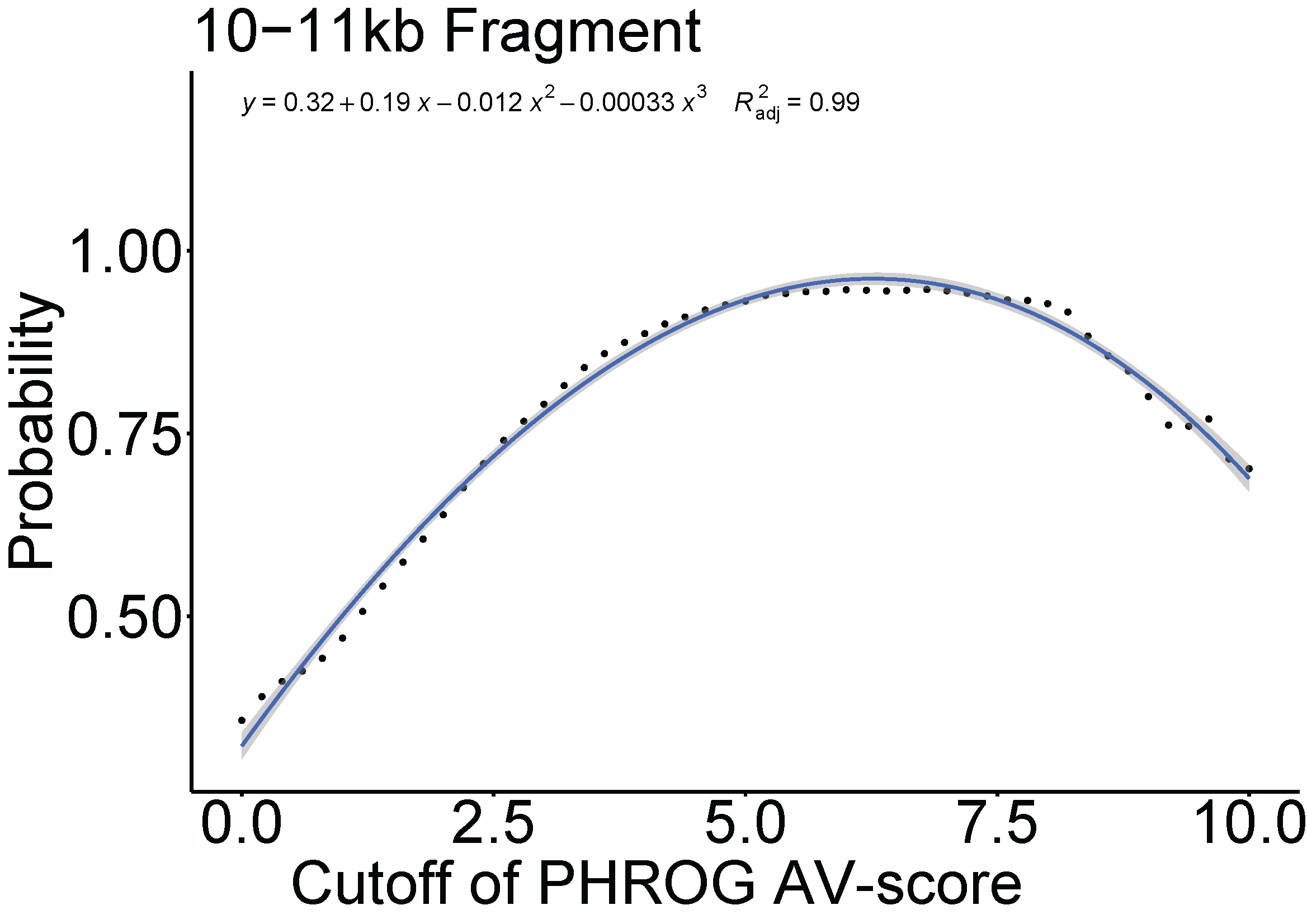

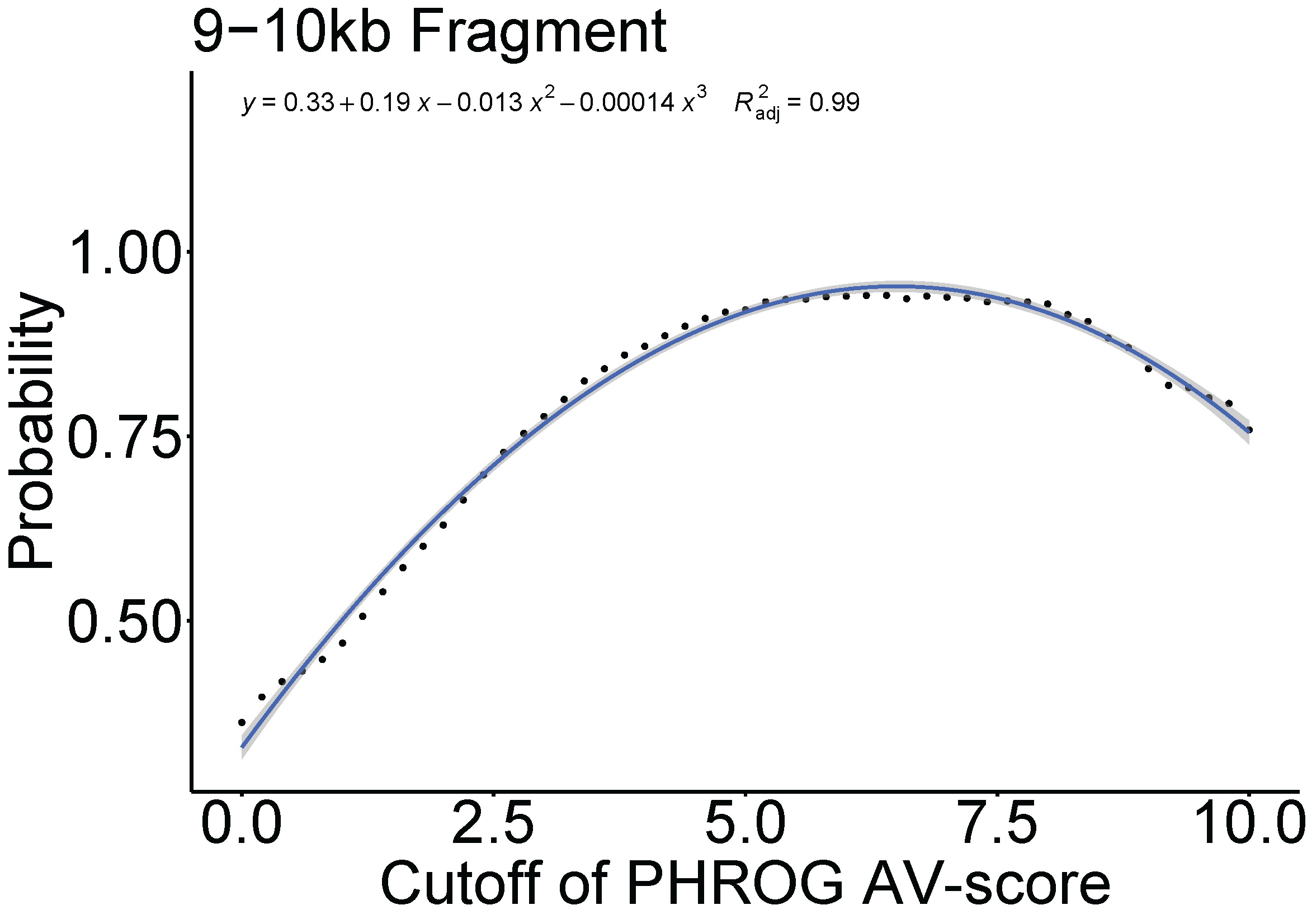

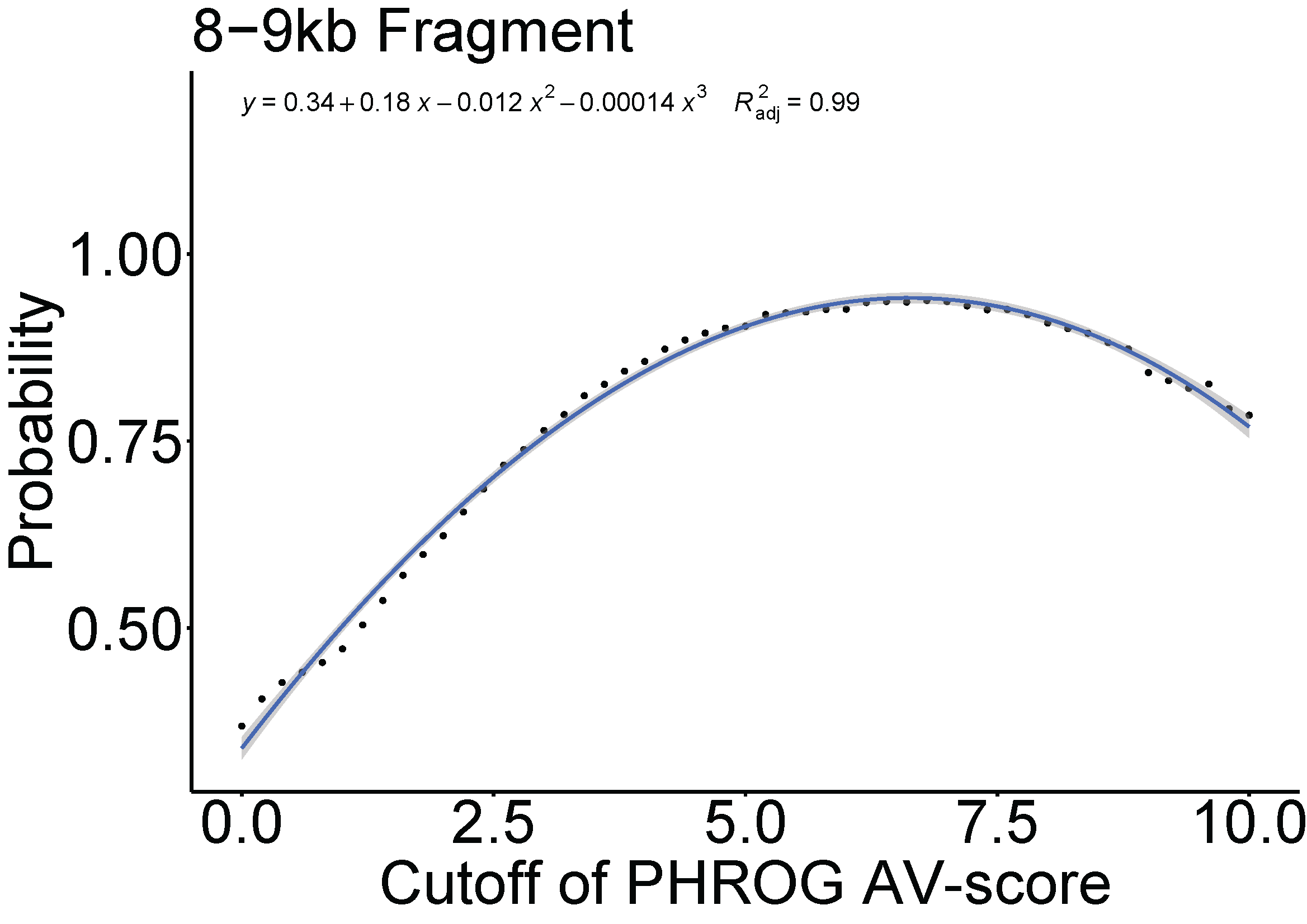

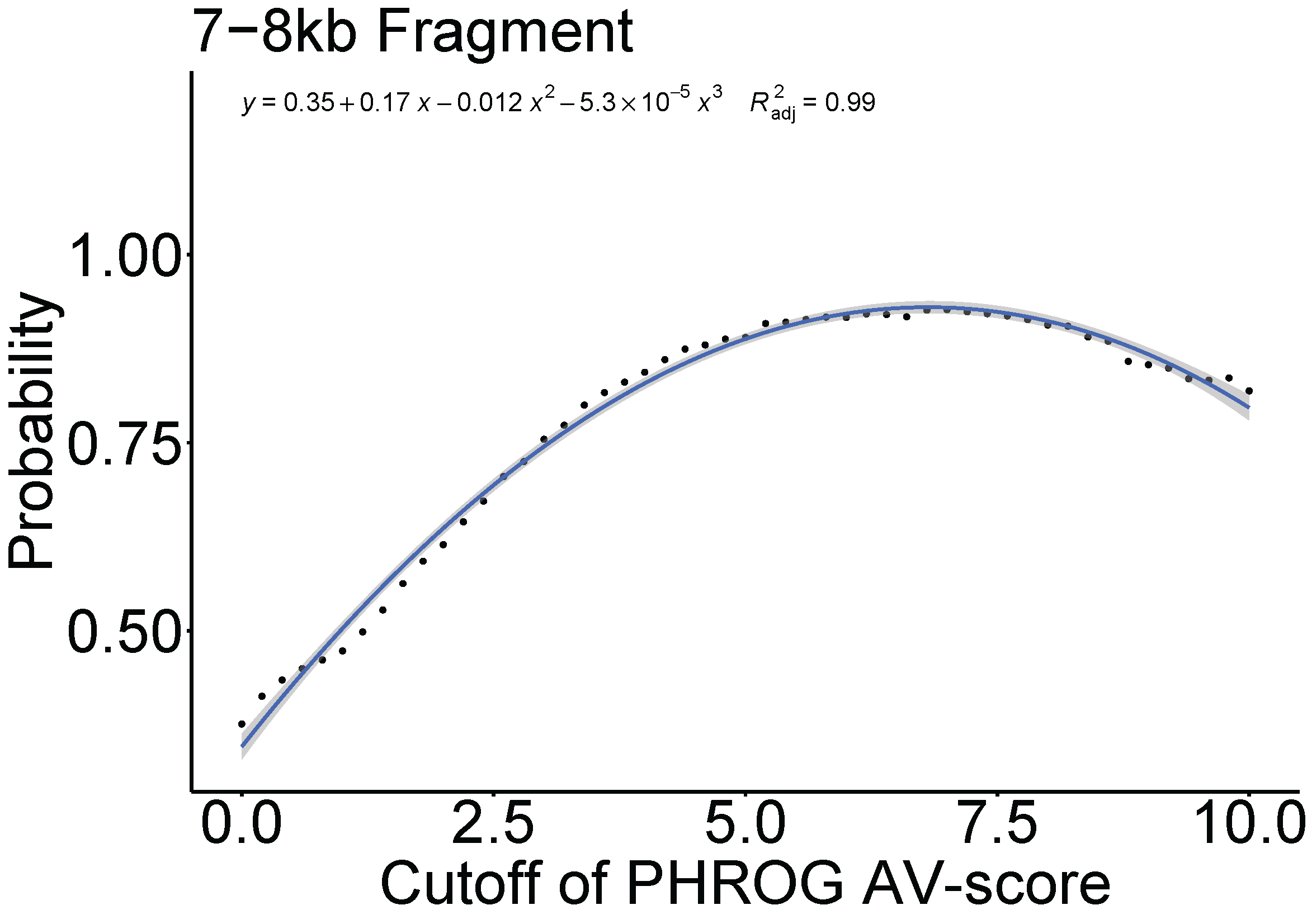

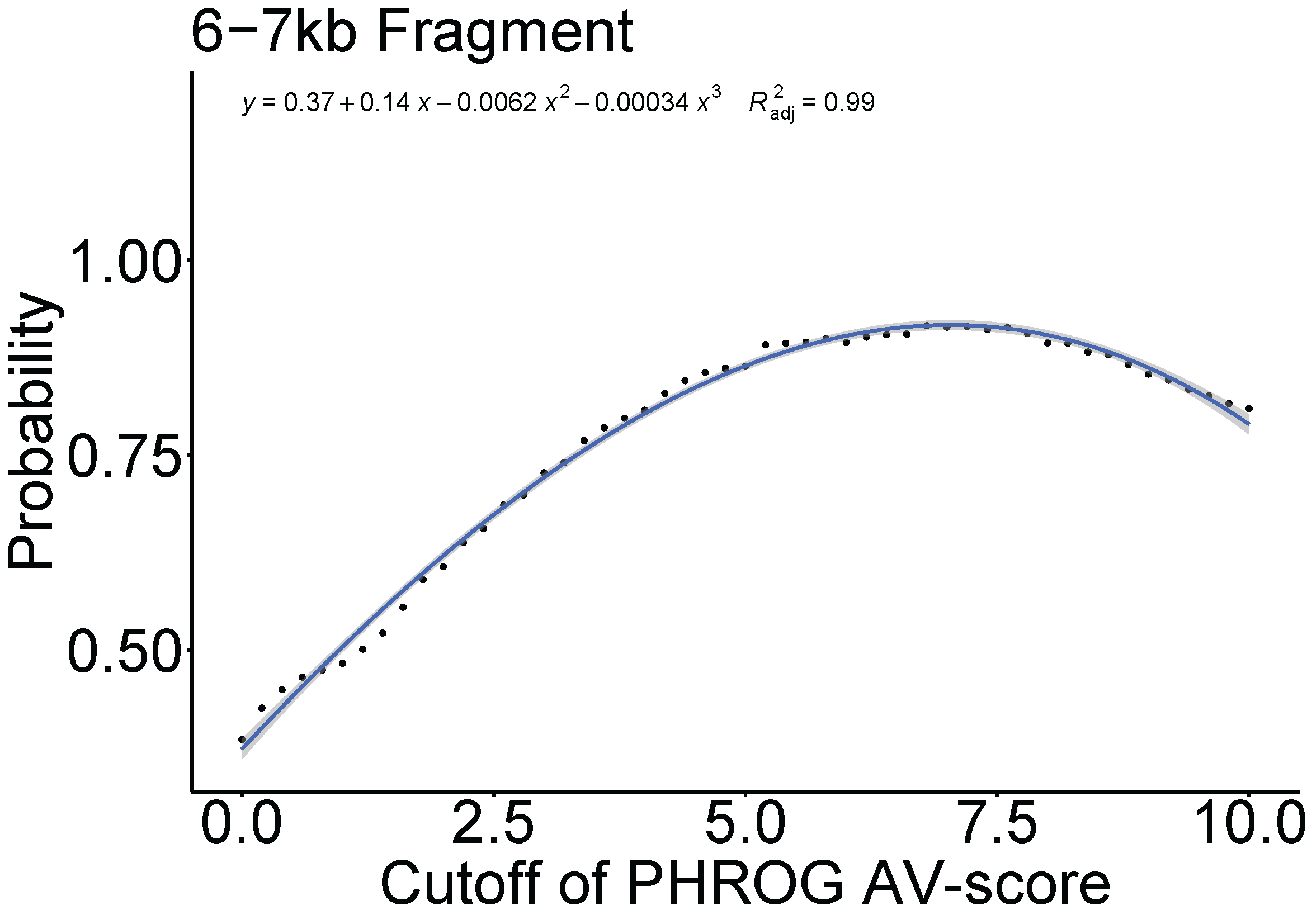

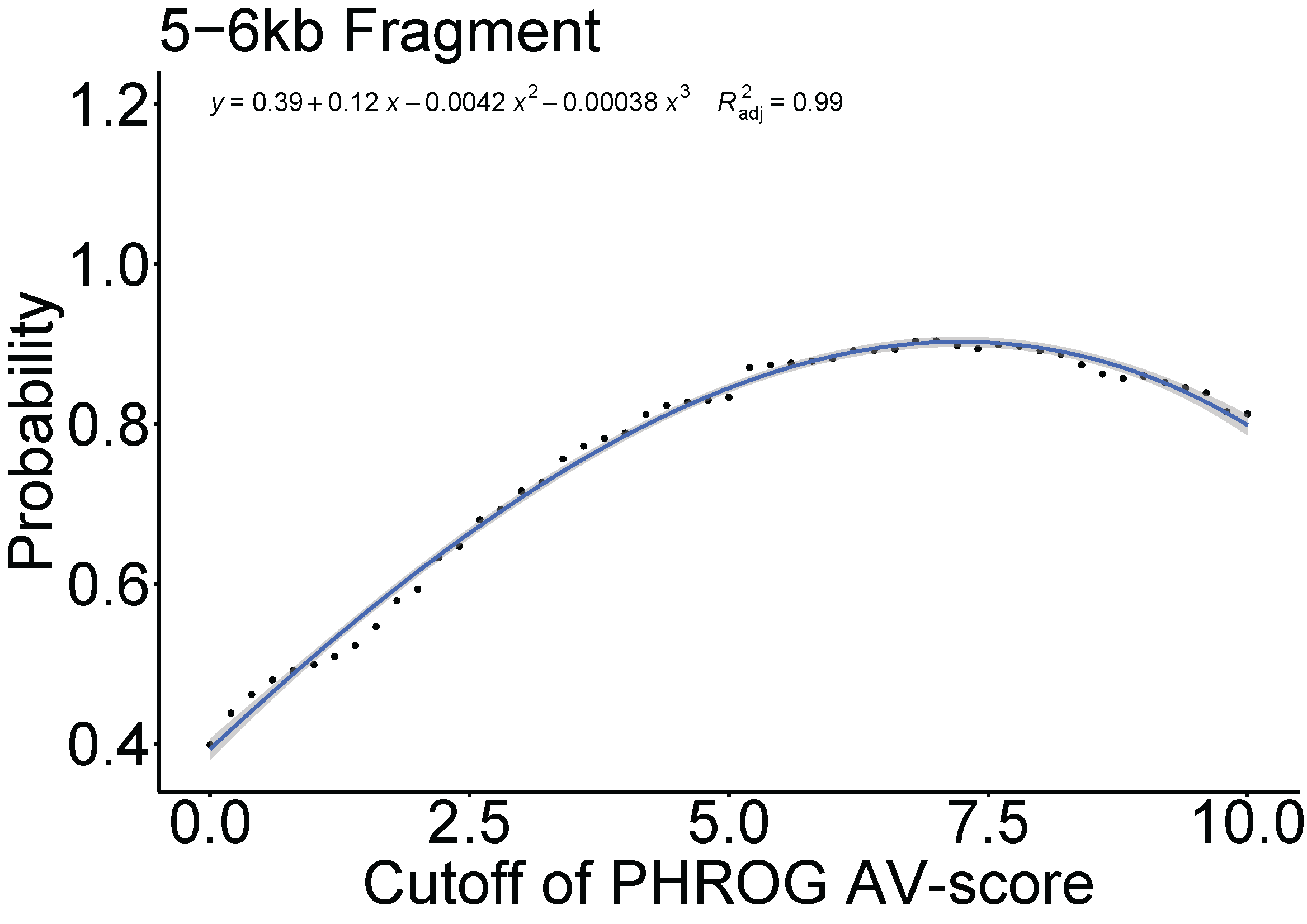

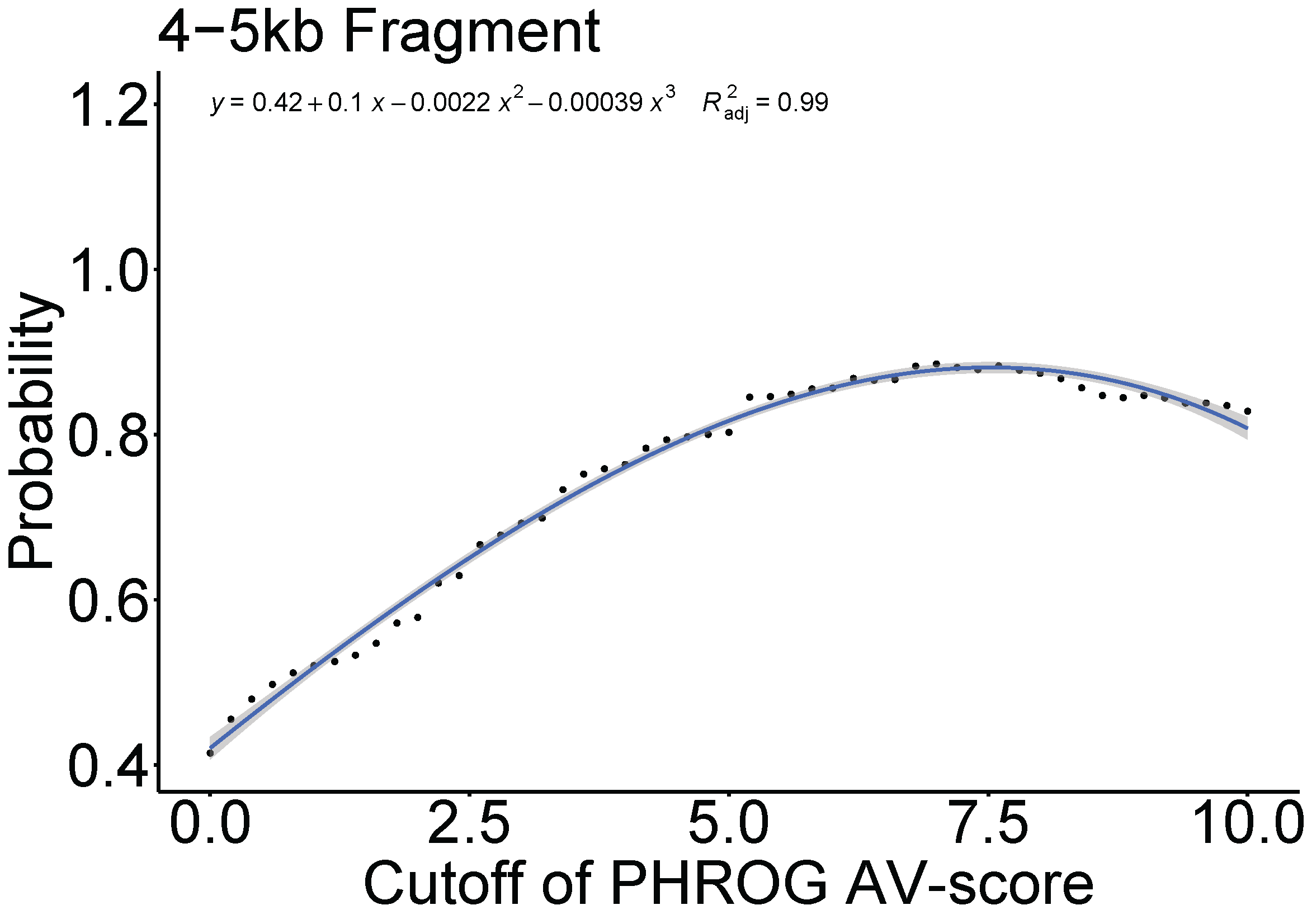

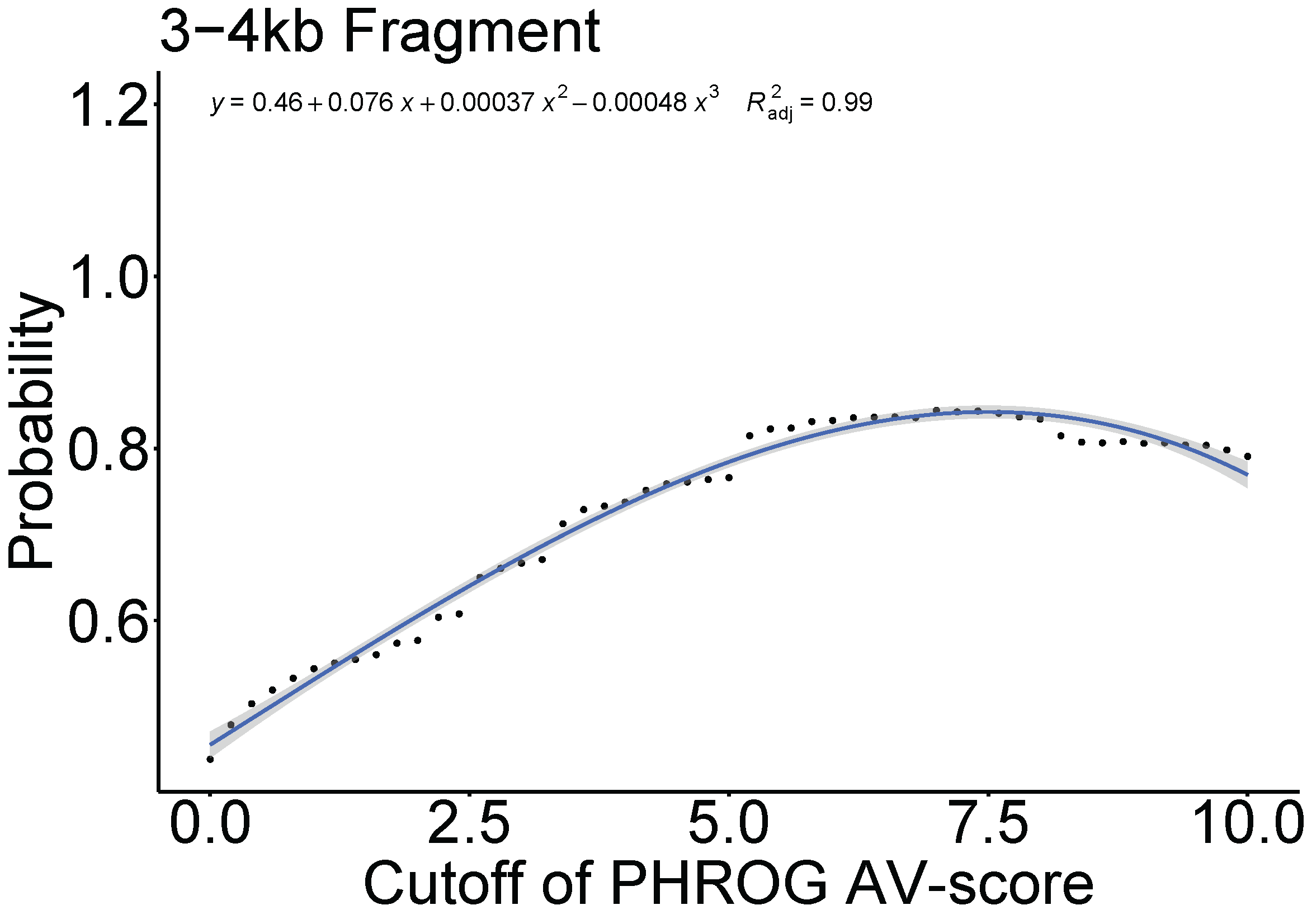

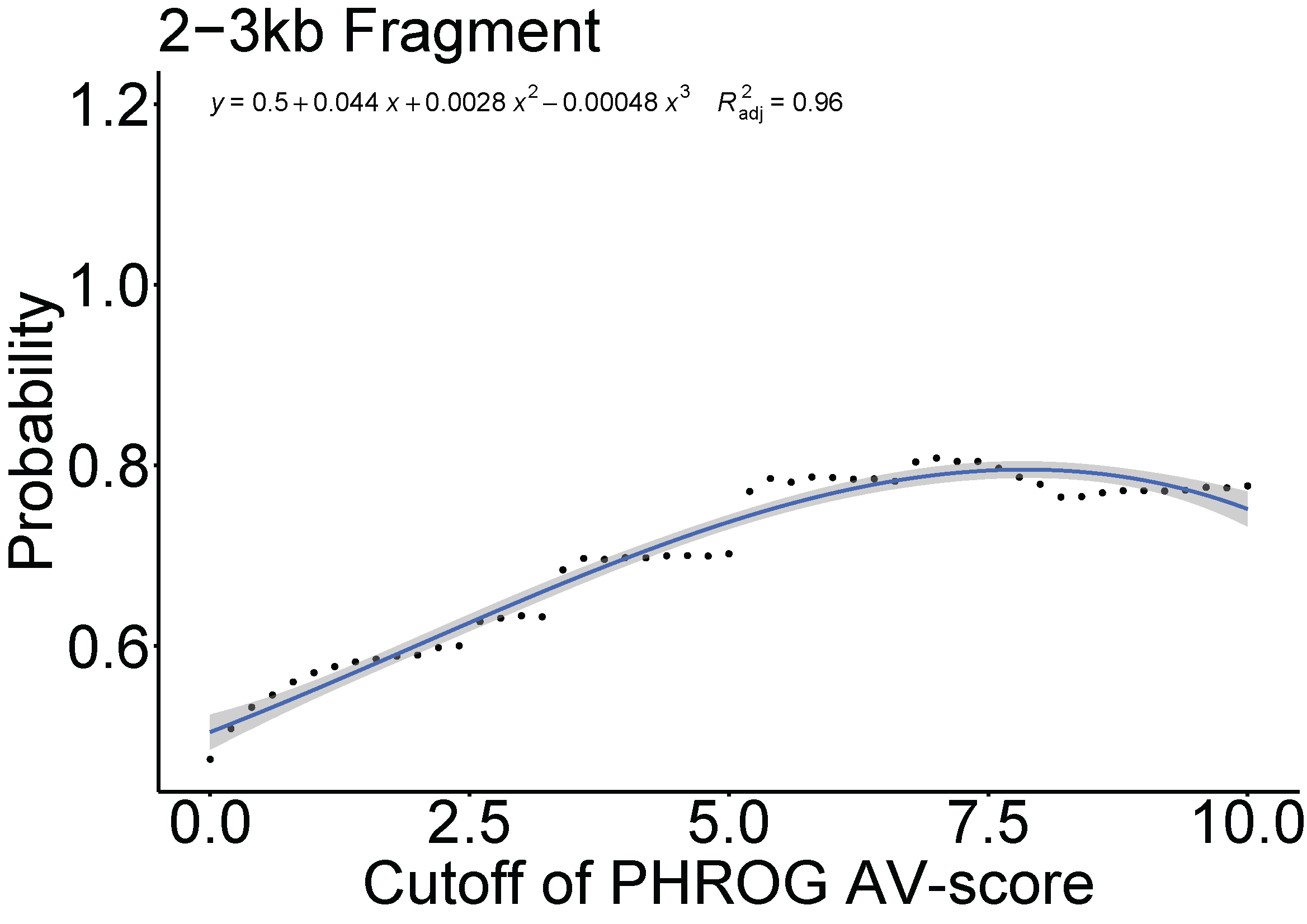

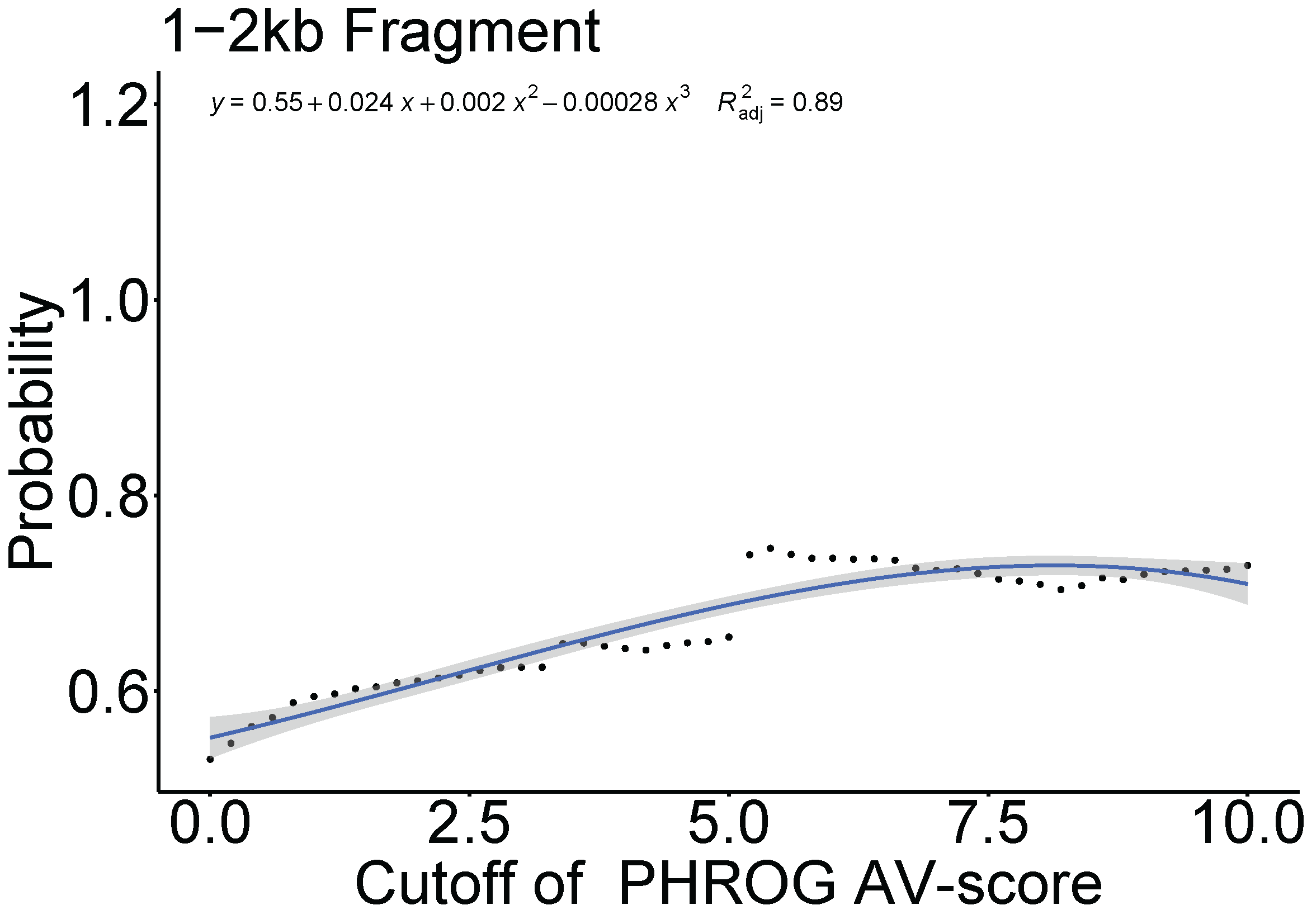

**Fig. S6.** Relationship between the cutoffs (see the definition of cutoff in Fig. S10) of the PHROG AV-score and AV_L_-score of whole genomes used in Fig. S2 and S3 and the fraction of viral genomes (here was defined as probability) above certain cutoffs.

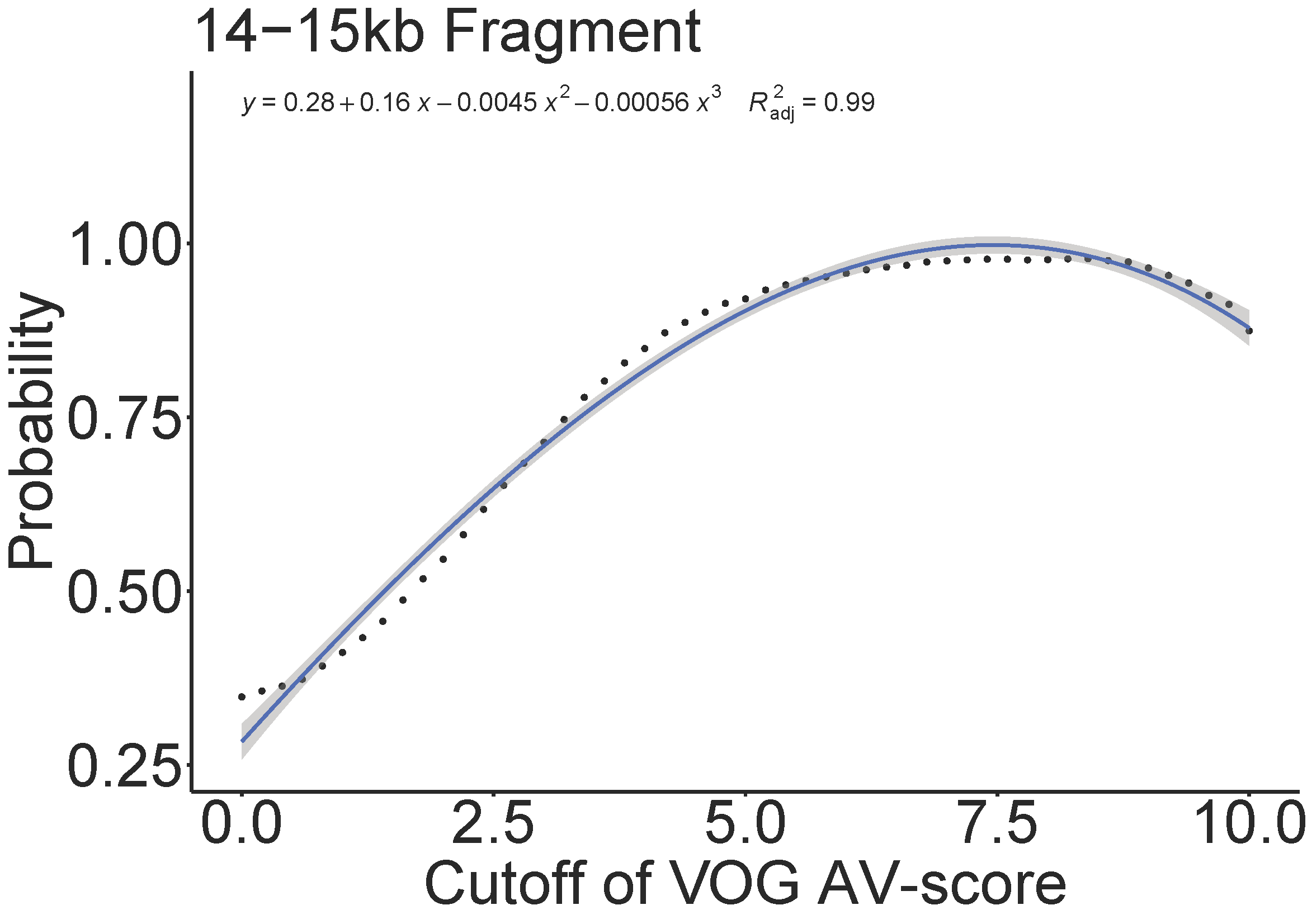

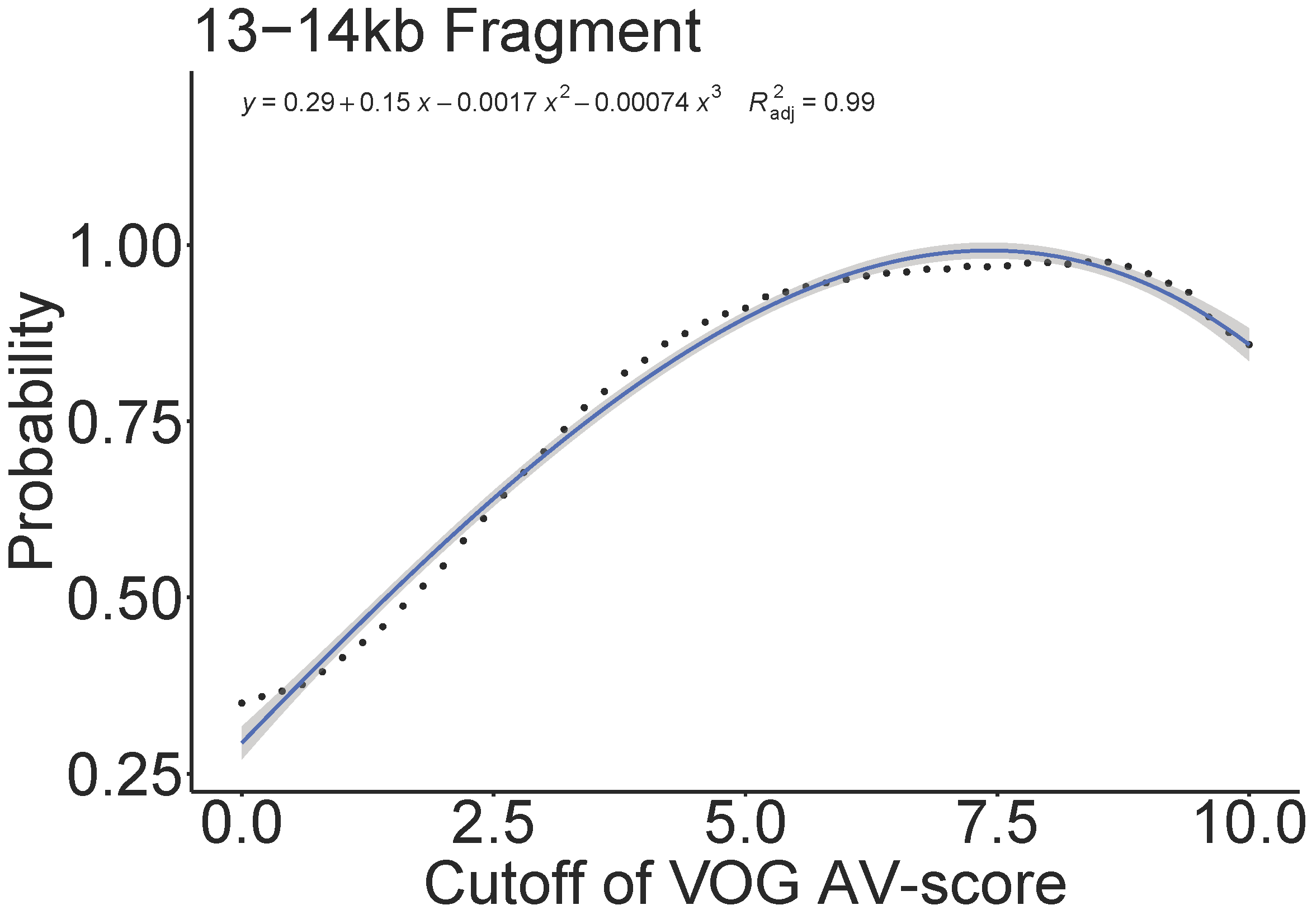

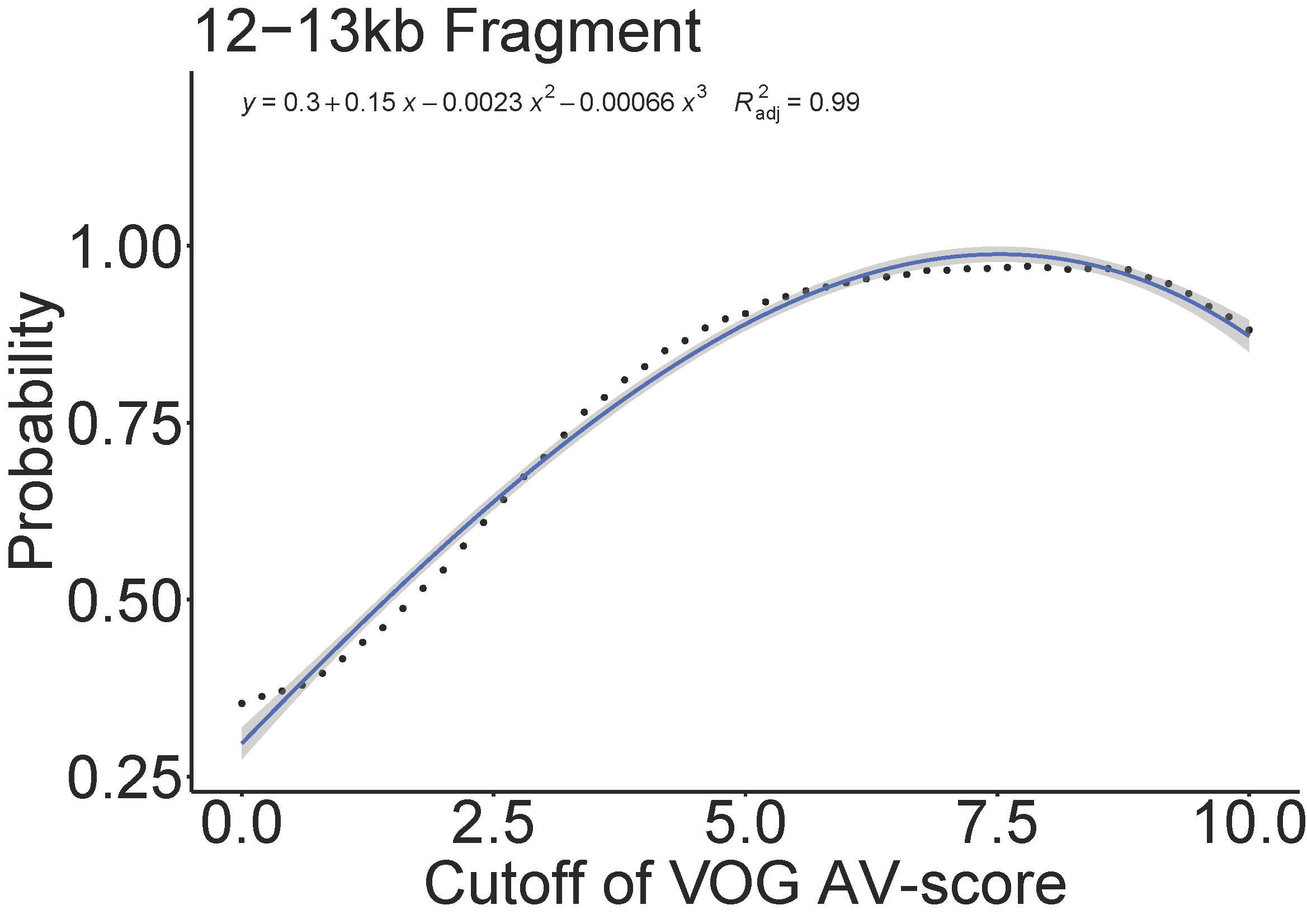

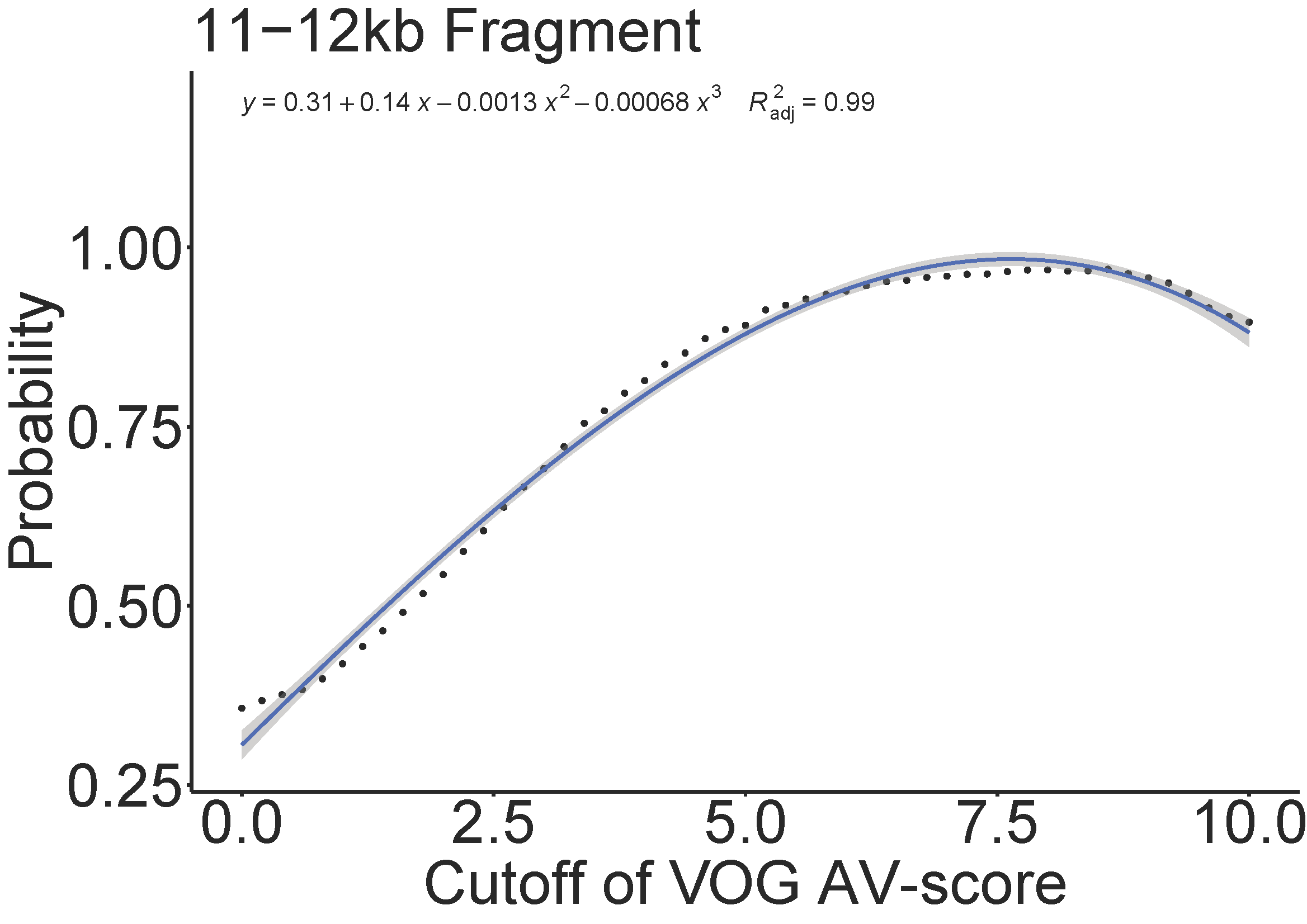

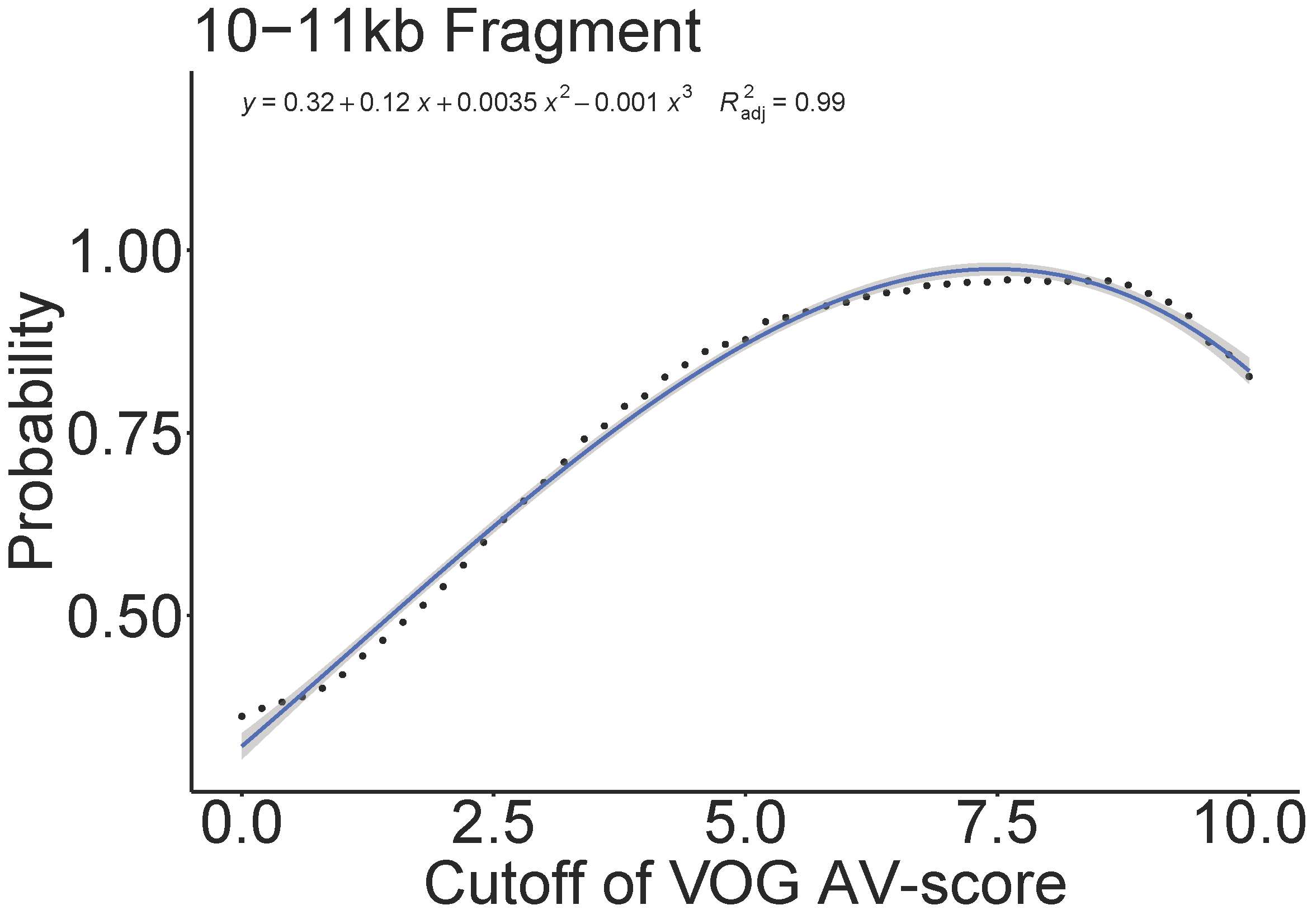

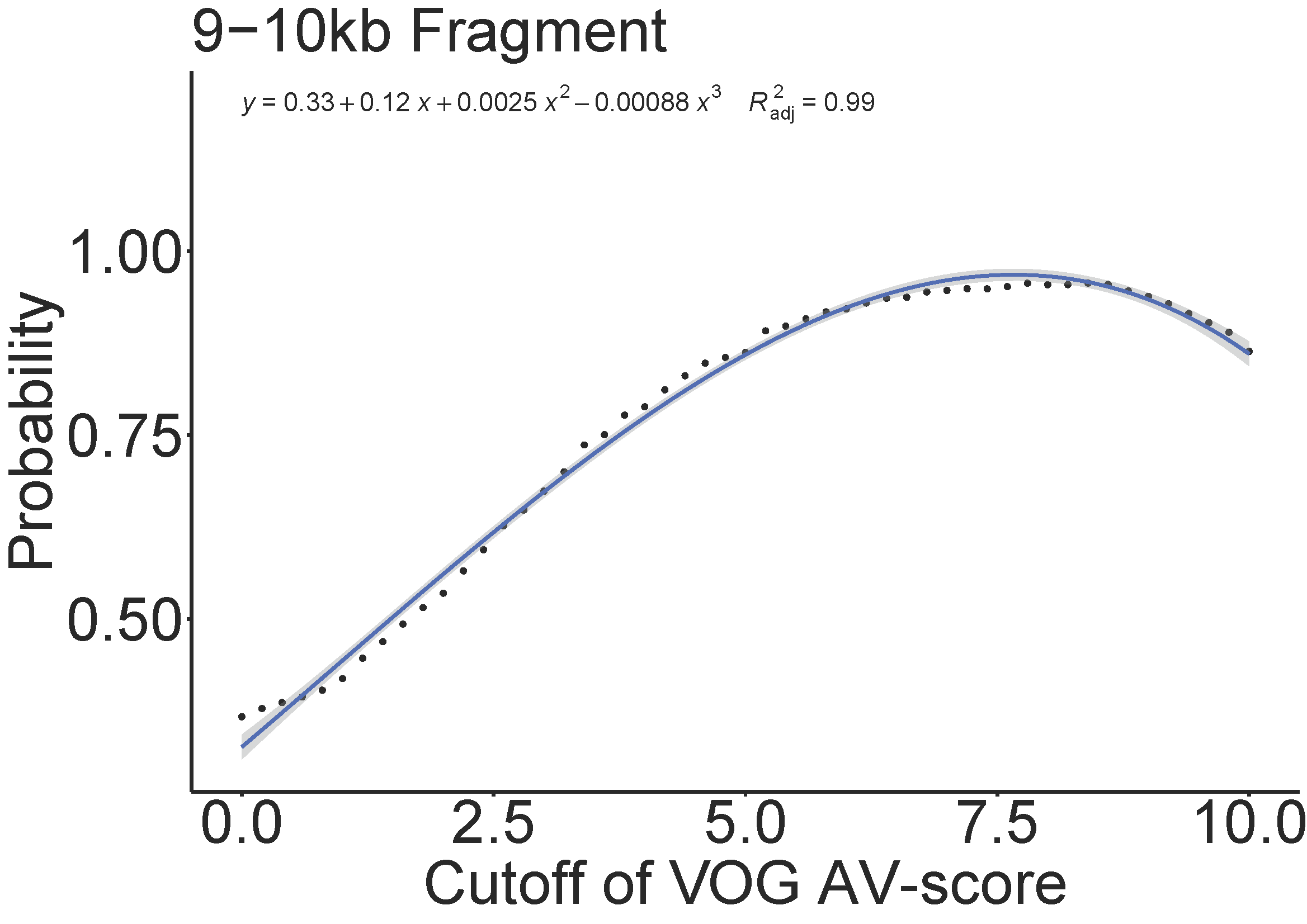

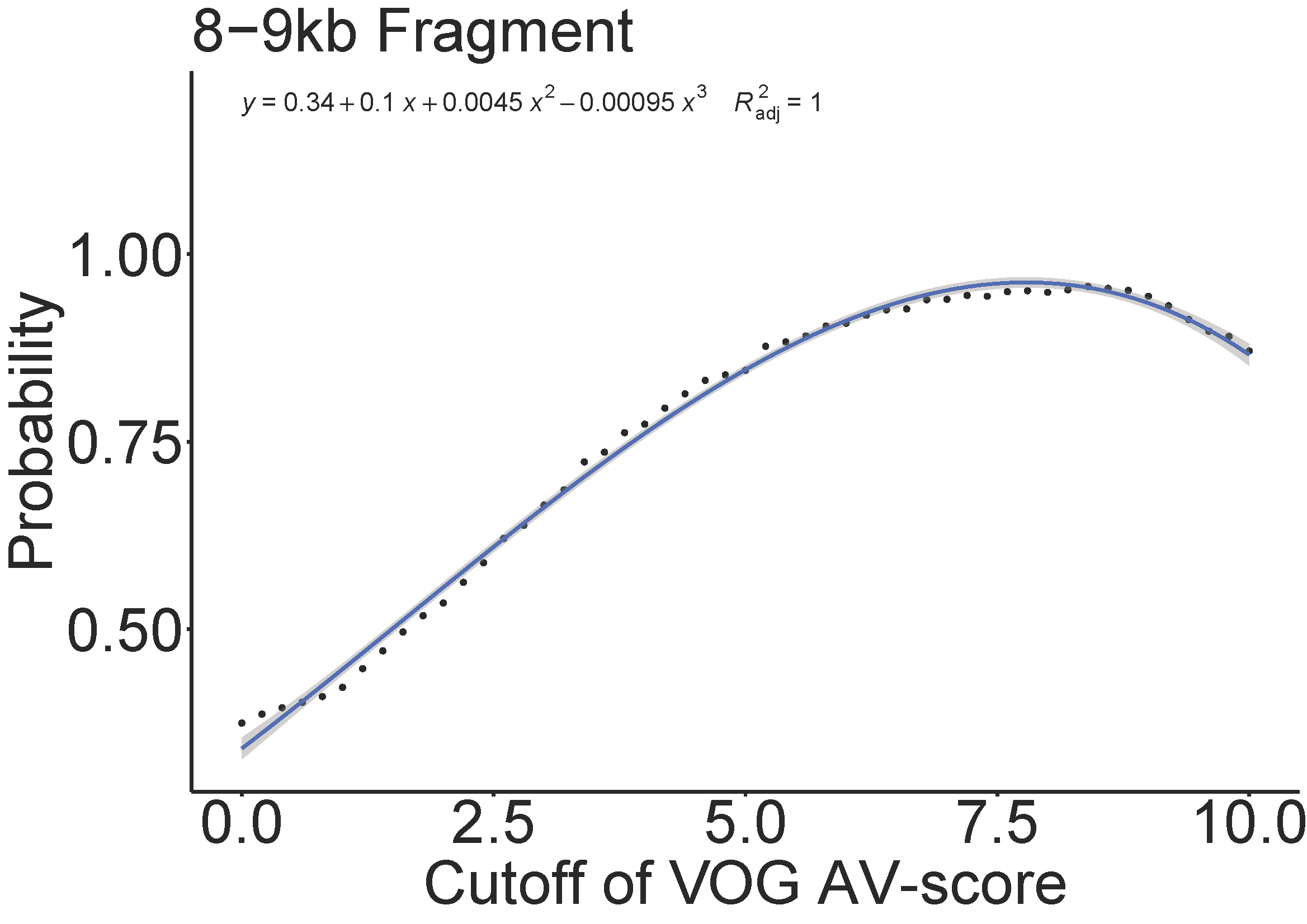

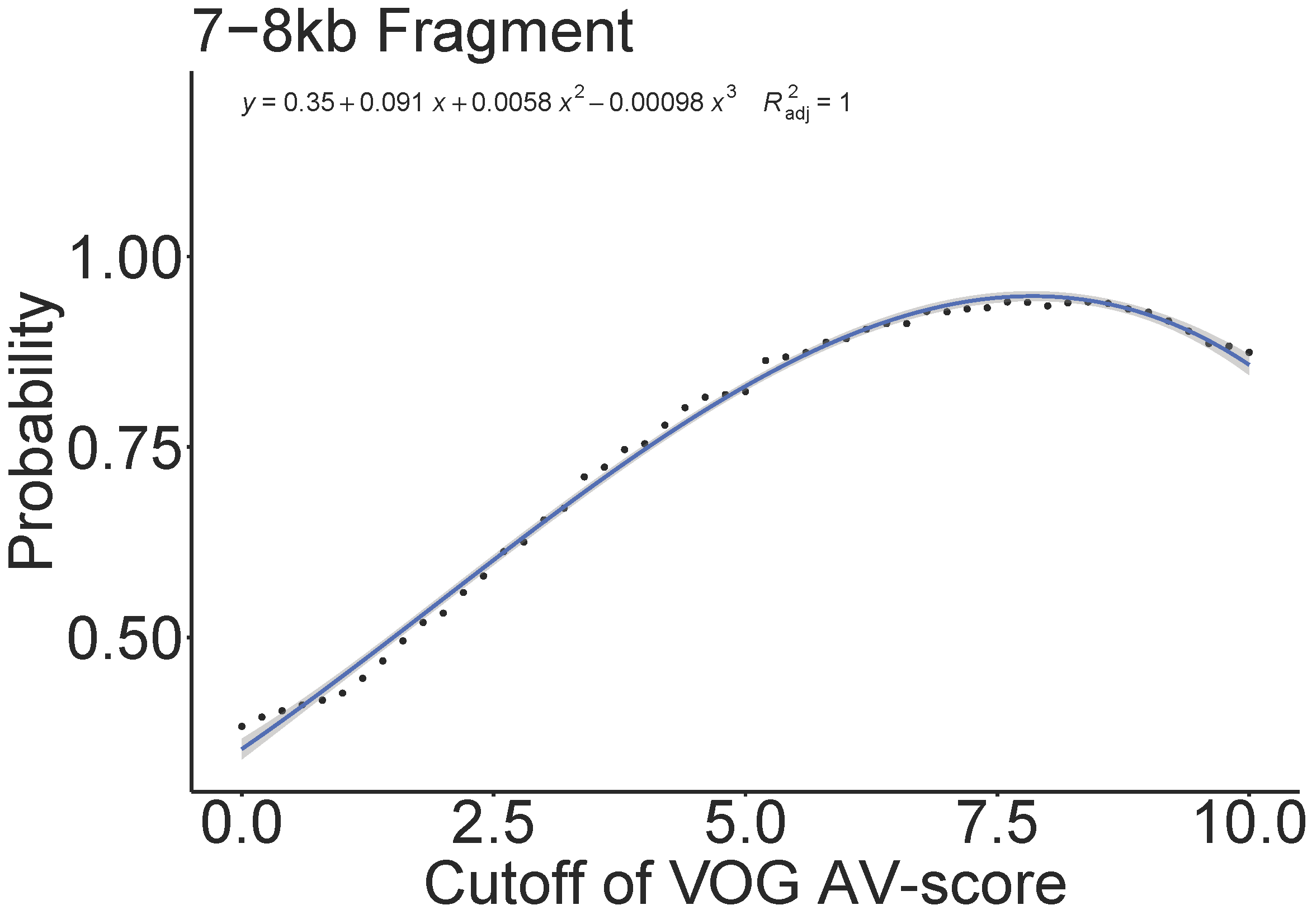

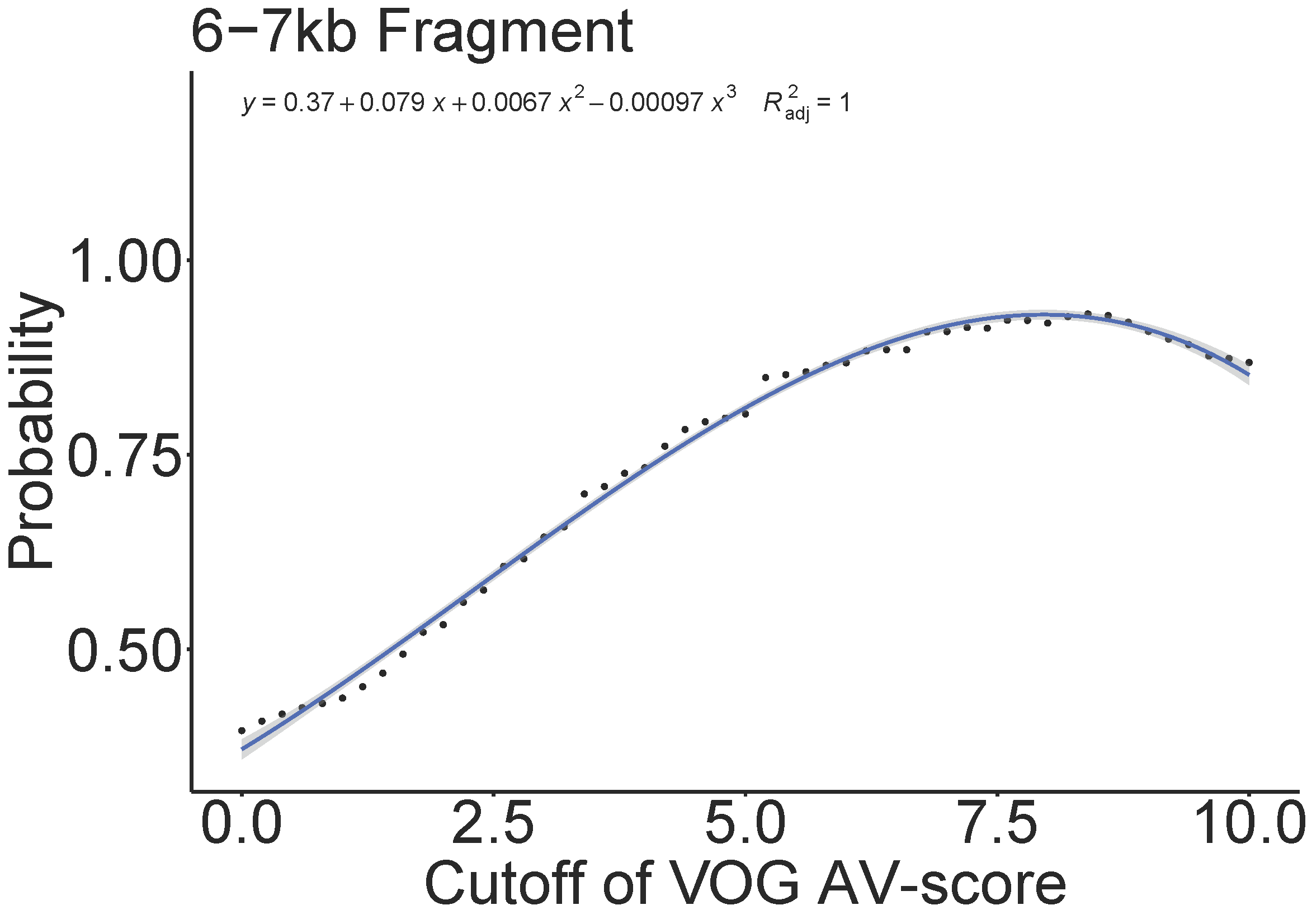

**Fig. S7.** Relationship between the cutoffs of the VOG AV-score and AV_L_-score of whole genomes used in Fig. S2 and S3 and the fraction of viral genomes (here was defined as probability) above certain cutoffs.

a	b

**Fig. S8.** Number of low-, medium-, and high-quality viral sequences identified using AV-scores and geNomad (a). A Venn diagram was used to display the number of sequences shared between the two approaches. (b) For medium- and high-quality sequences, as assessed by CheckV, the overlap between the two approaches (geNomad and AV-score) was illustrated using Venn diagrams, showing the number of shared sequences identified by both methods.

**Fig. S9.** Distribution of AV-scores of Pfam, KEGG, eggNOG, VOG, and eggNOG of the *E. coli* host and its provirus genomes. The asterisks (***) indicate a significance level of *p* < 10^-3^.

c

b

a

d

e

f

**Fig. S10.** Illustration of the generation of the fraction of viral proteins from the comparison between plasmids, chromosomes, and viruses. In Fig. **S10a−e**, red boxes are used to highlight the variation in the ratios of viral, plasmid, and chromosomal proteins. These plots demonstrate the process of calculation of the fraction of viral proteins. The top of each red box remains constant (with the upper edge corresponding to a V_L_-score of 5), while the position of the bottom varies. We observed that as the bottom of the red boxes rises, indicating an increase in V_L_-scores, the fraction of viral proteins within the red box also increases. We tested more than 40 different V_L_-scores, each corresponding to a different position for the bottom of the red boxes. A dot plot was generated to show the relationship between the fraction of viral proteins and these 40+ V_L_-scores (**Fig. S10f**). A clear pattern emerged, showing that the fraction of viral proteins increases as the V_L_-scores rise, indicated by the upward movement of the bottom of the red boxes. A polynomial analysis of the dot plot produced a formula that relates the fraction of viral proteins to the V_L_-scores (**Fig. S10f**). We define the fraction of viral proteins as the probability of a protein being viral, and the V_L_-score corresponding to the bottom of the red boxes as the cutoff for identifying viral-like proteins (the definitions of probability and cutoff are also applicable to AV_L_-score and AV-score). This allows us to predict viral proteins based on V_L_-scores. For **Fig. S10f**, the dots on the dotted line represent the actual values of the fraction of viral proteins, while the blue lines indicate the predicted values.
